# Metabolic modeling of the International Space Station microbiome reveals key microbial interactions

**DOI:** 10.1101/2021.09.03.458819

**Authors:** Rachita K. Kumar, Nitin K. Singh, Sanjaay Balakrishnan, Ceth W. Parker, Karthik Raman, Kasthuri Venkateswaran

## Abstract

**Background:** Recent studies have provided insights into the persistence and succession of microbes aboard the International Space Station (ISS), notably the dominance of *Klebsiella pneumoniae*. However, the interactions between the various microbes aboard the ISS, and how it shapes the microbiome remain to be clearly understood. In this study, we apply a computational approach to predict possible metabolic interactions in the ISS microbiome and shed further light on its organization.

**Results:** Through a combination of a systems-based graph-theoretical approach, and a constraint-based community metabolic modelling approach, we demonstrated several key interactions in the ISS microbiome. These complementary approaches provided insights into the metabolic interactions and dependencies present amongst various microbes in a community, highlighting key interactions and keystone species. Our results showed that the presence of *K. pneumoniae* is beneficial to many other microorganisms it coexists with, notably those from the *Pantoea* genus. Species belonging to the *Enterobacteriaceae* family were often found to be the most beneficial for the survival of other microorganisms in the ISS microbiome. However, *K. pneumoniae* was found to exhibit parasitic and amensalistic interactions with *Aspergillus* and *Penicillium* species, respectively. To prove this metabolic prediction, *K. pneumoniae* and *Aspergillus fumigatus* were co-cultured under normal and simulated microgravity, where *K. pneumoniae* cells showed parasitic characteristics to the fungus. The electron micrography revealed that the presence of *K. pneumoniae* compromised the morphology of fungal conidia and its biofilm biofilm-forming structures.

**Conclusions:** Our study underscores the importance of *K. pneumoniae* in the ISS, and its potential contribution to the survival (mutualism) and eradication (parasitism) of other microbes, including potential pathogens. This integrated modelling approach, combined with experiments, demonstrates immense potential for understanding the organization of other such microbiomes, unravelling key organisms and their interdependencies.

## Background

Microorganisms are ubiquitous and exist in diverse communities around us. They form complex dynamic assemblages in every ecosystem, and their interactions shape the biotic and abiotic environments [1–3]. Beyond natural environments such as the soil or the ocean, microorganisms abound in all human habitats, where they directly impact human health [4]. The human-made International Space Station (ISS) is a unique and controlled system to study the interplay between the human microbiome and the microbiome of their habitats. The ISS is a hermetically sealed closed system; yet it harbors many microorganisms that survive extreme environmental conditions such as microgravity, radiation, and elevated CO_2_ levels [5–8]. Recent studies have demonstrated the unique link between crew member microbiomes and surface microbiomes on the ISS [9]. These microorganisms might have been brought into the ISS via routine payloads and astronauts [9].

Several studies have focused on microbial isolation [10] and molecular microbial community analyses of the ISS microbiome, through experiments performed on vacuum filter debris [11], high-efficiency particle arrestance (HEPA) filters [12, 13], the ISS environmental surfaces [8, 14] and astronaut’s microbiome [9]. A recent report using a shotgun metagenomic sequencing approach on intact cells of the ISS environmental microbiome revealed the succession and persistence of certain microbial populations on the ISS environmental surfaces [15]. The study posited a dominant, viable presence of Biosafety Level – 2 (BSL-2) pathogens such as *Klebsiella pneumoniae, Staphylococcus aureus, Enterococcus faecalis*, and *Salmonella enterica* (Figure S1).

*K. pneumoniae* is well-known for its ability to cause pneumonia and other nosocomial infections and is largely studied for its known resistance to a wide spectrum of antibiotics, such as carbapenems, and its hypervirulence [16–20]. The present study is motivated by the evidence presented about the dominance of *K. pneumoniae* at multiple locations of the ISS, its succession over time [15], and the potential clinical implications it could have on the health of the astronauts inhabiting the ISS.

Metabolic interactions are a key driver in shaping microbial communities [21]. Studying these metabolic interactions can be instrumental in understanding the interplay between various microorganisms in diverse communities [22–24]. Genome-scale metabolic modelling is a powerful tool to study and understand microbial metabolism [25]. These modelling approaches capture in detail, the known metabolic reactions happening in a microorganism, along with the enzymes that catalyze them, representing them in a mathematical form amenable to simulations [26]. Beyond single microorganisms, metabolic modelling can also be extended to study microbial communities; many paradigms have been developed, including those based on graph theory, and constraint-based modelling, as reviewed elsewhere [27, 28].

In this study, we leverage the metagenome datasets that have captured the microbial composition of the ISS surfaces [15] to specifically predict how various species in these communities can influence one another’s metabolism, leading to mutually beneficial interactions and stable microbiomes. In particular, we focus our study on *K. pneumoniae* and its coexisting species, to elucidate the possible underlying metabolic interactions that drive these communities. Through a combination of a systems-based graph-theoretical approach, MetQuest [29, 30], and a constraint-based community modelling approach, SteadyCom [31], we illustrate the central role of metabolic interactions and dependencies in shaping the ISS microbiome.

## Methodology

In this study, a two-pronged computational approach was adopted to investigate and decipher the prevailing microbial interactions in the ISS. In the first graph-based approach [29], bipartite graphs were created to represent the metabolic networks of microorganisms taken individually, as well as when in pairs or larger communities. These networks were used to identify possible metabolic dependencies between different microorganisms. In the second constraint-based approach [31], microbial growth rates were predicted, individually and in communities, to gauge the nature of their interactions, as we describe in the following sections. Figure 1 shows a broad overview of the approach of this study, outlining the steps beginning with the identification of microorganisms coexisting with *K. pneumoniae* to various metabolic network analyses.

**Figure 1.**
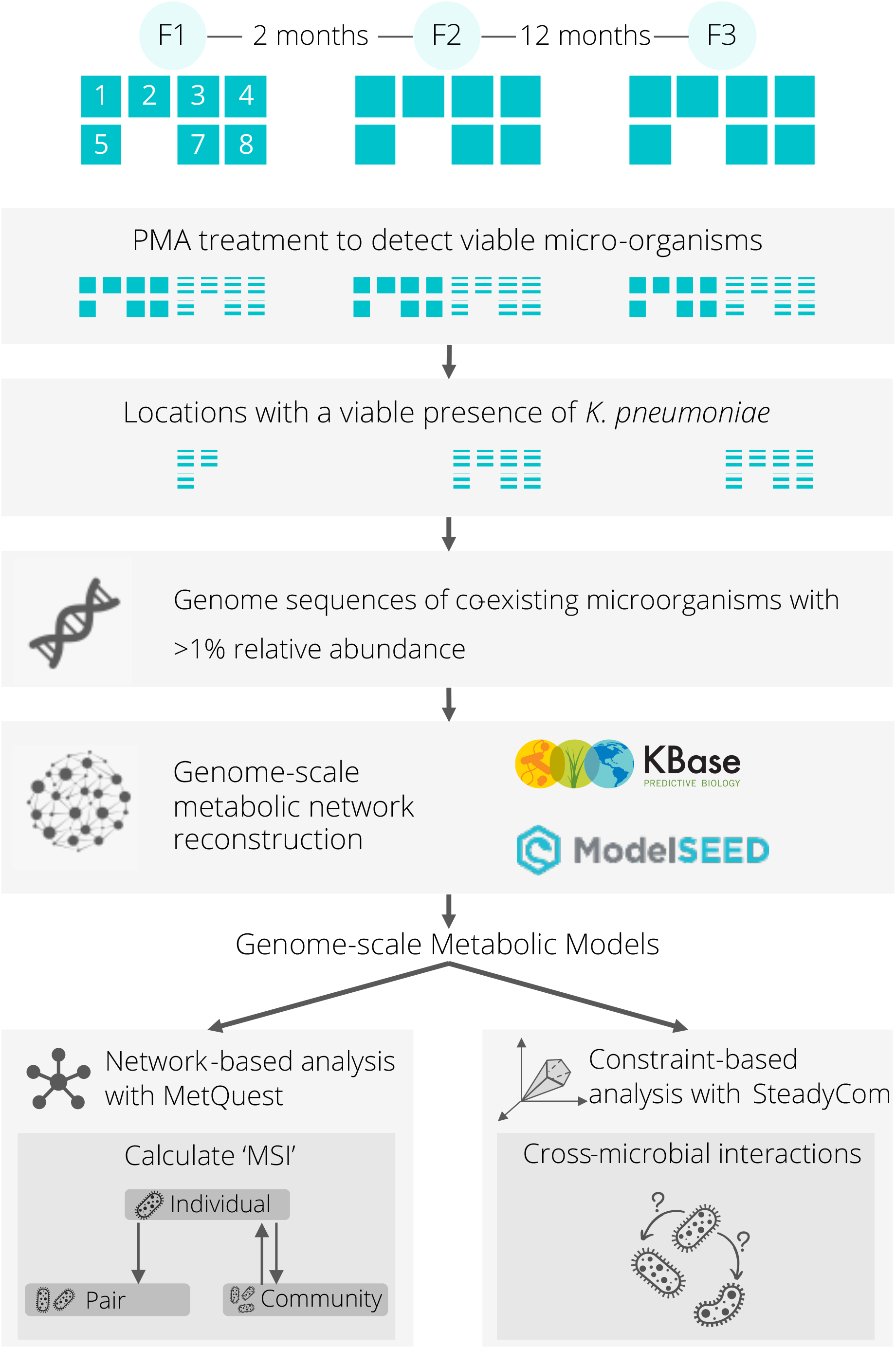
Overview of the analysis pipeline. Beginning with identifying a list of microorganisms that coexist with *K. pneumoniae*, the genome-scale metabolic models were built using KBase and the extent of benefit derived by each individual microorganism in different scenarios was computed using MetQuest and the Metabolic Support Index (MSI) as a metric. SteadyCom, a constraint-based approach, was also employed here to determine the effect of a microorganism on the growth of another, and to further classify the nature of their interactions into their respective types based on the observed change in growth rates.

## Data

Microbial abundances associated with samples taken across three flights at eight ISS locations were previously measured using metagenomic sequencing [15]. Among eight locations, only seven locations yielded measurable metagenomics data. In the present study, the data were further pruned such that only shotgun metagenome sequences associated with propidium monoazide (PMA)-treated samples that recorded a presence of *K. pneumoniae* were retained. The PMA-treatment removed naked DNA and dead cells, thus retaining sequences associated with viable and intact cells only [32]. Further, microorganisms that were found to coexist with *K. pneumoniae* with >1% relative abundance at the location only were used in this study.

For this study, the available whole-genome sequences (WGS) of microorganisms from the ISS, which were compatible with KBase [33] along with reference genomes for the relevant microorganisms taken from the NCBI database, were collected. The RefSeq and GenBank Accession identification numbers, along with the underlying reasoning for the choice of these sequences, is provided in Supplementary Table S1. Although the relative abundance of Pantoea sp. PSNIH2 was >1% in some of the locations considered in this study, as its reference genome sequence was removed from the NCBI database, this microorganism therefore had to be excluded. Metabolic models were constructed for the 52 microorganisms using ModelSEED [34]. The detailed process is elaborated in Supplementary Methods M1. The details of the reconstructions, templates and includes information about reaction counts are specified in Supplementary Table S2.

### Predicting metabolic dependencies in the community

To investigate the microbial interactions underlying the chosen microbial communities of the ISS, MetQuest [29], a graph-theoretic algorithm was first employed. Through a novel dynamic-programming based enumeration, MetQuest assembles reactions into pathways of a specified size producing a given target from a specified set of source molecules (‘seed’). Using this algorithm, the present study presents a top-down view of metabolic dependencies in these communities, beginning with predicting the most supportive families, then the most dependent genera, and finally the beneficial and dependent microorganisms by looking at pairs of potentially interacting microorganisms. In each scenario, the extent of benefit derived was calculated using a measure known as the Metabolic Support Index (MSI) [30].

The composition of the seed metabolites was chosen so as to incorporate the co-factors and coenzymes that would be present in the environment and required by a living cell, and capture the minimal nutrient content offered by the ISS. The minimal medium of each metabolic model was identified by minimizing the components required to achieve a minimum growth rate of 0.1 h^-1^, using built-in functions in COBRApy [35]. This approach is adopted on the premise that each microorganism is capable of individual growth. For each location at each flight, the minimal medium sets corresponding to each constituent microorganism were pooled together to form a consolidated medium. Additionally, a base set of co-factors and coenzymes were added to each of these sets. The same media were used for both the graph-based and constraint-based analyses. Supplementary Table S3 contains the list of all such metabolites.

### Higher-order Interactions

The enhancement of metabolic capabilities rendered by a microorganism by virtue of it thriving in a community was derived from the metabolic support index, here referred to as the community support index (CSI). CSI was calculated as the fraction of reactions relieved in the rest of the community over the reactions “stuck” in the same in the presence of the microorganism. For a community *X* and an inhabitant microorganism *A*, the effect of *A* on the rest of the community *Ã* = (*X* − *A*) on *A* is given by:

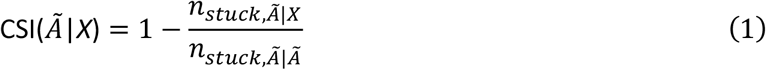

Here, *n*_*stuck,Ã*| *Ã*_ refers to the number of stuck reactions in the community *X* without microorganism *A* for the given set of seed metabolites. A reaction is considered stuck when it does not have the necessary precursors for the reaction to happen. In other words, these are reactions that cannot occur under the present conditions, without extraneous help in terms of other metabolites from other microorganism(s), or the environment itself. Similarly, *n*_*stuck,Ã*| *X*_ refers to the number of reactions in *Ã* that remain stuck, even in the presence of *A*. We consider only internal reactions (as against transport or extracellular reactions) while computing these numbers.

For each of the sites considered in this study, bipartite graphs were first constructed (*n* = 129) for communities devoid of one constituent microorganism, and their respective stuck reactions were determined in the appropriate environment (i.e., the seed metabolites described above, and listed in Supplementary Table S3). This approach provided insight into the intrinsic metabolic capabilities of each community devoid of that microorganism. Following this, bipartite graphs were constructed, taking all 11 inhabitant microorganisms. The stuck reactions were determined for each community to gather the effect of the interactions by calculating the CSI. A CSI of unity indicates that *A* completely supports *Ã* and relieves all its stuck reactions, whereas a value of zero indicates that *A* has no effect on *Ã*.

Microorganisms could potentially be metabolically similar, and therefore redundant to the community, offering little or no metabolic support individually. Therefore, microbial interactions were also studied by grouping them by their respective families (*n* = 7), at each location. The metabolic support provided by groups of microorganisms (e.g., a family) can be readily computed by replacing *A* in *X* − *A* with the set of microorganisms comprising the family in Eq. (1).

It is also possible to estimate the community benefit, in terms of enhancement of metabolic capabilities, an individual microorganism might receive by virtue of it thriving in a community. Here, the community support index was calculated as the fraction of reactions relieved in the microorganism when in a community assemblage over the reactions stuck in the individual. For a community *X* and an inhabitant microorganism *A*, the effect of the rest of the community *Ã* on *A* is given by:

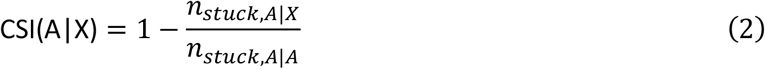

The CSI thus captures the “benefit” a microorganism or a group of microorganisms receives from another microorganism or group, by virtue of its stuck reactions being relieved via metabolic exchanges with its coexisting microorganism(s).

### Pairwise Interactions

For a pair of microorganisms, *A* and *B*, MSI is calculated as:

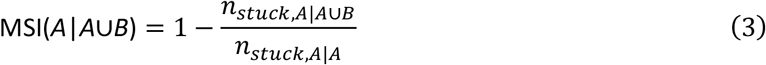

Similar to the approach used above, individual bipartite graphs were constructed (*n* = 52), and the stuck reactions were first determined for every individual microorganism in the dataset, which provided insight into the intrinsic metabolic capabilities of each individual microorganism. Following this, for every pair of coexisting microorganisms that may potentially interact, bipartite graphs were constructed for the two-membered community (*n* = 426), and the stuck reactions were determined for each constituent member of each community, in a flight-location-specific approach (*n* = 761), to derive the effect of its partner by virtue of being in this community, through the calculation of the metabolic support index.

Building further on these interactions, as captured by the metabolic support indices between various microbial pairs, microbial association networks were constructed and visualized using Cytoscape [36]. These networks capture the extent to which a microorganism is able to relieve the stuck reactions of another through metabolic exchanges.

For clarity, all community support indices and metabolic support indices have been represented as percentages.

### Determining the nature of interactions

Microbial interactions have major relevance in building and shaping the community. Based on the effects a microorganism has on another, the interactions can be categorized into six types, namely amensalism, commensalism, competition, neutralism, mutualism, and parasitism [37]. The present study reports a constraint-based community modelling study using the algorithm, SteadyCom [31].

SteadyCom was employed to determine the biomass rates of all microorganisms that are a part of a community under steady state. The seed metabolites described earlier, in the network-based approach were used here as the medium (see Supplementary Table S3). The lower bounds of the exchange reactions for the uptake of these metabolites, were constrained to −10 mmol/gDW-h [38]. Joint models were then created for all pairs of microorganisms that can potentially interact by virtue of being at the location at that point in time. These joint models were optimized using SteadyCom [31] to find the individual biomass rates when existing as a community.

The growth rate observed for each microorganism in community was compared with that observed for the individual microorganism. A 10% or higher increase or decrease in growth rate, when in a community, was taken to be a significant effect [39]. The interactions were accordingly classified into the types listed above. The equation defined below was used to determine the effect of microorganism *B* on *A*, in a two-membered community constituting *A* and *B*.

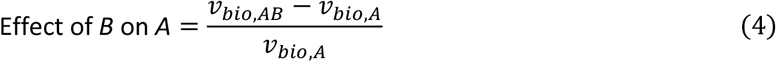

Depending on the value (positive [+], negative [-], or neutral [0]) obtained, the interaction was classified as mutualism (the effect of B on A, and the effect of A on B, both being positive, i.e. +, +), neutralism (effects on each other being not significant, denoted 0,0), competition (a negative effect on each other, denoted by -,-), amensalism (-,0 and 0,-), commensalism (0,+ and +,0), and parasitism (-,-).

### *K. pneumoniae* and *A. fumigatus* interactions

Overnight grown cultures of *K. pneumoniae* (10^6^ cells per 100 µL) and purified conidial spores of *A. fumigatus* (10^6^ per 100 µL) were either mixed or grown alone in 10 mL sterile potato extract (0.4%; w/v) dextrose (2%; w/v) liquid medium and grown under normal and simulated microgravity conditions using

High Aspect Ratio Vessel (HARV; Synthecon inc., Houston, TX) culturing units [40]. The autoclavable HARVs provided oxygenation to the culture media via their large-diameter gas permeable membrane. The HARVs with microbes were incubated at 30°C; 150 rpm, for 48 hours. HARVs were run in parallel in both horizontal rotation (normal gravity control) and vertical rotation (simulated microgravity) orientation during growth. After growth, the bacterial and fungal cells were harvested, then fixed by incubating the cells in 2.5% glutaraldehyde in 0.1M Sodium Cacodylate (Sigma) buffer at 4°C for 1 hour for scanning electron microscopy (SEM) study. Samples were then washed in 0.1M Sodium Cacodylate buffer three times. Fixed cells were then dehydrated in isopropyl alcohol (IPA, Sigma), using a stepped series of increasing concentrations (50%, 70%, 80%, 90%, 95 to 100%). Each step consisted of a 10 min incubation at 4°C, followed by 3 x replacements with 100% IPA, and finally stored at 4°C. Samples were critically point dried in an Automegasamdri 915B critical point dryer (Tousimis, Rockville, MD). Samples were attached in SEM stubs with carbon tape (Ted Pella Inc., Redding, CA), followed by carbon coating with a Leica EM ACE600 Carbon Evaporator (Leica, Wetzlar, Germany) to a thickness of ∼12nm. SEM analysis was performed with an FEI Quanta 200F (Thermo Fisher, Waltham, MA).

## Results

The key results of this study are four-fold. First, among the families considered, species belonging to the *Enterobacteriaceae* family were often found to be the most beneficial, among the ISS microbiome. Secondly, *Pantoea* species were predicted to be extensively dependent on their coexisting microorganisms. Third, through metabolic network analysis of microorganisms taken pairwise, *K. pneumoniae* was found to be beneficial to many of its coexisting microorganisms, especially to those species belonging to the *Pantoea*. Fourth, metabolic interactions in the community broadly fell under the categories of amensalism and parasitism. The parasitic interaction under normal and simulated microgravity between *K. pneumoniae* and *A. fumigatus* was experimentally checked.

### Data acquisition and genome-scale metabolic network reconstruction

According to the data published by Singh et al. 2018 [15], among the PMA-treated samples, reads of *K. pneumoniae* were detected in a total of eleven sites – locations #1, #2 and #5 in Flight 1, location #5 in Flight 2 and locations #1, #2, #3, #4, #5, and #7 in Flight 3 (Figure 2). The relative abundances of microorganisms were calculated at the respective sites, and upon further pruning of this data to include solely those with >1% abundance, a total of 50 different strains of microorganisms that coexist with *K. pneumoniae* at varied locations and time points were chosen for further study.

**Figure 2.**
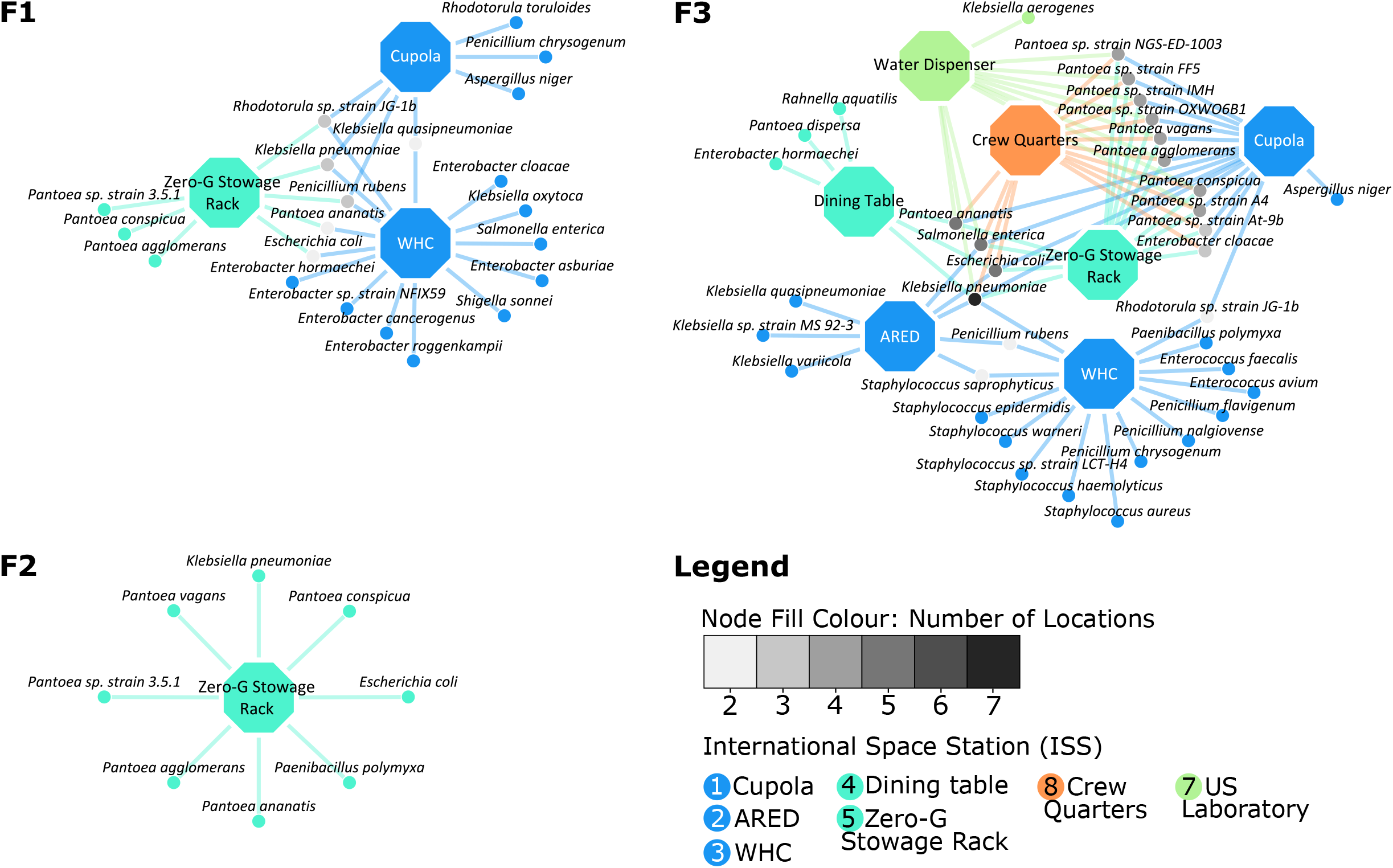
Locations with a recorded viable presence of *K. pneumoniae*. The figure describes the locations of the ISS at which *K. pneumoniae* was detected from the respective PMA treated samples, and also features its coexisting microorganisms that inhabit the location at >1% relative abundance. The networks have been drawn on Cytoscape for all three time points of the study by Singh et al. 2018: Flight 1 (F1), Flight 2 (F2) and Flight 3 (F3). The octagonal location nodes have been color coded according to the ISS Node they belong to. Locations #1, #2, and #3 belong to Node 3, locations #4 and #5 to Node 1, location #6 (not shown here) to PMM and location #7 to the US Laboratory. The microbial nodes have been color-coded such that those existing at more than one location with *K. pneumoniae* are shown in grey, implying that a microorganism found in a large number of locations (such as *K. pneumoniae* in F3) will be filled with a darker shade of grey.

Analysis of the diversity of these microorganisms at each of these locations over time (Figure 2) reveals the following trends. In Flight 1, among the three locations that were colonized by *K. pneumoniae*, there was an observed dominance of fungi in location #1, *Enterobacteriaceae* in location #3, and *Erwiniaceae* in location #5. In Flight 2, *K. pneumoniae* was documented to be present only in location #5, coexisting with extant and dominant *Erwiniaceae*, a member from the *Enterobacteriaceae* family and another from the *Paenibacilleae* family. In Flight 3 however, *K. pneumoniae* spread across the ISS, inhabiting nearly all sampled locations. Additionally, many more members of *Erwiniaceae*, specifically the *Pantoea* genus, emerged into view at a majority of these locations, as shown in Figure 2. Further, with the exception of *A. niger* and *Rhodoturula* sp. strain JG-1b, all fungi considered in the dataset appear to be concentrated in location #3. Location #3 is also dominated by a large number of *Staphylococcaceae*. The list of microorganisms, along with their relative abundances at each location in each flight, is provided in Supplementary Table S4.

### *Enterobacteriaceae* are pivotal contributors to community metabolism

Through a “leave-one-out” approach, microorganisms were knocked out one at a time from the community to estimate the extent of benefit provided by that single microorganism to the remaining coexisting microorganisms. From the analysis, it was evident that some microorganisms do have an influence on the metabolic capabilities on the rest of the community, albeit low in most cases. Notable among these, were *P. rubens* at location #3 during Flight 3 (CSI = 1.15%), and at locations #2 (CSI = 0.40%) and #5 (CSI = 0.36%) during Flight 1, and *K. oxytoca* at location #2 in Flight 1 (CSI = 0.49%) (Supplementary Table S5 and Supplementary Figure S2).

These low values of CSI are somewhat not surprising, given the possible high metabolic overlap between members within a community, thereby rendering individual organisms less important to a community. To examine this, we grouped organisms based on their phylogenetic affiliation, and removed these groups, one at a time, from the community, to study the importance of a given family to the community.

Overall, with respect to the locations considered in this study, the *Enterobacteriaceae* family comprising species of the *Klebsiella, Escherichia, Salmonella, Shigella*, and *Enterobacter* genera, was often found to be the most beneficial of all clusters, with the highest CSIs in six out of a total eleven communities under study (Figure 3 and Supplementary Table S6), five of which are during Flight 3. With regards to the remaining five communities, *Trichocomaceae* have the highest CSIs at locations #2 and #5 during Flight 1, *Erwiniaceae* have the highest CSIs at location #1 in Flight 3 and at location #5 during Flight 2, and *Paenibacillaceae* has the highest CSI at location #2 in Flight 3.

**Figure 3.**
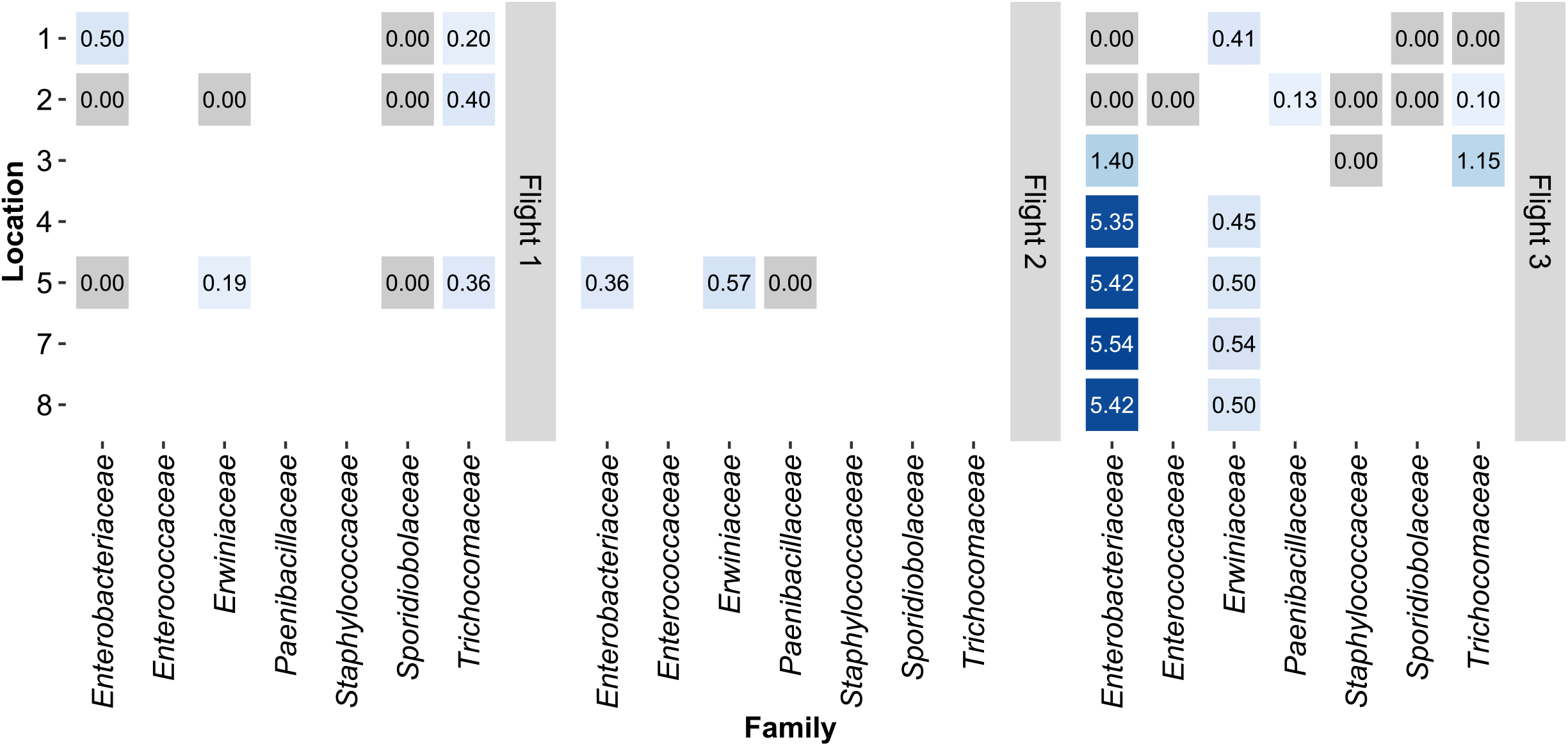
Metabolic support provided by each family to the remaining community. The values on the tiles denote the Community Support Index (CSI) calculated in the presence of the family. The X-axis denotes the families present in the dataset, and the Y-axis denotes the respective locations. The heatmap has been facetted to indicate the three timepoints, i.e. flights.

### *Pantoea* species thrive in the support of the microbiome

The CSI of an individual microorganism in the presence of its coexisting microorganisms, was calculated to gain insight into the combined effect of the community on an individual. The CSI values of *Pantoea* were found to be among the highest (Supplementary Table S7 and **Error! Reference source not found**.), indicating that these species are benefitted to a comparatively greater extent from the remaining members of the community. Across all locations and flights in the dataset, the CSIs of these species ranged between 0 and 6.52%, wherein the values at the lower end were predominantly observed in those *Pantoea* species found at location #1 in Flight 3. *P. vagans* was found to exhibit the highest dependency amongst all the *Pantoea*, at location #5 at Flight 2. (Figure 4).

**Figure 4.**
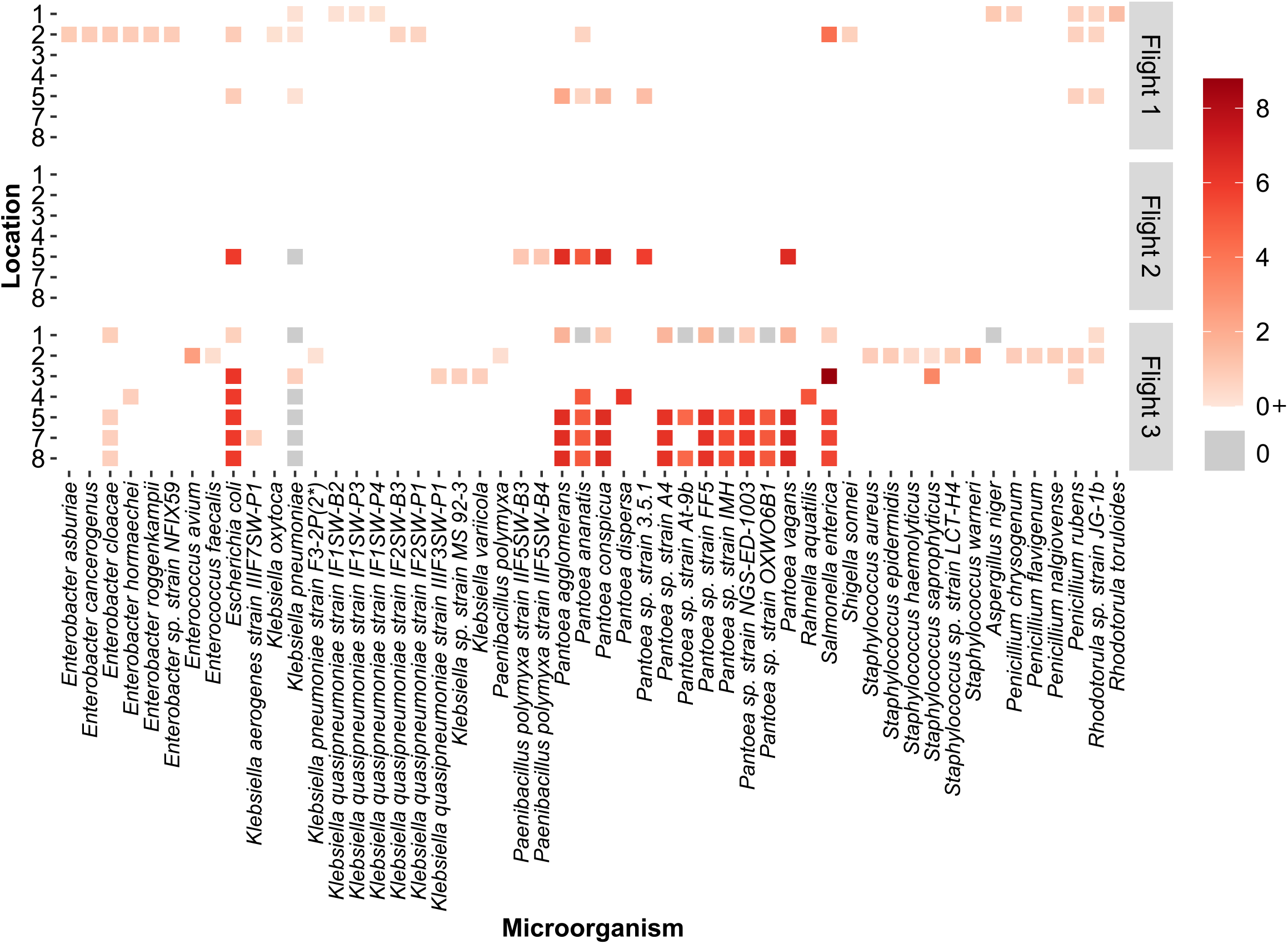
Extent of metabolic benefit derived by an individual microorganism from its coexisting microorganisms. The heatmap depicts the range of metabolic support indices that indicate the metabolic support rendered to an individual by virtue of it being in that location. On the X-axis is the list of microorganisms in consideration, and on the Y-axis is the flight number and the concerned location number. A darker red tile indicates the microorganism is benefitted to a greater extent in that flight and location.

With the exception of *E. coli* and *S. enterica*, the CSI values of other *Enterobacteriaceae* fell in the range of 0 - 0.91%, suggesting an overall lower dependence of members of the *Enterobacteriaceae* on their coexisting microorganisms (**Error! Reference source not found**.). *K. pneumoniae* showed no dependency (CSI = 0%) in six out of the 11 locations that they were found. In addition, *S. enterica* exhibited the highest community dependence amongst all microorganisms in consideration, with a CSI = 8.79% at location #3 during Flight 3, and along with *E. coli*, showed high CSI values (Figure 4).

Analysis across the three flights at location #5 reveals that four out of five organisms present at all instances are benefitted to a lesser extent at Flight 1 compared to Flights 2 and 3 (**Error! Reference source not found**. and Supplementary Table S7).

### *K. pneumoniae* are beneficial to its coexisting microorganisms

In every community, microorganisms exchange metabolites, influencing one another’s growth and survival. We performed a pairwise analysis of a total of 761 flight-location-specific communities to identify how every pair of microorganisms interacts with one another, at a given location, during a given flight, and through various metabolic exchanges support each other.

In communities with *K. pneumoniae* and a member outside of the *Enterobacteriaceae*, the MSI values in the presence of *K. pneumoniae* were found to fall in the range of 0 – 6.46%. Most *Pantoea* species were found to be highly dependent on *K. pneumoniae*, with MSIs at the higher end (4.41 – 6.46%) of the spectrum (Supplementary Table S8). With the exception of *E. coli* and *S. enterica*, the MSI values of all other *Enterobacteriaceae* members were found to be 0% in the presence of *K. pneumoniae* (Supplementary Figures S3 and S4). Out of 118 pairs, *K. pneumoniae* was found to have a non-zero MSI value in only nine such pairs. In these nine pairs, the other interacting member belonged to a species of the *Penicillium* genus.

Since *K. pneumoniae* was present across the three flights only at location #5, the corresponding microbial association networks were constructed to gain insights into microbial interaction patterns over time (Figure 5). In Flight 1, fungi such as *P. rubens* and *Rhodotorula* sp. JG-1b are prevalent and shown to decline in abundance over time (Flight 2 and Flight 3). On the contrary, members of the *Panteoa* genus were observed to dominate on this location over time, with many more *Pantoea* species surfacing. In agreement with earlier results discussed, many species of the *Enterobacteriaceae* family shown here such as *K. pneumoniae* and *E. cloacae*, were beneficial to members of the *Erwiniacea* family to a great extent (Figure 5).

**Figure 5.**
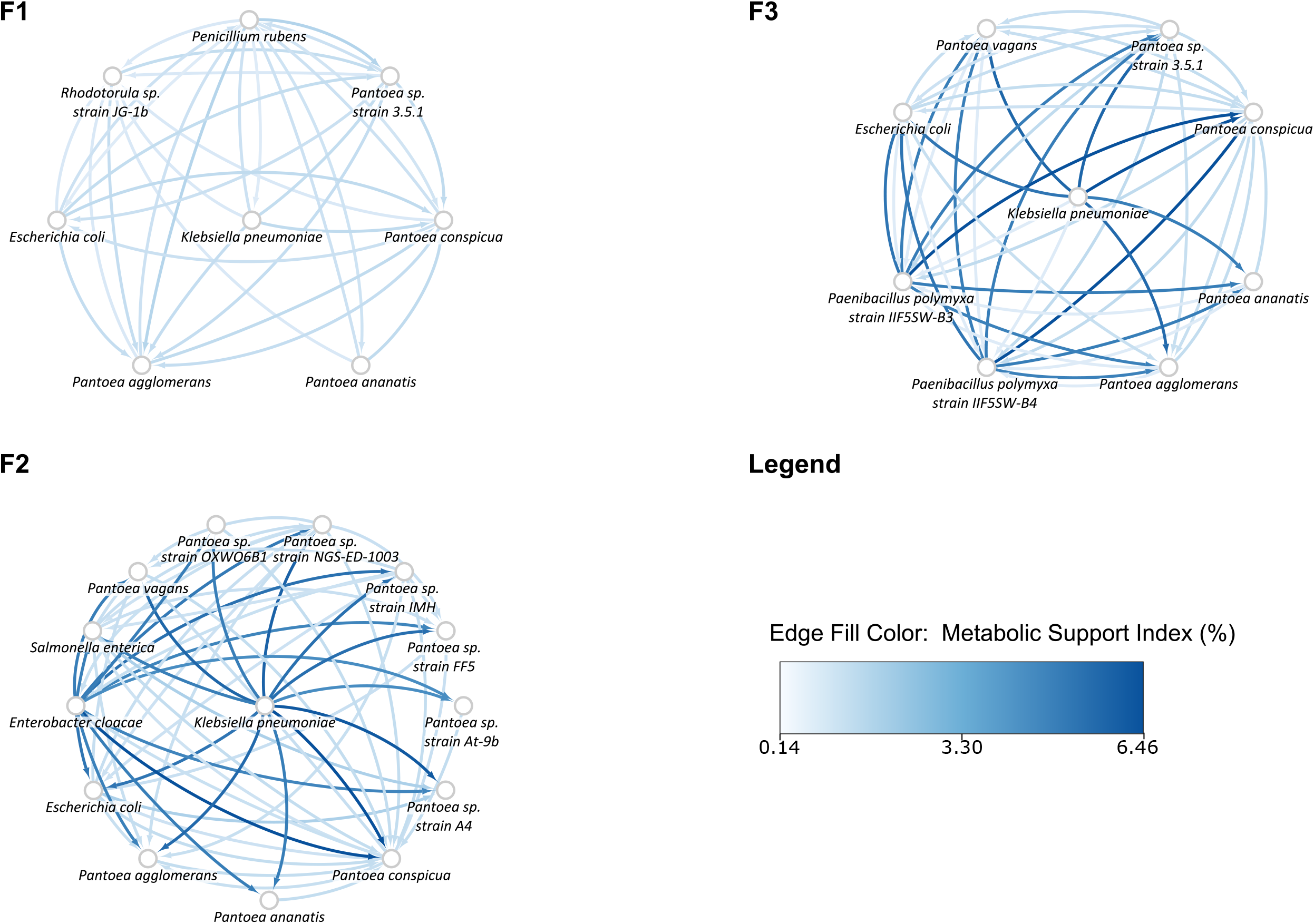
Microbial association networks for pairs of microorganisms inhabiting location 5, across the three flights. Cytoscape was used to construct and visualize these networks across three flights: Flight 1 (F1), Flight 2 (F2) and Flight 3 (F3). The nodes are labelled with microorganisms that inhabit the site at that respective time point. The directed edges are directed from the metabolically supportive microorganism to the metabolically dependent microorganism. The color of these directed edges are mapped to the Metabolic Support Indices (MSI), represented as percentages, such that the gradient from light blue to dark blue represents an increasing MSI.

### Constraint-based modelling suggests the domination of amensalistic and parasitic interactions

The graph-theoretic approaches described earlier do not explicitly account for the growth of microorganisms in the presence of one another or their rates, although a higher MSI does point towards a possible higher biomass in the community [30]. Complementary to the above approaches, constraint-based modelling approaches such as SteadyCom [31] can shed light on the ability of microorganisms to grow in a steady-state. Based on these predicted growth rates, it is also possible to classify the nature of the interaction between various pairs of microorganisms [37].

A total of 761 flight-location specific pairs of microorganisms were studied. Among these, 458 pairs exhibited amensalism and 166 were predicted to be parasitism. Of the remaining, 48 were commensal, 36 were competitive, 34 were mutualistic, and 19 were neutral.

Looking closely into the nature of interactions of *K. pneumoniae* and its coexisting microorganisms, in 118 such pairs, 72 were predicted to be amensal, 14 commensal, 13 involved in competition, 10 were parasitic interactions, 7 were mutual, and 2 were neutral. In communities with the *Enterobacter* species, *K. pneumoniae* was involved in amensalistic interactions as observed across all such flight-location pairs. While in the interactions with *E. cancerogenus* and *Enterobacter* sp. strain NFIX59, the growth rate of *K. pneumoniae* was predicted to decrease, in the interactions with all other *Enterobacter* species, there was no significant effect on the predicted growth rate of *K. pneumoniae*.

In eight out of nine flight-location-specific communities of *K. pneumoniae* with *E. coli, E. coli* was predicted to have no significant change in growth rate while that of *K. pneumoniae* was predicted to decrease, suggesting an amensalistic interaction. Further, in Flight 3, at location #3, *E. coli* was predicted to be parasitic towards *K. pneumoniae*. In the interactions with *S. enterica*, in four out of the six locations, the growth rate of *K. pneumoniae* was decreased in its presence, whereas that of *S. enterica* was increased, resulting in a parasitic interaction. This is in accordance with the previously observed high MSI values at locations #3, #5, #7 and #8 during Flight 3. In the other cases during Flight 1 at location #2 and Flight 3 at location #1, the interaction was predicted to be amensalistic, wherein the growth rate of *K. pneumoniae* was reduced.

With other *Klebsiella* species, the interactions were predicted to be predominantly amensalistic, with the growth rate of *K. pneumoniae* reduced, except in the interaction with *Klebsiella* sp. strain MS 93-3, where the reverse was observed. *K. pneumoniae* was predicted to be parasitic towards K. aerogenes strain IIIF7SW-P1.

The interactions with *Pantoea* species, such as *P. agglomerans, P. conspicua, Pantoea* sp. strain 3.5.1, *Pantoea* sp. strain A4, *Pantoea* sp. strain At-9b, *Pantoea* sp. strain FF5, *Pantoea* sp. strain NGS-ED-1003, *Pantoea* sp. strain OXWO6B1 and *P. vagans*, were amensalistic with a decreased growth rate of the *Pantoea* species, in all cases except in Flight 3 at location #1, where the interactions were predicted to be competitive. Interactions with *P. ananatis* however were found to be commensal, with an observed increased growth rate. *P. dispersa* and *Pantoea* sp. strain IMH were found to compete with *K. pneumoniae* at their coexisting locations (Figure 6 and Supplementary Table S9).

**Figure 6.**
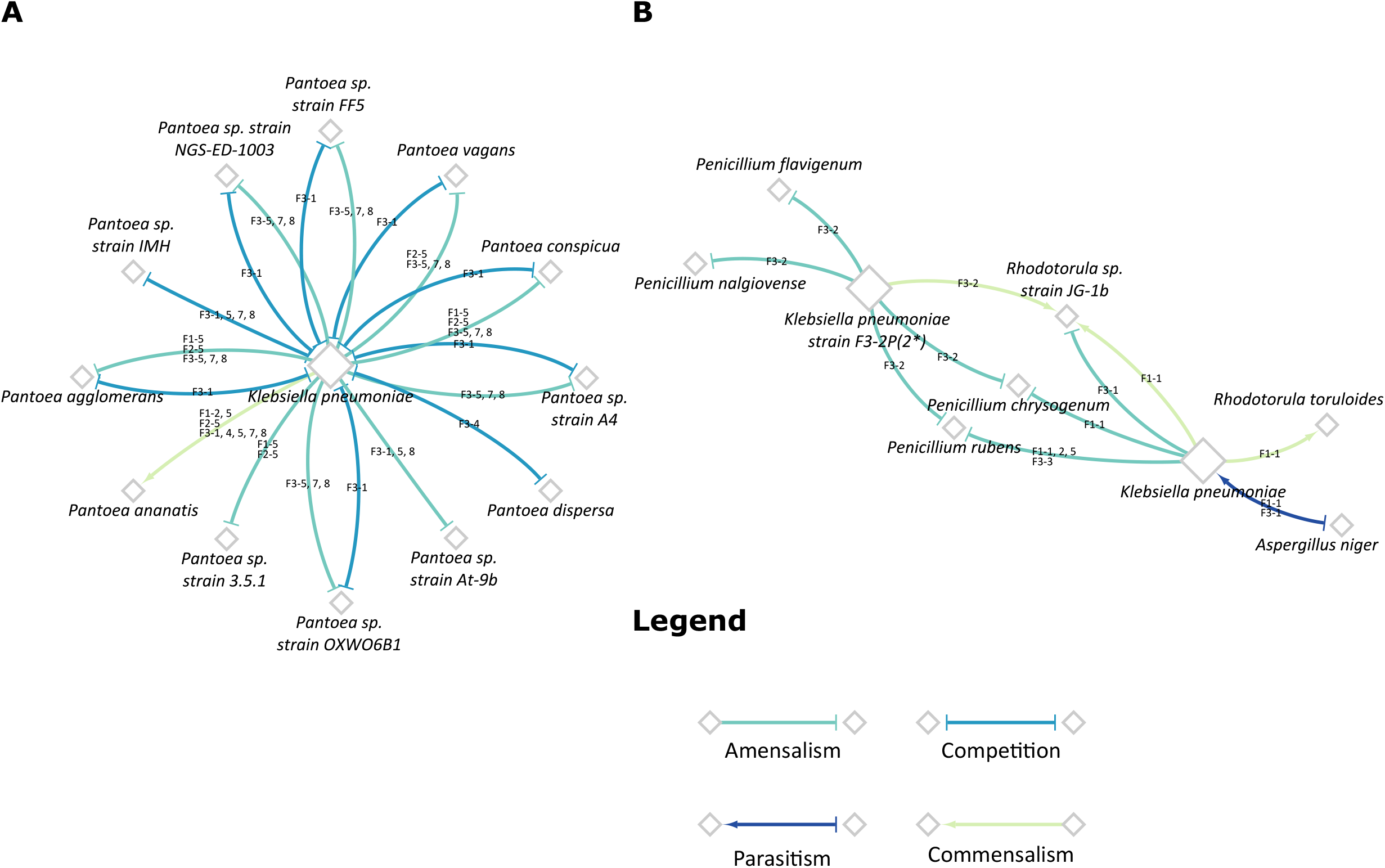
Key interactions with *Klebsiella pneumoniae*. The network diagrams depict the nature of interactions pertaining to those of A) *Klebsiella pneumoniae* with coexisting *Pantoea* species B) *Klebsiella pneumoniae* with the coexisting fungi. The color and arrowheads of the edge indicate the nature of the observed interaction, and the edge labels correspond to the flight-location in which that type of interaction was observed. The nodes of *K. pneumoniae* and *K. pneumoniae* strain F3-2P(2*) have been increased in size solely for the purpose of clarity.

*Staphylococcus* sp. in Flight 3 at location #2 were found to be mutualistic with *K. pneumoniae* strain F3-2P(2*), with the exception of *S. saprophyticus* that was observed to be involved in commensalism. During Flight 3, location #3 however, this interaction of *S. saprophyticus* with *K. pneumoniae* was predicted as mutualistic (Supplementary Table S9).

The interactions of *K. pneumoniae* with *A. niger* were observed to be parasitic, with an observed decreased growth rate of *A. niger*, and an increased growth rate of *K. pneumoniae*. With *Penicillium* species, *K. pneumoniae* had an amensalistic effect on them, resulting in their decreased growth rates. The interactions with *Rhodotorula* sp. strain JG-1b were found to depend on the flight and location in which they coexisted. During Flight 1, at location #1, an increased growth rate was observed in *Rhodotorula* sp. strain JG-1b, whereas there was no significant effect on that of *K. pneumoniae*, thereby suggesting a commensal interaction. In locations #2 and #5, however, a neutral interaction was predicted, with no significant change in growth rates of either species. At Flight 3, *K. pneumoniae* was observed to be amensalistic towards *Rhodotorula* sp. strain JG-1b at location #1, and commensal at location #2. *K. pneumoniae* increased the growth rate of *R. toruloides* at Flight 1 at location #1 (Figure 6 and Supplementary Table S9). Details regarding the nature of interactions of other microorganisms in the community have been delineated in Supplementary Table S9.

### Detrimental effect of *K. pneumoniae* on *Aspergillus fumigatus*

The scanning electron micrographs confirmed the parasitic effect of *K. pneumoniae* over *A. fumigatus* (Figure 7). Under normal gravity conditions, *A. fumigatus* was found to be healthy where conidiophore holding vesicle with healthy metulae and phialides harboring conidia were noticed (Figure 7A), whereas simulated microgravity affected the phialides structure (Figure 7C) and subsequently conidial spores were not clearly visible. When *K. pneumoniae* was co-cultured with the fungus, under normal gravity conditions, the bacterial cells (artificially colored in red) were shown to destroy the conidial spores and also partially degraded metulae and phialides (Figure 7B). Furthermore, when *K. pneumoniae* cells were grown with the fungus and incubated in simulated microgravity conditions, bacterial cells destroyed the fungal structure where only vesicle was seen along with the spores and phialides, additionally the metulae were disformed and the hyphae (Figure 7D) appeared to be disfigured compared to those seen in Figure 7A or 7B. This demonstrates that *K. pneumoniae* in combination with simulated microgravity negatively influenced the growth of the fungi. Both these bacterial and fungal strains were isolated from ISS and detrimental effect of *K. pneumoniae* as predicted by the metabolic model during this study was experimentally demonstrated, confirming the degeneration of *A. fumigatus* morphological architecture. In addition to these morphological modifications, *K. pneumoniae* also reduced the biofilm forming capability of the fungus when grown together (data not shown).

**Figure 7:**
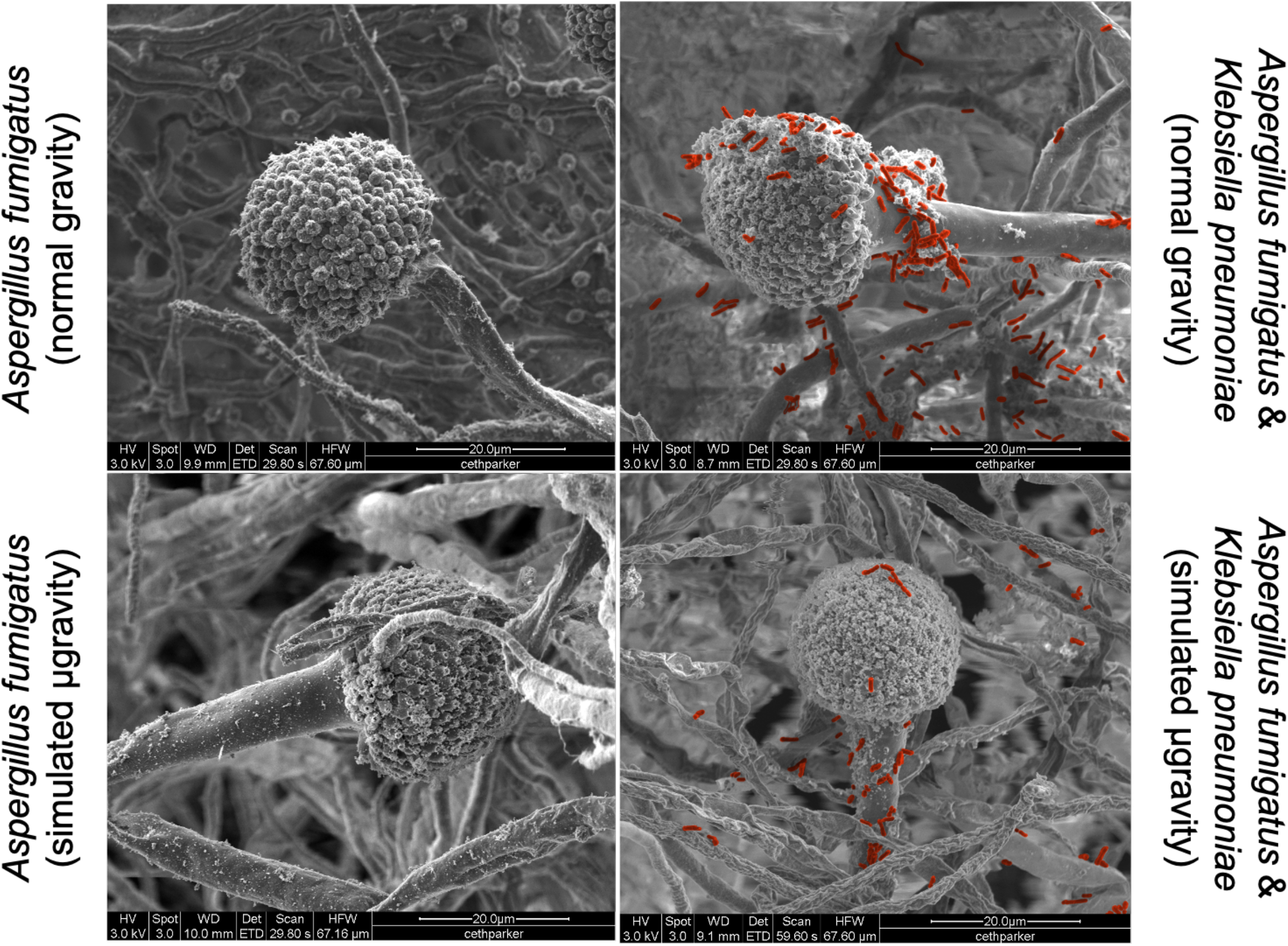
Antagonistic characteristics of *K. pneumoniae* when co-cultured with *A. fumigatus* under simulated microgravity. (A) *A. fumigatus* grown under normal gravity, (B) Both bacteria and fungus grown under normal gravity, (C) *A. fumigatus* grown under simulated gravity, and (D) Both bacteria and fungus grown under simulated gravity. *K. pneumoniae* cells were artificially colored to show their presence on and around fungus culture.

## Discussion

In this study, we used a metabolic perspective to analyze the ISS microbiome with a focus on *K. pneumoniae*, a BSL-2 human pathogen, and its coexisting microorganisms. In our two-pronged approach, we first used a graph-theoretical approach to unravel the metabolic interdependencies of the microbiome, and we further complemented this with constraint-based analyses, to identify the consequences of microbial interactions on growth. We begin by looking at higher-order interactions in the community and further delve deeper and look at pairwise interactions, that have been previously shown to be major drivers of community dynamics [41]. We traverse three taxonomic levels – family, genus, and species, and provide insights into the keystone species of the community through metabolic network analyses.

Even though <7% of the metagenomic reads of ISS environmental surfaces constituted BSL-2 microorganisms, the dominant and persistent human microbial pathogens were *Pantoea, Klebsiella, Staphylococcus, Erwinia*, and *Penicillium* [15]. Among the ISS surfaces, Zero-G Stowage Rack had more *Pantoea* reads compared to other locations of all flights. When metagenomic reads of all BSL-2 microorganisms were compiled, *K. pneumoniae* was found to be persistent and dominant in Zero-G Stowage Rack of all three flights (Figure 2). In general, *K. pneumoniae* reads were more and dominant in all seven locations sampled in Flight 3. Unlike in Flight 3, *K. pneumoniae* reads were retrieved sparingly in Flight 1 (Cupola and WHC) and Flight 2 (Zero-G Stowage Rack). Among fungi, reads associated with *Penicillium* species were dominant and persistent in Flight 1 and Flight 2, but *K. pneumoniae* succeeded its dominance in Flight 3 [15]. Unlike *K. pneumoniae* dominance (e.g. Zero-G Stowage Rack), fungi did not persist in certain locations or flights examined but were sporadically present in other locations as well. However, reads associated with yeast (*Rhodotorula* species) were high only in a certain location (WHC). The cleaning reagent used in ISS consists of benzalkonium chloride, which might be able to eradicate the fungal population; thus fungal succession was not observed, unlike bacterial members which were persistent in the ISS locations examined [42]. It is also documented that *K. pneumoniae* cells were resistant to the quaternary ammonium compound concentration used as cleaning agents in ISS [43].

The *in silico* metabolic model results from our analyses were primarily four-fold. On applying graph-theoretical algorithms and surveying the microbiome at multiple levels, we predicted that among the families in the dataset, members of the *Enterobacteriaceae* family were perhaps the most beneficial to other microorganisms in the community, whereas in most cases, members of the families such as *Erwiniaceae, Staphylococcaceae*, and *Sporidiobolaceae* (yeast) offered little or no additional metabolic benefit to the community. It has been documented that some *Rhodotorula* species produced silver nanoparticles containing antimicrobial properties against a wide variety of Gram-positive and negative microorganisms [44]. The members of the family *Sporidiobolaceae* (*Rhodotorula*, pink yeast) were isolated from the ISS locations [8], their antagonistic and parasitic behavior predicted during this study should be tested against opportunistic microbial pathogens to aid in the development of appropriate countermeasures. Similarly, the amensalism predicted towards *Erwiniaceae* and mutualism of *Staphylococcaceae* members with *K. pneumoniae* need further study utilizing the ISS strains.

At the genus level, we observed that *Pantoea* genus typically derives the most benefit in the community. On looking at the pairwise interactions, we observed that amongst other *Enterobacteriaceae, K. pneumoniae* provides metabolic support to many of its coexisting species, notably those from the *Pantoea* genus. The members of the genus *Pantoea* were primarily considered as plant pathogens, but subsequently, they have been isolated from many aquatic and terrestrial environments, including ISS [8], as well as in association with insects, animals, and humans [45–49]. During this study [8], the metagenomic sequences of the opportunistic human pathogens associated with *Pantoea* genus were *P. agglomerans, P. conspicua, P. brenneri, P. ananatis*, and *P. dispersa*, whereas the plant pathogenic *Pantoea* species were not observed. The competitive metabolomic properties predicted during this study by *K. pneumoniae* with *Pantoea* members (Figure 6A) might be due to the assimilation of similar compounds for sustaining their growth. Detailed phenotypic metabolic profiles of these microbes are needed to confirm the competition and amensalism predicted in this study.

Finally, from the constraint-based analyses, many amensal and parasitic interactions were noticed. This is corroborated by earlier findings that *K. pneumoniae* can inhibit fungal spore germination and hyphal growth, as well as biofilm formation of *Aspergillus* species [50]. Recent reports documented that *K. pneumoniae* shifted to a pathogenic state, potentially leading to septic infections when cooccurred with *A. fumigatus* in immunocompromised individuals [50]. In addition, this study shows that *K. pneumoniae* cells prevented conidial spore germination and hyphal development of the fungi. It has been further shown that *K. pneumoniae* bacterial cells induced the fungal cell wall stress response mechanisms and suppressed the filamentous growth of fungi [50]. The simple *in silico* metabolic model of this study predicted the antagonistic (parasitic) metabolic interaction between *K. pneumoniae* and *A. niger* (Figure 6B), which further enabled to validate parasitism *in vivo*, using the strains isolated from the ISS.

Our study does have some limitations. First, the metabolic models used in the study are automated reconstructions, which despite gap-filling, could potentially contain gaps and blocked reactions and need to be curated [51]. Nevertheless, such automated reconstructions are being widely used in many studies as they serve as useful predictive tools and representations of the metabolism of microorganisms [52]. Secondly, wherever it is applicable, genomes generated from the cultured ISS microorganisms were utilized. Conversely, when strains were not isolated from the ISS environment, and thus genomes were not available, the reference genomes of the corresponding type strains were used. However, reference genomes provide a useful perspective of an organism’s metabolic capabilities and have also been used in many other studies that incorporate metabolic models [53, 54]. The graph-based approach provides a somewhat static snapshot of the metabolic interactions happening between the organisms. As reported for *Candida albicans*, in addition to the genome-scale metabolic model, refining the model using phenotypic microarray and other wet-lab confirmation are needed [55]. Yet, it is highly useful and offers valuable insights into the potential microbial interactions in the community [30, 56], and well complements the constraint-based methods. Importantly, this study offers a first glimpse into the metabolic interactions of the ISS microbiome, upon which several hypotheses can be formulated for future experimental design.

In summary, our analyses show the key role played by *Klebsiella* and other *Enterobacteriaceae* in mediating the metabolic interactions taking place between microorganisms in the ISS. Metabolic modeling, through a combination of graph-theoretic approaches and steady-state constraint-based modelling, paints a more comprehensive picture of possible microbial interactions, which are as yet inscrutable to this extent by experimental approaches. Our results point towards key dependencies of microorganisms in various locations on the ISS and can pave the way for possible interventions that may rely on targeted disinfection of surfaces aboard the ISS. Our approach also underscores the importance of complementary modelling approaches in dissecting a fairly complex microbiome, and understanding various possible interactions. Our methodology is fairly generic and can be readily extended to predict microbial interactions in other interesting milieu and generate testable hypotheses for wet lab experiments, as we have demonstrated in this study with *Klebsiella and Aspergillus* species.

## List of abbreviations

ISS: International Space Station
BSL – 2: Biosafety Level 2
CSI: Community Support Index
MSI: Metabolic Support Index

## Declaration

## Acknowledgements

RKK acknowledges the post-baccalaureate fellowship from RBCDSAI. SB acknowledges support from the Ministry of Human Resource and Development, Government of India. Part of the research described in this publication was carried out at the Jet Propulsion Laboratory, California Institute of Technology, under a contract with NASA. © 2021 California Institute of Technology. Government sponsorship acknowledged.

## Funding

This research was funded by a 2012 Space Biology NNH12ZTT001N grant nos. 19-12829-26 under Task Order NNN13D111T award to KV, which also funded post-doctoral fellowship for NKS and CEP. KR acknowledges support from the Science and Engineering Board (SERB) MATRICS Grant MTR/2020/000490, IIT Madras, Centre for Integrative Biology and Systems mEdicine (IBSE) and Robert Bosch Center for Data Science and Artificial Intelligence (RBC-DSAI).

## Availability of data and materials

The metagenomic sequence data generated from this study can be found under NCBI Short Read Archive (SRA) under the bio-project number PRJNA438545 and GeneLab dataset GLDS-69 (https://genelab-data.ndc.nasa.gov/genelab/accession/GLDS-69/).

## Authors’ contributions

KV and KR were involved in early organization, study design, and planning of the project, and providing direct feedback to all authors throughout the project and during write-up of the manuscript. RKK and SB wrote the manuscript. RKK generated all figures in the manuscript, processed the data and constructed metabolic models, and performed the MSI-based analyses. SB performed the constraint-based analysis and determined the nature of interactions. NKS was involved in study design, helped interpret and write the manuscript, generated metagenome data and bioinformatic analysis. CWP carried out the microscopy and confirmed the parasitic aspect of *Klebsiella* and *Aspergillus* species. All authors reviewed, read, and approved the final manuscript.

## Ethics approval and consent to participate

No human subjects were analyzed and only environmental samples were collected.

## Consent for publication

All authors participated in this study and given their consent for publishing the results.

## Competing interests

The authors declare that they have no competing interests.

## Supplementary Method Legends

**Supplementary Methods M1:** Protocols involved in the genome-scale metabolic reconstruction

## Supplementary Figure Legends

**Figure S1 BSL-2 pathogens**. The word cloud shows the dominant and persistent BSL-2 pathogens at each location in each flight.

**Figure S2 Extent of metabolic benefit conferred by an individual microorganism to its coexisting microorganisms**. The heatmap depicts the range of community support indices that indicate the metabolic support rendered by an individual microorganism to its coexisting microorganisms, by virtue of it being in that location. On the X-axis is the list of microorganisms in consideration, and on the Y-axis is the flight number and the concerned location number. A darker blue tile indicates the microorganism is highly beneficial.

**Figure S3 Microbial association networks depicting the metabolic dependencies of *E. coli* and its coexisting microorganisms on each other, during Flight 3**. Cytoscape was used to construct and visualize these networks across all locations during Flight 3 (F3). The nodes are labelled with microorganisms that co-inhabit that location with *E. coli*. The directed edges are directed from the metabolically supportive microorganism to the metabolically dependent microorganism. The color of these directed edges are mapped to the Metabolic Support Indices (MSI), represented as percentages, such that the gradient from light blue to dark blue represents an increasing MSI.

**Figure S4 Microbial association networks depicting the metabolic dependencies of *S. enterica* and its coexisting microorganisms on each other, during Flight 3**. Cytoscape was used to construct and visualize these networks across all locations during Flight 3 (F3). The nodes are labelled with microorganisms that co-inhabit that location with *S. enterica*. The directed edges are directed from the metabolically supportive microorganism to the metabolically dependent microorganism. The color of these directed edges are mapped to the Metabolic Support Indices (MSI), represented as percentages, such that the gradient from light blue to dark blue represents an increasing MSI.

## Supplementary Tables

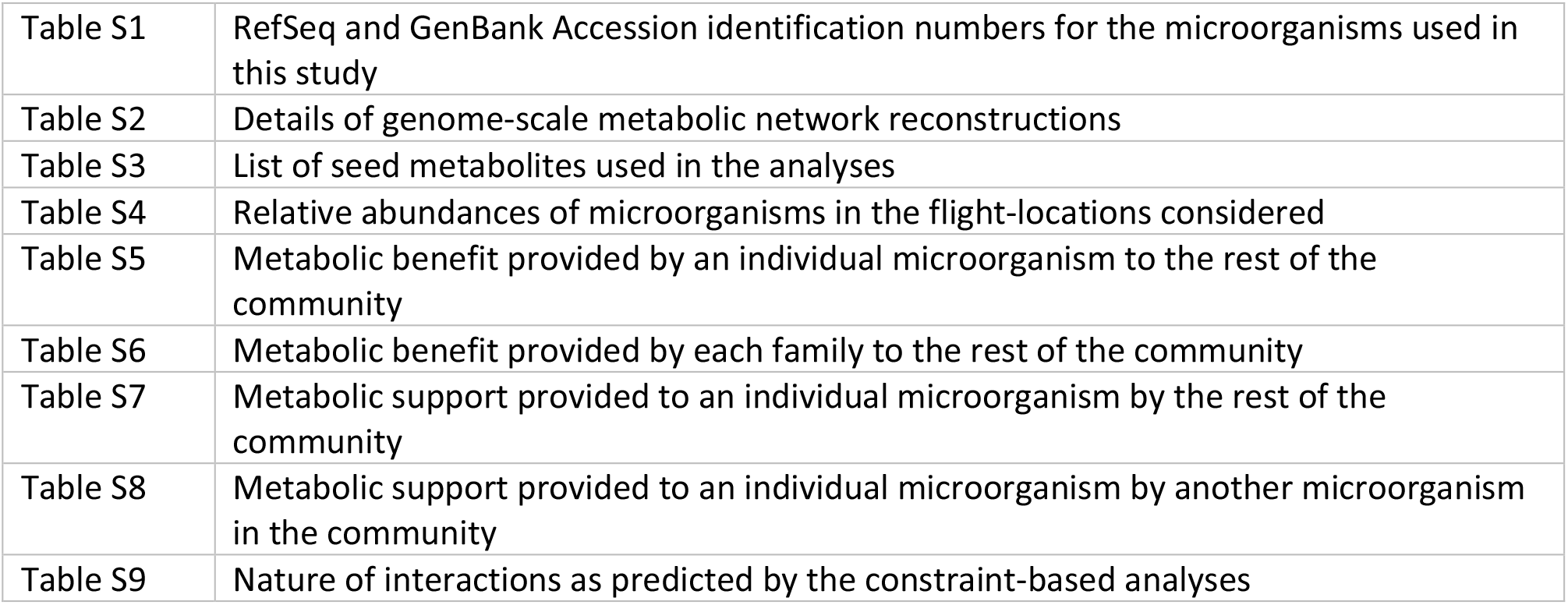

## Notes

### Competing Interest Statement

The authors have declared no competing interest.

