## Supplementary Material for "Metabolic modeling of the International Space Station microbiome reveals key microbial interactions"

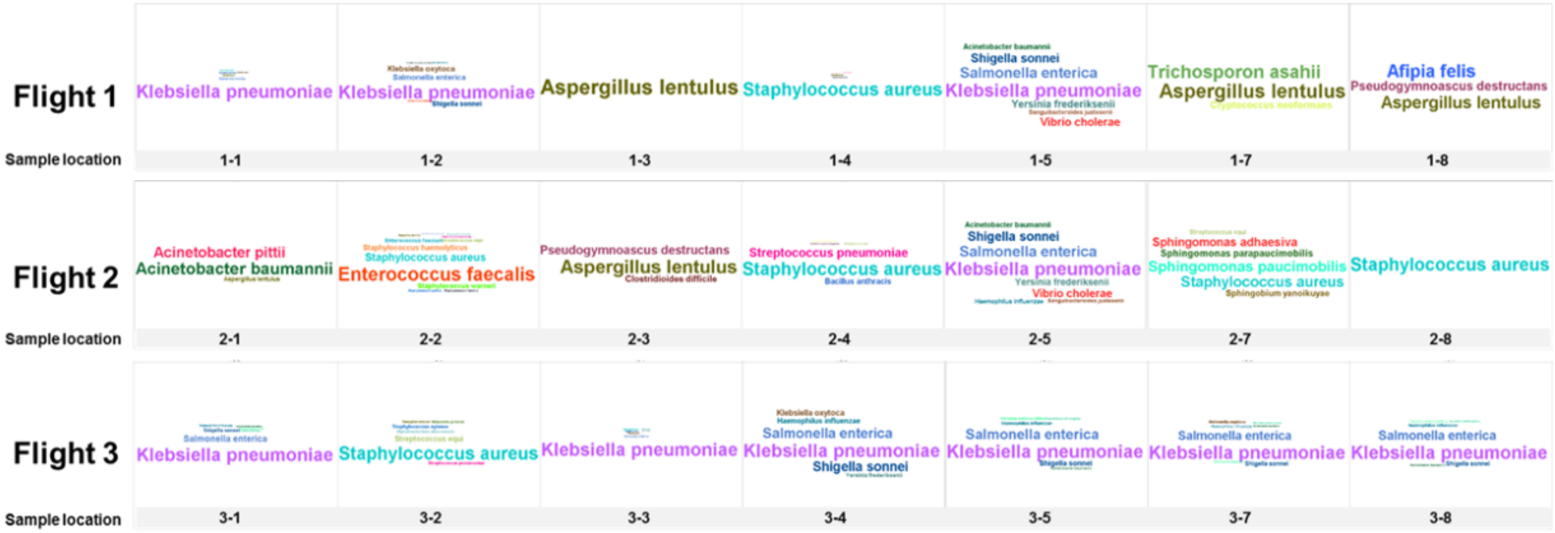


Supplementary Figure S1

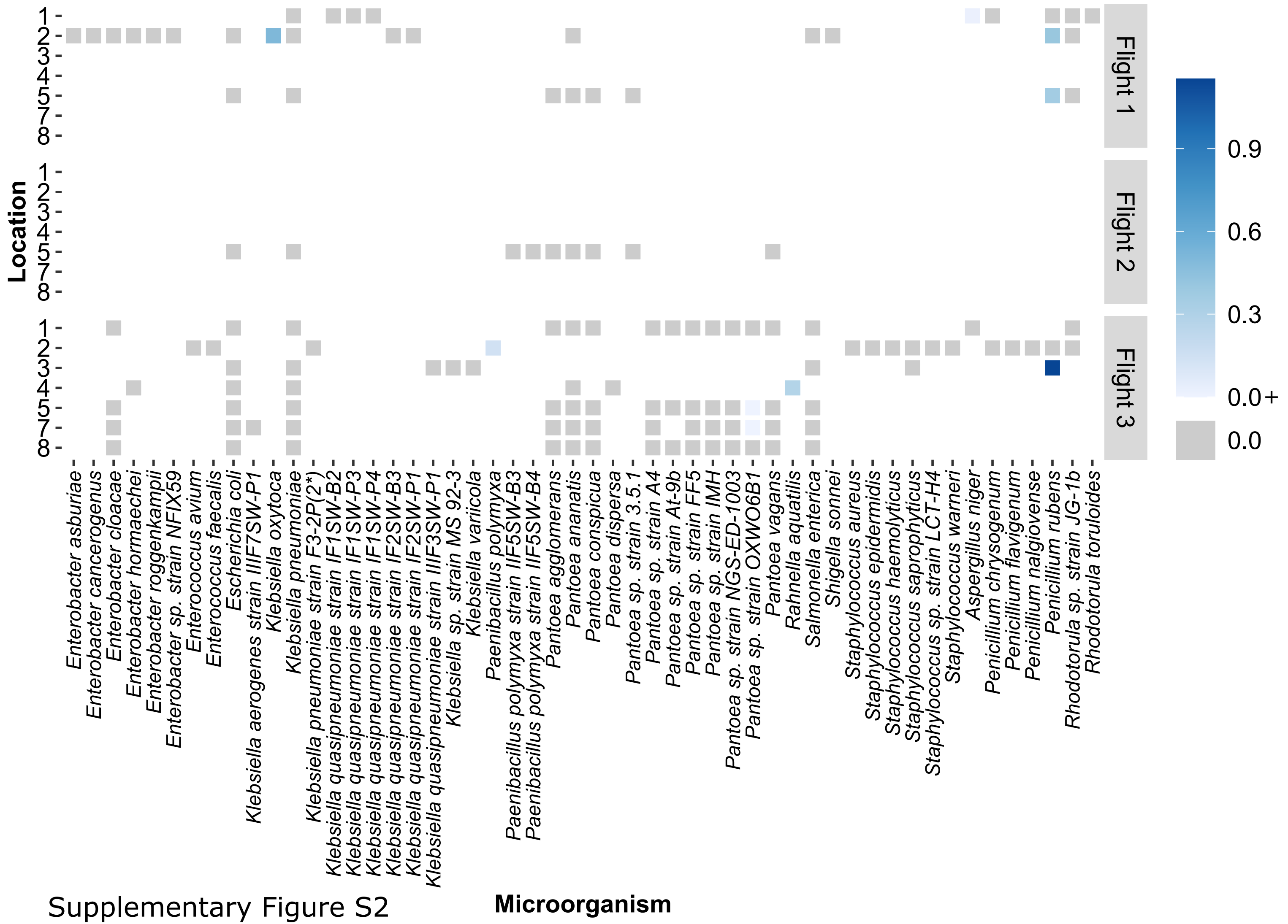


Supplementary Figure S2

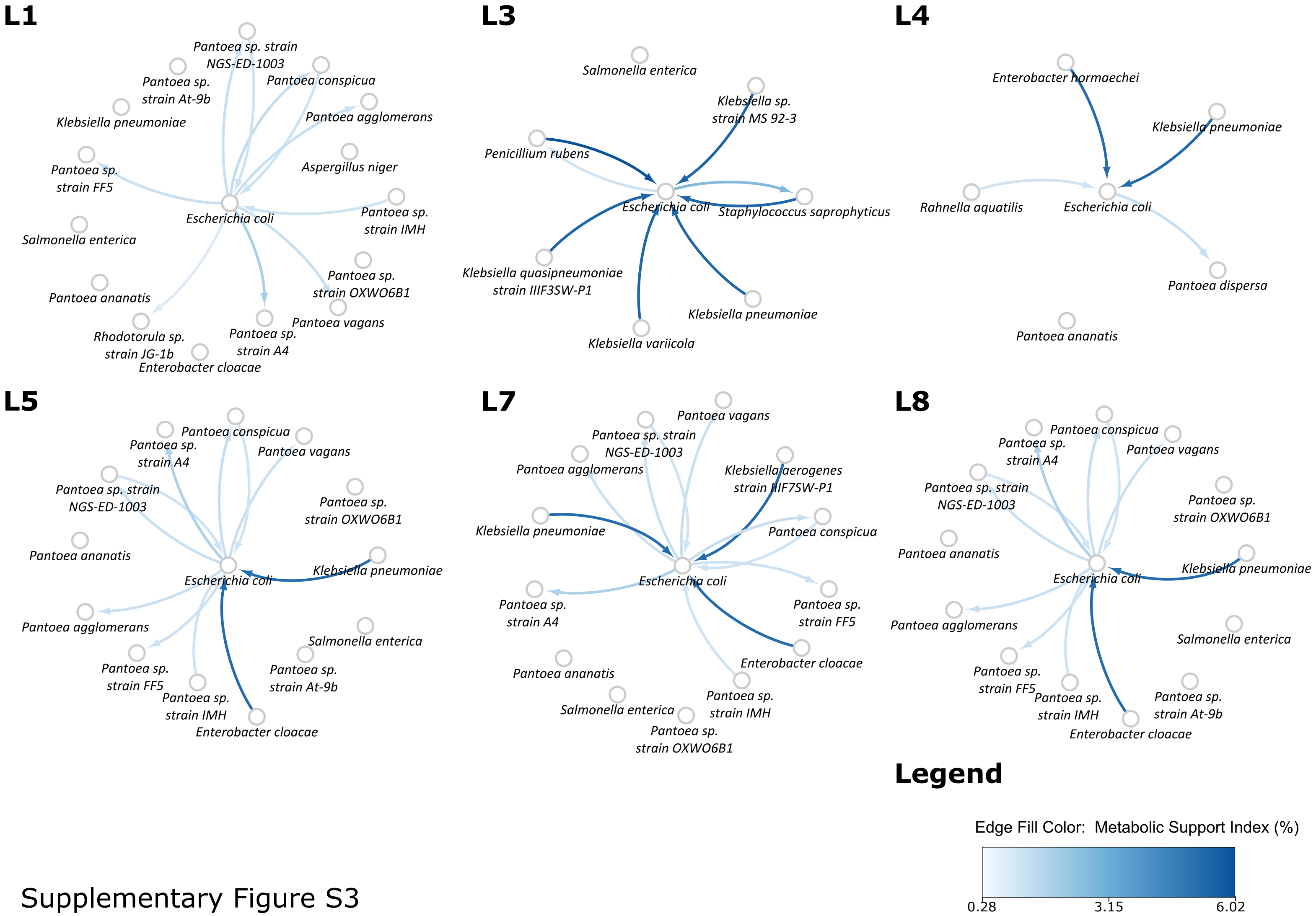


Supplementary Figure S3

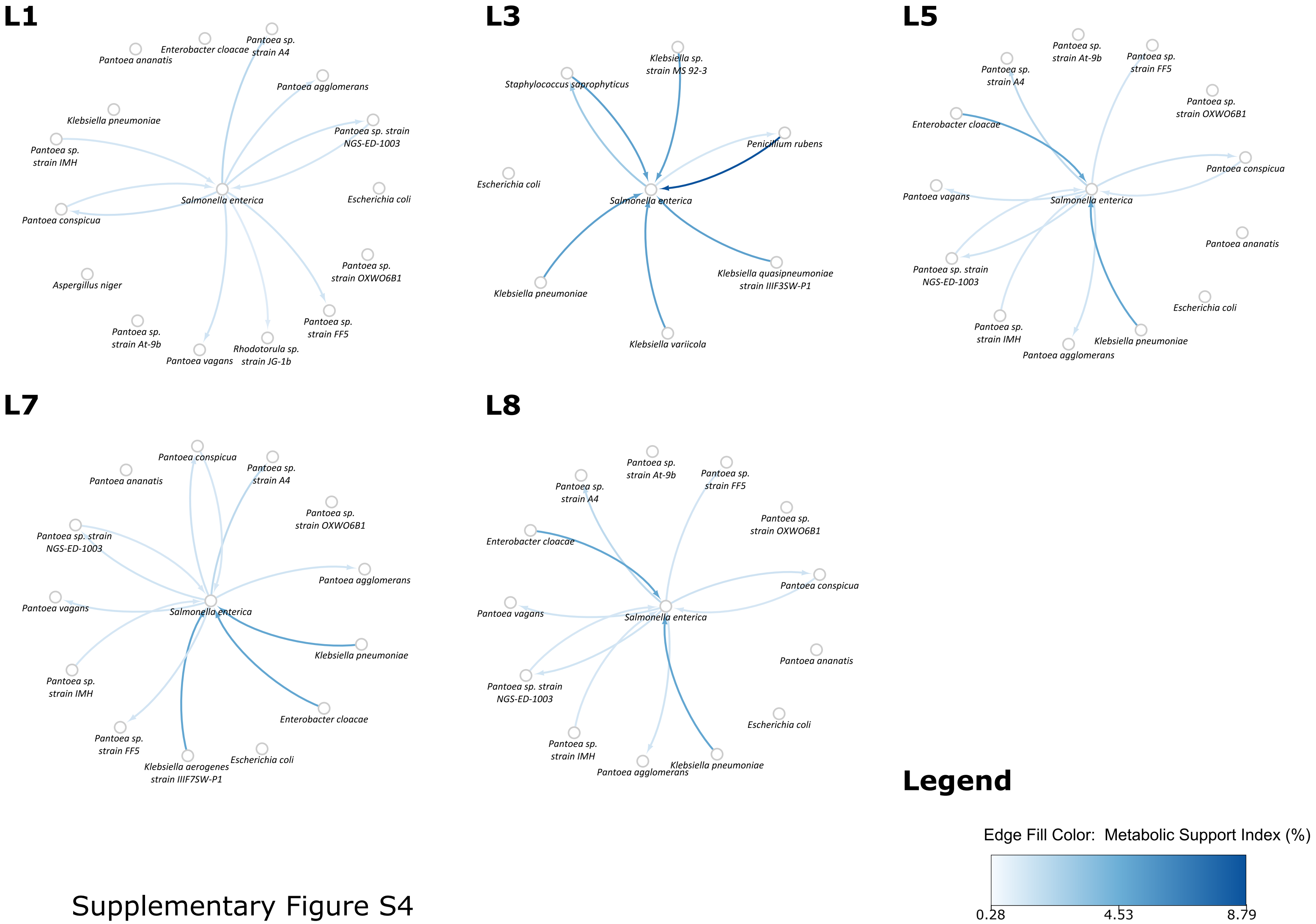


Supplementary Figure S4

### Supplementary Methods M1

#### Protocols involved in the genome-scale metabolic network reconstruction

The genome sequences were, either via direct import from the NCBI RefSeq database or via upload in the case of GenBank genomes, as input to KBase [1] to reconstruct automated genome-scale metabolic models (GSMMs). These GSMMs capture the metabolic network of these microorganisms, to the extent known across published literature and reaction databases [2]. The bacterial genomes were first annotated with RASTtk toolkit v1.073 [3–5]. The metabolic models were then reconstructed using the Build Metabolic Model v2.0.0 app, and a Gram positive/Gram negative template was appropriately selected for reconstruction. The gap-filling was done with a minimal medium readily available in the KBase public media database, RefGlucoseMinimal, and increasing the maximum uptake of Glucose to 10 mmol/gDW-h. For the reconstruction of fungal models, the Build Fungal Model v1.0.0 was used and templates were selected from the available fourteen templates, and gap-filling was also carried out with the default settings.

These models were used for both graph-theoretic analyses and constraint-based analyses. Constraint-based model artifacts, such as the reaction node for biomass, the nodes corresponding to the compounds for biomass, DNA replication, RNA transcription, and Protein biosynthesis, were removed from all the graphs. Supplementary Table S2 contains information on the reconstructions.

**This document contains supplementary information**

**Table S1** **RefSeq and GenBank Accession identification numbers for microorganisms used in this study**

RefSeq and GenBank accession IDs used in this study. As detailed in the comments, the sequences fall into three categories, namely: a) ISS strain sequences b) ISS strain sequence not compatible with KBase hence reference sequence used, and c) ISS strain sequence not available, hence reference sequence used

**Table S2** **Details of genome-scale metabolic network reconstructions**

The table contains information regarding the template for reconstruction, as well as the number of reactions in the model as reported by KBase.

**Table S3** **List of seed metabolites used in the analyses**

The list contains components of minimal medium as obtained from cobrapy, as well as other co-factors and coenzymes.

**Table S4** **Relative abundances of microorganisms in the flight-locations considered**

The table contains the relative abundance of microorganisms in the corresponding location and Flight. Only microorganisms that coexist with *Klebsiella pneumoniae* with a relative abundance of >1% have been shown here.

**Table S5** **Metabolic benefit provided by an individual microorganism to the rest of the community**

Community Support Index (X-A|X) (%)

**Table S6** **Metabolic benefit provided by each family to the rest of the community**

Community Support Index (X-Family|X) (%)

**Table S7** **Metabolic support provided to an individual microorganism by the rest of the community**

Community Support Index (A|X) (%)

**Table S8** **Metabolic support provided to an individual microorganism by another microorganism in the community**

Metabolic Support Index (A|AUB) (%)

**Table S9**

**Nature of interactions as predicted by the constraint-based analyses**

vbio here refers to the growth rate of a microorganism. The growth rates in two scenarios -- individual and in the two-membered community have been calculated. The effect has been determined by calculating the percentage increase in growth rate. The effect was considered significant if the growth rates increased or decreased by 10% or higher.

**Table S1: RefSeq and GenBank Accession identification numbers for microorganisms used in this study**

| Microorganism | Strain | Assembly | Representative | Comment |
|---|---|---|---|---|
| *Aspergillus niger* | CBS 513.88 | GCF_000002855.3_ASM285v2 | yes | ISS sequence not compatible with KBase |
| *Enterobacter asburiae* | PDN3 | GCF_000799205.1_ASM79920v1 | yes | ISS sequence unavailable, reference genome used |
| *Enterobacter cancerogenus* | MiY-F | GCF_009648915.1_ASM964891v1 | yes | ISS sequence unavailable, reference genome used |
| *Enterobacter cloacae* | GGT036 | GCF_000770155.1_ASM77015v1 | yes | ISS sequence unavailable, reference genome used |
| *Enterobacter hormaechei* | YT3 | GCF_000328885.1_SOAPdenovo | yes | ISS sequence unavailable, reference genome used |
| *Enterobacter roggenkampii* | DSM 16690 | GCF_001729805.1_ASM172980v1 | yes | ISS sequence unavailable, reference genome used |
| *Enterobacter* sp. strain NFIX59 | NFIX59 | GCF_900113755.1_IMG-taxon_2602042047 | no | ISS sequence unavailable, reference genome used |
| *Enterococcus avium* | ATCC 14025 | GCF_000407245.1_Ente_aviu_ATCC14025_V2 | yes | ISS sequence unavailable, reference genome used |
| *Enterococcus faecalis* | 39EA1 | GCF_003319815.1_ASM331981v1 | yes | ISS sequence unavailable, reference genome used |
| *Escherichia coli* | K-12 substr. MG1655 | GCF_000005845.2_ASM584v2 | yes | ISS sequence unavailable, reference genome used |
| *Klebsiella aerogenes* strain IIIF7SW-P1 | IIIF7SW-P1 | GCF_013403435.1_ASM1340343v1 | ISS | ISS sequence |
| *Klebsiella oxytoca* | 10-5243 | GCF_000247855.1_Kleb_oxyt_10-5243_V1 | yes | ISS sequence unavailable, reference genome used |
| *Klebsiella pneumoniae* | HS11286 | GCF_000240185.1_ASM24018v2 | yes | ISS sequence unavailable, reference genome used |
| *Klebsiella pneumoniae* strain F3-2P(2*) | F3-2P(2*) | GCF_014162935.1_ASM1416293v1 | ISS | ISS sequence |
| *Klebsiella quasipneumoniae* strain IF1SW-B2 | IF1SW-B2 | GCF_013366635.1_ASM1336663v1 | ISS | ISS sequence |
| *Klebsiella quasipneumoniae* strain IF1SW-P3 | IF1SW-P3 | GCF_013366555.1_ASM1336655v1 | ISS | ISS sequence |
| *Klebsiella quasipneumoniae* strain IF1SW-P4 | IF1SW-P4 | GCF_013366575.1_ASM1336657v1 | ISS | ISS sequence |
| *Klebsiella quasipneumoniae* strain IF2SW-B3 | IF2SW-B3 | GCF_013366565.1_ASM1336656v1 | ISS | ISS sequence |
| *Klebsiella quasipneumoniae* strain IF2SW-P1 | IF2SW-P1 | GCF_013366585.1_ASM1336658v1 | ISS | ISS sequence |
| *Klebsiella quasipneumoniae* strain IIIF3SW-P1 | IIIF3SW-P1 | GCF_013377655.1_ASM1337765v1 | ISS | ISS sequence |
| *Klebsiella* sp. strain MS 92-3 | MS 92-3 | GCF_000195655.1_ASM19565v1 | no | ISS sequence unavailable, reference genome used |
| *Klebsiella variicola* | FH-1 | GCF_013305245.1_ASM1330524v1 | yes | ISS sequence unavailable, reference genome used |
| *Paenibacillus polymyxa* | CF05 | GCF_000785455.1_ASM78545v1 | yes | ISS sequence unavailable, reference genome used |
| *Paenibacillus polymyxa* strain IIF5SW-B3 | IIF5SW-B3 | GCF_013346085.1_ASM1334608v1 | ISS | ISS sequence |
| *Paenibacillus polymyxa* strain IIF5SW-B4 | IIF5SW-B4 | GCF_013345675.1_ASM1334567v1 | ISS | ISS sequence |
| *Pantoea agglomerans* | L15 | GCF_003860325.1_ASM386032v1 | yes | ISS sequence unavailable, reference genome used |
| *Pantoea ananatis* | PA13 | GCF_000233595.1_ASM23359v1 | yes | ISS sequence unavailable, reference genome used |
| *Pantoea conspicua* | LMG 24534 | GCF_002095315.1_ASM209531v1 | yes | ISS sequence unavailable, reference genome used |
| *Pantoea dispersa* | 625 | GCF_009866445.1_ASM986644v1 | yes | ISS sequence unavailable, reference genome used |
| *Pantoea* sp. strain 3.5.1 | 3.5.1 | GCF_000731025.2_ASM73102v2 | no | ISS sequence unavailable, reference genome used |
| *Pantoea* sp. strain A4 | A4 | GCF_000295955.2_ASM29595v2 | no | ISS sequence unavailable, reference genome used |
| *Pantoea* sp. strain At-9b | At-9b | GCF_000175935.2_ASM17593v2 | no | ISS sequence unavailable, reference genome used |
| *Pantoea* sp. strain FF5 | FF5 | GCF_000612605.1_FF5 | yes | ISS sequence unavailable, reference genome used |
| *Pantoea* sp. strain IMH | IMH | GCF_000599885.1_IMH | no | ISS sequence unavailable, reference genome used |
| *Pantoea* sp. strain NGS-ED-1003 | NGS-ED-1003 | GCF_000738765.1_ASM73876v1 | no | ISS sequence unavailable, reference genome used |
| *Pantoea* sp. strain OXWO6B1 | OXWO6B1 | GCF_001641135.1_ASM164113v1 | no | ISS sequence unavailable, reference genome used |
| *Pantoea vagans* | LMG 24199 | GCF_004792415.1_ASM479241v1 | yes | ISS sequence unavailable, reference genome used |
| *Penicillium chrysogenum* | P2niaD18 | GCA_000710275.1_ASM71027v1 | yes | ISS sequence not compatible with KBase |
| *Penicillium flavigenum* | IBT 14082 | GCA_002072365.1_ASM207236v1 | yes | ISS sequence unavailable, reference genome used |
| *Penicillium nalgiovense* | IBT 13039 | GCA_002072425.1_ASM207242v1 | yes | ISS sequence unavailable, reference genome used |
| *Penicillium rubens* | Wisconsin 54-1255 | GCF_000226395.1_PenChr_Nov2007 | yes | ISS sequence unavailable, reference genome used |
| *Rahnella aquatilis* | CIP 78.65 = ATCC 33071 | GCF_000241955.1_ASM24195v1 | yes | ISS sequence unavailable, reference genome used |
| *Rhodotorula* sp. strain JG-1b | JG-1b | GCA_001541205.1_Rhosp1 | yes | ISS sequence unavailable, reference genome used |
| *Rhodotorula toruloides* | NP11 | GCF_000320785.1_RHOziaDV1.0 | yes | ISS sequence unavailable, reference genome used |
| *Salmonella enterica* | LT2 | GCF_000006945.2_ASM694v2 | yes | ISS sequence unavailable, reference genome used |
| *Shigella sonnei* | ECH+12 | GCF_002247485.1_ASM224748v1 | yes | ISS sequence unavailable, reference genome used |
| *Staphylococcus aureus* | NCTC 8325 | GCF_000013425.1_ASM1342v1 | yes (reference) | ISS sequence unavailable, reference genome used |
| *Staphylococcus epidermidis* | ATCC 14990 | GCF_006094375.1_ASM609437v1 | yes | ISS sequence unavailable, reference genome used |
| *Staphylococcus haemolyticus* | S167 | GCF_001611955.1_ASM161195v1 | yes | ISS sequence unavailable, reference genome used |
| *Staphylococcus saprophyticus* | NCTC7666 | GCF_900635295.1_40853_E01 | yes | ISS sequence unavailable, reference genome used |
| *Staphylococcus* sp. strain LCT-H4 | LCT-H4 | GCF_001866145.1_ASM186614v1 | no | ISS sequence unavailable, reference genome used |
| *Staphylococcus warneri* | NCTC4133 | GCF_900636255.1_40295_A02 | yes | ISS sequence unavailable, reference genome used |

**Table S2: Details of genome-scale metabolic network reconstructions**

| Microorganism | Size of the annotated genome (bp) | KBase Rxns | KBase Compounds | KBase Genes | Reactions added during gap-filling |
|---|---|---|---|---|---|
| *Aspergillus niger* | 34,006,671 | 1149 | | | |
| *Enterobacter asburiae* | 4,625,174 | 1541 | 1423 | 1270 | 27 |
| *Enterobacter cancerogenus* | 4,992,573 | 1541 | 1430 | 1281 | 26 |
| *Enterobacter cloacae* | 4,848,754 | 1556 | 1433 | 1306 | 28 |
| *Enterobacter hormaechei* | 4,809,358 | 1547 | 1426 | 1297 | 29 |
| *Enterobacter roggenkampii* | 4,899,997 | 1535 | 1435 | 1277 | 27 |
| *Enterobacter* sp. strain NFIX59 | 4,697,674 | 1567 | 1451 | 1324 | 24 |
| *Enterococcus avium* | 4,614,282 | 1092 | 1085 | 1186 | 77 |
| *Enterococcus faecalis* | 2,707,177 | 993 | 1010 | 727 | 114 |
| *Escherichia coli* | 4,641,652 | 1591 | 1457 | 1299 | 14 |
| *Klebsiella aerogenes* strain IIIF7SW-P1 | 5,147,257 | 1604 | 1462 | 1414 | 22 |
| *Klebsiella oxytoca* | 5,994,455 | 1665 | 1509 | 1678 | 21 |
| *Klebsiella pneumoniae* | 5,682,322 | 1628 | 1490 | 1590 | 21 |
| *Klebsiella pneumoniae* strain F3-2P(2*) | 5,491,924 | 1627 | 1493 | 1527 | 24 |
| *Klebsiella quasipneumoniae* strain IF1SW-B2 | 5,195,913 | 1645 | 1497 | 1554 | 20 |
| *Klebsiella quasipneumoniae* strain IF1SW-P3 | 5,196,283 | 1645 | 1497 | 1554 | 20 |
| *Klebsiella quasipneumoniae* strain IF1SW-P4 | 5,196,206 | 1646 | 1497 | 1554 | 21 |
| *Klebsiella quasipneumoniae* strain IF2SW-B3 | 5,196,325 | 1645 | 1497 | 1554 | 20 |
| *Klebsiella quasipneumoniae* strain IF2SW-P1 | 5,196,423 | 1647 | 1498 | 1554 | 22 |
| *Klebsiella quasipneumoniae* strain IIIF3SW-P1 | 5,157,465 | 1633 | 1494 | 1542 | 22 |
| *Klebsiella* sp. strain MS 92-3 | 5,646,513 | 1617 | 1486 | 1569 | 24 |
| *Klebsiella variicola* | 5,652,418 | 1644 | 1504 | 1587 | 23 |
| *Paenibacillus polymyxa* | 5,762,608 | 1265 | 1247 | 1146 | 52 |
| *Paenibacillus polymyxa* strain IIF5SW-B3 | 5,803,136 | 1283 | 1276 | 1131 | 53 |
| *Paenibacillus polymyxa* strain IIF5SW-B4 | 5,803,259 | 1283 | 1276 | 1131 | 53 |
| *Pantoea agglomerans* | 4,858,869 | 1491 | 1389 | 1235 | 31 |
| *Pantoea ananatis* | 4,867,131 | 1518 | 1404 | 1265 | 32 |
| *Pantoea conspicua* | 4,308,858 | 1388 | 1330 | 1115 | 39 |
| *Pantoea dispersa* | 4,740,205 | 1488 | 1376 | 1247 | 32 |
| *Pantoea* sp. strain 3.5.1 | 4,964,649 | 1514 | 1423 | 1274 | 32 |
| *Pantoea* sp. strain A4 | 5,329,719 | 1495 | 1404 | 1301 | 31 |
| *Pantoea* sp. strain At-9b | 6,312,783 | 1573 | 1473 | 1564 | 29 |
| *Pantoea* sp. strain FF5 | 4,548,084 | 1500 | 1410 | 1222 | 33 |
| *Pantoea* sp. strain IMH | 4,091,152 | 1415 | 1300 | 1122 | 29 |
| *Pantoea* sp. strain NGS-ED-1003 | 4,809,062 | 1506 | 1416 | 1241 | 32 |
| *Pantoea* sp. strain OXWO6B1 | 5,236,140 | 1498 | 1380 | 1288 | 32 |
| *Pantoea vagans* | 4,790,329 | 1483 | 1374 | 1254 | 33 |
| *Penicillium chrysogenum* | 32,525,541 | 1216 | | | |
| *Penicillium flavigenum* | 32,903,267 | 1147 | | | |
| *Penicillium nalgiovense* | 33,006,161 | 1153 | | | |
| *Penicillium rubens* | 32,223,735 | 1221 | | | |
| *Rahnella aquatilis* | 5,448,900 | 1530 | 1394 | 1398 | 20 |
| *Rhodotorula* sp. strain JG-1b | 19,393,393 | 876 | | | |
| *Rhodotorula toruloides* | 20,223,942 | 920 | | | |
| *Salmonella enterica* | 4,951,383 | 1546 | 1422 | 1272 | 19 |
| *Shigella sonnei* | 4,822,833 | 1593 | 1462 | 1299 | 17 |
| *Staphylococcus aureus* | 2,821,361 | 1108 | 1100 | 772 | 67 |
| *Staphylococcus epidermidis* | 2,491,058 | 1086 | 1079 | 753 | 62 |
| *Staphylococcus haemolyticus* | 2,560,146 | 1106 | 1103 | 751 | 73 |
| *Staphylococcus saprophyticus* | 2,610,800 | 1117 | 1104 | 785 | 69 |
| *Staphylococcus* sp. strain LCT-H4 | 2,782,699 | 1175 | 1155 | 863 | 60 |
| *Staphylococcus warneri* | 2,430,678 | 1104 | 1080 | 744 | 76 |

**Table S3: List of seed metabolites used in the analyses**

| Compound name | Compound ID | F1-1 | F1-2 | F1-5 | F2-5 | F3-1 | F3-2 | F3-3 | F3-4 | F3-5 | F3-7 | F3-8 |
|---|---|---|---|---|---|---|---|---|---|---|---|---|
| 2-Oxoglutarate | cpd00024 | * | * | * | * | * | * | * | * | * | * | * |
| 4-Hydroxybenzoate | cpd00136 | | | | | | * | | | | | |
| 5,10-Methylenetetrahydrofolate | cpd00125 | * | * | * | * | * | * | * | * | * | * | * |
| 5-Aminolevulinate | cpd00338 | * | * | * | | | * | * | | | | |
| Acetyl-CoA | cpd00022 | * | * | * | * | * | * | * | * | * | * | * |
| Acyl Carrier Protein | cpd11493 | * | * | * | * | * | * | * | * | * | * | * |
| Acyl-CoA | cpd11611 | * | * | * | | * | * | * | | | | |
| Adenine | cpd00128 | * | * | * | * | * | * | * | * | * | * | * |
| Adenosine | cpd00182 | * | * | * | * | * | * | * | * | * | * | * |
| ADP | cpd00008 | * | * | * | * | * | * | * | * | * | * | * |
| Ala-Gln | cpd11587 | | | | * | | | | | | | |
| Aminoethanol | cpd00162 | | | | | | * | | | | | |
| AMP | cpd00018 | * | * | * | * | * | * | * | * | * | * | * |
| ATP | cpd00002 | * | * | * | * | * | * | * | * | * | * | * |
| BIOT | cpd00104 | * | * | * | * | * | * | * | * | * | * | * |
| Ca2+ | cpd00063 | * | * | * | * | * | * | * | * | * | * | * |
| CDP | cpd00096 | * | * | * | * | * | * | * | * | * | * | * |
| Choline | cpd00098 | * | * | * | | * | * | | | | | |
| Ciliatine | cpd02233 | | | | | | * | | | | | |
| Cl- | cpd00099 | * | * | * | * | * | * | * | * | * | * | * |
| CMP | cpd00046 | * | * | * | * | * | * | * | * | * | * | * |
| Co2+ | cpd00149 | * | * | * | * | * | * | * | * | * | * | * |
| CoA | cpd00010 | * | * | * | * | * | * | * | * | * | * | * |
| Coproporphyrinogen III | cpd02083 | * | * | * | * | * | * | * | * | * | * | * |
| CTP | cpd00052 | * | * | * | * | * | * | * | * | * | * | * |
| Cu2+ | cpd00058 | * | * | * | * | * | * | * | * | * | * | * |
| Cytidine | cpd00367 | * | * | * | * | * | * | * | * | * | * | * |
| Cytosine | cpd00307 | * | * | * | * | * | * | * | * | * | * | * |
| dADP | cpd00177 | * | * | * | * | * | * | * | * | * | * | * |
| dAMP | cpd00294 | * | * | * | * | * | * | * | * | * | * | * |
| dATP | cpd00115 | * | * | * | * | * | * | * | * | * | * | * |
| dCDP | cpd00533 | * | * | * | * | * | * | * | * | * | * | * |
| dCMP | cpd00206 | * | * | * | * | * | * | * | * | * | * | * |
| dCTP | cpd00356 | * | * | * | * | * | * | * | * | * | * | * |
| Deoxyadenosine | cpd00438 | * | * | * | * | * | * | * | * | * | * | * |
| Deoxycytidine | cpd00654 | * | * | * | * | * | * | * | * | * | * | * |
| Deoxyguanosine | cpd00277 | * | * | * | * | * | * | * | * | * | * | * |
| Deoxyuridine | cpd00412 | * | * | * | * | * | * | * | * | * | * | * |
| D-fructose | dfru-e | * | | | | * | | | | | | |
| dGDP | cpd00295 | * | * | * | * | * | * | * | * | * | * | * |
| D-Glucose | cpd00027 | * | * | * | | * | * | | | | | |
| dGMP | cpd00296 | * | * | * | * | * | * | * | * | * | * | * |
| dGTP | cpd00241 | * | * | * | * | * | * | * | * | * | * | * |
| dIDP | cpd00976 | * | * | * | * | * | * | * | * | * | * | * |
| dIMP | cpd03704 | * | * | * | * | * | * | * | * | * | * | * |
| dITP | cpd00977 | * | * | * | * | * | * | * | * | * | * | * |
| dUDP | cpd00978 | * | * | * | * | * | * | * | * | * | * | * |
| dUMP | cpd00299 | * | * | * | * | * | * | * | * | * | * | * |
| dUTP | cpd00358 | * | * | * | * | * | * | * | * | * | * | * |
| Ergosterol | cpd01170 | * | * | * | | * | * | | | | | |
| FAD | cpd00015 | * | * | * | * | * | * | * | * | * | * | * |
| Fe2+ | cpd10515 | * | * | * | * | * | * | * | * | * | * | * |
| fe3 | cpd10516 | * | * | * | * | * | * | * | * | * | * | * |
| Flavin adenine dinucleotide reduced | cpd00982 | * | * | * | * | * | * | * | * | * | * | * |
| FMN | cpd00050 | * | * | * | * | * | * | * | * | * | * | * |
| Folate | cpd00393 | * | * | * | * | * | * | * | * | * | * | * |
| Galactose | cpd00108 | * | * | * | | | * | * | | | | |
| GDP | cpd00031 | * | * | * | * | * | * | * | * | * | * | * |
| GDP-mannose | cpd00083 | * | | | | * | | | | | | |
| GLUM | cpd00276 | | | | | | * | | | | | |
| gly-asn-L | cpd11581 | | | | * | | | | | | | |
| Glycerol | cpd00100 | * | * | * | | * | * | | | | | |
| Glycerol-3-phosphate | cpd00080 | | | | * | | * | | | | | |
| Glycine | cpd00033 | * | * | * | | * | * | | | | | |
| GMP | cpd00126 | * | * | * | * | * | * | * | * | * | * | * |
| GTP | cpd00038 | * | * | * | * | * | * | * | * | * | * | * |
| Guanine | cpd00207 | * | * | * | * | * | * | * | * | * | * | * |
| Guanosine | cpd00311 | * | * | * | * | * | * | * | * | * | * | * |
| H+ | cpd00067 | * | * | * | * | * | * | * | * | * | * | * |
| H2O | cpd00001 | * | * | * | * | * | * | * | * | * | * | * |

| Compound name | Compound ID | F1-1 | F1-2 | F1-5 | F2-5 | F3-1 | F3-2 | F3-3 | F3-4 | F3-5 | F3-7 | F3-8 |
|---|---|---|---|---|---|---|---|---|---|---|---|---|
| HCO3 | cpd00242 | * | * | * | * | * | * | * | * | * | * | * |
| Heme | cpd00028 | * | * | * | * | * | * | * | * | * | * | * |
| IDP | cpd00090 | * | * | * | * | * | * | * | * | * | * | * |
| IMP | cpd00114 | * | * | * | * | * | * | * | * | * | * | * |
| ITP | cpd00068 | * | * | * | * | * | * | * | * | * | * | * |
| K+ | cpd00205 | * | * | * | * | * | * | * | * | * | * | * |
| L-Alanine | cpd00035 | * | | | | | | | | | | |
| L-Arginine | cpd00051 | * | * | * | | * | * | * | | | | |
| L-Asparagine | cpd00132 | * | | | | * | | | | | | |
| L-Cysteine | cpd00084 | * | * | * | | * | * | | | | | |
| Lecithin | cpd11624 | * | | | | | | | | | | |
| L-Glutamine | cpd00053 | * | * | * | | * | * | | | | | |
| L-Histidine | cpd00119 | * | * | * | | * | * | * | | | | |
| L-Inositol | cpd00121 | * | * | * | | * | * | | | | | |
| L-Isoleucine | cpd00322 | * | * | * | | * | * | * | | | | |
| L-Leucine | cpd00107 | * | * | * | | * | * | * | | | | |
| L-Lysine | cpd00039 | * | * | * | | * | * | * | | | | |
| L-Methionine | cpd00060 | * | * | * | | * | * | * | | | | |
| L-Phenylalanine | cpd00066 | | | | | | * | | | | | |
| L-Proline | cpd00129 | * | * | * | | * | * | * | | | | |
| L-Serine | cpd00054 | * | * | * | | * | * | | | | | |
| L-Threonine | cpd00161 | | | | | | * | | | | | |
| L-Tryptophan | cpd00065 | * | * | * | | * | * | | | | | |
| L-Tyrosine | cpd00069 | | | | | | * | | | | | |
| L-Valine | cpd00156 | * | * | * | | * | * | * | | | | |
| Mannan | cpd11685 | * | * | * | | * | * | | | | | |
| Mg | cpd00254 | * | * | * | * | * | * | * | * | * | * | * |
| Mn2+ | cpd00030 | * | * | * | * | * | * | * | * | * | * | * |
| Na+ | cpd00971 | * | * | * | * | * | * | * | * | * | * | * |
| N-Acetylmuramate | cpd01757 | | | | | | * | | | | | |
| NAD | cpd00003 | * | * | * | * | * | * | * | * | * | * | * |
| NADH | cpd00004 | * | * | * | * | * | * | * | * | * | * | * |
| NADP | cpd00006 | * | * | * | * | * | * | * | * | * | * | * |
| NADPH | cpd00005 | * | * | * | * | * | * | * | * | * | * | * |
| NH4+ | cpd19013 | * | * | * | * | * | * | * | * | * | * | * |
| Niacin | cpd00218 | * | * | * | * | * | * | * | * | * | * | * |
| Nicotinamide ribonucleotide | cpd00355 | * | * | * | * | * | * | * | * | * | * | * |
| Nitrate | cpd00209 | * | * | * | * | * | * | * | * | * | * | * |
| Nitrite | cpd00075 | * | * | * | * | * | * | * | * | * | * | * |
| O2 | cpd00007 | * | * | * | * | * | * | * | * | * | * | * |
| ocdca | cpd01080 | | | | | | * | | | | | |
| Octodecanoyl-ACP | cpd11573 | * | | | | * | | | | | | |
| Ornithine | cpd00064 | | | | | | * | | | | | |
| Phosphate | cpd00009 | * | * | * | * | * | * | * | * | * | * | * |
| Phosphoethanolamine | cpd00285 | * | | | | | | | | | | |
| Phospholipids | cpd27865 | * | * | * | | | * | * | | | | |
| PPi | cpd00012 | * | * | * | * | * | * | * | * | * | * | * |
| Pyridoxal | cpd00215 | * | * | * | * | * | * | * | * | * | * | * |
| Riboflavin | cpd00220 | * | * | * | * | * | * | * | * | * | * | * |
| S-Adenosyl-L-homocysteine | cpd00019 | * | * | * | * | * | * | * | * | * | * | * |
| S-Adenosyl-L-methionine | cpd00017 | * | * | * | * | * | * | * | * | * | * | * |
| Sulfate | cpd00048 | * | * | * | * | * | * | * | * | * | * | * |
| TDP | cpd00297 | * | * | * | * | * | * | * | * | * | * | * |
| Thiamin | cpd00305 | * | * | * | * | * | * | * | * | * | * | * |
| Thymidine | cpd00184 | * | * | * | * | * | * | * | * | * | * | * |
| Thymine | cpd00151 | * | * | * | * | * | * | * | * | * | * | * |
| TMP | cpd00298 | * | * | * | * | * | * | * | * | * | * | * |
| TTP | cpd00357 | * | * | * | * | * | * | * | * | * | * | * |
| Ubiquinol-8 | cpd15561 | * | * | * | * | * | * | * | * | * | * | * |
| Ubiquinone-8 | cpd15560 | * | * | * | * | * | * | * | * | * | * | * |
| UDP | cpd00014 | * | * | * | * | * | * | * | * | * | * | * |
| UMP | cpd00091 | * | * | * | * | * | * | * | * | * | * | * |
| Uracil | cpd00092 | * | * | * | * | * | * | * | * | * | * | * |
| Uridine | cpd00249 | * | * | * | * | * | * | * | * | * | * | * |
| UTP | cpd00062 | * | * | * | * | * | * | * | * | * | * | * |
| Zn2+ | cpd00034 | * | * | * | * | * | * | * | * | * | * | * |
| Zymosterol | cpd03221 | * | * | * | | * | * | | | | | |

**Table S4: Relative abundances of microorganisms in the flight-locations considered**

| Microorganism | F1-1P | F1-2P | F1-5P | F2-5P | F3-1P | F3-2P | F3-3P | F3-4P | F3-5P | F3-7P | F3-8P |
|---|---|---|---|---|---|---|---|---|---|---|---|
| *Aspergillus niger* | 2.6950 | | | | 1.7809 | | | | | | |
| *Enterobacter asburiae* | | 3.1692 | | | | | | | | | |
| *Enterobacter cancerogenus* | | 1.6799 | | | | | | | | | |
| *Enterobacter cloacae* | | 40.4563 | | | 1.2416 | | | | 1.4452 | 1.1758 | 1.4222 |
| *Enterobacter hormaechei* | | 2.6764 | | | | | | 1.3348 | | | |
| *Enterobacter roggenkampii* | | 1.5267 | | | | | | | | | |
| *Enterobacter* sp. strain NFIX59 | | 1.3482 | | | | | | | | | |
| *Enterococcus avium* | | | | | | 3.9084 | | | | | |
| *Enterococcus faecalis* | | | | | | 11.8369 | | | | | |
| *Escherichia coli* | | 4.0754 | 1.1100 | 1.4012 | 3.8304 | | 2.7405 | 2.5528 | 4.4036 | 3.0074 | 4.0842 |
| *Klebsiella aerogenes* strain IIIF7SW-P1 | | | | | | | | | | 39.1339 | |
| *Klebsiella oxytoca* | | 2.9988 | | | | | | | | | |
| *Klebsiella pneumoniae* | 9.4072 | 22.1549 | 0.6361 | 0.8923 | 12.8160 | | 69.7300 | 1.1991 | 5.8714 | 6.3196 | 7.2692 |
| *Klebsiella pneumoniae* strain F3-2P(2*) | | | | | | 0.0003 | | | | | |
| *Klebsiella quasipneumoniae* strain IF1SW-B2 | | | | | | | | | | | |
| *Klebsiella quasipneumoniae* strain IF1SW-P3 | 1.0634 | | | | | | | | | | |
| *Klebsiella quasipneumoniae* strain IF1SW-P4 | | | | | | | | | | | |
| *Klebsiella quasipneumoniae* strain IF2SW-B3 | | 2.1124 | | | | | | | | | |
| *Klebsiella quasipneumoniae* strain IF2SW-P1 | | | | | | | | | | | |
| *Klebsiella quasipneumoniae* strain IIIF3SW-P1 | | | | | | | 7.7465 | | | | |
| *Klebsiella* sp. strain MS 92-3 | | | | | | | 1.2438 | | | | |
| *Klebsiella variicola* | | | | | | | 1.9551 | | | | |
| *Paenibacillus polymyxa* | | | | | | 6.1113 | | | | | |
| *Paenibacillus polymyxa* strain IIF5SW-B3 | | | | 2.1013 | | | | | | | |
| *Paenibacillus polymyxa* strain IIF5SW-B4 | | | | | | | | | | | |
| *Pantoea agglomerans* | | | 2.8332 | | 2.1787 | | | | 2.5743 | 1.3857 | 2.3939 |
| *Pantoea ananatis* | | 1.0429 | 3.4213 | 5.1279 | 9.4144 | | | 5.2446 | 10.1496 | 5.5660 | 9.8517 |
| *Pantoea conspicua* | | | 56.1206 | 73.9494 | 1.3448 | | | | 1.6095 | 1.1505 | 1.5426 |
| *Pantoea dispersa* | | | | | | | | 69.0180 | | | |
| *Pantoea* sp. strain 3.5.1 | | | 4.7175 | 6.6654 | | | | | | | |
| *Pantoea* sp. strain A4 | | | | | 2.2646 | | | | 2.4550 | 1.3064 | 2.4980 |
| *Pantoea* sp. strain At-9b | | | | | 1.0249 | | | | 1.2000 | | 1.1336 |
| *Pantoea* sp. strain FF5 | | | | | 14.4662 | | | | 15.7446 | 8.5105 | 16.1284 |
| *Pantoea* sp. strain IMH | | | | | 1.7618 | | | | 1.8758 | 1.0455 | 1.8996 |
| *Pantoea* sp. strain NGS-ED-1003 | | | | | 10.6271 | | | | 12.0618 | 6.4060 | 11.6334 |
| *Pantoea* sp. strain OXWO6B1 | | | | | 2.8918 | | | | 3.4149 | 1.7835 | 3.2142 |
| *Pantoea vagans* | | | | 1.1478 | 10.8985 | | | | 12.0321 | 7.0203 | 12.0197 |
| *Penicillium chrysogenum* | 2.0348 | | | | | 2.7061 | | | | | |
| *Penicillium flavigenum* | | | | | | 1.3604 | | | | | |
| *Penicillium nalgiovense* | | | | | | 2.0064 | | | | | |
| *Penicillium rubens* | 16.4952 | 1.1793 | 7.6740 | | | 13.7434 | 2.2977 | | | | |
| *Rahnella aquatilis* | | | | | | | | 1.3954 | | | |
| *Rhodotorula* sp. strain JG-1b | 59.3254 | 2.0681 | 16.5376 | | 1.6914 | 5.3056 | | | | | |
| *Rhodotorula toruloides* | 1.3452 | | | | | | | | | | |
| *Salmonella enterica* | | 3.3968 | | | 2.5982 | | 1.3580 | | 2.8379 | 2.2623 | 2.8345 |
| *Shigella sonnei* | | 1.6180 | | | | | | | | | |
| *Staphylococcus aureus* | | | | | | 2.2957 | | | | | |
| *Staphylococcus epidermidis* | | | | | | 1.0787 | | | | | |
| *Staphylococcus haemolyticus* | | | | | | 2.2603 | | | | | |
| *Staphylococcus saprophyticus* | | | | | | 30.8971 | 7.1234 | | | | |
| *Staphylococcus* sp. strain LCT-H4 | | | | | | 1.2604 | | | | | |
| *Staphylococcus warneri* | | | | | | 1.9677 | | | | | |

**Table S5: Metabolic benefit provided by an individual microorganism to the rest of the community**

| Microorganism | Flight | Location | Community Support Index (X-A\|X) (%) |
|---|---|---|---|
| *Penicillium rubens* | Flight 3 | 3 | 1.15 |
| *Klebsiella oxytoca* | Flight 1 | 2 | 0.49 |
| *Penicillium rubens* | Flight 1 | 2 | 0.4 |
| *Penicillium rubens* | Flight 1 | 5 | 0.36 |
| *Rahnella aquatilis* | Flight 3 | 4 | 0.27 |
| *Paenibacillus polymyxa* | Flight 3 | 2 | 0.13 |
| *Aspergillus niger* | Flight 1 | 1 | 0.02 |
| *Aspergillus niger* | Flight 3 | 1 | 0 |
| *Enterobacter asburiae* | Flight 1 | 2 | 0 |
| *Enterobacter cancerogenus* | Flight 1 | 2 | 0 |
| *Enterobacter cloacae* | Flight 1 | 2 | 0 |
| *Enterobacter cloacae* | Flight 3 | 1 | 0 |
| *Enterobacter cloacae* | Flight 3 | 5 | 0 |
| *Enterobacter cloacae* | Flight 3 | 7 | 0 |
| *Enterobacter cloacae* | Flight 3 | 8 | 0 |
| *Enterobacter hormaechei* | Flight 1 | 2 | 0 |
| *Enterobacter hormaechei* | Flight 3 | 4 | 0 |
| *Enterobacter roggenkampii* | Flight 1 | 2 | 0 |
| *Enterobacter* sp. strain NFIX59 | Flight 1 | 2 | 0 |
| *Enterococcus avium* | Flight 3 | 2 | 0 |
| *Enterococcus faecalis* | Flight 3 | 2 | 0 |
| *Escherichia coli* | Flight 1 | 2 | 0 |
| *Escherichia coli* | Flight 1 | 5 | 0 |
| *Escherichia coli* | Flight 2 | 5 | 0 |
| *Escherichia coli* | Flight 3 | 1 | 0 |
| *Escherichia coli* | Flight 3 | 3 | 0 |
| *Escherichia coli* | Flight 3 | 4 | 0 |
| *Escherichia coli* | Flight 3 | 5 | 0 |
| *Escherichia coli* | Flight 3 | 7 | 0 |
| *Escherichia coli* | Flight 3 | 8 | 0 |
| *Klebsiella aerogenes* strain IIIF7SW-P1 | Flight 3 | 7 | 0 |
| *Klebsiella pneumoniae* | Flight 1 | 1 | 0 |
| *Klebsiella pneumoniae* | Flight 1 | 2 | 0 |
| *Klebsiella pneumoniae* | Flight 1 | 5 | 0 |
| *Klebsiella pneumoniae* | Flight 2 | 5 | 0 |
| *Klebsiella pneumoniae* | Flight 3 | 1 | 0 |
| *Klebsiella pneumoniae* | Flight 3 | 3 | 0 |
| *Klebsiella pneumoniae* | Flight 3 | 4 | 0 |
| *Klebsiella pneumoniae* | Flight 3 | 5 | 0 |
| *Klebsiella pneumoniae* | Flight 3 | 7 | 0 |
| *Klebsiella pneumoniae* | Flight 3 | 8 | 0 |
| *Klebsiella pneumoniae* strain F3-2P(2*) | Flight 3 | 2 | 0 |
| *Klebsiella quasipneumoniae* strain IF1SW-B2 | Flight 1 | 1 | 0 |
| *Klebsiella quasipneumoniae* strain IF1SW-P3 | Flight 1 | 1 | 0 |
| *Klebsiella quasipneumoniae* strain IF1SW-P4 | Flight 1 | 1 | 0 |
| *Klebsiella quasipneumoniae* strain IF2SW-B3 | Flight 1 | 2 | 0 |
| *Klebsiella quasipneumoniae* strain IF2SW-P1 | Flight 1 | 2 | 0 |
| *Klebsiella quasipneumoniae* strain IIIF3SW-P1 | Flight 3 | 3 | 0 |
| *Klebsiella* sp. strain MS 92-3 | Flight 3 | 3 | 0 |
| *Klebsiella variicola* | Flight 3 | 3 | 0 |
| *Paenibacillus polymyxa* strain IIF5SW-B3 | Flight 2 | 5 | 0 |
| *Paenibacillus polymyxa* strain IIF5SW-B4 | Flight 2 | 5 | 0 |
| *Pantoea agglomerans* | Flight 1 | 5 | 0 |
| *Pantoea agglomerans* | Flight 2 | 5 | 0 |
| *Pantoea agglomerans* | Flight 3 | 1 | 0 |
| *Pantoea agglomerans* | Flight 3 | 5 | 0 |
| *Pantoea agglomerans* | Flight 3 | 7 | 0 |
| *Pantoea agglomerans* | Flight 3 | 8 | 0 |
| *Pantoea ananatis* | Flight 1 | 2 | 0 |
| *Pantoea ananatis* | Flight 1 | 5 | 0 |
| *Pantoea ananatis* | Flight 2 | 5 | 0 |
| *Pantoea ananatis* | Flight 3 | 1 | 0 |
| *Pantoea ananatis* | Flight 3 | 4 | 0 |
| *Pantoea ananatis* | Flight 3 | 5 | 0 |
| *Pantoea ananatis* | Flight 3 | 7 | 0 |

| Microorganism | Flight | Location | Community Support Index (X-A\|X) (%) |
|---|---|---|---|
| *Pantoea ananatis* | Flight 3 | 8 | 0 |
| *Pantoea conspicua* | Flight 1 | 5 | 0 |
| *Pantoea conspicua* | Flight 2 | 5 | 0 |
| *Pantoea conspicua* | Flight 3 | 1 | 0 |
| *Pantoea conspicua* | Flight 3 | 5 | 0 |
| *Pantoea conspicua* | Flight 3 | 7 | 0 |
| *Pantoea conspicua* | Flight 3 | 8 | 0 |
| *Pantoea dispersa* | Flight 3 | 4 | 0 |
| *Pantoea* sp. strain 3.5.1 | Flight 1 | 5 | 0 |
| *Pantoea* sp. strain 3.5.1 | Flight 2 | 5 | 0 |
| *Pantoea* sp. strain A4 | Flight 3 | 1 | 0 |
| *Pantoea* sp. strain A4 | Flight 3 | 5 | 0 |
| *Pantoea* sp. strain A4 | Flight 3 | 7 | 0 |
| *Pantoea* sp. strain A4 | Flight 3 | 8 | 0 |
| *Pantoea* sp. strain At-9b | Flight 3 | 1 | 0 |
| *Pantoea* sp. strain At-9b | Flight 3 | 5 | 0 |
| *Pantoea* sp. strain At-9b | Flight 3 | 8 | 0 |
| *Pantoea* sp. strain FF5 | Flight 3 | 1 | 0 |
| *Pantoea* sp. strain FF5 | Flight 3 | 5 | 0 |
| *Pantoea* sp. strain FF5 | Flight 3 | 7 | 0 |
| *Pantoea* sp. strain FF5 | Flight 3 | 8 | 0 |
| *Pantoea* sp. strain IMH | Flight 3 | 1 | 0 |
| *Pantoea* sp. strain IMH | Flight 3 | 5 | 0 |
| *Pantoea* sp. strain IMH | Flight 3 | 7 | 0 |
| *Pantoea* sp. strain IMH | Flight 3 | 8 | 0 |
| *Pantoea* sp. strain NGS-ED-1003 | Flight 3 | 1 | 0 |
| *Pantoea* sp. strain NGS-ED-1003 | Flight 3 | 5 | 0 |
| *Pantoea* sp. strain NGS-ED-1003 | Flight 3 | 7 | 0 |
| *Pantoea* sp. strain NGS-ED-1003 | Flight 3 | 8 | 0 |
| *Pantoea* sp. strain OXWO6B1 | Flight 3 | 1 | 0 |
| *Pantoea* sp. strain OXWO6B1 | Flight 3 | 5 | 0 |
| *Pantoea* sp. strain OXWO6B1 | Flight 3 | 7 | 0 |
| *Pantoea* sp. strain OXWO6B1 | Flight 3 | 8 | 0 |
| *Pantoea vagans* | Flight 2 | 5 | 0 |
| *Pantoea vagans* | Flight 3 | 1 | 0 |
| *Pantoea vagans* | Flight 3 | 5 | 0 |
| *Pantoea vagans* | Flight 3 | 7 | 0 |
| *Pantoea vagans* | Flight 3 | 8 | 0 |
| *Penicillium chrysogenum* | Flight 1 | 1 | 0 |
| *Penicillium chrysogenum* | Flight 3 | 2 | 0 |
| *Penicillium flavigenum* | Flight 3 | 2 | 0 |
| *Penicillium nalgiovense* | Flight 3 | 2 | 0 |
| *Penicillium rubens* | Flight 1 | 1 | 0 |
| *Penicillium rubens* | Flight 3 | 2 | 0 |
| *Rhodotorula* sp. strain JG-1b | Flight 1 | 1 | 0 |
| *Rhodotorula* sp. strain JG-1b | Flight 1 | 2 | 0 |
| *Rhodotorula* sp. strain JG-1b | Flight 1 | 5 | 0 |
| *Rhodotorula* sp. strain JG-1b | Flight 3 | 1 | 0 |
| *Rhodotorula* sp. strain JG-1b | Flight 3 | 2 | 0 |
| *Rhodotorula toruloides* | Flight 1 | 1 | 0 |
| *Salmonella enterica* | Flight 1 | 2 | 0 |
| *Salmonella enterica* | Flight 3 | 1 | 0 |
| *Salmonella enterica* | Flight 3 | 3 | 0 |
| *Salmonella enterica* | Flight 3 | 5 | 0 |
| *Salmonella enterica* | Flight 3 | 7 | 0 |
| *Salmonella enterica* | Flight 3 | 8 | 0 |
| *Shigella sonnei* | Flight 1 | 2 | 0 |
| *Staphylococcus aureus* | Flight 3 | 2 | 0 |
| *Staphylococcus epidermidis* | Flight 3 | 2 | 0 |
| *Staphylococcus haemolyticus* | Flight 3 | 2 | 0 |
| *Staphylococcus saprophyticus* | Flight 3 | 2 | 0 |
| *Staphylococcus saprophyticus* | Flight 3 | 3 | 0 |
| *Staphylococcus* sp. strain LCT-H4 | Flight 3 | 2 | 0 |
| *Staphylococcus warneri* | Flight 3 | 2 | 0 |

**Table S6: Metabolic benefit provided by each family to the rest of the community**

| Family | Flight | Location | Community Support Index (X-Family\|X) (%) |
|---|---|---|---|
| *Enterobacteriaceae* | Flight 1 | 1 | 0.50 |
| *Sporidiobolaceae* | Flight 1 | 1 | 0.00 |
| *Trichocomaceae* | Flight 1 | 1 | 0.20 |
| *Enterobacteriaceae* | Flight 1 | 2 | 0.00 |
| *Erwiniaceae* | Flight 1 | 2 | 0.00 |
| *Sporidiobolaceae* | Flight 1 | 2 | 0.00 |
| *Trichocomaceae* | Flight 1 | 2 | 0.40 |
| *Enterobacteriaceae* | Flight 1 | 5 | 0.00 |
| *Erwiniaceae* | Flight 1 | 5 | 0.19 |
| *Sporidiobolaceae* | Flight 1 | 5 | 0.00 |
| *Trichocomaceae* | Flight 1 | 5 | 0.36 |
| *Enterobacteriaceae* | Flight 2 | 5 | 0.36 |
| *Erwiniaceae* | Flight 2 | 5 | 0.57 |
| *Paenibacillaceae* | Flight 2 | 5 | 0.00 |
| *Enterobacteriaceae* | Flight 3 | 1 | 0.00 |
| *Erwiniaceae* | Flight 3 | 1 | 0.41 |
| *Sporidiobolaceae* | Flight 3 | 1 | 0.00 |
| *Trichocomaceae* | Flight 3 | 1 | 0.00 |
| *Enterobacteriaceae* | Flight 3 | 2 | 0.00 |
| *Enterococcaceae* | Flight 3 | 2 | 0.00 |
| *Paenibacillaceae* | Flight 3 | 2 | 0.13 |
| *Sporidiobolaceae* | Flight 3 | 2 | 0.00 |
| *Staphylococcaceae* | Flight 3 | 2 | 0.00 |
| *Trichocomaceae* | Flight 3 | 2 | 0.10 |
| *Enterobacteriaceae* | Flight 3 | 3 | 1.40 |
| *Staphylococcaceae* | Flight 3 | 3 | 0.00 |
| *Trichocomaceae* | Flight 3 | 3 | 1.15 |
| *Enterobacteriaceae* | Flight 3 | 4 | 5.35 |
| *Erwiniaceae* | Flight 3 | 4 | 0.45 |
| *Enterobacteriaceae* | Flight 3 | 5 | 5.42 |
| *Erwiniaceae* | Flight 3 | 5 | 0.50 |
| *Enterobacteriaceae* | Flight 3 | 7 | 5.54 |
| *Erwiniaceae* | Flight 3 | 7 | 0.54 |
| *Enterobacteriaceae* | Flight 3 | 8 | 5.42 |
| *Erwiniaceae* | Flight 3 | 8 | 0.50 |

**Table S7: Metabolic support provided to an individual microorganism by the rest of the community**

| Microorganism | Flight | Location | Community Support Index (A\|X) (%) |
|---|---|---|---|
| *Aspergillus niger* | Flight 1 | 1 | 0.93 |
| *Aspergillus niger* | Flight 3 | 1 | 0.00 |
| *Enterobacter asburiae* | Flight 1 | 2 | 0.88 |
| *Enterobacter cancerogenus* | Flight 1 | 2 | 0.83 |
| *Enterobacter cloacae* | Flight 1 | 2 | 0.91 |
| *Enterobacter cloacae* | Flight 3 | 1 | 0.74 |
| *Enterobacter cloacae* | Flight 3 | 5 | 0.73 |
| *Enterobacter cloacae* | Flight 3 | 7 | 0.73 |
| *Enterobacter cloacae* | Flight 3 | 8 | 0.73 |
| *Enterobacter hormaechei* | Flight 1 | 2 | 0.86 |
| *Enterobacter hormaechei* | Flight 3 | 4 | 0.69 |
| *Enterobacter roggenkampii* | Flight 1 | 2 | 0.86 |
| *Enterobacter* sp. strain NFIX59 | Flight 1 | 2 | 0.85 |
| *Enterococcus avium* | Flight 3 | 2 | 2.45 |
| *Enterococcus faecalis* | Flight 3 | 2 | 0.22 |
| *Escherichia coli* | Flight 1 | 2 | 0.83 |
| *Escherichia coli* | Flight 1 | 5 | 0.83 |
| *Escherichia coli* | Flight 2 | 5 | 5.87 |
| *Escherichia coli* | Flight 3 | 1 | 0.68 |
| *Escherichia coli* | Flight 3 | 3 | 6.02 |
| *Escherichia coli* | Flight 3 | 4 | 5.87 |
| *Escherichia coli* | Flight 3 | 5 | 5.87 |
| *Escherichia coli* | Flight 3 | 7 | 5.87 |
| *Escherichia coli* | Flight 3 | 8 | 5.87 |
| *Klebsiella aerogenes* strain IIIF7SW-P1 | Flight 3 | 7 | 0.67 |
| *Klebsiella oxytoca* | Flight 1 | 2 | 0.15 |
| *Klebsiella pneumoniae* | Flight 1 | 1 | 0.14 |
| *Klebsiella pneumoniae* | Flight 1 | 2 | 0.14 |
| *Klebsiella pneumoniae* | Flight 1 | 5 | 0.14 |
| *Klebsiella pneumoniae* | Flight 2 | 5 | 0.00 |
| *Klebsiella pneumoniae* | Flight 3 | 1 | 0.00 |
| *Klebsiella pneumoniae* | Flight 3 | 3 | 0.69 |
| *Klebsiella pneumoniae* | Flight 3 | 4 | 0.00 |
| *Klebsiella pneumoniae* | Flight 3 | 5 | 0.00 |
| *Klebsiella pneumoniae* | Flight 3 | 7 | 0.00 |
| *Klebsiella pneumoniae* | Flight 3 | 8 | 0.00 |
| *Klebsiella pneumoniae* strain F3-2P(2*) | Flight 3 | 2 | 0.14 |
| *Klebsiella quasipneumoniae* strain IF1SW-B2 | Flight 1 | 1 | 0.14 |
| *Klebsiella quasipneumoniae* strain IF1SW-P3 | Flight 1 | 1 | 0.14 |
| *Klebsiella quasipneumoniae* strain IF1SW-P4 | Flight 1 | 1 | 0.15 |
| *Klebsiella quasipneumoniae* strain IF2SW-B3 | Flight 1 | 2 | 0.55 |
| *Klebsiella quasipneumoniae* strain IF2SW-P1 | Flight 1 | 2 | 0.55 |
| *Klebsiella quasipneumoniae* strain IIIF3SW-P1 | Flight 3 | 3 | 0.68 |
| *Klebsiella* sp. strain MS 92-3 | Flight 3 | 3 | 0.69 |
| *Klebsiella variicola* | Flight 3 | 3 | 0.68 |
| *Paenibacillus polymyxa* | Flight 3 | 2 | 0.16 |

| Microorganism | Flight | Location | Community Support Index (A\|X) (%) |
|---|---|---|---|
| *Paenibacillus polymyxa* strain IIF5SW-B3 | Flight 2 | 5 | 0.97 |
| *Paenibacillus polymyxa* strain IIF5SW-B4 | Flight 2 | 5 | 0.97 |
| *Pantoea agglomerans* | Flight 1 | 5 | 2.13 |
| *Pantoea agglomerans* | Flight 2 | 5 | 6.40 |
| *Pantoea agglomerans* | Flight 3 | 1 | 1.67 |
| *Pantoea agglomerans* | Flight 3 | 5 | 6.40 |
| *Pantoea agglomerans* | Flight 3 | 7 | 6.40 |
| *Pantoea agglomerans* | Flight 3 | 8 | 6.40 |
| *Pantoea ananatis* | Flight 1 | 2 | 0.57 |
| *Pantoea ananatis* | Flight 1 | 5 | 0.57 |
| *Pantoea ananatis* | Flight 2 | 5 | 4.95 |
| *Pantoea ananatis* | Flight 3 | 1 | 0.00 |
| *Pantoea ananatis* | Flight 3 | 4 | 4.95 |
| *Pantoea ananatis* | Flight 3 | 5 | 4.95 |
| *Pantoea ananatis* | Flight 3 | 7 | 4.95 |
| *Pantoea ananatis* | Flight 3 | 8 | 4.95 |
| *Pantoea conspicua* | Flight 1 | 5 | 1.46 |
| *Pantoea conspicua* | Flight 2 | 5 | 6.46 |
| *Pantoea conspicua* | Flight 3 | 1 | 0.95 |
| *Pantoea conspicua* | Flight 3 | 5 | 6.46 |
| *Pantoea conspicua* | Flight 3 | 7 | 6.46 |
| *Pantoea conspicua* | Flight 3 | 8 | 6.46 |
| *Pantoea dispersa* | Flight 3 | 4 | 6.05 |
| *Pantoea* sp. strain 3.5.1 | Flight 1 | 5 | 1.31 |
| *Pantoea* sp. strain 3.5.1 | Flight 2 | 5 | 5.76 |
| *Pantoea* sp. strain A4 | Flight 3 | 1 | 1.61 |
| *Pantoea* sp. strain A4 | Flight 3 | 5 | 6.19 |
| *Pantoea* sp. strain A4 | Flight 3 | 7 | 6.19 |
| *Pantoea* sp. strain A4 | Flight 3 | 8 | 6.19 |
| *Pantoea* sp. strain At-9b | Flight 3 | 1 | 0.00 |
| *Pantoea* sp. strain At-9b | Flight 3 | 5 | 4.41 |
| *Pantoea sp. strain At-9b* | Flight 3 | 8 | 4.41 |
| *Pantoea* sp. strain FF5 | Flight 3 | 1 | 1.57 |
| *Pantoea* sp. strain FF5 | Flight 3 | 5 | 6.17 |
| *Pantoea* sp. strain FF5 | Flight 3 | 7 | 6.17 |
| *Pantoea* sp. strain FF5 | Flight 3 | 8 | 6.17 |
| *Pantoea* sp. strain IMH | Flight 3 | 1 | 0.00 |
| *Pantoea* sp. strain IMH | Flight 3 | 5 | 5.39 |
| *Pantoea* sp. strain IMH | Flight 3 | 7 | 5.39 |
| *Pantoea* sp. strain IMH | Flight 3 | 8 | 5.39 |
| *Pantoea* sp. strain NGS-ED-1003 | Flight 3 | 1 | 0.85 |
| *Pantoea* sp. strain NGS-ED-1003 | Flight 3 | 5 | 5.87 |
| *Pantoea* sp. strain NGS-ED-1003 | Flight 3 | 7 | 5.87 |
| *Pantoea* sp. strain NGS-ED-1003 | Flight 3 | 8 | 5.87 |
| *Pantoea* sp. strain OXWO6B1 | Flight 3 | 1 | 0.00 |
| *Pantoea* sp. strain OXWO6B1 | Flight 3 | 5 | 4.94 |
| *Pantoea* sp. strain OXWO6B1 | Flight 3 | 7 | 4.94 |

| Microorganism | Flight | Location | Community Support Index (A\|X) (%) |
|---|---|---|---|
| *Pantoea* sp. strain OXWO6B1 | Flight 3 | 8 | 4.94 |
| *Pantoea vagans* | Flight 2 | 5 | 6.52 |
| *Pantoea vagans* | Flight 3 | 1 | 1.70 |
| *Pantoea vagans* | Flight 3 | 5 | 6.52 |
| *Pantoea vagans* | Flight 3 | 7 | 6.52 |
| *Pantoea vagans* | Flight 3 | 8 | 6.52 |
| *Penicillium chrysogenum* | Flight 1 | 1 | 0.64 |
| *Penicillium chrysogenum* | Flight 3 | 2 | 0.79 |
| *Penicillium flavigenum* | Flight 3 | 2 | 0.63 |
| *Penicillium nalgiovense* | Flight 3 | 2 | 0.78 |
| *Penicillium rubens* | Flight 1 | 1 | 0.64 |
| *Penicillium rubens* | Flight 1 | 2 | 0.64 |
| *Penicillium rubens* | Flight 1 | 5 | 0.64 |
| *Penicillium rubens* | Flight 3 | 2 | 0.78 |
| *Penicillium rubens* | Flight 3 | 3 | 0.64 |
| *Rahnella aquatilis* | Flight 3 | 4 | 5.09 |
| *Rhodotorula* sp. strain JG-1b | Flight 1 | 1 | 0.56 |
| *Rhodotorula* sp. strain JG-1b | Flight 1 | 2 | 0.56 |
| *Rhodotorula* sp. strain JG-1b | Flight 1 | 5 | 0.56 |
| *Rhodotorula* sp. strain JG-1b | Flight 3 | 1 | 0.28 |
| *Rhodotorula* sp. strain JG-1b | Flight 3 | 2 | 0.57 |
| *Rhodotorula toruloides* | Flight 1 | 1 | 1.33 |
| *Salmonella enterica* | Flight 1 | 2 | 4.12 |
| *Salmonella enterica* | Flight 3 | 1 | 0.65 |
| *Salmonella enterica* | Flight 3 | 3 | 8.79 |
| *Salmonella enterica* | Flight 3 | 5 | 5.52 |
| *Salmonella enterica* | Flight 3 | 7 | 5.52 |
| *Salmonella enterica* | Flight 3 | 8 | 5.52 |
| *Shigella sonnei* | Flight 1 | 2 | 0.67 |
| *Staphylococcus aureus* | Flight 3 | 2 | 0.78 |
| *Staphylococcus epidermidis* | Flight 3 | 2 | 0.81 |
| *Staphylococcus haemolyticus* | Flight 3 | 2 | 0.38 |
| *Staphylococcus saprophyticus* | Flight 3 | 2 | 0.19 |
| *Staphylococcus saprophyticus* | Flight 3 | 3 | 3.29 |
| *Staphylococcus* sp. strain LCT-H4 | Flight 3 | 2 | 0.88 |
| *Staphylococcus warneri* | Flight 3 | 2 | 2.20 |

**Table S8: Metabolic support provided to an individual microorganism by another microorganism in the community**

| Microorganism | In the presence of | Flight | Location | Metabolic Support Index (A\|AUB) (%) |
|---|---|---|---|---|
| *Aspergillus niger* | *Enterobacter cloacae* | Flight 3 | 1 | 0.0000 |
| *Aspergillus niger* | *Escherichia coli* | Flight 3 | 1 | 0.0000 |
| *Aspergillus niger* | *Klebsiella pneumoniae* | Flight 1 | 1 | 0.0000 |
| *Aspergillus niger* | *Klebsiella pneumoniae* | Flight 3 | 1 | 0.0000 |
| *Aspergillus niger* | *Klebsiella quasipneumoniae strain IF1SW-B2* | Flight 1 | 1 | 0.0000 |
| *Aspergillus niger* | *Klebsiella quasipneumoniae strain IF1SW-P3* | Flight 1 | 1 | 0.0000 |
| *Aspergillus niger* | *Klebsiella quasipneumoniae strain IF1SW-P4* | Flight 1 | 1 | 0.0000 |
| *Aspergillus niger* | *Pantoea agglomerans* | Flight 3 | 1 | 0.0000 |
| *Aspergillus niger* | *Pantoea ananatis* | Flight 3 | 1 | 0.0000 |
| *Aspergillus niger* | *Pantoea conspicua* | Flight 3 | 1 | 0.0000 |
| *Aspergillus niger* | *Pantoea sp. strain A4* | Flight 3 | 1 | 0.0000 |
| *Aspergillus niger* | *Pantoea sp. strain At-9b* | Flight 3 | 1 | 0.0000 |
| *Aspergillus niger* | *Pantoea sp. strain FF5* | Flight 3 | 1 | 0.0000 |
| *Aspergillus niger* | *Pantoea sp. strain IMH* | Flight 3 | 1 | 0.0000 |
| *Aspergillus niger* | *Pantoea sp. strain NGS-ED-1003* | Flight 3 | 1 | 0.0000 |
| *Aspergillus niger* | *Pantoea sp. strain OXWO6B1* | Flight 3 | 1 | 0.0000 |
| *Aspergillus niger* | *Pantoea vagans* | Flight 3 | 1 | 0.0000 |
| *Aspergillus niger* | *Penicillium chrysogenum* | Flight 1 | 1 | 0.9317 |
| *Aspergillus niger* | *Penicillium rubens* | Flight 1 | 1 | 0.9317 |
| *Aspergillus niger* | *Rhodotorula sp. strain JG-1b* | Flight 1 | 1 | 0.0000 |
| *Aspergillus niger* | *Rhodotorula sp. strain JG-1b* | Flight 3 | 1 | 0.0000 |
| *Aspergillus niger* | *Rhodotorula toruloides* | Flight 1 | 1 | 0.0000 |
| *Aspergillus niger* | *Salmonella enterica* | Flight 3 | 1 | 0.0000 |
| *Enterobacter asburiae* | *Enterobacter cancerogenus* | Flight 1 | 2 | 0.0000 |
| *Enterobacter asburiae* | *Enterobacter cloacae* | Flight 1 | 2 | 0.0000 |
| *Enterobacter asburiae* | *Enterobacter hormaechei* | Flight 1 | 2 | 0.0000 |
| *Enterobacter asburiae* | *Enterobacter roggenkampii* | Flight 1 | 2 | 0.0000 |
| *Enterobacter asburiae* | *Enterobacter sp. strain NFIX59* | Flight 1 | 2 | 0.0000 |
| *Enterobacter asburiae* | *Escherichia coli* | Flight 1 | 2 | 0.0000 |
| *Enterobacter asburiae* | *Klebsiella oxytoca* | Flight 1 | 2 | 0.7310 |
| *Enterobacter asburiae* | *Klebsiella pneumoniae* | Flight 1 | 2 | 0.0000 |
| *Enterobacter asburiae* | *Klebsiella quasipneumoniae strain IF2SW-B3* | Flight 1 | 2 | 0.0000 |
| *Enterobacter asburiae* | *Klebsiella quasipneumoniae strain IF2SW-P1* | Flight 1 | 2 | 0.0000 |
| *Enterobacter asburiae* | *Pantoea ananatis* | Flight 1 | 2 | 0.0000 |
| *Enterobacter asburiae* | *Penicillium rubens* | Flight 1 | 2 | 0.1462 |
| *Enterobacter asburiae* | *Rhodotorula sp. strain JG-1b* | Flight 1 | 2 | 0.0000 |
| *Enterobacter asburiae* | *Salmonella enterica* | Flight 1 | 2 | 0.0000 |
| *Enterobacter asburiae* | *Shigella sonnei* | Flight 1 | 2 | 0.0000 |
| *Enterobacter cancerogenus* | *Enterobacter asburiae* | Flight 1 | 2 | 0.0000 |
| *Enterobacter cancerogenus* | *Enterobacter cloacae* | Flight 1 | 2 | 0.0000 |
| *Enterobacter cancerogenus* | *Enterobacter hormaechei* | Flight 1 | 2 | 0.0000 |
| *Enterobacter cancerogenus* | *Enterobacter roggenkampii* | Flight 1 | 2 | 0.0000 |
| *Enterobacter cancerogenus* | *Enterobacter sp. strain NFIX59* | Flight 1 | 2 | 0.0000 |
| *Enterobacter cancerogenus* | *Escherichia coli* | Flight 1 | 2 | 0.0000 |
| *Enterobacter cancerogenus* | *Klebsiella oxytoca* | Flight 1 | 2 | 0.6887 |
| *Enterobacter cancerogenus* | *Klebsiella pneumoniae* | Flight 1 | 2 | 0.0000 |
| *Enterobacter cancerogenus* | *Klebsiella quasipneumoniae strain IF2SW-B3* | Flight 1 | 2 | 0.0000 |
| *Enterobacter cancerogenus* | *Klebsiella quasipneumoniae strain IF2SW-P1* | Flight 1 | 2 | 0.0000 |
| *Enterobacter cancerogenus* | *Pantoea ananatis* | Flight 1 | 2 | 0.0000 |
| *Enterobacter cancerogenus* | *Penicillium rubens* | Flight 1 | 2 | 0.1377 |
| *Enterobacter cancerogenus* | *Rhodotorula sp. strain JG-1b* | Flight 1 | 2 | 0.0000 |
| *Enterobacter cancerogenus* | *Salmonella enterica* | Flight 1 | 2 | 0.0000 |
| *Enterobacter cancerogenus* | *Shigella sonnei* | Flight 1 | 2 | 0.0000 |
| *Enterobacter cloacae* | *Aspergillus niger* | Flight 3 | 1 | 0.0000 |
| *Enterobacter cloacae* | *Enterobacter asburiae* | Flight 1 | 2 | 0.0000 |
| *Enterobacter cloacae* | *Enterobacter cancerogenus* | Flight 1 | 2 | 0.0000 |
| *Enterobacter cloacae* | *Enterobacter hormaechei* | Flight 1 | 2 | 0.0000 |
| *Enterobacter cloacae* | *Enterobacter roggenkampii* | Flight 1 | 2 | 0.0000 |
| *Enterobacter cloacae* | *Enterobacter sp. strain NFIX59* | Flight 1 | 2 | 0.0000 |
| *Enterobacter cloacae* | *Escherichia coli* | Flight 1 | 2 | 0.0000 |
| *Enterobacter cloacae* | *Escherichia coli* | Flight 3 | 1 | 0.0000 |
| *Enterobacter cloacae* | *Escherichia coli* | Flight 3 | 5 | 0.0000 |
| *Enterobacter cloacae* | *Escherichia coli* | Flight 3 | 7 | 0.0000 |
| *Enterobacter cloacae* | *Escherichia coli* | Flight 3 | 8 | 0.0000 |
| *Enterobacter cloacae* | *Klebsiella aerogenes strain IIIF7SW-P1* | Flight 3 | 7 | 0.0000 |
| *Enterobacter cloacae* | *Klebsiella oxytoca* | Flight 1 | 2 | 0.7610 |
| *Enterobacter cloacae* | *Klebsiella pneumoniae* | Flight 1 | 2 | 0.0000 |
| *Enterobacter cloacae* | *Klebsiella pneumoniae* | Flight 3 | 1 | 0.0000 |
| *Enterobacter cloacae* | *Klebsiella pneumoniae* | Flight 3 | 5 | 0.0000 |
| *Enterobacter cloacae* | *Klebsiella pneumoniae* | Flight 3 | 7 | 0.0000 |
| *Enterobacter cloacae* | *Klebsiella pneumoniae* | Flight 3 | 8 | 0.0000 |
| *Enterobacter cloacae* | *Klebsiella quasipneumoniae strain IF2SW-B3* | Flight 1 | 2 | 0.0000 |
| *Enterobacter cloacae* | *Klebsiella quasipneumoniae strain IF2SW-P1* | Flight 1 | 2 | 0.0000 |
| *Enterobacter cloacae* | *Pantoea agglomerans* | Flight 3 | 1 | 0.0000 |
| *Enterobacter cloacae* | *Pantoea agglomerans* | Flight 3 | 5 | 0.0000 |
| *Enterobacter cloacae* | *Pantoea agglomerans* | Flight 3 | 7 | 0.0000 |
| *Enterobacter cloacae* | *Pantoea agglomerans* | Flight 3 | 8 | 0.0000 |
| *Enterobacter cloacae* | *Pantoea ananatis* | Flight 1 | 2 | 0.0000 |
| *Enterobacter cloacae* | *Pantoea ananatis* | Flight 3 | 1 | 0.0000 |
| *Enterobacter cloacae* | *Pantoea ananatis* | Flight 3 | 5 | 0.0000 |
| *Enterobacter cloacae* | *Pantoea ananatis* | Flight 3 | 7 | 0.0000 |
| *Enterobacter cloacae* | *Pantoea ananatis* | Flight 3 | 8 | 0.0000 |
| *Enterobacter cloacae* | *Pantoea conspicua* | Flight 3 | 1 | 0.7386 |
| *Enterobacter cloacae* | *Pantoea conspicua* | Flight 3 | 5 | 0.7310 |
| *Enterobacter cloacae* | *Pantoea conspicua* | Flight 3 | 7 | 0.7310 |
| *Enterobacter cloacae* | *Pantoea conspicua* | Flight 3 | 8 | 0.7310 |
| *Enterobacter cloacae* | *Pantoea sp. strain A4* | Flight 3 | 1 | 0.0000 |
| *Enterobacter cloacae* | *Pantoea sp. strain A4* | Flight 3 | 5 | 0.0000 |
| *Enterobacter cloacae* | *Pantoea sp. strain A4* | Flight 3 | 7 | 0.0000 |
| *Enterobacter cloacae* | *Pantoea sp. strain A4* | Flight 3 | 8 | 0.0000 |
| *Enterobacter cloacae* | *Pantoea sp. strain At-9b* | Flight 3 | 1 | 0.0000 |
| *Enterobacter cloacae* | *Pantoea sp. strain At-9b* | Flight 3 | 5 | 0.0000 |
| *Enterobacter cloacae* | *Pantoea sp. strain At-9b* | Flight 3 | 8 | 0.0000 |
| *Enterobacter cloacae* | *Pantoea sp. strain FF5* | Flight 3 | 1 | 0.0000 |
| *Enterobacter cloacae* | *Pantoea sp. strain FF5* | Flight 3 | 5 | 0.0000 |

| Microorganism | In the presence of | Flight | Location | Metabolic Support Index (A\|AUB) (%) |
|---|---|---|---|---|
| *Enterobacter cloacae* | *Pantoea sp. strain FF5* | Flight 3 | 7 | 0.0000 |
| *Enterobacter cloacae* | *Pantoea sp. strain FF5* | Flight 3 | 8 | 0.0000 |
| *Enterobacter cloacae* | *Pantoea sp. strain IMH* | Flight 3 | 1 | 0.7386 |
| *Enterobacter cloacae* | *Pantoea sp. strain IMH* | Flight 3 | 5 | 0.7310 |
| *Enterobacter cloacae* | *Pantoea sp. strain IMH* | Flight 3 | 7 | 0.7310 |
| *Enterobacter cloacae* | *Pantoea sp. strain IMH* | Flight 3 | 8 | 0.7310 |
| *Enterobacter cloacae* | *Pantoea sp. strain NGS-ED-1003* | Flight 3 | 1 | 0.7386 |
| *Enterobacter cloacae* | *Pantoea sp. strain NGS-ED-1003* | Flight 3 | 5 | 0.7310 |
| *Enterobacter cloacae* | *Pantoea sp. strain NGS-ED-1003* | Flight 3 | 7 | 0.7310 |
| *Enterobacter cloacae* | *Pantoea sp. strain NGS-ED-1003* | Flight 3 | 8 | 0.7310 |
| *Enterobacter cloacae* | *Pantoea sp. strain OXWO6B1* | Flight 3 | 1 | 0.0000 |
| *Enterobacter cloacae* | *Pantoea sp. strain OXWO6B1* | Flight 3 | 5 | 0.0000 |
| *Enterobacter cloacae* | *Pantoea sp. strain OXWO6B1* | Flight 3 | 7 | 0.0000 |
| *Enterobacter cloacae* | *Pantoea sp. strain OXWO6B1* | Flight 3 | 8 | 0.0000 |
| *Enterobacter cloacae* | *Pantoea vagans* | Flight 3 | 1 | 0.0000 |
| *Enterobacter cloacae* | *Pantoea vagans* | Flight 3 | 5 | 0.0000 |
| *Enterobacter cloacae* | *Pantoea vagans* | Flight 3 | 7 | 0.0000 |
| *Enterobacter cloacae* | *Pantoea vagans* | Flight 3 | 8 | 0.0000 |
| *Enterobacter cloacae* | *Penicillium rubens* | Flight 1 | 2 | 0.1522 |
| *Enterobacter cloacae* | *Rhodotorula sp. strain JG-1b* | Flight 1 | 2 | 0.0000 |
| *Enterobacter cloacae* | *Rhodotorula sp. strain JG-1b* | Flight 3 | 1 | 0.0000 |
| *Enterobacter cloacae* | *Salmonella enterica* | Flight 1 | 2 | 0.0000 |
| *Enterobacter cloacae* | *Salmonella enterica* | Flight 3 | 1 | 0.0000 |
| *Enterobacter cloacae* | *Salmonella enterica* | Flight 3 | 5 | 0.0000 |
| *Enterobacter cloacae* | *Salmonella enterica* | Flight 3 | 7 | 0.0000 |
| *Enterobacter cloacae* | *Salmonella enterica* | Flight 3 | 8 | 0.0000 |
| *Enterobacter cloacae* | *Shigella sonnei* | Flight 1 | 2 | 0.0000 |
| *Enterobacter hormaechei* | *Enterobacter asburiae* | Flight 1 | 2 | 0.0000 |
| *Enterobacter hormaechei* | *Enterobacter cancerogenus* | Flight 1 | 2 | 0.0000 |
| *Enterobacter hormaechei* | *Enterobacter cloacae* | Flight 1 | 2 | 0.0000 |
| *Enterobacter hormaechei* | *Enterobacter roggenkampii* | Flight 1 | 2 | 0.0000 |
| *Enterobacter hormaechei* | *Enterobacter sp. strain NFIX59* | Flight 1 | 2 | 0.0000 |
| *Enterobacter hormaechei* | *Escherichia coli* | Flight 1 | 2 | 0.0000 |
| *Enterobacter hormaechei* | *Escherichia coli* | Flight 3 | 4 | 0.0000 |
| *Enterobacter hormaechei* | *Klebsiella oxytoca* | Flight 1 | 2 | 0.7133 |
| *Enterobacter hormaechei* | *Klebsiella pneumoniae* | Flight 1 | 2 | 0.0000 |
| *Enterobacter hormaechei* | *Klebsiella pneumoniae* | Flight 3 | 4 | 0.0000 |
| *Enterobacter hormaechei* | *Klebsiella quasipneumoniae strain IF2SW-B3* | Flight 1 | 2 | 0.0000 |
| *Enterobacter hormaechei* | *Klebsiella quasipneumoniae strain IF2SW-P1* | Flight 1 | 2 | 0.0000 |
| *Enterobacter hormaechei* | *Pantoea ananatis* | Flight 1 | 2 | 0.0000 |
| *Enterobacter hormaechei* | *Pantoea ananatis* | Flight 3 | 4 | 0.0000 |
| *Enterobacter hormaechei* | *Pantoea dispersa* | Flight 3 | 4 | 0.0000 |
| *Enterobacter hormaechei* | *Penicillium rubens* | Flight 1 | 2 | 0.1427 |
| *Enterobacter hormaechei* | *Rahnella aquatilis* | Flight 3 | 4 | 0.6897 |
| *Enterobacter hormaechei* | *Rhodotorula sp. strain JG-1b* | Flight 1 | 2 | 0.0000 |
| *Enterobacter hormaechei* | *Salmonella enterica* | Flight 1 | 2 | 0.0000 |
| *Enterobacter hormaechei* | *Shigella sonnei* | Flight 1 | 2 | 0.0000 |
| *Enterobacter roggenkampii* | *Enterobacter asburiae* | Flight 1 | 2 | 0.0000 |
| *Enterobacter roggenkampii* | *Enterobacter cancerogenus* | Flight 1 | 2 | 0.0000 |
| *Enterobacter roggenkampii* | *Enterobacter cloacae* | Flight 1 | 2 | 0.0000 |
| *Enterobacter roggenkampii* | *Enterobacter hormaechei* | Flight 1 | 2 | 0.0000 |
| *Enterobacter roggenkampii* | *Enterobacter sp. strain NFIX59* | Flight 1 | 2 | 0.0000 |
| *Enterobacter roggenkampii* | *Escherichia coli* | Flight 1 | 2 | 0.0000 |
| *Enterobacter roggenkampii* | *Klebsiella oxytoca* | Flight 1 | 2 | 0.7205 |
| *Enterobacter roggenkampii* | *Klebsiella pneumoniae* | Flight 1 | 2 | 0.0000 |
| *Enterobacter roggenkampii* | *Klebsiella quasipneumoniae strain IF2SW-B3* | Flight 1 | 2 | 0.0000 |
| *Enterobacter roggenkampii* | *Klebsiella quasipneumoniae strain IF2SW-P1* | Flight 1 | 2 | 0.0000 |
| *Enterobacter roggenkampii* | *Pantoea ananatis* | Flight 1 | 2 | 0.0000 |
| *Enterobacter roggenkampii* | *Penicillium rubens* | Flight 1 | 2 | 0.1441 |
| *Enterobacter roggenkampii* | *Rhodotorula sp. strain JG-1b* | Flight 1 | 2 | 0.0000 |
| *Enterobacter roggenkampii* | *Salmonella enterica* | Flight 1 | 2 | 0.0000 |
| *Enterobacter roggenkampii* | *Shigella sonnei* | Flight 1 | 2 | 0.0000 |
| *Enterobacter sp. strain NFIX59* | *Enterobacter asburiae* | Flight 1 | 2 | 0.0000 |
| *Enterobacter sp. strain NFIX59* | *Enterobacter cancerogenus* | Flight 1 | 2 | 0.0000 |
| *Enterobacter sp. strain NFIX59* | *Enterobacter cloacae* | Flight 1 | 2 | 0.0000 |
| *Enterobacter sp. strain NFIX59* | *Enterobacter hormaechei* | Flight 1 | 2 | 0.0000 |
| *Enterobacter sp. strain NFIX59* | *Enterobacter roggenkampii* | Flight 1 | 2 | 0.0000 |
| *Enterobacter sp. strain NFIX59* | *Escherichia coli* | Flight 1 | 2 | 0.0000 |
| *Enterobacter sp. strain NFIX59* | *Klebsiella oxytoca* | Flight 1 | 2 | 0.7082 |
| *Enterobacter sp. strain NFIX59* | *Klebsiella pneumoniae* | Flight 1 | 2 | 0.0000 |
| *Enterobacter sp. strain NFIX59* | *Klebsiella quasipneumoniae strain IF2SW-B3* | Flight 1 | 2 | 0.0000 |
| *Enterobacter sp. strain NFIX59* | *Klebsiella quasipneumoniae strain IF2SW-P1* | Flight 1 | 2 | 0.0000 |
| *Enterobacter sp. strain NFIX59* | *Pantoea ananatis* | Flight 1 | 2 | 0.0000 |
| *Enterobacter sp. strain NFIX59* | *Penicillium rubens* | Flight 1 | 2 | 0.1416 |
| *Enterobacter sp. strain NFIX59* | *Rhodotorula sp. strain JG-1b* | Flight 1 | 2 | 0.0000 |
| *Enterobacter sp. strain NFIX59* | *Salmonella enterica* | Flight 1 | 2 | 0.0000 |
| *Enterobacter sp. strain NFIX59* | *Shigella sonnei* | Flight 1 | 2 | 0.0000 |
| *Enterococcus avium* | *Enterococcus faecalis* | Flight 3 | 2 | 0.0000 |
| *Enterococcus avium* | *Klebsiella pneumoniae strain F3-2P(2*)* | Flight 3 | 2 | 0.2045 |
| *Enterococcus avium* | *Paenibacillus polymyxa* | Flight 3 | 2 | 2.2495 |
| *Enterococcus avium* | *Penicillium chrysogenum* | Flight 3 | 2 | 0.2045 |
| *Enterococcus avium* | *Penicillium flavigenum* | Flight 3 | 2 | 0.2045 |
| *Enterococcus avium* | *Penicillium nalgiovense* | Flight 3 | 2 | 0.2045 |
| *Enterococcus avium* | *Penicillium rubens* | Flight 3 | 2 | 0.2045 |
| *Enterococcus avium* | *Rhodotorula sp. strain JG-1b* | Flight 3 | 2 | 0.0000 |
| *Enterococcus avium* | *Staphylococcus aureus* | Flight 3 | 2 | 0.0000 |
| *Enterococcus avium* | *Staphylococcus epidermidis* | Flight 3 | 2 | 0.0000 |
| *Enterococcus avium* | *Staphylococcus haemolyticus* | Flight 3 | 2 | 0.0000 |
| *Enterococcus avium* | *Staphylococcus saprophyticus* | Flight 3 | 2 | 0.0000 |
| *Enterococcus avium* | *Staphylococcus sp. strain LCT-H4* | Flight 3 | 2 | 0.2045 |
| *Enterococcus avium* | *Staphylococcus warneri* | Flight 3 | 2 | 0.0000 |
| *Enterococcus faecalis* | *Enterococcus avium* | Flight 3 | 2 | 0.2169 |
| *Enterococcus faecalis* | *Klebsiella pneumoniae strain F3-2P(2*)* | Flight 3 | 2 | 0.2169 |
| *Enterococcus faecalis* | *Paenibacillus polymyxa* | Flight 3 | 2 | 0.2169 |
| *Enterococcus faecalis* | *Penicillium chrysogenum* | Flight 3 | 2 | 0.0000 |
| *Enterococcus faecalis* | *Penicillium flavigenum* | Flight 3 | 2 | 0.0000 |

| Microorganism | In the presence of | Flight | Location | Metabolic Support Index (A\|AUB) (%) |
|---|---|---|---|---|
| *Enterococcus faecalis* | *Penicillium nalgiovense* | Flight 3 | 2 | 0.0000 |
| *Enterococcus faecalis* | *Penicillium rubens* | Flight 3 | 2 | 0.0000 |
| *Enterococcus faecalis* | *Rhodotorula sp. strain JG-1b* | Flight 3 | 2 | 0.0000 |
| *Enterococcus faecalis* | *Staphylococcus aureus* | Flight 3 | 2 | 0.2169 |
| *Enterococcus faecalis* | *Staphylococcus epidermidis* | Flight 3 | 2 | 0.0000 |
| *Enterococcus faecalis* | *Staphylococcus haemolyticus* | Flight 3 | 2 | 0.2169 |
| *Enterococcus faecalis* | *Staphylococcus saprophyticus* | Flight 3 | 2 | 0.0000 |
| *Enterococcus faecalis* | *Staphylococcus sp. strain LCT-H4* | Flight 3 | 2 | 0.2169 |
| *Enterococcus faecalis* | *Staphylococcus warneri* | Flight 3 | 2 | 0.2169 |
| *Escherichia coli* | *Aspergillus niger* | Flight 3 | 1 | 0.0000 |
| *Escherichia coli* | *Enterobacter asburiae* | Flight 1 | 2 | 0.0000 |
| *Escherichia coli* | *Enterobacter cancerogenus* | Flight 1 | 2 | 0.0000 |
| *Escherichia coli* | *Enterobacter cloacae* | Flight 1 | 2 | 0.0000 |
| *Escherichia coli* | *Enterobacter cloacae* | Flight 3 | 1 | 0.0000 |
| *Escherichia coli* | *Enterobacter cloacae* | Flight 3 | 5 | 5.2296 |
| *Escherichia coli* | *Enterobacter cloacae* | Flight 3 | 7 | 5.2296 |
| *Escherichia coli* | *Enterobacter cloacae* | Flight 3 | 8 | 5.2296 |
| *Escherichia coli* | *Enterobacter hormaechei* | Flight 1 | 2 | 0.0000 |
| *Escherichia coli* | *Enterobacter hormaechei* | Flight 3 | 4 | 5.2296 |
| *Escherichia coli* | *Enterobacter roggenkampii* | Flight 1 | 2 | 0.0000 |
| *Escherichia coli* | *Enterobacter sp. strain NFIX59* | Flight 1 | 2 | 0.0000 |
| *Escherichia coli* | *Klebsiella aerogenes strain IIIF7SW-P1* | Flight 3 | 7 | 5.2296 |
| *Escherichia coli* | *Klebsiella oxytoca* | Flight 1 | 2 | 0.6954 |
| *Escherichia coli* | *Klebsiella pneumoniae* | Flight 1 | 2 | 0.0000 |
| *Escherichia coli* | *Klebsiella pneumoniae* | Flight 1 | 5 | 0.0000 |
| *Escherichia coli* | *Klebsiella pneumoniae* | Flight 2 | 5 | 5.2296 |
| *Escherichia coli* | *Klebsiella pneumoniae* | Flight 3 | 1 | 0.0000 |
| *Escherichia coli* | *Klebsiella pneumoniae* | Flight 3 | 3 | 5.3665 |
| *Escherichia coli* | *Klebsiella pneumoniae* | Flight 3 | 4 | 5.2296 |
| *Escherichia coli* | *Klebsiella pneumoniae* | Flight 3 | 5 | 5.2296 |
| *Escherichia coli* | *Klebsiella pneumoniae* | Flight 3 | 7 | 5.2296 |
| *Escherichia coli* | *Klebsiella pneumoniae* | Flight 3 | 8 | 5.2296 |
| *Escherichia coli* | *Klebsiella quasipneumoniae strain IF2SW-B3* | Flight 1 | 2 | 0.0000 |
| *Escherichia coli* | *Klebsiella quasipneumoniae strain IF2SW-P1* | Flight 1 | 2 | 0.0000 |
| *Escherichia coli* | *Klebsiella quasipneumoniae strain IIIF3SW-P1* | Flight 3 | 3 | 5.3665 |
| *Escherichia coli* | *Klebsiella sp. strain MS 92-3* | Flight 3 | 3 | 5.3665 |
| *Escherichia coli* | *Klebsiella variicola* | Flight 3 | 3 | 5.3665 |
| *Escherichia coli* | *Paenibacillus polymyxa strain IIF5SW-B3* | Flight 2 | 5 | 5.2296 |
| *Escherichia coli* | *Paenibacillus polymyxa strain IIF5SW-B4* | Flight 2 | 5 | 5.2296 |
| *Escherichia coli* | *Pantoea agglomerans* | Flight 1 | 5 | 0.0000 |
| *Escherichia coli* | *Pantoea agglomerans* | Flight 2 | 5 | 0.0000 |
| *Escherichia coli* | *Pantoea agglomerans* | Flight 3 | 1 | 0.0000 |
| *Escherichia coli* | *Pantoea agglomerans* | Flight 3 | 5 | 0.0000 |
| *Escherichia coli* | *Pantoea agglomerans* | Flight 3 | 7 | 0.0000 |
| *Escherichia coli* | *Pantoea agglomerans* | Flight 3 | 8 | 0.0000 |
| *Escherichia coli* | *Pantoea ananatis* | Flight 1 | 2 | 0.0000 |
| *Escherichia coli* | *Pantoea ananatis* | Flight 1 | 5 | 0.0000 |
| *Escherichia coli* | *Pantoea ananatis* | Flight 2 | 5 | 0.0000 |
| *Escherichia coli* | *Pantoea ananatis* | Flight 3 | 1 | 0.0000 |
| *Escherichia coli* | *Pantoea ananatis* | Flight 3 | 4 | 0.0000 |
| *Escherichia coli* | *Pantoea ananatis* | Flight 3 | 5 | 0.0000 |
| *Escherichia coli* | *Pantoea ananatis* | Flight 3 | 7 | 0.0000 |
| *Escherichia coli* | *Pantoea ananatis* | Flight 3 | 8 | 0.0000 |
| *Escherichia coli* | *Pantoea conspicua* | Flight 1 | 5 | 0.6954 |
| *Escherichia coli* | *Pantoea conspicua* | Flight 2 | 5 | 0.6378 |
| *Escherichia coli* | *Pantoea conspicua* | Flight 3 | 1 | 0.6766 |
| *Escherichia coli* | *Pantoea conspicua* | Flight 3 | 5 | 0.6378 |
| *Escherichia coli* | *Pantoea conspicua* | Flight 3 | 7 | 0.6378 |
| *Escherichia coli* | *Pantoea conspicua* | Flight 3 | 8 | 0.6378 |
| *Escherichia coli* | *Pantoea dispersa* | Flight 3 | 4 | 0.0000 |
| *Escherichia coli* | *Pantoea sp. strain 3.5.1* | Flight 1 | 5 | 0.6954 |
| *Escherichia coli* | *Pantoea sp. strain 3.5.1* | Flight 2 | 5 | 0.6378 |
| *Escherichia coli* | *Pantoea sp. strain A4* | Flight 3 | 1 | 0.0000 |
| *Escherichia coli* | *Pantoea sp. strain A4* | Flight 3 | 5 | 0.0000 |
| *Escherichia coli* | *Pantoea sp. strain A4* | Flight 3 | 7 | 0.0000 |
| *Escherichia coli* | *Pantoea sp. strain A4* | Flight 3 | 8 | 0.0000 |
| *Escherichia coli* | *Pantoea sp. strain At-9b* | Flight 3 | 1 | 0.0000 |
| *Escherichia coli* | *Pantoea sp. strain At-9b* | Flight 3 | 5 | 0.0000 |
| *Escherichia coli* | *Pantoea sp. strain At-9b* | Flight 3 | 8 | 0.0000 |
| *Escherichia coli* | *Pantoea sp. strain FF5* | Flight 3 | 1 | 0.0000 |
| *Escherichia coli* | *Pantoea sp. strain FF5* | Flight 3 | 5 | 0.0000 |
| *Escherichia coli* | *Pantoea sp. strain FF5* | Flight 3 | 7 | 0.0000 |
| *Escherichia coli* | *Pantoea sp. strain FF5* | Flight 3 | 8 | 0.0000 |
| *Escherichia coli* | *Pantoea sp. strain IMH* | Flight 3 | 1 | 0.6766 |
| *Escherichia coli* | *Pantoea sp. strain IMH* | Flight 3 | 5 | 0.6378 |
| *Escherichia coli* | *Pantoea sp. strain IMH* | Flight 3 | 7 | 0.6378 |
| *Escherichia coli* | *Pantoea sp. strain IMH* | Flight 3 | 8 | 0.6378 |
| *Escherichia coli* | *Pantoea sp. strain NGS-ED-1003* | Flight 3 | 1 | 0.6766 |
| *Escherichia coli* | *Pantoea sp. strain NGS-ED-1003* | Flight 3 | 5 | 0.6378 |
| *Escherichia coli* | *Pantoea sp. strain NGS-ED-1003* | Flight 3 | 7 | 0.6378 |
| *Escherichia coli* | *Pantoea sp. strain NGS-ED-1003* | Flight 3 | 8 | 0.6378 |
| *Escherichia coli* | *Pantoea sp. strain OXWO6B1* | Flight 3 | 1 | 0.0000 |
| *Escherichia coli* | *Pantoea sp. strain OXWO6B1* | Flight 3 | 5 | 0.0000 |
| *Escherichia coli* | *Pantoea sp. strain OXWO6B1* | Flight 3 | 7 | 0.0000 |
| *Escherichia coli* | *Pantoea sp. strain OXWO6B1* | Flight 3 | 8 | 0.0000 |
| *Escherichia coli* | *Pantoea vagans* | Flight 2 | 5 | 0.0000 |
| *Escherichia coli* | *Pantoea vagans* | Flight 3 | 1 | 0.0000 |
| *Escherichia coli* | *Pantoea vagans* | Flight 3 | 5 | 0.0000 |
| *Escherichia coli* | *Pantoea vagans* | Flight 3 | 7 | 0.0000 |
| *Escherichia coli* | *Pantoea vagans* | Flight 3 | 8 | 0.0000 |
| *Escherichia coli* | *Penicillium rubens* | Flight 1 | 2 | 0.1391 |
| *Escherichia coli* | *Penicillium rubens* | Flight 1 | 5 | 0.1391 |
| *Escherichia coli* | *Penicillium rubens* | Flight 3 | 3 | 6.0209 |
| *Escherichia coli* | *Rahnella aquatilis* | Flight 3 | 4 | 0.6378 |
| *Escherichia coli* | *Rhodotorula sp. strain JG-1b* | Flight 1 | 2 | 0.0000 |
| *Escherichia coli* | *Rhodotorula sp. strain JG-1b* | Flight 1 | 5 | 0.0000 |

| Microorganism | In the presence of | Flight | Location | Metabolic Support Index (A\|AUB) (%) |
|---|---|---|---|---|
| *Escherichia coli* | *Rhodotorula sp. strain JG-1b* | Flight 3 | 1 | 0.0000 |
| *Escherichia coli* | *Salmonella enterica* | Flight 1 | 2 | 0.0000 |
| *Escherichia coli* | *Salmonella enterica* | Flight 3 | 1 | 0.0000 |
| *Escherichia coli* | *Salmonella enterica* | Flight 3 | 3 | 0.0000 |
| *Escherichia coli* | *Salmonella enterica* | Flight 3 | 5 | 0.0000 |
| *Escherichia coli* | *Salmonella enterica* | Flight 3 | 7 | 0.0000 |
| *Escherichia coli* | *Salmonella enterica* | Flight 3 | 8 | 0.0000 |
| *Escherichia coli* | *Shigella sonnei* | Flight 1 | 2 | 0.0000 |
| *Escherichia coli* | *Staphylococcus saprophyticus* | Flight 3 | 3 | 5.3665 |
| *Klebsiella aerogenes strain IIIF7SW-P1* | *Enterobacter cloacae* | Flight 3 | 7 | 0.0000 |
| *Klebsiella aerogenes strain IIIF7SW-P1* | *Escherichia coli* | Flight 3 | 7 | 0.0000 |
| *Klebsiella aerogenes strain IIIF7SW-P1* | *Klebsiella pneumoniae* | Flight 3 | 7 | 0.0000 |
| *Klebsiella aerogenes strain IIIF7SW-P1* | *Pantoea agglomerans* | Flight 3 | 7 | 0.0000 |
| *Klebsiella aerogenes strain IIIF7SW-P1* | *Pantoea ananatis* | Flight 3 | 7 | 0.0000 |
| *Klebsiella aerogenes strain IIIF7SW-P1* | *Pantoea conspicua* | Flight 3 | 7 | 0.6684 |
| *Klebsiella aerogenes strain IIIF7SW-P1* | *Pantoea sp. strain A4* | Flight 3 | 7 | 0.0000 |
| *Klebsiella aerogenes strain IIIF7SW-P1* | *Pantoea sp. strain FF5* | Flight 3 | 7 | 0.0000 |
| *Klebsiella aerogenes strain IIIF7SW-P1* | *Pantoea sp. strain IMH* | Flight 3 | 7 | 0.6684 |
| *Klebsiella aerogenes strain IIIF7SW-P1* | *Pantoea sp. strain NGS-ED-1003* | Flight 3 | 7 | 0.6684 |
| *Klebsiella aerogenes strain IIIF7SW-P1* | *Pantoea sp. strain OXWO6B1* | Flight 3 | 7 | 0.0000 |
| *Klebsiella aerogenes strain IIIF7SW-P1* | *Pantoea vagans* | Flight 3 | 7 | 0.0000 |
| *Klebsiella aerogenes strain IIIF7SW-P1* | *Salmonella enterica* | Flight 3 | 7 | 0.0000 |
| *Klebsiella oxytoca* | *Enterobacter asburiae* | Flight 1 | 2 | 0.0000 |
| *Klebsiella oxytoca* | *Enterobacter cancerogenus* | Flight 1 | 2 | 0.0000 |
| *Klebsiella oxytoca* | *Enterobacter cloacae* | Flight 1 | 2 | 0.0000 |
| *Klebsiella oxytoca* | *Enterobacter hormaechei* | Flight 1 | 2 | 0.0000 |
| *Klebsiella oxytoca* | *Enterobacter roggenkampii* | Flight 1 | 2 | 0.0000 |
| *Klebsiella oxytoca* | *Enterobacter sp. strain NFIX59* | Flight 1 | 2 | 0.0000 |
| *Klebsiella oxytoca* | *Escherichia coli* | Flight 1 | 2 | 0.0000 |
| *Klebsiella oxytoca* | *Klebsiella pneumoniae* | Flight 1 | 2 | 0.0000 |
| *Klebsiella oxytoca* | *Klebsiella quasipneumoniae strain IF2SW-B3* | Flight 1 | 2 | 0.0000 |
| *Klebsiella oxytoca* | *Klebsiella quasipneumoniae strain IF2SW-P1* | Flight 1 | 2 | 0.0000 |
| *Klebsiella oxytoca* | *Pantoea ananatis* | Flight 1 | 2 | 0.0000 |
| *Klebsiella oxytoca* | *Penicillium rubens* | Flight 1 | 2 | 0.1477 |
| *Klebsiella oxytoca* | *Rhodotorula sp. strain JG-1b* | Flight 1 | 2 | 0.0000 |
| *Klebsiella oxytoca* | *Salmonella enterica* | Flight 1 | 2 | 0.0000 |
| *Klebsiella oxytoca* | *Shigella sonnei* | Flight 1 | 2 | 0.0000 |
| *Klebsiella pneumoniae* | *Aspergillus niger* | Flight 1 | 1 | 0.0000 |
| *Klebsiella pneumoniae* | *Aspergillus niger* | Flight 3 | 1 | 0.0000 |
| *Klebsiella pneumoniae* | *Enterobacter asburiae* | Flight 1 | 2 | 0.0000 |
| *Klebsiella pneumoniae* | *Enterobacter cancerogenus* | Flight 1 | 2 | 0.0000 |
| *Klebsiella pneumoniae* | *Enterobacter cloacae* | Flight 1 | 2 | 0.0000 |
| *Klebsiella pneumoniae* | *Enterobacter cloacae* | Flight 3 | 1 | 0.0000 |
| *Klebsiella pneumoniae* | *Enterobacter cloacae* | Flight 3 | 5 | 0.0000 |
| *Klebsiella pneumoniae* | *Enterobacter cloacae* | Flight 3 | 7 | 0.0000 |
| *Klebsiella pneumoniae* | *Enterobacter cloacae* | Flight 3 | 8 | 0.0000 |
| *Klebsiella pneumoniae* | *Enterobacter hormaechei* | Flight 1 | 2 | 0.0000 |
| *Klebsiella pneumoniae* | *Enterobacter hormaechei* | Flight 3 | 4 | 0.0000 |
| *Klebsiella pneumoniae* | *Enterobacter roggenkampii* | Flight 1 | 2 | 0.0000 |
| *Klebsiella pneumoniae* | *Enterobacter sp. strain NFIX59* | Flight 1 | 2 | 0.0000 |
| *Klebsiella pneumoniae* | *Escherichia coli* | Flight 1 | 2 | 0.0000 |
| *Klebsiella pneumoniae* | *Escherichia coli* | Flight 1 | 5 | 0.0000 |
| *Klebsiella pneumoniae* | *Escherichia coli* | Flight 2 | 5 | 0.0000 |
| *Klebsiella pneumoniae* | *Escherichia coli* | Flight 3 | 1 | 0.0000 |
| *Klebsiella pneumoniae* | *Escherichia coli* | Flight 3 | 3 | 0.0000 |
| *Klebsiella pneumoniae* | *Escherichia coli* | Flight 3 | 4 | 0.0000 |
| *Klebsiella pneumoniae* | *Escherichia coli* | Flight 3 | 5 | 0.0000 |
| *Klebsiella pneumoniae* | *Escherichia coli* | Flight 3 | 7 | 0.0000 |
| *Klebsiella pneumoniae* | *Escherichia coli* | Flight 3 | 8 | 0.0000 |
| *Klebsiella pneumoniae* | *Klebsiella aerogenes strain IIIF7SW-P1* | Flight 3 | 7 | 0.0000 |
| *Klebsiella pneumoniae* | *Klebsiella oxytoca* | Flight 1 | 2 | 0.0000 |
| *Klebsiella pneumoniae* | *Klebsiella quasipneumoniae strain IF1SW-B2* | Flight 1 | 1 | 0.0000 |
| *Klebsiella pneumoniae* | *Klebsiella quasipneumoniae strain IF1SW-P3* | Flight 1 | 1 | 0.0000 |
| *Klebsiella pneumoniae* | *Klebsiella quasipneumoniae strain IF1SW-P4* | Flight 1 | 1 | 0.0000 |
| *Klebsiella pneumoniae* | *Klebsiella quasipneumoniae strain IF2SW-B3* | Flight 1 | 2 | 0.0000 |
| *Klebsiella pneumoniae* | *Klebsiella quasipneumoniae strain IF2SW-P1* | Flight 1 | 2 | 0.0000 |
| *Klebsiella pneumoniae* | *Klebsiella quasipneumoniae strain IIIF3SW-P1* | Flight 3 | 3 | 0.0000 |
| *Klebsiella pneumoniae* | *Klebsiella sp. strain MS 92-3* | Flight 3 | 3 | 0.0000 |
| *Klebsiella pneumoniae* | *Klebsiella variicola* | Flight 3 | 3 | 0.0000 |
| *Klebsiella pneumoniae* | *Paenibacillus polymyxa strain IIF5SW-B3* | Flight 2 | 5 | 0.0000 |
| *Klebsiella pneumoniae* | *Paenibacillus polymyxa strain IIF5SW-B4* | Flight 2 | 5 | 0.0000 |
| *Klebsiella pneumoniae* | *Pantoea agglomerans* | Flight 1 | 5 | 0.0000 |
| *Klebsiella pneumoniae* | *Pantoea agglomerans* | Flight 2 | 5 | 0.0000 |
| *Klebsiella pneumoniae* | *Pantoea agglomerans* | Flight 3 | 1 | 0.0000 |
| *Klebsiella pneumoniae* | *Pantoea agglomerans* | Flight 3 | 5 | 0.0000 |
| *Klebsiella pneumoniae* | *Pantoea agglomerans* | Flight 3 | 7 | 0.0000 |
| *Klebsiella pneumoniae* | *Pantoea agglomerans* | Flight 3 | 8 | 0.0000 |
| *Klebsiella pneumoniae* | *Pantoea ananatis* | Flight 1 | 2 | 0.0000 |
| *Klebsiella pneumoniae* | *Pantoea ananatis* | Flight 1 | 5 | 0.0000 |
| *Klebsiella pneumoniae* | *Pantoea ananatis* | Flight 2 | 5 | 0.0000 |
| *Klebsiella pneumoniae* | *Pantoea ananatis* | Flight 3 | 1 | 0.0000 |
| *Klebsiella pneumoniae* | *Pantoea ananatis* | Flight 3 | 4 | 0.0000 |
| *Klebsiella pneumoniae* | *Pantoea ananatis* | Flight 3 | 5 | 0.0000 |
| *Klebsiella pneumoniae* | *Pantoea ananatis* | Flight 3 | 7 | 0.0000 |
| *Klebsiella pneumoniae* | *Pantoea ananatis* | Flight 3 | 8 | 0.0000 |
| *Klebsiella pneumoniae* | *Pantoea conspicua* | Flight 1 | 5 | 0.0000 |
| *Klebsiella pneumoniae* | *Pantoea conspicua* | Flight 2 | 5 | 0.0000 |
| *Klebsiella pneumoniae* | *Pantoea conspicua* | Flight 3 | 1 | 0.0000 |
| *Klebsiella pneumoniae* | *Pantoea conspicua* | Flight 3 | 5 | 0.0000 |
| *Klebsiella pneumoniae* | *Pantoea conspicua* | Flight 3 | 7 | 0.0000 |
| *Klebsiella pneumoniae* | *Pantoea conspicua* | Flight 3 | 8 | 0.0000 |
| *Klebsiella pneumoniae* | *Pantoea dispersa* | Flight 3 | 4 | 0.0000 |
| *Klebsiella pneumoniae* | *Pantoea sp. strain 3.5.1* | Flight 1 | 5 | 0.0000 |
| *Klebsiella pneumoniae* | *Pantoea sp. strain 3.5.1* | Flight 2 | 5 | 0.0000 |
| *Klebsiella pneumoniae* | *Pantoea sp. strain A4* | Flight 3 | 1 | 0.0000 |
| *Klebsiella pneumoniae* | *Pantoea sp. strain A4* | Flight 3 | 5 | 0.0000 |

| Microorganism | In the presence of | Flight | Location | Metabolic Support Index (A\|AUB) (%) |
|---|---|---|---|---|
| *Klebsiella pneumoniae* | *Pantoea sp. strain A4* | Flight 3 | 7 | 0.0000 |
| *Klebsiella pneumoniae* | *Pantoea sp. strain A4* | Flight 3 | 8 | 0.0000 |
| *Klebsiella pneumoniae* | *Pantoea sp. strain At-9b* | Flight 3 | 1 | 0.0000 |
| *Klebsiella pneumoniae* | *Pantoea sp. strain At-9b* | Flight 3 | 5 | 0.0000 |
| *Klebsiella pneumoniae* | *Pantoea sp. strain At-9b* | Flight 3 | 8 | 0.0000 |
| *Klebsiella pneumoniae* | *Pantoea sp. strain FF5* | Flight 3 | 1 | 0.0000 |
| *Klebsiella pneumoniae* | *Pantoea sp. strain FF5* | Flight 3 | 5 | 0.0000 |
| *Klebsiella pneumoniae* | *Pantoea sp. strain FF5* | Flight 3 | 7 | 0.0000 |
| *Klebsiella pneumoniae* | *Pantoea sp. strain FF5* | Flight 3 | 8 | 0.0000 |
| *Klebsiella pneumoniae* | *Pantoea sp. strain IMH* | Flight 3 | 1 | 0.0000 |
| *Klebsiella pneumoniae* | *Pantoea sp. strain IMH* | Flight 3 | 5 | 0.0000 |
| *Klebsiella pneumoniae* | *Pantoea sp. strain IMH* | Flight 3 | 7 | 0.0000 |
| *Klebsiella pneumoniae* | *Pantoea sp. strain IMH* | Flight 3 | 8 | 0.0000 |
| *Klebsiella pneumoniae* | *Pantoea sp. strain NGS-ED-1003* | Flight 3 | 1 | 0.0000 |
| *Klebsiella pneumoniae* | *Pantoea sp. strain NGS-ED-1003* | Flight 3 | 5 | 0.0000 |
| *Klebsiella pneumoniae* | *Pantoea sp. strain NGS-ED-1003* | Flight 3 | 7 | 0.0000 |
| *Klebsiella pneumoniae* | *Pantoea sp. strain NGS-ED-1003* | Flight 3 | 8 | 0.0000 |
| *Klebsiella pneumoniae* | *Pantoea sp. strain OXWO6B1* | Flight 3 | 1 | 0.0000 |
| *Klebsiella pneumoniae* | *Pantoea sp. strain OXWO6B1* | Flight 3 | 5 | 0.0000 |
| *Klebsiella pneumoniae* | *Pantoea sp. strain OXWO6B1* | Flight 3 | 7 | 0.0000 |
| *Klebsiella pneumoniae* | *Pantoea sp. strain OXWO6B1* | Flight 3 | 8 | 0.0000 |
| *Klebsiella pneumoniae* | *Pantoea vagans* | Flight 2 | 5 | 0.0000 |
| *Klebsiella pneumoniae* | *Pantoea vagans* | Flight 3 | 1 | 0.0000 |
| *Klebsiella pneumoniae* | *Pantoea vagans* | Flight 3 | 5 | 0.0000 |
| *Klebsiella pneumoniae* | *Pantoea vagans* | Flight 3 | 7 | 0.0000 |
| *Klebsiella pneumoniae* | *Pantoea vagans* | Flight 3 | 8 | 0.0000 |
| *Klebsiella pneumoniae* | *Penicillium chrysogenum* | Flight 1 | 1 | 0.1393 |
| *Klebsiella pneumoniae* | *Penicillium rubens* | Flight 1 | 1 | 0.1393 |
| *Klebsiella pneumoniae* | *Penicillium rubens* | Flight 1 | 2 | 0.1389 |
| *Klebsiella pneumoniae* | *Penicillium rubens* | Flight 1 | 5 | 0.1389 |
| *Klebsiella pneumoniae* | *Penicillium rubens* | Flight 3 | 3 | 0.6878 |
| *Klebsiella pneumoniae* | *Rahnella aquatilis* | Flight 3 | 4 | 0.0000 |
| *Klebsiella pneumoniae* | *Rhodotorula sp. strain JG-1b* | Flight 1 | 1 | 0.0000 |
| *Klebsiella pneumoniae* | *Rhodotorula sp. strain JG-1b* | Flight 1 | 2 | 0.0000 |
| *Klebsiella pneumoniae* | *Rhodotorula sp. strain JG-1b* | Flight 1 | 5 | 0.0000 |
| *Klebsiella pneumoniae* | *Rhodotorula sp. strain JG-1b* | Flight 3 | 1 | 0.0000 |
| *Klebsiella pneumoniae* | *Rhodotorula toruloides* | Flight 1 | 1 | 0.0000 |
| *Klebsiella pneumoniae* | *Salmonella enterica* | Flight 1 | 2 | 0.0000 |
| *Klebsiella pneumoniae* | *Salmonella enterica* | Flight 3 | 1 | 0.0000 |
| *Klebsiella pneumoniae* | *Salmonella enterica* | Flight 3 | 3 | 0.0000 |
| *Klebsiella pneumoniae* | *Salmonella enterica* | Flight 3 | 5 | 0.0000 |
| *Klebsiella pneumoniae* | *Salmonella enterica* | Flight 3 | 7 | 0.0000 |
| *Klebsiella pneumoniae* | *Salmonella enterica* | Flight 3 | 8 | 0.0000 |
| *Klebsiella pneumoniae* | *Shigella sonnei* | Flight 1 | 2 | 0.0000 |
| *Klebsiella pneumoniae* | *Staphylococcus saprophyticus* | Flight 3 | 3 | 0.0000 |
| *Klebsiella pneumoniae strain F3-2P(2*)* | *Enterococcus avium* | Flight 3 | 2 | 0.0000 |
| *Klebsiella pneumoniae strain F3-2P(2*)* | *Enterococcus faecalis* | Flight 3 | 2 | 0.0000 |
| *Klebsiella pneumoniae strain F3-2P(2*)* | *Paenibacillus polymyxa* | Flight 3 | 2 | 0.0000 |
| *Klebsiella pneumoniae strain F3-2P(2*)* | *Penicillium chrysogenum* | Flight 3 | 2 | 0.1395 |
| *Klebsiella pneumoniae strain F3-2P(2*)* | *Penicillium flavigenum* | Flight 3 | 2 | 0.1395 |
| *Klebsiella pneumoniae strain F3-2P(2*)* | *Penicillium nalgiovense* | Flight 3 | 2 | 0.1395 |
| *Klebsiella pneumoniae strain F3-2P(2*)* | *Penicillium rubens* | Flight 3 | 2 | 0.1395 |
| *Klebsiella pneumoniae strain F3-2P(2*)* | *Rhodotorula sp. strain JG-1b* | Flight 3 | 2 | 0.0000 |
| *Klebsiella pneumoniae strain F3-2P(2*)* | *Staphylococcus aureus* | Flight 3 | 2 | 0.0000 |
| *Klebsiella pneumoniae strain F3-2P(2*)* | *Staphylococcus epidermidis* | Flight 3 | 2 | 0.0000 |
| *Klebsiella pneumoniae strain F3-2P(2*)* | *Staphylococcus haemolyticus* | Flight 3 | 2 | 0.0000 |
| *Klebsiella pneumoniae strain F3-2P(2*)* | *Staphylococcus saprophyticus* | Flight 3 | 2 | 0.0000 |
| *Klebsiella pneumoniae strain F3-2P(2*)* | *Staphylococcus sp. strain LCT-H4* | Flight 3 | 2 | 0.0000 |
| *Klebsiella pneumoniae strain F3-2P(2*)* | *Staphylococcus warneri* | Flight 3 | 2 | 0.0000 |
| *Klebsiella quasipneumoniae strain IF1SW-B2* | *Aspergillus niger* | Flight 1 | 1 | 0.0000 |
| *Klebsiella quasipneumoniae strain IF1SW-B2* | *Klebsiella pneumoniae* | Flight 1 | 1 | 0.0000 |
| *Klebsiella quasipneumoniae strain IF1SW-B2* | *Klebsiella quasipneumoniae strain IF1SW-P3* | Flight 1 | 1 | 0.0000 |
| *Klebsiella quasipneumoniae strain IF1SW-B2* | *Klebsiella quasipneumoniae strain IF1SW-P4* | Flight 1 | 1 | 0.0000 |
| *Klebsiella quasipneumoniae strain IF1SW-B2* | *Penicillium chrysogenum* | Flight 1 | 1 | 0.1374 |
| *Klebsiella quasipneumoniae strain IF1SW-B2* | *Penicillium rubens* | Flight 1 | 1 | 0.1374 |
| *Klebsiella quasipneumoniae strain IF1SW-B2* | *Rhodotorula sp. strain JG-1b* | Flight 1 | 1 | 0.0000 |
| *Klebsiella quasipneumoniae strain IF1SW-B2* | *Rhodotorula toruloides* | Flight 1 | 1 | 0.0000 |
| *Klebsiella quasipneumoniae strain IF1SW-P3* | *Aspergillus niger* | Flight 1 | 1 | 0.0000 |
| *Klebsiella quasipneumoniae strain IF1SW-P3* | *Klebsiella pneumoniae* | Flight 1 | 1 | 0.0000 |
| *Klebsiella quasipneumoniae strain IF1SW-P3* | *Klebsiella quasipneumoniae strain IF1SW-B2* | Flight 1 | 1 | 0.0000 |
| *Klebsiella quasipneumoniae strain IF1SW-P3* | *Klebsiella quasipneumoniae strain IF1SW-P4* | Flight 1 | 1 | 0.0000 |
| *Klebsiella quasipneumoniae strain IF1SW-P3* | *Penicillium chrysogenum* | Flight 1 | 1 | 0.1374 |
| *Klebsiella quasipneumoniae strain IF1SW-P3* | *Penicillium rubens* | Flight 1 | 1 | 0.1374 |
| *Klebsiella quasipneumoniae strain IF1SW-P3* | *Rhodotorula sp. strain JG-1b* | Flight 1 | 1 | 0.0000 |
| *Klebsiella quasipneumoniae strain IF1SW-P3* | *Rhodotorula toruloides* | Flight 1 | 1 | 0.0000 |
| *Klebsiella quasipneumoniae strain IF1SW-P4* | *Aspergillus niger* | Flight 1 | 1 | 0.0000 |
| *Klebsiella quasipneumoniae strain IF1SW-P4* | *Klebsiella pneumoniae* | Flight 1 | 1 | 0.0000 |
| *Klebsiella quasipneumoniae strain IF1SW-P4* | *Klebsiella quasipneumoniae strain IF1SW-B2* | Flight 1 | 1 | 0.0000 |
| *Klebsiella quasipneumoniae strain IF1SW-P4* | *Klebsiella quasipneumoniae strain IF1SW-P3* | Flight 1 | 1 | 0.0000 |
| *Klebsiella quasipneumoniae strain IF1SW-P4* | *Penicillium chrysogenum* | Flight 1 | 1 | 0.1466 |
| *Klebsiella quasipneumoniae strain IF1SW-P4* | *Penicillium rubens* | Flight 1 | 1 | 0.1466 |
| *Klebsiella quasipneumoniae strain IF1SW-P4* | *Rhodotorula sp. strain JG-1b* | Flight 1 | 1 | 0.0000 |
| *Klebsiella quasipneumoniae strain IF1SW-P4* | *Rhodotorula toruloides* | Flight 1 | 1 | 0.0000 |
| *Klebsiella quasipneumoniae strain IF2SW-B3* | *Enterobacter asburiae* | Flight 1 | 2 | 0.0000 |
| *Klebsiella quasipneumoniae strain IF2SW-B3* | *Enterobacter cancerogenus* | Flight 1 | 2 | 0.0000 |
| *Klebsiella quasipneumoniae strain IF2SW-B3* | *Enterobacter cloacae* | Flight 1 | 2 | 0.0000 |
| *Klebsiella quasipneumoniae strain IF2SW-B3* | *Enterobacter hormaechei* | Flight 1 | 2 | 0.0000 |
| *Klebsiella quasipneumoniae strain IF2SW-B3* | *Enterobacter roggenkampii* | Flight 1 | 2 | 0.0000 |
| *Klebsiella quasipneumoniae strain IF2SW-B3* | *Enterobacter sp. strain NFIX59* | Flight 1 | 2 | 0.0000 |
| *Klebsiella quasipneumoniae strain IF2SW-B3* | *Escherichia coli* | Flight 1 | 2 | 0.0000 |
| *Klebsiella quasipneumoniae strain IF2SW-B3* | *Klebsiella oxytoca* | Flight 1 | 2 | 0.4121 |
| *Klebsiella quasipneumoniae strain IF2SW-B3* | *Klebsiella pneumoniae* | Flight 1 | 2 | 0.0000 |
| *Klebsiella quasipneumoniae strain IF2SW-B3* | *Klebsiella quasipneumoniae strain IF2SW-P1* | Flight 1 | 2 | 0.0000 |
| *Klebsiella quasipneumoniae strain IF2SW-B3* | *Pantoea ananatis* | Flight 1 | 2 | 0.0000 |
| *Klebsiella quasipneumoniae strain IF2SW-B3* | *Penicillium rubens* | Flight 1 | 2 | 0.1374 |
| *Klebsiella quasipneumoniae strain IF2SW-B3* | *Rhodotorula sp. strain JG-1b* | Flight 1 | 2 | 0.0000 |

| Microorganism | In the presence of | Flight | Location | Metabolic Support Index (A\|AUB) (%) |
|---|---|---|---|---|
| *Klebsiella quasipneumoniae strain IF2SW-B3* | *Salmonella enterica* | Flight 1 | 2 | 0.0000 |
| *Klebsiella quasipneumoniae strain IF2SW-B3* | *Shigella sonnei* | Flight 1 | 2 | 0.0000 |
| *Klebsiella quasipneumoniae strain IF2SW-P1* | *Enterobacter asburiae* | Flight 1 | 2 | 0.0000 |
| *Klebsiella quasipneumoniae strain IF2SW-P1* | *Enterobacter cancerogenus* | Flight 1 | 2 | 0.0000 |
| *Klebsiella quasipneumoniae strain IF2SW-P1* | *Enterobacter cloacae* | Flight 1 | 2 | 0.0000 |
| *Klebsiella quasipneumoniae strain IF2SW-P1* | *Enterobacter hormaechei* | Flight 1 | 2 | 0.0000 |
| *Klebsiella quasipneumoniae strain IF2SW-P1* | *Enterobacter roggenkampii* | Flight 1 | 2 | 0.0000 |
| *Klebsiella quasipneumoniae strain IF2SW-P1* | *Enterobacter sp. strain NFIX59* | Flight 1 | 2 | 0.0000 |
| *Klebsiella quasipneumoniae strain IF2SW-P1* | *Escherichia coli* | Flight 1 | 2 | 0.0000 |
| *Klebsiella quasipneumoniae strain IF2SW-P1* | *Klebsiella oxytoca* | Flight 1 | 2 | 0.4121 |
| *Klebsiella quasipneumoniae strain IF2SW-P1* | *Klebsiella pneumoniae* | Flight 1 | 2 | 0.0000 |
| *Klebsiella quasipneumoniae strain IF2SW-P1* | *Klebsiella quasipneumoniae strain IF2SW-B3* | Flight 1 | 2 | 0.0000 |
| *Klebsiella quasipneumoniae strain IF2SW-P1* | *Pantoea ananatis* | Flight 1 | 2 | 0.0000 |
| *Klebsiella quasipneumoniae strain IF2SW-P1* | *Penicillium rubens* | Flight 1 | 2 | 0.1374 |
| *Klebsiella quasipneumoniae strain IF2SW-P1* | *Rhodotorula sp. strain JG-1b* | Flight 1 | 2 | 0.0000 |
| *Klebsiella quasipneumoniae strain IF2SW-P1* | *Salmonella enterica* | Flight 1 | 2 | 0.0000 |
| *Klebsiella quasipneumoniae strain IF2SW-P1* | *Shigella sonnei* | Flight 1 | 2 | 0.0000 |
| *Klebsiella quasipneumoniae strain IIIF3SW-P1* | *Escherichia coli* | Flight 3 | 3 | 0.0000 |
| *Klebsiella quasipneumoniae strain IIIF3SW-P1* | *Klebsiella pneumoniae* | Flight 3 | 3 | 0.0000 |
| *Klebsiella quasipneumoniae strain IIIF3SW-P1* | *Klebsiella sp. strain MS 92-3* | Flight 3 | 3 | 0.0000 |
| *Klebsiella quasipneumoniae strain IIIF3SW-P1* | *Klebsiella variicola* | Flight 3 | 3 | 0.0000 |
| *Klebsiella quasipneumoniae strain IIIF3SW-P1* | *Penicillium rubens* | Flight 3 | 3 | 0.6766 |
| *Klebsiella quasipneumoniae strain IIIF3SW-P1* | *Salmonella enterica* | Flight 3 | 3 | 0.0000 |
| *Klebsiella quasipneumoniae strain IIIF3SW-P1* | *Staphylococcus saprophyticus* | Flight 3 | 3 | 0.0000 |
| *Klebsiella sp. strain MS 92-3* | *Escherichia coli* | Flight 3 | 3 | 0.0000 |
| *Klebsiella sp. strain MS 92-3* | *Klebsiella pneumoniae* | Flight 3 | 3 | 0.0000 |
| *Klebsiella sp. strain MS 92-3* | *Klebsiella quasipneumoniae strain IIIF3SW-P1* | Flight 3 | 3 | 0.0000 |
| *Klebsiella sp. strain MS 92-3* | *Klebsiella variicola* | Flight 3 | 3 | 0.0000 |
| *Klebsiella sp. strain MS 92-3* | *Penicillium rubens* | Flight 3 | 3 | 0.6859 |
| *Klebsiella sp. strain MS 92-3* | *Salmonella enterica* | Flight 3 | 3 | 0.0000 |
| *Klebsiella sp. strain MS 92-3* | *Staphylococcus saprophyticus* | Flight 3 | 3 | 0.0000 |
| *Klebsiella variicola* | *Escherichia coli* | Flight 3 | 3 | 0.0000 |
| *Klebsiella variicola* | *Klebsiella pneumoniae* | Flight 3 | 3 | 0.0000 |
| *Klebsiella variicola* | *Klebsiella quasipneumoniae strain IIIF3SW-P1* | Flight 3 | 3 | 0.0000 |
| *Klebsiella variicola* | *Klebsiella sp. strain MS 92-3* | Flight 3 | 3 | 0.0000 |
| *Klebsiella variicola* | *Penicillium rubens* | Flight 3 | 3 | 0.6821 |
| *Klebsiella variicola* | *Salmonella enterica* | Flight 3 | 3 | 0.0000 |
| *Klebsiella variicola* | *Staphylococcus saprophyticus* | Flight 3 | 3 | 0.0000 |
| *Paenibacillus polymyxa* | *Enterococcus avium* | Flight 3 | 2 | 0.0000 |
| *Paenibacillus polymyxa* | *Enterococcus faecalis* | Flight 3 | 2 | 0.0000 |
| *Paenibacillus polymyxa* | *Klebsiella pneumoniae strain F3-2P(2*)* | Flight 3 | 2 | 0.1629 |
| *Paenibacillus polymyxa* | *Penicillium chrysogenum* | Flight 3 | 2 | 0.1629 |
| *Paenibacillus polymyxa* | *Penicillium flavigenum* | Flight 3 | 2 | 0.1629 |
| *Paenibacillus polymyxa* | *Penicillium nalgiovense* | Flight 3 | 2 | 0.1629 |
| *Paenibacillus polymyxa* | *Penicillium rubens* | Flight 3 | 2 | 0.1629 |
| *Paenibacillus polymyxa* | *Rhodotorula sp. strain JG-1b* | Flight 3 | 2 | 0.0000 |
| *Paenibacillus polymyxa* | *Staphylococcus aureus* | Flight 3 | 2 | 0.0000 |
| *Paenibacillus polymyxa* | *Staphylococcus epidermidis* | Flight 3 | 2 | 0.0000 |
| *Paenibacillus polymyxa* | *Staphylococcus haemolyticus* | Flight 3 | 2 | 0.0000 |
| *Paenibacillus polymyxa* | *Staphylococcus saprophyticus* | Flight 3 | 2 | 0.0000 |
| *Paenibacillus polymyxa* | *Staphylococcus sp. strain LCT-H4* | Flight 3 | 2 | 0.0000 |
| *Paenibacillus polymyxa* | *Staphylococcus warneri* | Flight 3 | 2 | 0.0000 |
| *Paenibacillus polymyxa strain IIF5SW-B3* | *Escherichia coli* | Flight 2 | 5 | 0.1383 |
| *Paenibacillus polymyxa strain IIF5SW-B3* | *Klebsiella pneumoniae* | Flight 2 | 5 | 0.1383 |
| *Paenibacillus polymyxa strain IIF5SW-B3* | *Paenibacillus polymyxa strain IIF5SW-B4* | Flight 2 | 5 | 0.0000 |
| *Paenibacillus polymyxa strain IIF5SW-B3* | *Pantoea agglomerans* | Flight 2 | 5 | 0.1383 |
| *Paenibacillus polymyxa strain IIF5SW-B3* | *Pantoea ananatis* | Flight 2 | 5 | 0.1383 |
| *Paenibacillus polymyxa strain IIF5SW-B3* | *Pantoea conspicua* | Flight 2 | 5 | 0.9682 |
| *Paenibacillus polymyxa strain IIF5SW-B3* | *Pantoea sp. strain 3.5.1* | Flight 2 | 5 | 0.9682 |
| *Paenibacillus polymyxa strain IIF5SW-B3* | *Pantoea vagans* | Flight 2 | 5 | 0.1383 |
| *Paenibacillus polymyxa strain IIF5SW-B4* | *Escherichia coli* | Flight 2 | 5 | 0.1383 |
| *Paenibacillus polymyxa strain IIF5SW-B4* | *Klebsiella pneumoniae* | Flight 2 | 5 | 0.1383 |
| *Paenibacillus polymyxa strain IIF5SW-B4* | *Paenibacillus polymyxa strain IIF5SW-B3* | Flight 2 | 5 | 0.0000 |
| *Paenibacillus polymyxa strain IIF5SW-B4* | *Pantoea agglomerans* | Flight 2 | 5 | 0.1383 |
| *Paenibacillus polymyxa strain IIF5SW-B4* | *Pantoea ananatis* | Flight 2 | 5 | 0.1383 |
| *Paenibacillus polymyxa strain IIF5SW-B4* | *Pantoea conspicua* | Flight 2 | 5 | 0.9682 |
| *Paenibacillus polymyxa strain IIF5SW-B4* | *Pantoea sp. strain 3.5.1* | Flight 2 | 5 | 0.9682 |
| *Paenibacillus polymyxa strain IIF5SW-B4* | *Pantoea vagans* | Flight 2 | 5 | 0.1383 |
| *Pantoea agglomerans* | *Aspergillus niger* | Flight 3 | 1 | 0.0000 |
| *Pantoea agglomerans* | *Enterobacter cloacae* | Flight 3 | 1 | 0.0000 |
| *Pantoea agglomerans* | *Enterobacter cloacae* | Flight 3 | 5 | 4.8303 |
| *Pantoea agglomerans* | *Enterobacter cloacae* | Flight 3 | 7 | 4.8303 |
| *Pantoea agglomerans* | *Enterobacter cloacae* | Flight 3 | 8 | 4.8303 |
| *Pantoea agglomerans* | *Escherichia coli* | Flight 1 | 5 | 0.8523 |
| *Pantoea agglomerans* | *Escherichia coli* | Flight 2 | 5 | 0.7833 |
| *Pantoea agglomerans* | *Escherichia coli* | Flight 3 | 1 | 0.8333 |
| *Pantoea agglomerans* | *Escherichia coli* | Flight 3 | 5 | 0.7833 |
| *Pantoea agglomerans* | *Escherichia coli* | Flight 3 | 7 | 0.7833 |
| *Pantoea agglomerans* | *Escherichia coli* | Flight 3 | 8 | 0.7833 |
| *Pantoea agglomerans* | *Klebsiella aerogenes strain IIIF7SW-P1* | Flight 3 | 7 | 5.6136 |
| *Pantoea agglomerans* | *Klebsiella pneumoniae* | Flight 1 | 5 | 0.8523 |
| *Pantoea agglomerans* | *Klebsiella pneumoniae* | Flight 2 | 5 | 5.6136 |
| *Pantoea agglomerans* | *Klebsiella pneumoniae* | Flight 3 | 1 | 0.8333 |
| *Pantoea agglomerans* | *Klebsiella pneumoniae* | Flight 3 | 5 | 5.6136 |
| *Pantoea agglomerans* | *Klebsiella pneumoniae* | Flight 3 | 7 | 5.6136 |
| *Pantoea agglomerans* | *Klebsiella pneumoniae* | Flight 3 | 8 | 5.6136 |
| *Pantoea agglomerans* | *Paenibacillus polymyxa strain IIF5SW-B3* | Flight 2 | 5 | 4.8303 |
| *Pantoea agglomerans* | *Paenibacillus polymyxa strain IIF5SW-B4* | Flight 2 | 5 | 4.8303 |
| *Pantoea agglomerans* | *Pantoea ananatis* | Flight 1 | 5 | 0.0000 |
| *Pantoea agglomerans* | *Pantoea ananatis* | Flight 2 | 5 | 0.0000 |
| *Pantoea agglomerans* | *Pantoea ananatis* | Flight 3 | 1 | 0.0000 |
| *Pantoea agglomerans* | *Pantoea ananatis* | Flight 3 | 5 | 0.0000 |
| *Pantoea agglomerans* | *Pantoea ananatis* | Flight 3 | 7 | 0.0000 |
| *Pantoea agglomerans* | *Pantoea ananatis* | Flight 3 | 8 | 0.0000 |
| *Pantoea agglomerans* | *Pantoea conspicua* | Flight 1 | 5 | 0.8523 |
| *Pantoea agglomerans* | *Pantoea conspicua* | Flight 2 | 5 | 0.7833 |

| Microorganism | In the presence of | Flight | Location | Metabolic Support Index (A\|AUB) (%) |
|---|---|---|---|---|
| *Pantoea agglomerans* | *Pantoea conspicua* | Flight 3 | 1 | 0.8333 |
| *Pantoea agglomerans* | *Pantoea conspicua* | Flight 3 | 5 | 0.7833 |
| *Pantoea agglomerans* | *Pantoea conspicua* | Flight 3 | 7 | 0.7833 |
| *Pantoea agglomerans* | *Pantoea conspicua* | Flight 3 | 8 | 0.7833 |
| *Pantoea agglomerans* | *Pantoea sp. strain 3.5.1* | Flight 1 | 5 | 0.8523 |
| *Pantoea agglomerans* | *Pantoea sp. strain 3.5.1* | Flight 2 | 5 | 0.7833 |
| *Pantoea agglomerans* | *Pantoea sp. strain A4* | Flight 3 | 1 | 0.0000 |
| *Pantoea agglomerans* | *Pantoea sp. strain A4* | Flight 3 | 5 | 0.0000 |
| *Pantoea agglomerans* | *Pantoea sp. strain A4* | Flight 3 | 7 | 0.0000 |
| *Pantoea agglomerans* | *Pantoea sp. strain A4* | Flight 3 | 8 | 0.0000 |
| *Pantoea agglomerans* | *Pantoea sp. strain At-9b* | Flight 3 | 1 | 0.0000 |
| *Pantoea agglomerans* | *Pantoea sp. strain At-9b* | Flight 3 | 5 | 0.0000 |
| *Pantoea agglomerans* | *Pantoea sp. strain At-9b* | Flight 3 | 8 | 0.0000 |
| *Pantoea agglomerans* | *Pantoea sp. strain FF5* | Flight 3 | 1 | 0.0000 |
| *Pantoea agglomerans* | *Pantoea sp. strain FF5* | Flight 3 | 5 | 0.0000 |
| *Pantoea agglomerans* | *Pantoea sp. strain FF5* | Flight 3 | 7 | 0.0000 |
| *Pantoea agglomerans* | *Pantoea sp. strain FF5* | Flight 3 | 8 | 0.0000 |
| *Pantoea agglomerans* | *Pantoea sp. strain IMH* | Flight 3 | 1 | 0.8333 |
| *Pantoea agglomerans* | *Pantoea sp. strain IMH* | Flight 3 | 5 | 0.7833 |
| *Pantoea agglomerans* | *Pantoea sp. strain IMH* | Flight 3 | 7 | 0.7833 |
| *Pantoea agglomerans* | *Pantoea sp. strain IMH* | Flight 3 | 8 | 0.7833 |
| *Pantoea agglomerans* | *Pantoea sp. strain NGS-ED-1003* | Flight 3 | 1 | 0.8333 |
| *Pantoea agglomerans* | *Pantoea sp. strain NGS-ED-1003* | Flight 3 | 5 | 0.7833 |
| *Pantoea agglomerans* | *Pantoea sp. strain NGS-ED-1003* | Flight 3 | 7 | 0.7833 |
| *Pantoea agglomerans* | *Pantoea sp. strain NGS-ED-1003* | Flight 3 | 8 | 0.7833 |
| *Pantoea agglomerans* | *Pantoea sp. strain OXWO6B1* | Flight 3 | 1 | 0.0000 |
| *Pantoea agglomerans* | *Pantoea sp. strain OXWO6B1* | Flight 3 | 5 | 0.0000 |
| *Pantoea agglomerans* | *Pantoea sp. strain OXWO6B1* | Flight 3 | 7 | 0.0000 |
| *Pantoea agglomerans* | *Pantoea sp. strain OXWO6B1* | Flight 3 | 8 | 0.0000 |
| *Pantoea agglomerans* | *Pantoea vagans* | Flight 2 | 5 | 0.0000 |
| *Pantoea agglomerans* | *Pantoea vagans* | Flight 3 | 1 | 0.0000 |
| *Pantoea agglomerans* | *Pantoea vagans* | Flight 3 | 5 | 0.0000 |
| *Pantoea agglomerans* | *Pantoea vagans* | Flight 3 | 7 | 0.0000 |
| *Pantoea agglomerans* | *Pantoea vagans* | Flight 3 | 8 | 0.0000 |
| *Pantoea agglomerans* | *Penicillium rubens* | Flight 1 | 5 | 1.2784 |
| *Pantoea agglomerans* | *Rhodotorula sp. strain JG-1b* | Flight 1 | 5 | 0.8523 |
| *Pantoea agglomerans* | *Rhodotorula sp. strain JG-1b* | Flight 3 | 1 | 0.8333 |
| *Pantoea agglomerans* | *Salmonella enterica* | Flight 3 | 1 | 0.8333 |
| *Pantoea agglomerans* | *Salmonella enterica* | Flight 3 | 5 | 0.7833 |
| *Pantoea agglomerans* | *Salmonella enterica* | Flight 3 | 7 | 0.7833 |
| *Pantoea agglomerans* | *Salmonella enterica* | Flight 3 | 8 | 0.7833 |
| *Pantoea ananatis* | *Aspergillus niger* | Flight 3 | 1 | 0.0000 |
| *Pantoea ananatis* | *Enterobacter asburiae* | Flight 1 | 2 | 0.0000 |
| *Pantoea ananatis* | *Enterobacter cancerogenus* | Flight 1 | 2 | 0.0000 |
| *Pantoea ananatis* | *Enterobacter cloacae* | Flight 1 | 2 | 0.0000 |
| *Pantoea ananatis* | *Enterobacter cloacae* | Flight 3 | 1 | 0.0000 |
| *Pantoea ananatis* | *Enterobacter cloacae* | Flight 3 | 5 | 4.9479 |
| *Pantoea ananatis* | *Enterobacter cloacae* | Flight 3 | 7 | 4.9479 |
| *Pantoea ananatis* | *Enterobacter cloacae* | Flight 3 | 8 | 4.9479 |
| *Pantoea ananatis* | *Enterobacter hormaechei* | Flight 1 | 2 | 0.0000 |
| *Pantoea ananatis* | *Enterobacter hormaechei* | Flight 3 | 4 | 4.9479 |
| *Pantoea ananatis* | *Enterobacter roggenkampii* | Flight 1 | 2 | 0.0000 |
| *Pantoea ananatis* | *Enterobacter sp. strain NFIX59* | Flight 1 | 2 | 0.0000 |
| *Pantoea ananatis* | *Escherichia coli* | Flight 1 | 2 | 0.0000 |
| *Pantoea ananatis* | *Escherichia coli* | Flight 1 | 5 | 0.0000 |
| *Pantoea ananatis* | *Escherichia coli* | Flight 2 | 5 | 0.0000 |
| *Pantoea ananatis* | *Escherichia coli* | Flight 3 | 1 | 0.0000 |
| *Pantoea ananatis* | *Escherichia coli* | Flight 3 | 4 | 0.0000 |
| *Pantoea ananatis* | *Escherichia coli* | Flight 3 | 5 | 0.0000 |
| *Pantoea ananatis* | *Escherichia coli* | Flight 3 | 7 | 0.0000 |
| *Pantoea ananatis* | *Escherichia coli* | Flight 3 | 8 | 0.0000 |
| *Pantoea ananatis* | *Klebsiella aerogenes strain IIIF7SW-P1* | Flight 3 | 7 | 4.9479 |
| *Pantoea ananatis* | *Klebsiella oxytoca* | Flight 1 | 2 | 0.0000 |
| *Pantoea ananatis* | *Klebsiella pneumoniae* | Flight 1 | 2 | 0.0000 |
| *Pantoea ananatis* | *Klebsiella pneumoniae* | Flight 1 | 5 | 0.0000 |
| *Pantoea ananatis* | *Klebsiella pneumoniae* | Flight 2 | 5 | 4.9479 |
| *Pantoea ananatis* | *Klebsiella pneumoniae* | Flight 3 | 1 | 0.0000 |
| *Pantoea ananatis* | *Klebsiella pneumoniae* | Flight 3 | 4 | 4.9479 |
| *Pantoea ananatis* | *Klebsiella pneumoniae* | Flight 3 | 5 | 4.9479 |
| *Pantoea ananatis* | *Klebsiella pneumoniae* | Flight 3 | 7 | 4.9479 |
| *Pantoea ananatis* | *Klebsiella pneumoniae* | Flight 3 | 8 | 4.9479 |
| *Pantoea ananatis* | *Klebsiella quasipneumoniae strain IF2SW-B3* | Flight 1 | 2 | 0.0000 |
| *Pantoea ananatis* | *Klebsiella quasipneumoniae strain IF2SW-P1* | Flight 1 | 2 | 0.0000 |
| *Pantoea ananatis* | *Paenibacillus polymyxa strain IIF5SW-B3* | Flight 2 | 5 | 4.9479 |
| *Pantoea ananatis* | *Paenibacillus polymyxa strain IIF5SW-B4* | Flight 2 | 5 | 4.9479 |
| *Pantoea ananatis* | *Pantoea agglomerans* | Flight 1 | 5 | 0.0000 |
| *Pantoea ananatis* | *Pantoea agglomerans* | Flight 2 | 5 | 0.0000 |
| *Pantoea ananatis* | *Pantoea agglomerans* | Flight 3 | 1 | 0.0000 |
| *Pantoea ananatis* | *Pantoea agglomerans* | Flight 3 | 5 | 0.0000 |
| *Pantoea ananatis* | *Pantoea agglomerans* | Flight 3 | 7 | 0.0000 |
| *Pantoea ananatis* | *Pantoea agglomerans* | Flight 3 | 8 | 0.0000 |
| *Pantoea ananatis* | *Pantoea conspicua* | Flight 1 | 5 | 0.0000 |
| *Pantoea ananatis* | *Pantoea conspicua* | Flight 2 | 5 | 0.0000 |
| *Pantoea ananatis* | *Pantoea conspicua* | Flight 3 | 1 | 0.0000 |
| *Pantoea ananatis* | *Pantoea conspicua* | Flight 3 | 5 | 0.0000 |
| *Pantoea ananatis* | *Pantoea conspicua* | Flight 3 | 7 | 0.0000 |
| *Pantoea ananatis* | *Pantoea conspicua* | Flight 3 | 8 | 0.0000 |
| *Pantoea ananatis* | *Pantoea dispersa* | Flight 3 | 4 | 0.0000 |
| *Pantoea ananatis* | *Pantoea sp. strain 3.5.1* | Flight 1 | 5 | 0.0000 |
| *Pantoea ananatis* | *Pantoea sp. strain 3.5.1* | Flight 2 | 5 | 0.0000 |
| *Pantoea ananatis* | *Pantoea sp. strain A4* | Flight 3 | 1 | 0.0000 |
| *Pantoea ananatis* | *Pantoea sp. strain A4* | Flight 3 | 5 | 0.0000 |
| *Pantoea ananatis* | *Pantoea sp. strain A4* | Flight 3 | 7 | 0.0000 |
| *Pantoea ananatis* | *Pantoea sp. strain A4* | Flight 3 | 8 | 0.0000 |
| *Pantoea ananatis* | *Pantoea sp. strain At-9b* | Flight 3 | 1 | 0.0000 |
| *Pantoea ananatis* | *Pantoea sp. strain At-9b* | Flight 3 | 5 | 0.0000 |

| Microorganism | In the presence of | Flight | Location | Metabolic Support Index (A\|AUB) (%) |
|---|---|---|---|---|
| *Pantoea ananatis* | *Pantoea sp. strain At-9b* | Flight 3 | 8 | 0.0000 |
| *Pantoea ananatis* | *Pantoea sp. strain FF5* | Flight 3 | 1 | 0.0000 |
| *Pantoea ananatis* | *Pantoea sp. strain FF5* | Flight 3 | 5 | 0.0000 |
| *Pantoea ananatis* | *Pantoea sp. strain FF5* | Flight 3 | 7 | 0.0000 |
| *Pantoea ananatis* | *Pantoea sp. strain FF5* | Flight 3 | 8 | 0.0000 |
| *Pantoea ananatis* | *Pantoea sp. strain IMH* | Flight 3 | 1 | 0.0000 |
| *Pantoea ananatis* | *Pantoea sp. strain IMH* | Flight 3 | 5 | 0.0000 |
| *Pantoea ananatis* | *Pantoea sp. strain IMH* | Flight 3 | 7 | 0.0000 |
| *Pantoea ananatis* | *Pantoea sp. strain IMH* | Flight 3 | 8 | 0.0000 |
| *Pantoea ananatis* | *Pantoea sp. strain NGS-ED-1003* | Flight 3 | 1 | 0.0000 |
| *Pantoea ananatis* | *Pantoea sp. strain NGS-ED-1003* | Flight 3 | 5 | 0.0000 |
| *Pantoea ananatis* | *Pantoea sp. strain NGS-ED-1003* | Flight 3 | 7 | 0.0000 |
| *Pantoea ananatis* | *Pantoea sp. strain NGS-ED-1003* | Flight 3 | 8 | 0.0000 |
| *Pantoea ananatis* | *Pantoea sp. strain OXWO6B1* | Flight 3 | 1 | 0.0000 |
| *Pantoea ananatis* | *Pantoea sp. strain OXWO6B1* | Flight 3 | 5 | 0.0000 |
| *Pantoea ananatis* | *Pantoea sp. strain OXWO6B1* | Flight 3 | 7 | 0.0000 |
| *Pantoea ananatis* | *Pantoea sp. strain OXWO6B1* | Flight 3 | 8 | 0.0000 |
| *Pantoea ananatis* | *Pantoea vagans* | Flight 2 | 5 | 0.0000 |
| *Pantoea ananatis* | *Pantoea vagans* | Flight 3 | 1 | 0.0000 |
| *Pantoea ananatis* | *Pantoea vagans* | Flight 3 | 5 | 0.0000 |
| *Pantoea ananatis* | *Pantoea vagans* | Flight 3 | 7 | 0.0000 |
| *Pantoea ananatis* | *Pantoea vagans* | Flight 3 | 8 | 0.0000 |
| *Pantoea ananatis* | *Penicillium rubens* | Flight 1 | 2 | 0.5690 |
| *Pantoea ananatis* | *Penicillium rubens* | Flight 1 | 5 | 0.5690 |
| *Pantoea ananatis* | *Rahnella aquatilis* | Flight 3 | 4 | 0.0000 |
| *Pantoea ananatis* | *Rhodotorula sp. strain JG-1b* | Flight 1 | 2 | 0.0000 |
| *Pantoea ananatis* | *Rhodotorula sp. strain JG-1b* | Flight 1 | 5 | 0.0000 |
| *Pantoea ananatis* | *Rhodotorula sp. strain JG-1b* | Flight 3 | 1 | 0.0000 |
| *Pantoea ananatis* | *Salmonella enterica* | Flight 1 | 2 | 0.0000 |
| *Pantoea ananatis* | *Salmonella enterica* | Flight 3 | 1 | 0.0000 |
| *Pantoea ananatis* | *Salmonella enterica* | Flight 3 | 5 | 0.0000 |
| *Pantoea ananatis* | *Salmonella enterica* | Flight 3 | 7 | 0.0000 |
| *Pantoea ananatis* | *Salmonella enterica* | Flight 3 | 8 | 0.0000 |
| *Pantoea ananatis* | *Shigella sonnei* | Flight 1 | 2 | 0.0000 |
| *Pantoea conspicua* | *Aspergillus niger* | Flight 3 | 1 | 0.0000 |
| *Pantoea conspicua* | *Enterobacter cloacae* | Flight 3 | 1 | 0.9464 |
| *Pantoea conspicua* | *Enterobacter cloacae* | Flight 3 | 5 | 6.4611 |
| *Pantoea conspicua* | *Enterobacter cloacae* | Flight 3 | 7 | 6.4611 |
| *Pantoea conspicua* | *Enterobacter cloacae* | Flight 3 | 8 | 6.4611 |
| *Pantoea conspicua* | *Escherichia coli* | Flight 1 | 5 | 0.9709 |
| *Pantoea conspicua* | *Escherichia coli* | Flight 2 | 5 | 0.8811 |
| *Pantoea conspicua* | *Escherichia coli* | Flight 3 | 1 | 0.9464 |
| *Pantoea conspicua* | *Escherichia coli* | Flight 3 | 5 | 0.8811 |
| *Pantoea conspicua* | *Escherichia coli* | Flight 3 | 7 | 0.8811 |
| *Pantoea conspicua* | *Escherichia coli* | Flight 3 | 8 | 0.8811 |
| *Pantoea conspicua* | *Klebsiella aerogenes strain IIIF7SW-P1* | Flight 3 | 7 | 6.4611 |
| *Pantoea conspicua* | *Klebsiella pneumoniae* | Flight 1 | 5 | 0.9709 |
| *Pantoea conspicua* | *Klebsiella pneumoniae* | Flight 2 | 5 | 6.4611 |
| *Pantoea conspicua* | *Klebsiella pneumoniae* | Flight 3 | 1 | 0.9464 |
| *Pantoea conspicua* | *Klebsiella pneumoniae* | Flight 3 | 5 | 6.4611 |
| *Pantoea conspicua* | *Klebsiella pneumoniae* | Flight 3 | 7 | 6.4611 |
| *Pantoea conspicua* | *Klebsiella pneumoniae* | Flight 3 | 8 | 6.4611 |
| *Pantoea conspicua* | *Paenibacillus polymyxa strain IIF5SW-B3* | Flight 2 | 5 | 6.4611 |
| *Pantoea conspicua* | *Paenibacillus polymyxa strain IIF5SW-B4* | Flight 2 | 5 | 6.4611 |
| *Pantoea conspicua* | *Pantoea agglomerans* | Flight 1 | 5 | 0.9709 |
| *Pantoea conspicua* | *Pantoea agglomerans* | Flight 2 | 5 | 0.8811 |
| *Pantoea conspicua* | *Pantoea agglomerans* | Flight 3 | 1 | 0.9464 |
| *Pantoea conspicua* | *Pantoea agglomerans* | Flight 3 | 5 | 0.8811 |
| *Pantoea conspicua* | *Pantoea agglomerans* | Flight 3 | 7 | 0.8811 |
| *Pantoea conspicua* | *Pantoea agglomerans* | Flight 3 | 8 | 0.8811 |
| *Pantoea conspicua* | *Pantoea ananatis* | Flight 1 | 5 | 0.9709 |
| *Pantoea conspicua* | *Pantoea ananatis* | Flight 2 | 5 | 0.8811 |
| *Pantoea conspicua* | *Pantoea ananatis* | Flight 3 | 1 | 0.9464 |
| *Pantoea conspicua* | *Pantoea ananatis* | Flight 3 | 5 | 0.8811 |
| *Pantoea conspicua* | *Pantoea ananatis* | Flight 3 | 7 | 0.8811 |
| *Pantoea conspicua* | *Pantoea ananatis* | Flight 3 | 8 | 0.8811 |
| *Pantoea conspicua* | *Pantoea sp. strain 3.5.1* | Flight 1 | 5 | 0.9709 |
| *Pantoea conspicua* | *Pantoea sp. strain 3.5.1* | Flight 2 | 5 | 0.8811 |
| *Pantoea conspicua* | *Pantoea sp. strain A4* | Flight 3 | 1 | 0.0000 |
| *Pantoea conspicua* | *Pantoea sp. strain A4* | Flight 3 | 5 | 0.0000 |
| *Pantoea conspicua* | *Pantoea sp. strain A4* | Flight 3 | 7 | 0.0000 |
| *Pantoea conspicua* | *Pantoea sp. strain A4* | Flight 3 | 8 | 0.0000 |
| *Pantoea conspicua* | *Pantoea sp. strain At-9b* | Flight 3 | 1 | 0.9464 |
| *Pantoea conspicua* | *Pantoea sp. strain At-9b* | Flight 3 | 5 | 0.8811 |
| *Pantoea conspicua* | *Pantoea sp. strain At-9b* | Flight 3 | 8 | 0.8811 |
| *Pantoea conspicua* | *Pantoea sp. strain FF5* | Flight 3 | 1 | 0.0000 |
| *Pantoea conspicua* | *Pantoea sp. strain FF5* | Flight 3 | 5 | 0.0000 |
| *Pantoea conspicua* | *Pantoea sp. strain FF5* | Flight 3 | 7 | 0.0000 |
| *Pantoea conspicua* | *Pantoea sp. strain FF5* | Flight 3 | 8 | 0.0000 |
| *Pantoea conspicua* | *Pantoea sp. strain IMH* | Flight 3 | 1 | 0.9464 |
| *Pantoea conspicua* | *Pantoea sp. strain IMH* | Flight 3 | 5 | 0.8811 |
| *Pantoea conspicua* | *Pantoea sp. strain IMH* | Flight 3 | 7 | 0.8811 |
| *Pantoea conspicua* | *Pantoea sp. strain IMH* | Flight 3 | 8 | 0.8811 |
| *Pantoea conspicua* | *Pantoea sp. strain NGS-ED-1003* | Flight 3 | 1 | 0.9464 |
| *Pantoea conspicua* | *Pantoea sp. strain NGS-ED-1003* | Flight 3 | 5 | 0.8811 |
| *Pantoea conspicua* | *Pantoea sp. strain NGS-ED-1003* | Flight 3 | 7 | 0.8811 |
| *Pantoea conspicua* | *Pantoea sp. strain NGS-ED-1003* | Flight 3 | 8 | 0.8811 |
| *Pantoea conspicua* | *Pantoea sp. strain OXWO6B1* | Flight 3 | 1 | 0.9464 |
| *Pantoea conspicua* | *Pantoea sp. strain OXWO6B1* | Flight 3 | 5 | 0.8811 |
| *Pantoea conspicua* | *Pantoea sp. strain OXWO6B1* | Flight 3 | 7 | 0.8811 |
| *Pantoea conspicua* | *Pantoea sp. strain OXWO6B1* | Flight 3 | 8 | 0.8811 |
| *Pantoea conspicua* | *Pantoea vagans* | Flight 2 | 5 | 0.8811 |
| *Pantoea conspicua* | *Pantoea vagans* | Flight 3 | 1 | 0.9464 |
| *Pantoea conspicua* | *Pantoea vagans* | Flight 3 | 5 | 0.8811 |
| *Pantoea conspicua* | *Pantoea vagans* | Flight 3 | 7 | 0.8811 |
| *Pantoea conspicua* | *Pantoea vagans* | Flight 3 | 8 | 0.8811 |

| Microorganism | In the presence of | Flight | Location | Metabolic Support Index (A\|AUB) (%) |
|---|---|---|---|---|
| *Pantoea conspicua* | *Penicillium rubens* | Flight 1 | 5 | 0.4854 |
| *Pantoea conspicua* | *Rhodotorula sp. strain JG-1b* | Flight 1 | 5 | 0.0000 |
| *Pantoea conspicua* | *Rhodotorula sp. strain JG-1b* | Flight 3 | 1 | 0.0000 |
| *Pantoea conspicua* | *Salmonella enterica* | Flight 3 | 1 | 0.9464 |
| *Pantoea conspicua* | *Salmonella enterica* | Flight 3 | 5 | 0.8811 |
| *Pantoea conspicua* | *Salmonella enterica* | Flight 3 | 7 | 0.8811 |
| *Pantoea conspicua* | *Salmonella enterica* | Flight 3 | 8 | 0.8811 |
| *Pantoea dispersa* | *Enterobacter hormaechei* | Flight 3 | 4 | 5.2039 |
| *Pantoea dispersa* | *Escherichia coli* | Flight 3 | 4 | 0.8439 |
| *Pantoea dispersa* | *Klebsiella pneumoniae* | Flight 3 | 4 | 6.0478 |
| *Pantoea dispersa* | *Pantoea ananatis* | Flight 3 | 4 | 0.0000 |
| *Pantoea dispersa* | *Rahnella aquatilis* | Flight 3 | 4 | 0.0000 |
| *Pantoea sp. strain 3.5.1* | *Escherichia coli* | Flight 1 | 5 | 0.8759 |
| *Pantoea sp. strain 3.5.1* | *Escherichia coli* | Flight 2 | 5 | 0.8032 |
| *Pantoea sp. strain 3.5.1* | *Klebsiella pneumoniae* | Flight 1 | 5 | 0.8759 |
| *Pantoea sp. strain 3.5.1* | *Klebsiella pneumoniae* | Flight 2 | 5 | 5.7564 |
| *Pantoea sp. strain 3.5.1* | *Paenibacillus polymyxa strain IIF5SW-B3* | Flight 2 | 5 | 4.9531 |
| *Pantoea sp. strain 3.5.1* | *Paenibacillus polymyxa strain IIF5SW-B4* | Flight 2 | 5 | 4.9531 |
| *Pantoea sp. strain 3.5.1* | *Pantoea agglomerans* | Flight 1 | 5 | 0.0000 |
| *Pantoea sp. strain 3.5.1* | *Pantoea agglomerans* | Flight 2 | 5 | 0.0000 |
| *Pantoea sp. strain 3.5.1* | *Pantoea ananatis* | Flight 1 | 5 | 0.0000 |
| *Pantoea sp. strain 3.5.1* | *Pantoea ananatis* | Flight 2 | 5 | 0.0000 |
| *Pantoea sp. strain 3.5.1* | *Pantoea conspicua* | Flight 1 | 5 | 0.0000 |
| *Pantoea sp. strain 3.5.1* | *Pantoea conspicua* | Flight 2 | 5 | 0.0000 |
| *Pantoea sp. strain 3.5.1* | *Pantoea vagans* | Flight 2 | 5 | 0.0000 |
| *Pantoea sp. strain 3.5.1* | *Penicillium rubens* | Flight 1 | 5 | 1.3139 |
| *Pantoea sp. strain 3.5.1* | *Rhodotorula sp. strain JG-1b* | Flight 1 | 5 | 0.8759 |
| *Pantoea sp. strain A4* | *Aspergillus niger* | Flight 3 | 1 | 0.0000 |
| *Pantoea sp. strain A4* | *Enterobacter cloacae* | Flight 3 | 1 | 0.0000 |
| *Pantoea sp. strain A4* | *Enterobacter cloacae* | Flight 3 | 5 | 4.6776 |
| *Pantoea sp. strain A4* | *Enterobacter cloacae* | Flight 3 | 7 | 4.6776 |
| *Pantoea sp. strain A4* | *Enterobacter cloacae* | Flight 3 | 8 | 4.6776 |
| *Pantoea sp. strain A4* | *Escherichia coli* | Flight 3 | 1 | 1.6107 |
| *Pantoea sp. strain A4* | *Escherichia coli* | Flight 3 | 5 | 1.5171 |
| *Pantoea sp. strain A4* | *Escherichia coli* | Flight 3 | 7 | 1.5171 |
| *Pantoea sp. strain A4* | *Escherichia coli* | Flight 3 | 8 | 1.5171 |
| *Pantoea sp. strain A4* | *Klebsiella aerogenes strain IIIF7SW-P1* | Flight 3 | 7 | 6.1947 |
| *Pantoea sp. strain A4* | *Klebsiella pneumoniae* | Flight 3 | 1 | 1.6107 |
| *Pantoea sp. strain A4* | *Klebsiella pneumoniae* | Flight 3 | 5 | 6.1947 |
| *Pantoea sp. strain A4* | *Klebsiella pneumoniae* | Flight 3 | 7 | 6.1947 |
| *Pantoea sp. strain A4* | *Klebsiella pneumoniae* | Flight 3 | 8 | 6.1947 |
| *Pantoea sp. strain A4* | *Pantoea agglomerans* | Flight 3 | 1 | 0.0000 |
| *Pantoea sp. strain A4* | *Pantoea agglomerans* | Flight 3 | 5 | 0.0000 |
| *Pantoea sp. strain A4* | *Pantoea agglomerans* | Flight 3 | 7 | 0.0000 |
| *Pantoea sp. strain A4* | *Pantoea agglomerans* | Flight 3 | 8 | 0.0000 |
| *Pantoea sp. strain A4* | *Pantoea ananatis* | Flight 3 | 1 | 0.0000 |
| *Pantoea sp. strain A4* | *Pantoea ananatis* | Flight 3 | 5 | 0.0000 |
| *Pantoea sp. strain A4* | *Pantoea ananatis* | Flight 3 | 7 | 0.0000 |
| *Pantoea sp. strain A4* | *Pantoea ananatis* | Flight 3 | 8 | 0.0000 |
| *Pantoea sp. strain A4* | *Pantoea conspicua* | Flight 3 | 1 | 0.0000 |
| *Pantoea sp. strain A4* | *Pantoea conspicua* | Flight 3 | 5 | 0.0000 |
| *Pantoea sp. strain A4* | *Pantoea conspicua* | Flight 3 | 7 | 0.0000 |
| *Pantoea sp. strain A4* | *Pantoea conspicua* | Flight 3 | 8 | 0.0000 |
| *Pantoea sp. strain A4* | *Pantoea sp. strain At-9b* | Flight 3 | 1 | 0.0000 |
| *Pantoea sp. strain A4* | *Pantoea sp. strain At-9b* | Flight 3 | 5 | 0.0000 |
| *Pantoea sp. strain A4* | *Pantoea sp. strain At-9b* | Flight 3 | 8 | 0.0000 |
| *Pantoea sp. strain A4* | *Pantoea sp. strain FF5* | Flight 3 | 1 | 0.0000 |
| *Pantoea sp. strain A4* | *Pantoea sp. strain FF5* | Flight 3 | 5 | 0.0000 |
| *Pantoea sp. strain A4* | *Pantoea sp. strain FF5* | Flight 3 | 7 | 0.0000 |
| *Pantoea sp. strain A4* | *Pantoea sp. strain FF5* | Flight 3 | 8 | 0.0000 |
| *Pantoea sp. strain A4* | *Pantoea sp. strain IMH* | Flight 3 | 1 | 0.0000 |
| *Pantoea sp. strain A4* | *Pantoea sp. strain IMH* | Flight 3 | 5 | 0.0000 |
| *Pantoea sp. strain A4* | *Pantoea sp. strain IMH* | Flight 3 | 7 | 0.0000 |
| *Pantoea sp. strain A4* | *Pantoea sp. strain IMH* | Flight 3 | 8 | 0.0000 |
| *Pantoea sp. strain A4* | *Pantoea sp. strain NGS-ED-1003* | Flight 3 | 1 | 0.0000 |
| *Pantoea sp. strain A4* | *Pantoea sp. strain NGS-ED-1003* | Flight 3 | 5 | 0.0000 |
| *Pantoea sp. strain A4* | *Pantoea sp. strain NGS-ED-1003* | Flight 3 | 7 | 0.0000 |
| *Pantoea sp. strain A4* | *Pantoea sp. strain NGS-ED-1003* | Flight 3 | 8 | 0.0000 |
| *Pantoea sp. strain A4* | *Pantoea sp. strain OXWO6B1* | Flight 3 | 1 | 0.0000 |
| *Pantoea sp. strain A4* | *Pantoea sp. strain OXWO6B1* | Flight 3 | 5 | 0.0000 |
| *Pantoea sp. strain A4* | *Pantoea sp. strain OXWO6B1* | Flight 3 | 7 | 0.0000 |
| *Pantoea sp. strain A4* | *Pantoea sp. strain OXWO6B1* | Flight 3 | 8 | 0.0000 |
| *Pantoea sp. strain A4* | *Pantoea vagans* | Flight 3 | 1 | 0.0000 |
| *Pantoea sp. strain A4* | *Pantoea vagans* | Flight 3 | 5 | 0.0000 |
| *Pantoea sp. strain A4* | *Pantoea vagans* | Flight 3 | 7 | 0.0000 |
| *Pantoea sp. strain A4* | *Pantoea vagans* | Flight 3 | 8 | 0.0000 |
| *Pantoea sp. strain A4* | *Rhodotorula sp. strain JG-1b* | Flight 3 | 1 | 1.6107 |
| *Pantoea sp. strain A4* | *Salmonella enterica* | Flight 3 | 1 | 1.6107 |
| *Pantoea sp. strain A4* | *Salmonella enterica* | Flight 3 | 5 | 1.5171 |
| *Pantoea sp. strain A4* | *Salmonella enterica* | Flight 3 | 7 | 1.5171 |
| *Pantoea sp. strain A4* | *Salmonella enterica* | Flight 3 | 8 | 1.5171 |
| *Pantoea sp. strain At-9b* | *Aspergillus niger* | Flight 3 | 1 | 0.0000 |
| *Pantoea sp. strain At-9b* | *Enterobacter cloacae* | Flight 3 | 1 | 0.0000 |
| *Pantoea sp. strain At-9b* | *Enterobacter cloacae* | Flight 3 | 5 | 4.4100 |
| *Pantoea sp. strain At-9b* | *Enterobacter cloacae* | Flight 3 | 8 | 4.4100 |
| *Pantoea sp. strain At-9b* | *Escherichia coli* | Flight 3 | 1 | 0.0000 |
| *Pantoea sp. strain At-9b* | *Escherichia coli* | Flight 3 | 5 | 0.0000 |
| *Pantoea sp. strain At-9b* | *Escherichia coli* | Flight 3 | 8 | 0.0000 |
| *Pantoea sp. strain At-9b* | *Klebsiella pneumoniae* | Flight 3 | 1 | 0.0000 |
| *Pantoea sp. strain At-9b* | *Klebsiella pneumoniae* | Flight 3 | 5 | 4.4100 |
| *Pantoea sp. strain At-9b* | *Klebsiella pneumoniae* | Flight 3 | 8 | 4.4100 |
| *Pantoea sp. strain At-9b* | *Pantoea agglomerans* | Flight 3 | 1 | 0.0000 |
| *Pantoea sp. strain At-9b* | *Pantoea agglomerans* | Flight 3 | 5 | 0.0000 |
| *Pantoea sp. strain At-9b* | *Pantoea agglomerans* | Flight 3 | 8 | 0.0000 |
| *Pantoea sp. strain At-9b* | *Pantoea ananatis* | Flight 3 | 1 | 0.0000 |
| *Pantoea sp. strain At-9b* | *Pantoea ananatis* | Flight 3 | 5 | 0.0000 |

| Microorganism | In the presence of | Flight | Location | Metabolic Support Index (A\|AUB) (%) |
|---|---|---|---|---|
| *Pantoea sp. strain At-9b* | *Pantoea ananatis* | Flight 3 | 8 | 0.0000 |
| *Pantoea sp. strain At-9b* | *Pantoea conspicua* | Flight 3 | 1 | 0.0000 |
| *Pantoea sp. strain At-9b* | *Pantoea conspicua* | Flight 3 | 5 | 0.0000 |
| *Pantoea sp. strain At-9b* | *Pantoea conspicua* | Flight 3 | 8 | 0.0000 |
| *Pantoea sp. strain At-9b* | *Pantoea sp. strain A4* | Flight 3 | 1 | 0.0000 |
| *Pantoea sp. strain At-9b* | *Pantoea sp. strain A4* | Flight 3 | 5 | 0.0000 |
| *Pantoea sp. strain At-9b* | *Pantoea sp. strain A4* | Flight 3 | 8 | 0.0000 |
| *Pantoea sp. strain At-9b* | *Pantoea sp. strain FF5* | Flight 3 | 1 | 0.0000 |
| *Pantoea sp. strain At-9b* | *Pantoea sp. strain FF5* | Flight 3 | 5 | 0.0000 |
| *Pantoea sp. strain At-9b* | *Pantoea sp. strain FF5* | Flight 3 | 8 | 0.0000 |
| *Pantoea sp. strain At-9b* | *Pantoea sp. strain IMH* | Flight 3 | 1 | 0.0000 |
| *Pantoea sp. strain At-9b* | *Pantoea sp. strain IMH* | Flight 3 | 5 | 0.0000 |
| *Pantoea sp. strain At-9b* | *Pantoea sp. strain IMH* | Flight 3 | 8 | 0.0000 |
| *Pantoea sp. strain At-9b* | *Pantoea sp. strain NGS-ED-1003* | Flight 3 | 1 | 0.0000 |
| *Pantoea sp. strain At-9b* | *Pantoea sp. strain NGS-ED-1003* | Flight 3 | 5 | 0.0000 |
| *Pantoea sp. strain At-9b* | *Pantoea sp. strain NGS-ED-1003* | Flight 3 | 8 | 0.0000 |
| *Pantoea sp. strain At-9b* | *Pantoea sp. strain OXWO6B1* | Flight 3 | 1 | 0.0000 |
| *Pantoea sp. strain At-9b* | *Pantoea sp. strain OXWO6B1* | Flight 3 | 5 | 0.0000 |
| *Pantoea sp. strain At-9b* | *Pantoea sp. strain OXWO6B1* | Flight 3 | 8 | 0.0000 |
| *Pantoea sp. strain At-9b* | *Pantoea vagans* | Flight 3 | 1 | 0.0000 |
| *Pantoea sp. strain At-9b* | *Pantoea vagans* | Flight 3 | 5 | 0.0000 |
| *Pantoea sp. strain At-9b* | *Pantoea vagans* | Flight 3 | 8 | 0.0000 |
| *Pantoea sp. strain At-9b* | *Rhodotorula sp. strain JG-1b* | Flight 3 | 1 | 0.0000 |
| *Pantoea sp. strain At-9b* | *Salmonella enterica* | Flight 3 | 1 | 0.0000 |
| *Pantoea sp. strain At-9b* | *Salmonella enterica* | Flight 3 | 5 | 0.0000 |
| *Pantoea sp. strain At-9b* | *Salmonella enterica* | Flight 3 | 8 | 0.0000 |
| *Pantoea sp. strain FF5* | *Aspergillus niger* | Flight 3 | 1 | 0.0000 |
| *Pantoea sp. strain FF5* | *Enterobacter cloacae* | Flight 3 | 1 | 0.0000 |
| *Pantoea sp. strain FF5* | *Enterobacter cloacae* | Flight 3 | 5 | 4.6914 |
| *Pantoea sp. strain FF5* | *Enterobacter cloacae* | Flight 3 | 7 | 4.6914 |
| *Pantoea sp. strain FF5* | *Enterobacter cloacae* | Flight 3 | 8 | 4.6914 |
| *Pantoea sp. strain FF5* | *Escherichia coli* | Flight 3 | 1 | 0.7864 |
| *Pantoea sp. strain FF5* | *Escherichia coli* | Flight 3 | 5 | 0.7407 |
| *Pantoea sp. strain FF5* | *Escherichia coli* | Flight 3 | 7 | 0.7407 |
| *Pantoea sp. strain FF5* | *Escherichia coli* | Flight 3 | 8 | 0.7407 |
| *Pantoea sp. strain FF5* | *Klebsiella aerogenes strain IIIF7SW-P1* | Flight 3 | 7 | 5.4321 |
| *Pantoea sp. strain FF5* | *Klebsiella pneumoniae* | Flight 3 | 1 | 0.7864 |
| *Pantoea sp. strain FF5* | *Klebsiella pneumoniae* | Flight 3 | 5 | 5.4321 |
| *Pantoea sp. strain FF5* | *Klebsiella pneumoniae* | Flight 3 | 7 | 5.4321 |
| *Pantoea sp. strain FF5* | *Klebsiella pneumoniae* | Flight 3 | 8 | 5.4321 |
| *Pantoea sp. strain FF5* | *Pantoea agglomerans* | Flight 3 | 1 | 0.0000 |
| *Pantoea sp. strain FF5* | *Pantoea agglomerans* | Flight 3 | 5 | 0.0000 |
| *Pantoea sp. strain FF5* | *Pantoea agglomerans* | Flight 3 | 7 | 0.0000 |
| *Pantoea sp. strain FF5* | *Pantoea agglomerans* | Flight 3 | 8 | 0.0000 |
| *Pantoea sp. strain FF5* | *Pantoea ananatis* | Flight 3 | 1 | 0.0000 |
| *Pantoea sp. strain FF5* | *Pantoea ananatis* | Flight 3 | 5 | 0.0000 |
| *Pantoea sp. strain FF5* | *Pantoea ananatis* | Flight 3 | 7 | 0.0000 |
| *Pantoea sp. strain FF5* | *Pantoea ananatis* | Flight 3 | 8 | 0.0000 |
| *Pantoea sp. strain FF5* | *Pantoea conspicua* | Flight 3 | 1 | 0.7864 |
| *Pantoea sp. strain FF5* | *Pantoea conspicua* | Flight 3 | 5 | 0.7407 |
| *Pantoea sp. strain FF5* | *Pantoea conspicua* | Flight 3 | 7 | 0.7407 |
| *Pantoea sp. strain FF5* | *Pantoea conspicua* | Flight 3 | 8 | 0.7407 |
| *Pantoea sp. strain FF5* | *Pantoea sp. strain A4* | Flight 3 | 1 | 0.0000 |
| *Pantoea sp. strain FF5* | *Pantoea sp. strain A4* | Flight 3 | 5 | 0.0000 |
| *Pantoea sp. strain FF5* | *Pantoea sp. strain A4* | Flight 3 | 7 | 0.0000 |
| *Pantoea sp. strain FF5* | *Pantoea sp. strain A4* | Flight 3 | 8 | 0.0000 |
| *Pantoea sp. strain FF5* | *Pantoea sp. strain At-9b* | Flight 3 | 1 | 0.0000 |
| *Pantoea sp. strain FF5* | *Pantoea sp. strain At-9b* | Flight 3 | 5 | 0.0000 |
| *Pantoea sp. strain FF5* | *Pantoea sp. strain At-9b* | Flight 3 | 8 | 0.0000 |
| *Pantoea sp. strain FF5* | *Pantoea sp. strain IMH* | Flight 3 | 1 | 0.7864 |
| *Pantoea sp. strain FF5* | *Pantoea sp. strain IMH* | Flight 3 | 5 | 0.7407 |
| *Pantoea sp. strain FF5* | *Pantoea sp. strain IMH* | Flight 3 | 7 | 0.7407 |
| *Pantoea sp. strain FF5* | *Pantoea sp. strain IMH* | Flight 3 | 8 | 0.7407 |
| *Pantoea sp. strain FF5* | *Pantoea sp. strain NGS-ED-1003* | Flight 3 | 1 | 0.7864 |
| *Pantoea sp. strain FF5* | *Pantoea sp. strain NGS-ED-1003* | Flight 3 | 5 | 0.7407 |
| *Pantoea sp. strain FF5* | *Pantoea sp. strain NGS-ED-1003* | Flight 3 | 7 | 0.7407 |
| *Pantoea sp. strain FF5* | *Pantoea sp. strain NGS-ED-1003* | Flight 3 | 8 | 0.7407 |
| *Pantoea sp. strain FF5* | *Pantoea sp. strain OXWO6B1* | Flight 3 | 1 | 0.0000 |
| *Pantoea sp. strain FF5* | *Pantoea sp. strain OXWO6B1* | Flight 3 | 5 | 0.0000 |
| *Pantoea sp. strain FF5* | *Pantoea sp. strain OXWO6B1* | Flight 3 | 7 | 0.0000 |
| *Pantoea sp. strain FF5* | *Pantoea sp. strain OXWO6B1* | Flight 3 | 8 | 0.0000 |
| *Pantoea sp. strain FF5* | *Pantoea vagans* | Flight 3 | 1 | 0.0000 |
| *Pantoea sp. strain FF5* | *Pantoea vagans* | Flight 3 | 5 | 0.0000 |
| *Pantoea sp. strain FF5* | *Pantoea vagans* | Flight 3 | 7 | 0.0000 |
| *Pantoea sp. strain FF5* | *Pantoea vagans* | Flight 3 | 8 | 0.0000 |
| *Pantoea sp. strain FF5* | *Rhodotorula sp. strain JG-1b* | Flight 3 | 1 | 0.7864 |
| *Pantoea sp. strain FF5* | *Salmonella enterica* | Flight 3 | 1 | 0.7864 |
| *Pantoea sp. strain FF5* | *Salmonella enterica* | Flight 3 | 5 | 0.7407 |
| *Pantoea sp. strain FF5* | *Salmonella enterica* | Flight 3 | 7 | 0.7407 |
| *Pantoea sp. strain FF5* | *Salmonella enterica* | Flight 3 | 8 | 0.7407 |
| *Pantoea sp. strain IMH* | *Aspergillus niger* | Flight 3 | 1 | 0.0000 |
| *Pantoea sp. strain IMH* | *Enterobacter cloacae* | Flight 3 | 1 | 0.0000 |
| *Pantoea sp. strain IMH* | *Enterobacter cloacae* | Flight 3 | 5 | 5.3901 |
| *Pantoea sp. strain IMH* | *Enterobacter cloacae* | Flight 3 | 7 | 5.3901 |
| *Pantoea sp. strain IMH* | *Enterobacter cloacae* | Flight 3 | 8 | 5.3901 |
| *Pantoea sp. strain IMH* | *Escherichia coli* | Flight 3 | 1 | 0.0000 |
| *Pantoea sp. strain IMH* | *Escherichia coli* | Flight 3 | 5 | 0.0000 |
| *Pantoea sp. strain IMH* | *Escherichia coli* | Flight 3 | 7 | 0.0000 |
| *Pantoea sp. strain IMH* | *Escherichia coli* | Flight 3 | 8 | 0.0000 |
| *Pantoea sp. strain IMH* | *Klebsiella aerogenes strain IIIF7SW-P1* | Flight 3 | 7 | 5.3901 |
| *Pantoea sp. strain IMH* | *Klebsiella pneumoniae* | Flight 3 | 1 | 0.0000 |
| *Pantoea sp. strain IMH* | *Klebsiella pneumoniae* | Flight 3 | 5 | 5.3901 |
| *Pantoea sp. strain IMH* | *Klebsiella pneumoniae* | Flight 3 | 7 | 5.3901 |
| *Pantoea sp. strain IMH* | *Klebsiella pneumoniae* | Flight 3 | 8 | 5.3901 |
| *Pantoea sp. strain IMH* | *Pantoea agglomerans* | Flight 3 | 1 | 0.0000 |
| *Pantoea sp. strain IMH* | *Pantoea agglomerans* | Flight 3 | 5 | 0.0000 |

| Microorganism | In the presence of | Flight | Location | Metabolic Support Index (A\|AUB) (%) |
|---|---|---|---|---|
| *Pantoea sp. strain IMH* | *Pantoea agglomerans* | Flight 3 | 7 | 0.0000 |
| *Pantoea sp. strain IMH* | *Pantoea agglomerans* | Flight 3 | 8 | 0.0000 |
| *Pantoea sp. strain IMH* | *Pantoea ananatis* | Flight 3 | 1 | 0.0000 |
| *Pantoea sp. strain IMH* | *Pantoea ananatis* | Flight 3 | 5 | 0.0000 |
| *Pantoea sp. strain IMH* | *Pantoea ananatis* | Flight 3 | 7 | 0.0000 |
| *Pantoea sp. strain IMH* | *Pantoea ananatis* | Flight 3 | 8 | 0.0000 |
| *Pantoea sp. strain IMH* | *Pantoea conspicua* | Flight 3 | 1 | 0.0000 |
| *Pantoea sp. strain IMH* | *Pantoea conspicua* | Flight 3 | 5 | 0.0000 |
| *Pantoea sp. strain IMH* | *Pantoea conspicua* | Flight 3 | 7 | 0.0000 |
| *Pantoea sp. strain IMH* | *Pantoea conspicua* | Flight 3 | 8 | 0.0000 |
| *Pantoea sp. strain IMH* | *Pantoea sp. strain A4* | Flight 3 | 1 | 0.0000 |
| *Pantoea sp. strain IMH* | *Pantoea sp. strain A4* | Flight 3 | 5 | 0.0000 |
| *Pantoea sp. strain IMH* | *Pantoea sp. strain A4* | Flight 3 | 7 | 0.0000 |
| *Pantoea sp. strain IMH* | *Pantoea sp. strain A4* | Flight 3 | 8 | 0.0000 |
| *Pantoea sp. strain IMH* | *Pantoea sp. strain At-9b* | Flight 3 | 1 | 0.0000 |
| *Pantoea sp. strain IMH* | *Pantoea sp. strain At-9b* | Flight 3 | 5 | 0.0000 |
| *Pantoea sp. strain IMH* | *Pantoea sp. strain At-9b* | Flight 3 | 8 | 0.0000 |
| *Pantoea sp. strain IMH* | *Pantoea sp. strain FF5* | Flight 3 | 1 | 0.0000 |
| *Pantoea sp. strain IMH* | *Pantoea sp. strain FF5* | Flight 3 | 5 | 0.0000 |
| *Pantoea sp. strain IMH* | *Pantoea sp. strain FF5* | Flight 3 | 7 | 0.0000 |
| *Pantoea sp. strain IMH* | *Pantoea sp. strain FF5* | Flight 3 | 8 | 0.0000 |
| *Pantoea sp. strain IMH* | *Pantoea sp. strain NGS-ED-1003* | Flight 3 | 1 | 0.0000 |
| *Pantoea sp. strain IMH* | *Pantoea sp. strain NGS-ED-1003* | Flight 3 | 5 | 0.0000 |
| *Pantoea sp. strain IMH* | *Pantoea sp. strain NGS-ED-1003* | Flight 3 | 7 | 0.0000 |
| *Pantoea sp. strain IMH* | *Pantoea sp. strain NGS-ED-1003* | Flight 3 | 8 | 0.0000 |
| *Pantoea sp. strain IMH* | *Pantoea sp. strain OXWO6B1* | Flight 3 | 1 | 0.0000 |
| *Pantoea sp. strain IMH* | *Pantoea sp. strain OXWO6B1* | Flight 3 | 5 | 0.0000 |
| *Pantoea sp. strain IMH* | *Pantoea sp. strain OXWO6B1* | Flight 3 | 7 | 0.0000 |
| *Pantoea sp. strain IMH* | *Pantoea sp. strain OXWO6B1* | Flight 3 | 8 | 0.0000 |
| *Pantoea sp. strain IMH* | *Pantoea vagans* | Flight 3 | 1 | 0.0000 |
| *Pantoea sp. strain IMH* | *Pantoea vagans* | Flight 3 | 5 | 0.0000 |
| *Pantoea sp. strain IMH* | *Pantoea vagans* | Flight 3 | 7 | 0.0000 |
| *Pantoea sp. strain IMH* | *Pantoea vagans* | Flight 3 | 8 | 0.0000 |
| *Pantoea sp. strain IMH* | *Rhodotorula sp. strain JG-1b* | Flight 3 | 1 | 0.0000 |
| *Pantoea sp. strain IMH* | *Salmonella enterica* | Flight 3 | 1 | 0.0000 |
| *Pantoea sp. strain IMH* | *Salmonella enterica* | Flight 3 | 5 | 0.0000 |
| *Pantoea sp. strain IMH* | *Salmonella enterica* | Flight 3 | 7 | 0.0000 |
| *Pantoea sp. strain IMH* | *Salmonella enterica* | Flight 3 | 8 | 0.0000 |
| *Pantoea sp. strain NGS-ED-1003* | *Aspergillus niger* | Flight 3 | 1 | 0.0000 |
| *Pantoea sp. strain NGS-ED-1003* | *Enterobacter cloacae* | Flight 3 | 1 | 0.0000 |
| *Pantoea sp. strain NGS-ED-1003* | *Enterobacter cloacae* | Flight 3 | 5 | 5.0667 |
| *Pantoea sp. strain NGS-ED-1003* | *Enterobacter cloacae* | Flight 3 | 7 | 5.0667 |
| *Pantoea sp. strain NGS-ED-1003* | *Enterobacter cloacae* | Flight 3 | 8 | 5.0667 |
| *Pantoea sp. strain NGS-ED-1003* | *Escherichia coli* | Flight 3 | 1 | 0.8535 |
| *Pantoea sp. strain NGS-ED-1003* | *Escherichia coli* | Flight 3 | 5 | 0.8000 |
| *Pantoea sp. strain NGS-ED-1003* | *Escherichia coli* | Flight 3 | 7 | 0.8000 |
| *Pantoea sp. strain NGS-ED-1003* | *Escherichia coli* | Flight 3 | 8 | 0.8000 |
| *Pantoea sp. strain NGS-ED-1003* | *Klebsiella aerogenes strain IIIF7SW-P1* | Flight 3 | 7 | 5.8667 |
| *Pantoea sp. strain NGS-ED-1003* | *Klebsiella pneumoniae* | Flight 3 | 1 | 0.8535 |
| *Pantoea sp. strain NGS-ED-1003* | *Klebsiella pneumoniae* | Flight 3 | 5 | 5.8667 |
| *Pantoea sp. strain NGS-ED-1003* | *Klebsiella pneumoniae* | Flight 3 | 7 | 5.8667 |
| *Pantoea sp. strain NGS-ED-1003* | *Klebsiella pneumoniae* | Flight 3 | 8 | 5.8667 |
| *Pantoea sp. strain NGS-ED-1003* | *Pantoea agglomerans* | Flight 3 | 1 | 0.0000 |
| *Pantoea sp. strain NGS-ED-1003* | *Pantoea agglomerans* | Flight 3 | 5 | 0.0000 |
| *Pantoea sp. strain NGS-ED-1003* | *Pantoea agglomerans* | Flight 3 | 7 | 0.0000 |
| *Pantoea sp. strain NGS-ED-1003* | *Pantoea agglomerans* | Flight 3 | 8 | 0.0000 |
| *Pantoea sp. strain NGS-ED-1003* | *Pantoea ananatis* | Flight 3 | 1 | 0.0000 |
| *Pantoea sp. strain NGS-ED-1003* | *Pantoea ananatis* | Flight 3 | 5 | 0.0000 |
| *Pantoea sp. strain NGS-ED-1003* | *Pantoea ananatis* | Flight 3 | 7 | 0.0000 |
| *Pantoea sp. strain NGS-ED-1003* | *Pantoea ananatis* | Flight 3 | 8 | 0.0000 |
| *Pantoea sp. strain NGS-ED-1003* | *Pantoea conspicua* | Flight 3 | 1 | 0.0000 |
| *Pantoea sp. strain NGS-ED-1003* | *Pantoea conspicua* | Flight 3 | 5 | 0.0000 |
| *Pantoea sp. strain NGS-ED-1003* | *Pantoea conspicua* | Flight 3 | 7 | 0.0000 |
| *Pantoea sp. strain NGS-ED-1003* | *Pantoea conspicua* | Flight 3 | 8 | 0.0000 |
| *Pantoea sp. strain NGS-ED-1003* | *Pantoea sp. strain A4* | Flight 3 | 1 | 0.0000 |
| *Pantoea sp. strain NGS-ED-1003* | *Pantoea sp. strain A4* | Flight 3 | 5 | 0.0000 |
| *Pantoea sp. strain NGS-ED-1003* | *Pantoea sp. strain A4* | Flight 3 | 7 | 0.0000 |
| *Pantoea sp. strain NGS-ED-1003* | *Pantoea sp. strain A4* | Flight 3 | 8 | 0.0000 |
| *Pantoea sp. strain NGS-ED-1003* | *Pantoea sp. strain At-9b* | Flight 3 | 1 | 0.0000 |
| *Pantoea sp. strain NGS-ED-1003* | *Pantoea sp. strain At-9b* | Flight 3 | 5 | 0.0000 |
| *Pantoea sp. strain NGS-ED-1003* | *Pantoea sp. strain At-9b* | Flight 3 | 8 | 0.0000 |
| *Pantoea sp. strain NGS-ED-1003* | *Pantoea sp. strain FF5* | Flight 3 | 1 | 0.0000 |
| *Pantoea sp. strain NGS-ED-1003* | *Pantoea sp. strain FF5* | Flight 3 | 5 | 0.0000 |
| *Pantoea sp. strain NGS-ED-1003* | *Pantoea sp. strain FF5* | Flight 3 | 7 | 0.0000 |
| *Pantoea sp. strain NGS-ED-1003* | *Pantoea sp. strain FF5* | Flight 3 | 8 | 0.0000 |
| *Pantoea sp. strain NGS-ED-1003* | *Pantoea sp. strain IMH* | Flight 3 | 1 | 0.0000 |
| *Pantoea sp. strain NGS-ED-1003* | *Pantoea sp. strain IMH* | Flight 3 | 5 | 0.0000 |
| *Pantoea sp. strain NGS-ED-1003* | *Pantoea sp. strain IMH* | Flight 3 | 7 | 0.0000 |
| *Pantoea sp. strain NGS-ED-1003* | *Pantoea sp. strain IMH* | Flight 3 | 8 | 0.0000 |
| *Pantoea sp. strain NGS-ED-1003* | *Pantoea sp. strain OXWO6B1* | Flight 3 | 1 | 0.0000 |
| *Pantoea sp. strain NGS-ED-1003* | *Pantoea sp. strain OXWO6B1* | Flight 3 | 5 | 0.0000 |
| *Pantoea sp. strain NGS-ED-1003* | *Pantoea sp. strain OXWO6B1* | Flight 3 | 7 | 0.0000 |
| *Pantoea sp. strain NGS-ED-1003* | *Pantoea sp. strain OXWO6B1* | Flight 3 | 8 | 0.0000 |
| *Pantoea sp. strain NGS-ED-1003* | *Pantoea vagans* | Flight 3 | 1 | 0.0000 |
| *Pantoea sp. strain NGS-ED-1003* | *Pantoea vagans* | Flight 3 | 5 | 0.0000 |
| *Pantoea sp. strain NGS-ED-1003* | *Pantoea vagans* | Flight 3 | 7 | 0.0000 |
| *Pantoea sp. strain NGS-ED-1003* | *Pantoea vagans* | Flight 3 | 8 | 0.0000 |
| *Pantoea sp. strain NGS-ED-1003* | *Rhodotorula sp. strain JG-1b* | Flight 3 | 1 | 0.8535 |
| *Pantoea sp. strain NGS-ED-1003* | *Salmonella enterica* | Flight 3 | 1 | 0.8535 |
| *Pantoea sp. strain NGS-ED-1003* | *Salmonella enterica* | Flight 3 | 5 | 0.8000 |
| *Pantoea sp. strain NGS-ED-1003* | *Salmonella enterica* | Flight 3 | 7 | 0.8000 |
| *Pantoea sp. strain NGS-ED-1003* | *Salmonella enterica* | Flight 3 | 8 | 0.8000 |
| *Pantoea sp. strain OXWO6B1* | *Aspergillus niger* | Flight 3 | 1 | 0.0000 |
| *Pantoea sp. strain OXWO6B1* | *Enterobacter cloacae* | Flight 3 | 1 | 0.0000 |
| *Pantoea sp. strain OXWO6B1* | *Enterobacter cloacae* | Flight 3 | 5 | 4.9351 |
| *Pantoea sp. strain OXWO6B1* | *Enterobacter cloacae* | Flight 3 | 7 | 4.9351 |

| Microorganism | In the presence of | Flight | Location | Metabolic Support Index (A\|AUB) (%) |
|---|---|---|---|---|
| *Pantoea sp. strain OXWO6B1* | *Enterobacter cloacae* | Flight 3 | 8 | 4.9351 |
| *Pantoea sp. strain OXWO6B1* | *Escherichia coli* | Flight 3 | 1 | 0.0000 |
| *Pantoea sp. strain OXWO6B1* | *Escherichia coli* | Flight 3 | 5 | 0.0000 |
| *Pantoea sp. strain OXWO6B1* | *Escherichia coli* | Flight 3 | 7 | 0.0000 |
| *Pantoea sp. strain OXWO6B1* | *Escherichia coli* | Flight 3 | 8 | 0.0000 |
| *Pantoea sp. strain OXWO6B1* | *Klebsiella aerogenes strain IIIF7SW-P1* | Flight 3 | 7 | 4.9351 |
| *Pantoea sp. strain OXWO6B1* | *Klebsiella pneumoniae* | Flight 3 | 1 | 0.0000 |
| *Pantoea sp. strain OXWO6B1* | *Klebsiella pneumoniae* | Flight 3 | 5 | 4.9351 |
| *Pantoea sp. strain OXWO6B1* | *Klebsiella pneumoniae* | Flight 3 | 7 | 4.9351 |
| *Pantoea sp. strain OXWO6B1* | *Klebsiella pneumoniae* | Flight 3 | 8 | 4.9351 |
| *Pantoea sp. strain OXWO6B1* | *Pantoea agglomerans* | Flight 3 | 1 | 0.0000 |
| *Pantoea sp. strain OXWO6B1* | *Pantoea agglomerans* | Flight 3 | 5 | 0.0000 |
| *Pantoea sp. strain OXWO6B1* | *Pantoea agglomerans* | Flight 3 | 7 | 0.0000 |
| *Pantoea sp. strain OXWO6B1* | *Pantoea agglomerans* | Flight 3 | 8 | 0.0000 |
| *Pantoea sp. strain OXWO6B1* | *Pantoea ananatis* | Flight 3 | 1 | 0.0000 |
| *Pantoea sp. strain OXWO6B1* | *Pantoea ananatis* | Flight 3 | 5 | 0.0000 |
| *Pantoea sp. strain OXWO6B1* | *Pantoea ananatis* | Flight 3 | 7 | 0.0000 |
| *Pantoea sp. strain OXWO6B1* | *Pantoea ananatis* | Flight 3 | 8 | 0.0000 |
| *Pantoea sp. strain OXWO6B1* | *Pantoea conspicua* | Flight 3 | 1 | 0.0000 |
| *Pantoea sp. strain OXWO6B1* | *Pantoea conspicua* | Flight 3 | 5 | 0.0000 |
| *Pantoea sp. strain OXWO6B1* | *Pantoea conspicua* | Flight 3 | 7 | 0.0000 |
| *Pantoea sp. strain OXWO6B1* | *Pantoea conspicua* | Flight 3 | 8 | 0.0000 |
| *Pantoea sp. strain OXWO6B1* | *Pantoea sp. strain A4* | Flight 3 | 1 | 0.0000 |
| *Pantoea sp. strain OXWO6B1* | *Pantoea sp. strain A4* | Flight 3 | 5 | 0.0000 |
| *Pantoea sp. strain OXWO6B1* | *Pantoea sp. strain A4* | Flight 3 | 7 | 0.0000 |
| *Pantoea sp. strain OXWO6B1* | *Pantoea sp. strain A4* | Flight 3 | 8 | 0.0000 |
| *Pantoea sp. strain OXWO6B1* | *Pantoea sp. strain At-9b* | Flight 3 | 1 | 0.0000 |
| *Pantoea sp. strain OXWO6B1* | *Pantoea sp. strain At-9b* | Flight 3 | 5 | 0.0000 |
| *Pantoea sp. strain OXWO6B1* | *Pantoea sp. strain At-9b* | Flight 3 | 8 | 0.0000 |
| *Pantoea sp. strain OXWO6B1* | *Pantoea sp. strain FF5* | Flight 3 | 1 | 0.0000 |
| *Pantoea sp. strain OXWO6B1* | *Pantoea sp. strain FF5* | Flight 3 | 5 | 0.0000 |
| *Pantoea sp. strain OXWO6B1* | *Pantoea sp. strain FF5* | Flight 3 | 7 | 0.0000 |
| *Pantoea sp. strain OXWO6B1* | *Pantoea sp. strain FF5* | Flight 3 | 8 | 0.0000 |
| *Pantoea sp. strain OXWO6B1* | *Pantoea sp. strain IMH* | Flight 3 | 1 | 0.0000 |
| *Pantoea sp. strain OXWO6B1* | *Pantoea sp. strain IMH* | Flight 3 | 5 | 0.0000 |
| *Pantoea sp. strain OXWO6B1* | *Pantoea sp. strain IMH* | Flight 3 | 7 | 0.0000 |
| *Pantoea sp. strain OXWO6B1* | *Pantoea sp. strain IMH* | Flight 3 | 8 | 0.0000 |
| *Pantoea sp. strain OXWO6B1* | *Pantoea sp. strain NGS-ED-1003* | Flight 3 | 1 | 0.0000 |
| *Pantoea sp. strain OXWO6B1* | *Pantoea sp. strain NGS-ED-1003* | Flight 3 | 5 | 0.0000 |
| *Pantoea sp. strain OXWO6B1* | *Pantoea sp. strain NGS-ED-1003* | Flight 3 | 7 | 0.0000 |
| *Pantoea sp. strain OXWO6B1* | *Pantoea sp. strain NGS-ED-1003* | Flight 3 | 8 | 0.0000 |
| *Pantoea sp. strain OXWO6B1* | *Pantoea vagans* | Flight 3 | 1 | 0.0000 |
| *Pantoea sp. strain OXWO6B1* | *Pantoea vagans* | Flight 3 | 5 | 0.0000 |
| *Pantoea sp. strain OXWO6B1* | *Pantoea vagans* | Flight 3 | 7 | 0.0000 |
| *Pantoea sp. strain OXWO6B1* | *Pantoea vagans* | Flight 3 | 8 | 0.0000 |
| *Pantoea sp. strain OXWO6B1* | *Rhodotorula sp. strain JG-1b* | Flight 3 | 1 | 0.0000 |
| *Pantoea sp. strain OXWO6B1* | *Salmonella enterica* | Flight 3 | 1 | 0.0000 |
| *Pantoea sp. strain OXWO6B1* | *Salmonella enterica* | Flight 3 | 5 | 0.0000 |
| *Pantoea sp. strain OXWO6B1* | *Salmonella enterica* | Flight 3 | 7 | 0.0000 |
| *Pantoea sp. strain OXWO6B1* | *Salmonella enterica* | Flight 3 | 8 | 0.0000 |
| *Pantoea vagans* | *Aspergillus niger* | Flight 3 | 1 | 0.0000 |
| *Pantoea vagans* | *Enterobacter cloacae* | Flight 3 | 1 | 0.0000 |
| *Pantoea vagans* | *Enterobacter cloacae* | Flight 3 | 5 | 4.9268 |
| *Pantoea vagans* | *Enterobacter cloacae* | Flight 3 | 7 | 4.9268 |
| *Pantoea vagans* | *Enterobacter cloacae* | Flight 3 | 8 | 4.9268 |
| *Pantoea vagans* | *Escherichia coli* | Flight 2 | 5 | 0.7989 |
| *Pantoea vagans* | *Escherichia coli* | Flight 3 | 1 | 0.8487 |
| *Pantoea vagans* | *Escherichia coli* | Flight 3 | 5 | 0.7989 |
| *Pantoea vagans* | *Escherichia coli* | Flight 3 | 7 | 0.7989 |
| *Pantoea vagans* | *Escherichia coli* | Flight 3 | 8 | 0.7989 |
| *Pantoea vagans* | *Klebsiella aerogenes strain IIIF7SW-P1* | Flight 3 | 7 | 5.7257 |
| *Pantoea vagans* | *Klebsiella pneumoniae* | Flight 2 | 5 | 5.7257 |
| *Pantoea vagans* | *Klebsiella pneumoniae* | Flight 3 | 1 | 0.8487 |
| *Pantoea vagans* | *Klebsiella pneumoniae* | Flight 3 | 5 | 5.7257 |
| *Pantoea vagans* | *Klebsiella pneumoniae* | Flight 3 | 7 | 5.7257 |
| *Pantoea vagans* | *Klebsiella pneumoniae* | Flight 3 | 8 | 5.7257 |
| *Pantoea vagans* | *Paenibacillus polymyxa strain IIF5SW-B3* | Flight 2 | 5 | 4.9268 |
| *Pantoea vagans* | *Paenibacillus polymyxa strain IIF5SW-B4* | Flight 2 | 5 | 4.9268 |
| *Pantoea vagans* | *Pantoea agglomerans* | Flight 2 | 5 | 0.0000 |
| *Pantoea vagans* | *Pantoea agglomerans* | Flight 3 | 1 | 0.0000 |
| *Pantoea vagans* | *Pantoea agglomerans* | Flight 3 | 5 | 0.0000 |
| *Pantoea vagans* | *Pantoea agglomerans* | Flight 3 | 7 | 0.0000 |
| *Pantoea vagans* | *Pantoea agglomerans* | Flight 3 | 8 | 0.0000 |
| *Pantoea vagans* | *Pantoea ananatis* | Flight 2 | 5 | 0.0000 |
| *Pantoea vagans* | *Pantoea ananatis* | Flight 3 | 1 | 0.0000 |
| *Pantoea vagans* | *Pantoea ananatis* | Flight 3 | 5 | 0.0000 |
| *Pantoea vagans* | *Pantoea ananatis* | Flight 3 | 7 | 0.0000 |
| *Pantoea vagans* | *Pantoea ananatis* | Flight 3 | 8 | 0.0000 |
| *Pantoea vagans* | *Pantoea conspicua* | Flight 2 | 5 | 0.7989 |
| *Pantoea vagans* | *Pantoea conspicua* | Flight 3 | 1 | 0.8487 |
| *Pantoea vagans* | *Pantoea conspicua* | Flight 3 | 5 | 0.7989 |
| *Pantoea vagans* | *Pantoea conspicua* | Flight 3 | 7 | 0.7989 |
| *Pantoea vagans* | *Pantoea conspicua* | Flight 3 | 8 | 0.7989 |
| *Pantoea vagans* | *Pantoea sp. strain 3.5.1* | Flight 2 | 5 | 0.7989 |
| *Pantoea vagans* | *Pantoea sp. strain A4* | Flight 3 | 1 | 0.0000 |
| *Pantoea vagans* | *Pantoea sp. strain A4* | Flight 3 | 5 | 0.0000 |
| *Pantoea vagans* | *Pantoea sp. strain A4* | Flight 3 | 7 | 0.0000 |
| *Pantoea vagans* | *Pantoea sp. strain A4* | Flight 3 | 8 | 0.0000 |
| *Pantoea vagans* | *Pantoea sp. strain At-9b* | Flight 3 | 1 | 0.0000 |
| *Pantoea vagans* | *Pantoea sp. strain At-9b* | Flight 3 | 5 | 0.0000 |
| *Pantoea vagans* | *Pantoea sp. strain At-9b* | Flight 3 | 8 | 0.0000 |
| *Pantoea vagans* | *Pantoea sp. strain FF5* | Flight 3 | 1 | 0.0000 |
| *Pantoea vagans* | *Pantoea sp. strain FF5* | Flight 3 | 5 | 0.0000 |
| *Pantoea vagans* | *Pantoea sp. strain FF5* | Flight 3 | 7 | 0.0000 |
| *Pantoea vagans* | *Pantoea sp. strain FF5* | Flight 3 | 8 | 0.0000 |
| *Pantoea vagans* | *Pantoea sp. strain IMH* | Flight 3 | 1 | 0.8487 |

| Microorganism | In the presence of | Flight | Location | Metabolic Support Index (A\|AUB) (%) |
|---|---|---|---|---|
| *Pantoea vagans* | *Pantoea sp. strain IMH* | Flight 3 | 5 | 0.7989 |
| *Pantoea vagans* | *Pantoea sp. strain IMH* | Flight 3 | 7 | 0.7989 |
| *Pantoea vagans* | *Pantoea sp. strain IMH* | Flight 3 | 8 | 0.7989 |
| *Pantoea vagans* | *Pantoea sp. strain NGS-ED-1003* | Flight 3 | 1 | 0.8487 |
| *Pantoea vagans* | *Pantoea sp. strain NGS-ED-1003* | Flight 3 | 5 | 0.7989 |
| *Pantoea vagans* | *Pantoea sp. strain NGS-ED-1003* | Flight 3 | 7 | 0.7989 |
| *Pantoea vagans* | *Pantoea sp. strain NGS-ED-1003* | Flight 3 | 8 | 0.7989 |
| *Pantoea vagans* | *Pantoea sp. strain OXWO6B1* | Flight 3 | 1 | 0.0000 |
| *Pantoea vagans* | *Pantoea sp. strain OXWO6B1* | Flight 3 | 5 | 0.0000 |
| *Pantoea vagans* | *Pantoea sp. strain OXWO6B1* | Flight 3 | 7 | 0.0000 |
| *Pantoea vagans* | *Pantoea sp. strain OXWO6B1* | Flight 3 | 8 | 0.0000 |
| *Pantoea vagans* | *Rhodotorula sp. strain JG-1b* | Flight 3 | 1 | 0.8487 |
| *Pantoea vagans* | *Salmonella enterica* | Flight 3 | 1 | 0.8487 |
| *Pantoea vagans* | *Salmonella enterica* | Flight 3 | 5 | 0.7989 |
| *Pantoea vagans* | *Salmonella enterica* | Flight 3 | 7 | 0.7989 |
| *Pantoea vagans* | *Salmonella enterica* | Flight 3 | 8 | 0.7989 |
| *Penicillium chrysogenum* | *Aspergillus niger* | Flight 1 | 1 | 0.0000 |
| *Penicillium chrysogenum* | *Enterococcus avium* | Flight 3 | 2 | 0.7874 |
| *Penicillium chrysogenum* | *Enterococcus faecalis* | Flight 3 | 2 | 0.7874 |
| *Penicillium chrysogenum* | *Klebsiella pneumoniae* | Flight 1 | 1 | 0.6418 |
| *Penicillium chrysogenum* | *Klebsiella pneumoniae strain F3-2P(2*)* | Flight 3 | 2 | 0.7874 |
| *Penicillium chrysogenum* | *Klebsiella quasipneumoniae strain IF1SW-B2* | Flight 1 | 1 | 0.6418 |
| *Penicillium chrysogenum* | *Klebsiella quasipneumoniae strain IF1SW-P3* | Flight 1 | 1 | 0.6418 |
| *Penicillium chrysogenum* | *Klebsiella quasipneumoniae strain IF1SW-P4* | Flight 1 | 1 | 0.6418 |
| *Penicillium chrysogenum* | *Paenibacillus polymyxa* | Flight 3 | 2 | 0.7874 |
| *Penicillium chrysogenum* | *Penicillium flavigenum* | Flight 3 | 2 | 0.0000 |
| *Penicillium chrysogenum* | *Penicillium nalgiovense* | Flight 3 | 2 | 0.0000 |
| *Penicillium chrysogenum* | *Penicillium rubens* | Flight 1 | 1 | 0.0000 |
| *Penicillium chrysogenum* | *Penicillium rubens* | Flight 3 | 2 | 0.0000 |
| *Penicillium chrysogenum* | *Rhodotorula sp. strain JG-1b* | Flight 1 | 1 | 0.0000 |
| *Penicillium chrysogenum* | *Rhodotorula sp. strain JG-1b* | Flight 3 | 2 | 0.0000 |
| *Penicillium chrysogenum* | *Rhodotorula toruloides* | Flight 1 | 1 | 0.0000 |
| *Penicillium chrysogenum* | *Staphylococcus aureus* | Flight 3 | 2 | 0.7874 |
| *Penicillium chrysogenum* | *Staphylococcus epidermidis* | Flight 3 | 2 | 0.7874 |
| *Penicillium chrysogenum* | *Staphylococcus haemolyticus* | Flight 3 | 2 | 0.7874 |
| *Penicillium chrysogenum* | *Staphylococcus saprophyticus* | Flight 3 | 2 | 0.7874 |
| *Penicillium chrysogenum* | *Staphylococcus sp. strain LCT-H4* | Flight 3 | 2 | 0.7874 |
| *Penicillium chrysogenum* | *Staphylococcus warneri* | Flight 3 | 2 | 0.0000 |
| *Penicillium flavigenum* | *Enterococcus avium* | Flight 3 | 2 | 0.6342 |
| *Penicillium flavigenum* | *Enterococcus faecalis* | Flight 3 | 2 | 0.6342 |
| *Penicillium flavigenum* | *Klebsiella pneumoniae strain F3-2P(2*)* | Flight 3 | 2 | 0.6342 |
| *Penicillium flavigenum* | *Paenibacillus polymyxa* | Flight 3 | 2 | 0.6342 |
| *Penicillium flavigenum* | *Penicillium chrysogenum* | Flight 3 | 2 | 0.0000 |
| *Penicillium flavigenum* | *Penicillium nalgiovense* | Flight 3 | 2 | 0.0000 |
| *Penicillium flavigenum* | *Penicillium rubens* | Flight 3 | 2 | 0.0000 |
| *Penicillium flavigenum* | *Rhodotorula sp. strain JG-1b* | Flight 3 | 2 | 0.0000 |
| *Penicillium flavigenum* | *Staphylococcus aureus* | Flight 3 | 2 | 0.6342 |
| *Penicillium flavigenum* | *Staphylococcus epidermidis* | Flight 3 | 2 | 0.6342 |
| *Penicillium flavigenum* | *Staphylococcus haemolyticus* | Flight 3 | 2 | 0.6342 |
| *Penicillium flavigenum* | *Staphylococcus saprophyticus* | Flight 3 | 2 | 0.6342 |
| *Penicillium flavigenum* | *Staphylococcus sp. strain LCT-H4* | Flight 3 | 2 | 0.6342 |
| *Penicillium flavigenum* | *Staphylococcus warneri* | Flight 3 | 2 | 0.0000 |
| *Penicillium nalgiovense* | *Enterococcus avium* | Flight 3 | 2 | 0.7764 |
| *Penicillium nalgiovense* | *Enterococcus faecalis* | Flight 3 | 2 | 0.7764 |
| *Penicillium nalgiovense* | *Klebsiella pneumoniae strain F3-2P(2*)* | Flight 3 | 2 | 0.7764 |
| *Penicillium nalgiovense* | *Paenibacillus polymyxa* | Flight 3 | 2 | 0.7764 |
| *Penicillium nalgiovense* | *Penicillium chrysogenum* | Flight 3 | 2 | 0.0000 |
| *Penicillium nalgiovense* | *Penicillium flavigenum* | Flight 3 | 2 | 0.0000 |
| *Penicillium nalgiovense* | *Penicillium rubens* | Flight 3 | 2 | 0.0000 |
| *Penicillium nalgiovense* | *Rhodotorula sp. strain JG-1b* | Flight 3 | 2 | 0.0000 |
| *Penicillium nalgiovense* | *Staphylococcus aureus* | Flight 3 | 2 | 0.7764 |
| *Penicillium nalgiovense* | *Staphylococcus epidermidis* | Flight 3 | 2 | 0.7764 |
| *Penicillium nalgiovense* | *Staphylococcus haemolyticus* | Flight 3 | 2 | 0.7764 |
| *Penicillium nalgiovense* | *Staphylococcus saprophyticus* | Flight 3 | 2 | 0.7764 |
| *Penicillium nalgiovense* | *Staphylococcus sp. strain LCT-H4* | Flight 3 | 2 | 0.7764 |
| *Penicillium nalgiovense* | *Staphylococcus warneri* | Flight 3 | 2 | 0.0000 |
| *Penicillium rubens* | *Aspergillus niger* | Flight 1 | 1 | 0.0000 |
| *Penicillium rubens* | *Enterobacter asburiae* | Flight 1 | 2 | 0.6394 |
| *Penicillium rubens* | *Enterobacter cancerogenus* | Flight 1 | 2 | 0.6394 |
| *Penicillium rubens* | *Enterobacter cloacae* | Flight 1 | 2 | 0.6394 |
| *Penicillium rubens* | *Enterobacter hormaechei* | Flight 1 | 2 | 0.6394 |
| *Penicillium rubens* | *Enterobacter roggenkampii* | Flight 1 | 2 | 0.6394 |
| *Penicillium rubens* | *Enterobacter sp. strain NFIX59* | Flight 1 | 2 | 0.6394 |
| *Penicillium rubens* | *Enterococcus avium* | Flight 3 | 2 | 0.7837 |
| *Penicillium rubens* | *Enterococcus faecalis* | Flight 3 | 2 | 0.7837 |
| *Penicillium rubens* | *Escherichia coli* | Flight 1 | 2 | 0.6394 |
| *Penicillium rubens* | *Escherichia coli* | Flight 1 | 5 | 0.6394 |
| *Penicillium rubens* | *Escherichia coli* | Flight 3 | 3 | 0.6361 |
| *Penicillium rubens* | *Klebsiella oxytoca* | Flight 1 | 2 | 0.6394 |
| *Penicillium rubens* | *Klebsiella pneumoniae* | Flight 1 | 1 | 0.6394 |
| *Penicillium rubens* | *Klebsiella pneumoniae* | Flight 1 | 2 | 0.6394 |
| *Penicillium rubens* | *Klebsiella pneumoniae* | Flight 1 | 5 | 0.6394 |
| *Penicillium rubens* | *Klebsiella pneumoniae* | Flight 3 | 3 | 0.6361 |
| *Penicillium rubens* | *Klebsiella pneumoniae strain F3-2P(2*)* | Flight 3 | 2 | 0.7837 |
| *Penicillium rubens* | *Klebsiella quasipneumoniae strain IF1SW-B2* | Flight 1 | 1 | 0.6394 |
| *Penicillium rubens* | *Klebsiella quasipneumoniae strain IF1SW-P3* | Flight 1 | 1 | 0.6394 |
| *Penicillium rubens* | *Klebsiella quasipneumoniae strain IF1SW-P4* | Flight 1 | 1 | 0.6394 |
| *Penicillium rubens* | *Klebsiella quasipneumoniae strain IF2SW-B3* | Flight 1 | 2 | 0.6394 |
| *Penicillium rubens* | *Klebsiella quasipneumoniae strain IF2SW-P1* | Flight 1 | 2 | 0.6394 |
| *Penicillium rubens* | *Klebsiella quasipneumoniae strain IIIF3SW-P1* | Flight 3 | 3 | 0.6361 |
| *Penicillium rubens* | *Klebsiella sp. strain MS 92-3* | Flight 3 | 3 | 0.6361 |
| *Penicillium rubens* | *Klebsiella variicola* | Flight 3 | 3 | 0.6361 |
| *Penicillium rubens* | *Paenibacillus polymyxa* | Flight 3 | 2 | 0.7837 |
| *Penicillium rubens* | *Pantoea agglomerans* | Flight 1 | 5 | 0.6394 |
| *Penicillium rubens* | *Pantoea ananatis* | Flight 1 | 2 | 0.6394 |
| *Penicillium rubens* | *Pantoea ananatis* | Flight 1 | 5 | 0.6394 |

| Microorganism | In the presence of | Flight | Location | Metabolic Support Index (A\|AUB) (%) |
|---|---|---|---|---|
| *Penicillium rubens* | *Pantoea conspicua* | Flight 1 | 5 | 0.6394 |
| *Penicillium rubens* | *Pantoea sp. strain 3.5.1* | Flight 1 | 5 | 0.6394 |
| *Penicillium rubens* | *Penicillium chrysogenum* | Flight 1 | 1 | 0.0000 |
| *Penicillium rubens* | *Penicillium chrysogenum* | Flight 3 | 2 | 0.0000 |
| *Penicillium rubens* | *Penicillium flavigenum* | Flight 3 | 2 | 0.0000 |
| *Penicillium rubens* | *Penicillium nalgiovense* | Flight 3 | 2 | 0.0000 |
| *Penicillium rubens* | *Rhodotorula sp. strain JG-1b* | Flight 1 | 1 | 0.0000 |
| *Penicillium rubens* | *Rhodotorula sp. strain JG-1b* | Flight 1 | 2 | 0.0000 |
| *Penicillium rubens* | *Rhodotorula sp. strain JG-1b* | Flight 1 | 5 | 0.0000 |
| *Penicillium rubens* | *Rhodotorula sp. strain JG-1b* | Flight 3 | 2 | 0.0000 |
| *Penicillium rubens* | *Rhodotorula toruloides* | Flight 1 | 1 | 0.0000 |
| *Penicillium rubens* | *Salmonella enterica* | Flight 1 | 2 | 0.6394 |
| *Penicillium rubens* | *Salmonella enterica* | Flight 3 | 3 | 0.6361 |
| *Penicillium rubens* | *Shigella sonnei* | Flight 1 | 2 | 0.6394 |
| *Penicillium rubens* | *Staphylococcus aureus* | Flight 3 | 2 | 0.7837 |
| *Penicillium rubens* | *Staphylococcus epidermidis* | Flight 3 | 2 | 0.7837 |
| *Penicillium rubens* | *Staphylococcus haemolyticus* | Flight 3 | 2 | 0.7837 |
| *Penicillium rubens* | *Staphylococcus saprophyticus* | Flight 3 | 2 | 0.7837 |
| *Penicillium rubens* | *Staphylococcus saprophyticus* | Flight 3 | 3 | 0.0000 |
| *Penicillium rubens* | *Staphylococcus sp. strain LCT-H4* | Flight 3 | 2 | 0.7837 |
| *Penicillium rubens* | *Staphylococcus warneri* | Flight 3 | 2 | 0.0000 |
| *Rahnella aquatilis* | *Enterobacter hormaechei* | Flight 3 | 4 | 5.0870 |
| *Rahnella aquatilis* | *Escherichia coli* | Flight 3 | 4 | 0.0000 |
| *Rahnella aquatilis* | *Klebsiella pneumoniae* | Flight 3 | 4 | 5.0870 |
| *Rahnella aquatilis* | *Pantoea ananatis* | Flight 3 | 4 | 0.0000 |
| *Rahnella aquatilis* | *Pantoea dispersa* | Flight 3 | 4 | 0.0000 |
| *Rhodotorula sp. strain JG-1b* | *Aspergillus niger* | Flight 1 | 1 | 0.0000 |
| *Rhodotorula sp. strain JG-1b* | *Aspergillus niger* | Flight 3 | 1 | 0.0000 |
| *Rhodotorula sp. strain JG-1b* | *Enterobacter asburiae* | Flight 1 | 2 | 0.2817 |
| *Rhodotorula sp. strain JG-1b* | *Enterobacter cancerogenus* | Flight 1 | 2 | 0.2817 |
| *Rhodotorula sp. strain JG-1b* | *Enterobacter cloacae* | Flight 1 | 2 | 0.2817 |
| *Rhodotorula sp. strain JG-1b* | *Enterobacter cloacae* | Flight 3 | 1 | 0.2793 |
| *Rhodotorula sp. strain JG-1b* | *Enterobacter hormaechei* | Flight 1 | 2 | 0.2817 |
| *Rhodotorula sp. strain JG-1b* | *Enterobacter roggenkampii* | Flight 1 | 2 | 0.2817 |
| *Rhodotorula sp. strain JG-1b* | *Enterobacter sp. strain NFIX59* | Flight 1 | 2 | 0.2817 |
| *Rhodotorula sp. strain JG-1b* | *Enterococcus avium* | Flight 3 | 2 | 0.2874 |
| *Rhodotorula sp. strain JG-1b* | *Enterococcus faecalis* | Flight 3 | 2 | 0.2874 |
| *Rhodotorula sp. strain JG-1b* | *Escherichia coli* | Flight 1 | 2 | 0.2817 |
| *Rhodotorula sp. strain JG-1b* | *Escherichia coli* | Flight 1 | 5 | 0.2817 |
| *Rhodotorula sp. strain JG-1b* | *Escherichia coli* | Flight 3 | 1 | 0.2793 |
| *Rhodotorula sp. strain JG-1b* | *Klebsiella oxytoca* | Flight 1 | 2 | 0.2817 |
| *Rhodotorula sp. strain JG-1b* | *Klebsiella pneumoniae* | Flight 1 | 1 | 0.2817 |
| *Rhodotorula sp. strain JG-1b* | *Klebsiella pneumoniae* | Flight 1 | 2 | 0.2817 |
| *Rhodotorula sp. strain JG-1b* | *Klebsiella pneumoniae* | Flight 1 | 5 | 0.2817 |
| *Rhodotorula sp. strain JG-1b* | *Klebsiella pneumoniae* | Flight 3 | 1 | 0.2793 |
| *Rhodotorula sp. strain JG-1b* | *Klebsiella pneumoniae strain F3-2P(2*)* | Flight 3 | 2 | 0.2874 |
| *Rhodotorula sp. strain JG-1b* | *Klebsiella quasipneumoniae strain IF1SW-B2* | Flight 1 | 1 | 0.2817 |
| *Rhodotorula sp. strain JG-1b* | *Klebsiella quasipneumoniae strain IF1SW-P3* | Flight 1 | 1 | 0.2817 |
| *Rhodotorula sp. strain JG-1b* | *Klebsiella quasipneumoniae strain IF1SW-P4* | Flight 1 | 1 | 0.2817 |
| *Rhodotorula sp. strain JG-1b* | *Klebsiella quasipneumoniae strain IF2SW-B3* | Flight 1 | 2 | 0.2817 |
| *Rhodotorula sp. strain JG-1b* | *Klebsiella quasipneumoniae strain IF2SW-P1* | Flight 1 | 2 | 0.2817 |
| *Rhodotorula sp. strain JG-1b* | *Paenibacillus polymyxa* | Flight 3 | 2 | 0.2874 |
| *Rhodotorula sp. strain JG-1b* | *Pantoea agglomerans* | Flight 1 | 5 | 0.2817 |
| *Rhodotorula sp. strain JG-1b* | *Pantoea agglomerans* | Flight 3 | 1 | 0.2793 |
| *Rhodotorula sp. strain JG-1b* | *Pantoea ananatis* | Flight 1 | 2 | 0.2817 |
| *Rhodotorula sp. strain JG-1b* | *Pantoea ananatis* | Flight 1 | 5 | 0.2817 |
| *Rhodotorula sp. strain JG-1b* | *Pantoea ananatis* | Flight 3 | 1 | 0.2793 |
| *Rhodotorula sp. strain JG-1b* | *Pantoea conspicua* | Flight 1 | 5 | 0.2817 |
| *Rhodotorula sp. strain JG-1b* | *Pantoea conspicua* | Flight 3 | 1 | 0.2793 |
| *Rhodotorula sp. strain JG-1b* | *Pantoea sp. strain 3.5.1* | Flight 1 | 5 | 0.2817 |
| *Rhodotorula sp. strain JG-1b* | *Pantoea sp. strain A4* | Flight 3 | 1 | 0.2793 |
| *Rhodotorula sp. strain JG-1b* | *Pantoea sp. strain At-9b* | Flight 3 | 1 | 0.2793 |
| *Rhodotorula sp. strain JG-1b* | *Pantoea sp. strain FF5* | Flight 3 | 1 | 0.2793 |
| *Rhodotorula sp. strain JG-1b* | *Pantoea sp. strain IMH* | Flight 3 | 1 | 0.2793 |
| *Rhodotorula sp. strain JG-1b* | *Pantoea sp. strain NGS-ED-1003* | Flight 3 | 1 | 0.2793 |
| *Rhodotorula sp. strain JG-1b* | *Pantoea sp. strain OXWO6B1* | Flight 3 | 1 | 0.2793 |
| *Rhodotorula sp. strain JG-1b* | *Pantoea vagans* | Flight 3 | 1 | 0.2793 |
| *Rhodotorula sp. strain JG-1b* | *Penicillium chrysogenum* | Flight 1 | 1 | 0.2817 |
| *Rhodotorula sp. strain JG-1b* | *Penicillium chrysogenum* | Flight 3 | 2 | 0.2874 |
| *Rhodotorula sp. strain JG-1b* | *Penicillium flavigenum* | Flight 3 | 2 | 0.2874 |
| *Rhodotorula sp. strain JG-1b* | *Penicillium nalgiovense* | Flight 3 | 2 | 0.2874 |
| *Rhodotorula sp. strain JG-1b* | *Penicillium rubens* | Flight 1 | 1 | 0.2817 |
| *Rhodotorula sp. strain JG-1b* | *Penicillium rubens* | Flight 1 | 2 | 0.2817 |
| *Rhodotorula sp. strain JG-1b* | *Penicillium rubens* | Flight 1 | 5 | 0.2817 |
| *Rhodotorula sp. strain JG-1b* | *Penicillium rubens* | Flight 3 | 2 | 0.2874 |
| *Rhodotorula sp. strain JG-1b* | *Rhodotorula toruloides* | Flight 1 | 1 | 0.0000 |
| *Rhodotorula sp. strain JG-1b* | *Salmonella enterica* | Flight 1 | 2 | 0.2817 |
| *Rhodotorula sp. strain JG-1b* | *Salmonella enterica* | Flight 3 | 1 | 0.2793 |
| *Rhodotorula sp. strain JG-1b* | *Shigella sonnei* | Flight 1 | 2 | 0.2817 |
| *Rhodotorula sp. strain JG-1b* | *Staphylococcus aureus* | Flight 3 | 2 | 0.2874 |
| *Rhodotorula sp. strain JG-1b* | *Staphylococcus epidermidis* | Flight 3 | 2 | 0.2874 |
| *Rhodotorula sp. strain JG-1b* | *Staphylococcus haemolyticus* | Flight 3 | 2 | 0.2874 |
| *Rhodotorula sp. strain JG-1b* | *Staphylococcus saprophyticus* | Flight 3 | 2 | 0.2874 |
| *Rhodotorula sp. strain JG-1b* | *Staphylococcus sp. strain LCT-H4* | Flight 3 | 2 | 0.2874 |
| *Rhodotorula sp. strain JG-1b* | *Staphylococcus warneri* | Flight 3 | 2 | 0.2874 |
| *Rhodotorula toruloides* | *Aspergillus niger* | Flight 1 | 1 | 0.5333 |
| *Rhodotorula toruloides* | *Klebsiella pneumoniae* | Flight 1 | 1 | 0.8000 |
| *Rhodotorula toruloides* | *Klebsiella quasipneumoniae strain IF1SW-B2* | Flight 1 | 1 | 0.8000 |
| *Rhodotorula toruloides* | *Klebsiella quasipneumoniae strain IF1SW-P3* | Flight 1 | 1 | 0.8000 |
| *Rhodotorula toruloides* | *Klebsiella quasipneumoniae strain IF1SW-P4* | Flight 1 | 1 | 0.8000 |
| *Rhodotorula toruloides* | *Penicillium chrysogenum* | Flight 1 | 1 | 0.5333 |
| *Rhodotorula toruloides* | *Penicillium rubens* | Flight 1 | 1 | 0.5333 |
| *Rhodotorula toruloides* | *Rhodotorula sp. strain JG-1b* | Flight 1 | 1 | 0.0000 |
| *Salmonella enterica* | *Aspergillus niger* | Flight 3 | 1 | 0.0000 |
| *Salmonella enterica* | *Enterobacter asburiae* | Flight 1 | 2 | 0.0000 |
| *Salmonella enterica* | *Enterobacter cancerogenus* | Flight 1 | 2 | 0.0000 |

| Microorganism | In the presence of | Flight | Location | Metabolic Support Index (A\|AUB) (%) |
|---|---|---|---|---|
| *Salmonella enterica* | *Enterobacter cloacae* | Flight 1 | 2 | 0.0000 |
| *Salmonella enterica* | *Enterobacter cloacae* | Flight 3 | 1 | 0.0000 |
| *Salmonella enterica* | *Enterobacter cloacae* | Flight 3 | 5 | 4.9080 |
| *Salmonella enterica* | *Enterobacter cloacae* | Flight 3 | 7 | 4.9080 |
| *Salmonella enterica* | *Enterobacter cloacae* | Flight 3 | 8 | 4.9080 |
| *Salmonella enterica* | *Enterobacter hormaechei* | Flight 1 | 2 | 0.0000 |
| *Salmonella enterica* | *Enterobacter roggenkampii* | Flight 1 | 2 | 0.0000 |
| *Salmonella enterica* | *Enterobacter sp. strain NFIX59* | Flight 1 | 2 | 0.0000 |
| *Salmonella enterica* | *Escherichia coli* | Flight 1 | 2 | 0.0000 |
| *Salmonella enterica* | *Escherichia coli* | Flight 3 | 1 | 0.0000 |
| *Salmonella enterica* | *Escherichia coli* | Flight 3 | 3 | 0.0000 |
| *Salmonella enterica* | *Escherichia coli* | Flight 3 | 5 | 0.0000 |
| *Salmonella enterica* | *Escherichia coli* | Flight 3 | 7 | 0.0000 |
| *Salmonella enterica* | *Escherichia coli* | Flight 3 | 8 | 0.0000 |
| *Salmonella enterica* | *Klebsiella aerogenes strain IIIF7SW-P1* | Flight 3 | 7 | 4.9080 |
| *Salmonella enterica* | *Klebsiella oxytoca* | Flight 1 | 2 | 0.6859 |
| *Salmonella enterica* | *Klebsiella pneumoniae* | Flight 1 | 2 | 0.0000 |
| *Salmonella enterica* | *Klebsiella pneumoniae* | Flight 3 | 1 | 0.0000 |
| *Salmonella enterica* | *Klebsiella pneumoniae* | Flight 3 | 3 | 5.1680 |
| *Salmonella enterica* | *Klebsiella pneumoniae* | Flight 3 | 5 | 4.9080 |
| *Salmonella enterica* | *Klebsiella pneumoniae* | Flight 3 | 7 | 4.9080 |
| *Salmonella enterica* | *Klebsiella pneumoniae* | Flight 3 | 8 | 4.9080 |
| *Salmonella enterica* | *Klebsiella quasipneumoniae strain IF2SW-B3* | Flight 1 | 2 | 0.0000 |
| *Salmonella enterica* | *Klebsiella quasipneumoniae strain IF2SW-P1* | Flight 1 | 2 | 0.0000 |
| *Salmonella enterica* | *Klebsiella quasipneumoniae strain IIIF3SW-P1* | Flight 3 | 3 | 5.1680 |
| *Salmonella enterica* | *Klebsiella sp. strain MS 92-3* | Flight 3 | 3 | 5.1680 |
| *Salmonella enterica* | *Klebsiella variicola* | Flight 3 | 3 | 5.1680 |
| *Salmonella enterica* | *Pantoea agglomerans* | Flight 3 | 1 | 0.0000 |
| *Salmonella enterica* | *Pantoea agglomerans* | Flight 3 | 5 | 0.0000 |
| *Salmonella enterica* | *Pantoea agglomerans* | Flight 3 | 7 | 0.0000 |
| *Salmonella enterica* | *Pantoea agglomerans* | Flight 3 | 8 | 0.0000 |
| *Salmonella enterica* | *Pantoea ananatis* | Flight 1 | 2 | 0.0000 |
| *Salmonella enterica* | *Pantoea ananatis* | Flight 3 | 1 | 0.0000 |
| *Salmonella enterica* | *Pantoea ananatis* | Flight 3 | 5 | 0.0000 |
| *Salmonella enterica* | *Pantoea ananatis* | Flight 3 | 7 | 0.0000 |
| *Salmonella enterica* | *Pantoea ananatis* | Flight 3 | 8 | 0.0000 |
| *Salmonella enterica* | *Pantoea conspicua* | Flight 3 | 1 | 0.6494 |
| *Salmonella enterica* | *Pantoea conspicua* | Flight 3 | 5 | 0.6135 |
| *Salmonella enterica* | *Pantoea conspicua* | Flight 3 | 7 | 0.6135 |
| *Salmonella enterica* | *Pantoea conspicua* | Flight 3 | 8 | 0.6135 |
| *Salmonella enterica* | *Pantoea sp. strain A4* | Flight 3 | 1 | 0.0000 |
| *Salmonella enterica* | *Pantoea sp. strain A4* | Flight 3 | 5 | 0.0000 |
| *Salmonella enterica* | *Pantoea sp. strain A4* | Flight 3 | 7 | 0.0000 |
| *Salmonella enterica* | *Pantoea sp. strain A4* | Flight 3 | 8 | 0.0000 |
| *Salmonella enterica* | *Pantoea sp. strain At-9b* | Flight 3 | 1 | 0.0000 |
| *Salmonella enterica* | *Pantoea sp. strain At-9b* | Flight 3 | 5 | 0.0000 |
| *Salmonella enterica* | *Pantoea sp. strain At-9b* | Flight 3 | 8 | 0.0000 |
| *Salmonella enterica* | *Pantoea sp. strain FF5* | Flight 3 | 1 | 0.0000 |
| *Salmonella enterica* | *Pantoea sp. strain FF5* | Flight 3 | 5 | 0.0000 |
| *Salmonella enterica* | *Pantoea sp. strain FF5* | Flight 3 | 7 | 0.0000 |
| *Salmonella enterica* | *Pantoea sp. strain FF5* | Flight 3 | 8 | 0.0000 |
| *Salmonella enterica* | *Pantoea sp. strain IMH* | Flight 3 | 1 | 0.6494 |
| *Salmonella enterica* | *Pantoea sp. strain IMH* | Flight 3 | 5 | 0.6135 |
| *Salmonella enterica* | *Pantoea sp. strain IMH* | Flight 3 | 7 | 0.6135 |
| *Salmonella enterica* | *Pantoea sp. strain IMH* | Flight 3 | 8 | 0.6135 |
| *Salmonella enterica* | *Pantoea sp. strain NGS-ED-1003* | Flight 3 | 1 | 0.6494 |
| *Salmonella enterica* | *Pantoea sp. strain NGS-ED-1003* | Flight 3 | 5 | 0.6135 |
| *Salmonella enterica* | *Pantoea sp. strain NGS-ED-1003* | Flight 3 | 7 | 0.6135 |
| *Salmonella enterica* | *Pantoea sp. strain NGS-ED-1003* | Flight 3 | 8 | 0.6135 |
| *Salmonella enterica* | *Pantoea sp. strain OXWO6B1* | Flight 3 | 1 | 0.0000 |
| *Salmonella enterica* | *Pantoea sp. strain OXWO6B1* | Flight 3 | 5 | 0.0000 |
| *Salmonella enterica* | *Pantoea sp. strain OXWO6B1* | Flight 3 | 7 | 0.0000 |
| *Salmonella enterica* | *Pantoea sp. strain OXWO6B1* | Flight 3 | 8 | 0.0000 |
| *Salmonella enterica* | *Pantoea vagans* | Flight 3 | 1 | 0.0000 |
| *Salmonella enterica* | *Pantoea vagans* | Flight 3 | 5 | 0.0000 |
| *Salmonella enterica* | *Pantoea vagans* | Flight 3 | 7 | 0.0000 |
| *Salmonella enterica* | *Pantoea vagans* | Flight 3 | 8 | 0.0000 |
| *Salmonella enterica* | *Penicillium rubens* | Flight 1 | 2 | 3.4294 |
| *Salmonella enterica* | *Penicillium rubens* | Flight 3 | 3 | 8.7855 |
| *Salmonella enterica* | *Rhodotorula sp. strain JG-1b* | Flight 1 | 2 | 0.0000 |
| *Salmonella enterica* | *Rhodotorula sp. strain JG-1b* | Flight 3 | 1 | 0.0000 |
| *Salmonella enterica* | *Shigella sonnei* | Flight 1 | 2 | 0.0000 |
| *Salmonella enterica* | *Staphylococcus saprophyticus* | Flight 3 | 3 | 5.1680 |
| *Shigella sonnei* | *Enterobacter asburiae* | Flight 1 | 2 | 0.0000 |
| *Shigella sonnei* | *Enterobacter cancerogenus* | Flight 1 | 2 | 0.0000 |
| *Shigella sonnei* | *Enterobacter cloacae* | Flight 1 | 2 | 0.0000 |
| *Shigella sonnei* | *Enterobacter hormaechei* | Flight 1 | 2 | 0.0000 |
| *Shigella sonnei* | *Enterobacter roggenkampii* | Flight 1 | 2 | 0.0000 |
| *Shigella sonnei* | *Enterobacter sp. strain NFIX59* | Flight 1 | 2 | 0.0000 |
| *Shigella sonnei* | *Escherichia coli* | Flight 1 | 2 | 0.0000 |
| *Shigella sonnei* | *Klebsiella oxytoca* | Flight 1 | 2 | 0.6729 |
| *Shigella sonnei* | *Klebsiella pneumoniae* | Flight 1 | 2 | 0.0000 |
| *Shigella sonnei* | *Klebsiella quasipneumoniae strain IF2SW-B3* | Flight 1 | 2 | 0.0000 |
| *Shigella sonnei* | *Klebsiella quasipneumoniae strain IF2SW-P1* | Flight 1 | 2 | 0.0000 |
| *Shigella sonnei* | *Pantoea ananatis* | Flight 1 | 2 | 0.0000 |
| *Shigella sonnei* | *Penicillium rubens* | Flight 1 | 2 | 0.0000 |
| *Shigella sonnei* | *Rhodotorula sp. strain JG-1b* | Flight 1 | 2 | 0.0000 |
| *Shigella sonnei* | *Salmonella enterica* | Flight 1 | 2 | 0.0000 |
| *Staphylococcus aureus* | *Enterococcus avium* | Flight 3 | 2 | 0.0000 |
| *Staphylococcus aureus* | *Enterococcus faecalis* | Flight 3 | 2 | 0.0000 |
| *Staphylococcus aureus* | *Klebsiella pneumoniae strain F3-2P(2*)* | Flight 3 | 2 | 0.3922 |
| *Staphylococcus aureus* | *Paenibacillus polymyxa* | Flight 3 | 2 | 0.1961 |
| *Staphylococcus aureus* | *Penicillium chrysogenum* | Flight 3 | 2 | 0.5882 |
| *Staphylococcus aureus* | *Penicillium flavigenum* | Flight 3 | 2 | 0.5882 |
| *Staphylococcus aureus* | *Penicillium nalgiovense* | Flight 3 | 2 | 0.5882 |
| *Staphylococcus aureus* | *Penicillium rubens* | Flight 3 | 2 | 0.5882 |

| Microorganism | In the presence of | Flight | Location | Metabolic Support Index (A\|AUB) (%) |
|---|---|---|---|---|
| *Staphylococcus aureus* | *Rhodotorula sp. strain JG-1b* | Flight 3 | 2 | 0.0000 |
| *Staphylococcus aureus* | *Staphylococcus epidermidis* | Flight 3 | 2 | 0.0000 |
| *Staphylococcus aureus* | *Staphylococcus haemolyticus* | Flight 3 | 2 | 0.0000 |
| *Staphylococcus aureus* | *Staphylococcus saprophyticus* | Flight 3 | 2 | 0.0000 |
| *Staphylococcus aureus* | *Staphylococcus sp. strain LCT-H4* | Flight 3 | 2 | 0.1961 |
| *Staphylococcus aureus* | *Staphylococcus warneri* | Flight 3 | 2 | 0.0000 |
| *Staphylococcus epidermidis* | *Enterococcus avium* | Flight 3 | 2 | 0.0000 |
| *Staphylococcus epidermidis* | *Enterococcus faecalis* | Flight 3 | 2 | 0.0000 |
| *Staphylococcus epidermidis* | *Klebsiella pneumoniae strain F3-2P(2*)* | Flight 3 | 2 | 0.4065 |
| *Staphylococcus epidermidis* | *Paenibacillus polymyxa* | Flight 3 | 2 | 0.2033 |
| *Staphylococcus epidermidis* | *Penicillium chrysogenum* | Flight 3 | 2 | 0.4065 |
| *Staphylococcus epidermidis* | *Penicillium flavigenum* | Flight 3 | 2 | 0.4065 |
| *Staphylococcus epidermidis* | *Penicillium nalgiovense* | Flight 3 | 2 | 0.4065 |
| *Staphylococcus epidermidis* | *Penicillium rubens* | Flight 3 | 2 | 0.4065 |
| *Staphylococcus epidermidis* | *Rhodotorula sp. strain JG-1b* | Flight 3 | 2 | 0.2033 |
| *Staphylococcus epidermidis* | *Staphylococcus aureus* | Flight 3 | 2 | 0.2033 |
| *Staphylococcus epidermidis* | *Staphylococcus haemolyticus* | Flight 3 | 2 | 0.0000 |
| *Staphylococcus epidermidis* | *Staphylococcus saprophyticus* | Flight 3 | 2 | 0.2033 |
| *Staphylococcus epidermidis* | *Staphylococcus sp. strain LCT-H4* | Flight 3 | 2 | 0.4065 |
| *Staphylococcus epidermidis* | *Staphylococcus warneri* | Flight 3 | 2 | 0.2033 |
| *Staphylococcus haemolyticus* | *Enterococcus avium* | Flight 3 | 2 | 0.0000 |
| *Staphylococcus haemolyticus* | *Enterococcus faecalis* | Flight 3 | 2 | 0.0000 |
| *Staphylococcus haemolyticus* | *Klebsiella pneumoniae strain F3-2P(2*)* | Flight 3 | 2 | 0.3817 |
| *Staphylococcus haemolyticus* | *Paenibacillus polymyxa* | Flight 3 | 2 | 0.1908 |
| *Staphylococcus haemolyticus* | *Penicillium chrysogenum* | Flight 3 | 2 | 0.1908 |
| *Staphylococcus haemolyticus* | *Penicillium flavigenum* | Flight 3 | 2 | 0.1908 |
| *Staphylococcus haemolyticus* | *Penicillium nalgiovense* | Flight 3 | 2 | 0.1908 |
| *Staphylococcus haemolyticus* | *Penicillium rubens* | Flight 3 | 2 | 0.1908 |
| *Staphylococcus haemolyticus* | *Rhodotorula sp. strain JG-1b* | Flight 3 | 2 | 0.0000 |
| *Staphylococcus haemolyticus* | *Staphylococcus aureus* | Flight 3 | 2 | 0.0000 |
| *Staphylococcus haemolyticus* | *Staphylococcus epidermidis* | Flight 3 | 2 | 0.0000 |
| *Staphylococcus haemolyticus* | *Staphylococcus saprophyticus* | Flight 3 | 2 | 0.0000 |
| *Staphylococcus haemolyticus* | *Staphylococcus sp. strain LCT-H4* | Flight 3 | 2 | 0.1908 |
| *Staphylococcus haemolyticus* | *Staphylococcus warneri* | Flight 3 | 2 | 0.0000 |
| *Staphylococcus saprophyticus* | *Enterococcus avium* | Flight 3 | 2 | 0.0000 |
| *Staphylococcus saprophyticus* | *Enterococcus faecalis* | Flight 3 | 2 | 0.0000 |
| *Staphylococcus saprophyticus* | *Escherichia coli* | Flight 3 | 3 | 2.5997 |
| *Staphylococcus saprophyticus* | *Klebsiella pneumoniae* | Flight 3 | 3 | 2.5997 |
| *Staphylococcus saprophyticus* | *Klebsiella pneumoniae strain F3-2P(2*)* | Flight 3 | 2 | 0.1876 |
| *Staphylococcus saprophyticus* | *Klebsiella quasipneumoniae strain IIIF3SW-P1* | Flight 3 | 3 | 2.5997 |
| *Staphylococcus saprophyticus* | *Klebsiella sp. strain MS 92-3* | Flight 3 | 3 | 2.5997 |
| *Staphylococcus saprophyticus* | *Klebsiella variicola* | Flight 3 | 3 | 2.5997 |
| *Staphylococcus saprophyticus* | *Paenibacillus polymyxa* | Flight 3 | 2 | 0.0000 |
| *Staphylococcus saprophyticus* | *Penicillium chrysogenum* | Flight 3 | 2 | 0.1876 |
| *Staphylococcus saprophyticus* | *Penicillium flavigenum* | Flight 3 | 2 | 0.1876 |
| *Staphylococcus saprophyticus* | *Penicillium nalgiovense* | Flight 3 | 2 | 0.1876 |
| *Staphylococcus saprophyticus* | *Penicillium rubens* | Flight 3 | 2 | 0.1876 |
| *Staphylococcus saprophyticus* | *Penicillium rubens* | Flight 3 | 3 | 0.8666 |
| *Staphylococcus saprophyticus* | *Rhodotorula sp. strain JG-1b* | Flight 3 | 2 | 0.0000 |
| *Staphylococcus saprophyticus* | *Salmonella enterica* | Flight 3 | 3 | 2.5997 |
| *Staphylococcus saprophyticus* | *Staphylococcus aureus* | Flight 3 | 2 | 0.0000 |
| *Staphylococcus saprophyticus* | *Staphylococcus epidermidis* | Flight 3 | 2 | 0.0000 |
| *Staphylococcus saprophyticus* | *Staphylococcus haemolyticus* | Flight 3 | 2 | 0.0000 |
| *Staphylococcus saprophyticus* | *Staphylococcus sp. strain LCT-H4* | Flight 3 | 2 | 0.0000 |
| *Staphylococcus saprophyticus* | *Staphylococcus warneri* | Flight 3 | 2 | 0.0000 |
| *Staphylococcus sp. strain LCT-H4* | *Enterococcus avium* | Flight 3 | 2 | 0.7018 |
| *Staphylococcus sp. strain LCT-H4* | *Enterococcus faecalis* | Flight 3 | 2 | 0.0000 |
| *Staphylococcus sp. strain LCT-H4* | *Klebsiella pneumoniae strain F3-2P(2*)* | Flight 3 | 2 | 0.1754 |
| *Staphylococcus sp. strain LCT-H4* | *Paenibacillus polymyxa* | Flight 3 | 2 | 0.0000 |
| *Staphylococcus sp. strain LCT-H4* | *Penicillium chrysogenum* | Flight 3 | 2 | 0.1754 |
| *Staphylococcus sp. strain LCT-H4* | *Penicillium flavigenum* | Flight 3 | 2 | 0.1754 |
| *Staphylococcus sp. strain LCT-H4* | *Penicillium nalgiovense* | Flight 3 | 2 | 0.1754 |
| *Staphylococcus sp. strain LCT-H4* | *Penicillium rubens* | Flight 3 | 2 | 0.1754 |
| *Staphylococcus sp. strain LCT-H4* | *Rhodotorula sp. strain JG-1b* | Flight 3 | 2 | 0.0000 |
| *Staphylococcus sp. strain LCT-H4* | *Staphylococcus aureus* | Flight 3 | 2 | 0.7018 |
| *Staphylococcus sp. strain LCT-H4* | *Staphylococcus epidermidis* | Flight 3 | 2 | 0.0000 |
| *Staphylococcus sp. strain LCT-H4* | *Staphylococcus haemolyticus* | Flight 3 | 2 | 0.0000 |
| *Staphylococcus sp. strain LCT-H4* | *Staphylococcus saprophyticus* | Flight 3 | 2 | 0.0000 |
| *Staphylococcus sp. strain LCT-H4* | *Staphylococcus warneri* | Flight 3 | 2 | 0.0000 |
| *Staphylococcus warneri* | *Enterococcus avium* | Flight 3 | 2 | 1.7964 |
| *Staphylococcus warneri* | *Enterococcus faecalis* | Flight 3 | 2 | 1.7964 |
| *Staphylococcus warneri* | *Klebsiella pneumoniae strain F3-2P(2*)* | Flight 3 | 2 | 2.1956 |
| *Staphylococcus warneri* | *Paenibacillus polymyxa* | Flight 3 | 2 | 1.9960 |
| *Staphylococcus warneri* | *Penicillium chrysogenum* | Flight 3 | 2 | 0.1996 |
| *Staphylococcus warneri* | *Penicillium flavigenum* | Flight 3 | 2 | 0.1996 |
| *Staphylococcus warneri* | *Penicillium nalgiovense* | Flight 3 | 2 | 0.1996 |
| *Staphylococcus warneri* | *Penicillium rubens* | Flight 3 | 2 | 0.1996 |
| *Staphylococcus warneri* | *Rhodotorula sp. strain JG-1b* | Flight 3 | 2 | 0.0000 |
| *Staphylococcus warneri* | *Staphylococcus aureus* | Flight 3 | 2 | 1.7964 |
| *Staphylococcus warneri* | *Staphylococcus epidermidis* | Flight 3 | 2 | 1.7964 |
| *Staphylococcus warneri* | *Staphylococcus haemolyticus* | Flight 3 | 2 | 1.7964 |
| *Staphylococcus warneri* | *Staphylococcus saprophyticus* | Flight 3 | 2 | 1.7964 |
| *Staphylococcus warneri* | *Staphylococcus sp. strain LCT-H4* | Flight 3 | 2 | 1.9960 |

**Table S9: Nature of interactions as predicted by the constraint-based analyses**

| Microorganism A | Microorganism B | Flight | Location | Vbio of A | Vbio of B | Vbio of A in AB | Vbio of B in AB | Ratio of A in AB | Ratio of B in AB | Effect of AB on A | Effect of AB on B | Significant effect on A | Significant effect on B | Interaction type |
|---|---|---|---|---|---|---|---|---|---|---|---|---|---|---|
| *Aspergillus niger* | *Enterobacter cloacae* | Flight 3 | 1 | 0.544 | 8.182 | 0.181 | 9.101 | 0.019 | 0.981 | -0.668 | 0.112 | -0.668 | 0.112 | Parasitism |
| *Aspergillus niger* | *Escherichia coli* | Flight 3 | 1 | 0.544 | 8.314 | 0.152 | 9.268 | 0.016 | 0.984 | -0.720 | 0.115 | -0.720 | 0.115 | Parasitism |
| *Aspergillus niger* | *Klebsiella pneumoniae* | Flight 1 | 1 | 0.544 | 9.634 | 0.363 | 10.845 | 0.032 | 0.968 | -0.333 | 0.126 | -0.333 | 0.126 | Parasitism |
| *Aspergillus niger* | *Klebsiella pneumoniae* | Flight 3 | 1 | 0.544 | 9.051 | 0.359 | 10.310 | 0.034 | 0.966 | -0.340 | 0.139 | -0.340 | 0.139 | Parasitism |
| *Aspergillus niger* | *Klebsiella quasipneumoniae strain IF1SW-B2* | Flight 1 | 1 | 0.544 | 9.634 | 0.363 | 10.845 | 0.032 | 0.968 | -0.333 | 0.126 | -0.333 | 0.126 | Parasitism |
| *Aspergillus niger* | *Klebsiella quasipneumoniae strain IF1SW-P3* | Flight 1 | 1 | 0.544 | 9.634 | 0.363 | 10.845 | 0.032 | 0.968 | -0.333 | 0.126 | -0.333 | 0.126 | Parasitism |
| *Aspergillus niger* | *Klebsiella quasipneumoniae strain IF1SW-P4* | Flight 1 | 1 | 0.544 | 9.634 | 0.363 | 10.845 | 0.032 | 0.968 | -0.333 | 0.126 | -0.333 | 0.126 | Parasitism |
| *Aspergillus niger* | *Pantoea agglomerans* | Flight 3 | 1 | 0.544 | 4.640 | 0.215 | 5.927 | 0.035 | 0.965 | -0.606 | 0.278 | -0.606 | 0.278 | Parasitism |
| *Aspergillus niger* | *Pantoea ananatis* | Flight 3 | 1 | 0.544 | 7.622 | 0.296 | 8.357 | 0.034 | 0.966 | -0.456 | 0.096 | -0.456 | 0.000 | Amensalism |
| *Aspergillus niger* | *Pantoea conspicua* | Flight 3 | 1 | 0.544 | 4.408 | 0.081 | 5.405 | 0.015 | 0.985 | -0.850 | 0.226 | -0.850 | 0.226 | Parasitism |
| *Aspergillus niger* | *Pantoea sp. strain A4* | Flight 3 | 1 | 0.544 | 4.256 | 0.192 | 5.252 | 0.035 | 0.965 | -0.647 | 0.234 | -0.647 | 0.234 | Parasitism |
| *Aspergillus niger* | *Pantoea sp. strain At-9b* | Flight 3 | 1 | 0.544 | 3.804 | 0.068 | 4.999 | 0.014 | 0.987 | -0.874 | 0.314 | -0.874 | 0.314 | Parasitism |
| *Aspergillus niger* | *Pantoea sp. strain FF5* | Flight 3 | 1 | 0.544 | 4.640 | 0.208 | 5.713 | 0.035 | 0.965 | -0.618 | 0.231 | -0.618 | 0.231 | Parasitism |
| *Aspergillus niger* | *Pantoea sp. strain IMH* | Flight 3 | 1 | 0.544 | 4.409 | 0.092 | 5.830 | 0.016 | 0.984 | -0.830 | 0.322 | -0.830 | 0.322 | Parasitism |
| *Aspergillus niger* | *Pantoea sp. strain NGS-ED-1003* | Flight 3 | 1 | 0.544 | 4.640 | 0.091 | 5.855 | 0.015 | 0.985 | -0.832 | 0.262 | -0.832 | 0.262 | Parasitism |
| *Aspergillus niger* | *Pantoea sp. strain OXWO6B1* | Flight 3 | 1 | 0.544 | 4.642 | 0.208 | 5.716 | 0.035 | 0.965 | -0.618 | 0.231 | -0.618 | 0.231 | Parasitism |
| *Aspergillus niger* | *Pantoea vagans* | Flight 3 | 1 | 0.544 | 4.642 | 0.215 | 5.930 | 0.035 | 0.965 | -0.605 | 0.278 | -0.605 | 0.278 | Parasitism |
| *Aspergillus niger* | *Penicillium chrysogenum* | Flight 1 | 1 | 0.544 | 0.071 | 0.718 | 0.017 | 0.977 | 0.023 | 0.320 | -0.759 | 0.320 | -0.759 | Parasitism |
| *Aspergillus niger* | *Penicillium rubens* | Flight 1 | 1 | 0.544 | 0.071 | 0.718 | 0.017 | 0.977 | 0.023 | 0.320 | -0.759 | 0.320 | -0.759 | Parasitism |
| *Aspergillus niger* | *Rhodotorula sp. strain JG-1b* | Flight 1 | 1 | 0.544 | 0.154 | 0.542 | 0.156 | 0.777 | 0.223 | -0.003 | 0.009 | 0.000 | 0.000 | Neutral |
| *Aspergillus niger* | *Rhodotorula sp. strain JG-1b* | Flight 3 | 1 | 0.544 | 0.154 | 0.542 | 0.156 | 0.777 | 0.223 | -0.003 | 0.009 | 0.000 | 0.000 | Neutral |
| *Aspergillus niger* | *Rhodotorula toruloides* | Flight 1 | 1 | 0.544 | 2.346 | 0.538 | 2.345 | 0.187 | 0.813 | -0.010 | 0.000 | 0.000 | 0.000 | Neutral |
| *Aspergillus niger* | *Salmonella enterica* | Flight 3 | 1 | 0.544 | 8.372 | 0.428 | 9.546 | 0.043 | 0.957 | -0.213 | 0.140 | -0.213 | 0.140 | Parasitism |
| *Enterobacter asburiae* | *Enterobacter cancerogenus* | Flight 1 | 2 | 8.200 | 7.602 | 8.200 | 0.000 | 1.000 | 0.000 | 0.000 | -1.000 | 0.000 | -1.000 | Amensalism |
| *Enterobacter asburiae* | *Enterobacter cloacae* | Flight 1 | 2 | 8.200 | 8.200 | 0.000 | 8.200 | 0.000 | 1.000 | -1.000 | 0.000 | -1.000 | 0.000 | Amensalism |
| *Enterobacter asburiae* | *Enterobacter hormaechei* | Flight 1 | 2 | 8.200 | 8.200 | 4.059 | 4.141 | 0.495 | 0.505 | -0.505 | -0.495 | -0.505 | -0.495 | Competitive |
| *Enterobacter asburiae* | *Enterobacter roggenkampii* | Flight 1 | 2 | 8.200 | 8.200 | 6.338 | 1.862 | 0.773 | 0.227 | -0.227 | -0.773 | -0.227 | -0.773 | Competitive |
| *Enterobacter asburiae* | *Enterobacter sp. strain NFIX59* | Flight 1 | 2 | 8.200 | 8.398 | 8.318 | 0.084 | 0.990 | 0.010 | 0.014 | -0.990 | 0.000 | -0.990 | Amensalism |
| *Enterobacter asburiae* | *Escherichia coli* | Flight 1 | 2 | 8.200 | 8.331 | 8.218 | 0.157 | 0.981 | 0.019 | 0.002 | -0.981 | 0.000 | -0.981 | Amensalism |
| *Enterobacter asburiae* | *Klebsiella oxytoca* | Flight 1 | 2 | 8.200 | 9.403 | 9.013 | 0.415 | 0.956 | 0.044 | 0.099 | -0.956 | 0.000 | -0.956 | Amensalism |
| *Enterobacter asburiae* | *Klebsiella pneumoniae* | Flight 1 | 2 | 8.200 | 9.251 | 8.835 | 0.439 | 0.953 | 0.047 | 0.077 | -0.953 | 0.000 | -0.953 | Amensalism |
| *Enterobacter asburiae* | *Klebsiella quasipneumoniae strain IF2SW-B3* | Flight 1 | 2 | 8.200 | 9.251 | 8.835 | 0.439 | 0.953 | 0.047 | 0.077 | -0.953 | 0.000 | -0.953 | Amensalism |
| *Enterobacter asburiae* | *Klebsiella quasipneumoniae strain IF2SW-P1* | Flight 1 | 2 | 8.200 | 9.251 | 8.835 | 0.439 | 0.953 | 0.047 | 0.077 | -0.953 | 0.000 | -0.953 | Amensalism |
| *Enterobacter asburiae* | *Pantoea ananatis* | Flight 1 | 2 | 8.200 | 7.840 | 9.050 | 10.087 | 0.473 | 0.527 | 0.104 | 0.287 | 0.104 | 0.287 | Mutualism |
| *Enterobacter asburiae* | *Penicillium rubens* | Flight 1 | 2 | 8.200 | 0.049 | 8.200 | 0.000 | 1.000 | 0.000 | 0.000 | -1.000 | 0.000 | -1.000 | Amensalism |
| *Enterobacter asburiae* | *Rhodotorula sp. strain JG-1b* | Flight 1 | 2 | 8.200 | 0.153 | 8.183 | 0.150 | 0.982 | 0.018 | -0.002 | -0.021 | 0.000 | 0.000 | Neutral |
| *Enterobacter asburiae* | *Salmonella enterica* | Flight 1 | 2 | 8.200 | 8.478 | 8.836 | 0.439 | 0.953 | 0.047 | 0.078 | -0.948 | 0.000 | -0.948 | Amensalism |
| *Enterobacter asburiae* | *Shigella sonnei* | Flight 1 | 2 | 8.200 | 8.331 | 8.218 | 0.157 | 0.981 | 0.019 | 0.002 | -0.981 | 0.000 | -0.981 | Amensalism |
| *Enterobacter cancerogenus* | *Enterobacter cloacae* | Flight 1 | 2 | 7.602 | 8.200 | 0.000 | 8.200 | 0.000 | 1.000 | -1.000 | 0.000 | -1.000 | 0.000 | Amensalism |
| *Enterobacter cancerogenus* | *Enterobacter hormaechei* | Flight 1 | 2 | 7.602 | 8.200 | 0.000 | 8.200 | 0.000 | 1.000 | -1.000 | 0.000 | -1.000 | 0.000 | Amensalism |
| *Enterobacter cancerogenus* | *Enterobacter roggenkampii* | Flight 1 | 2 | 7.602 | 8.200 | 0.000 | 8.200 | 0.000 | 1.000 | -1.000 | 0.000 | -1.000 | 0.000 | Amensalism |
| *Enterobacter cancerogenus* | *Enterobacter sp. strain NFIX59* | Flight 1 | 2 | 7.602 | 8.398 | 7.141 | 1.258 | 0.850 | 0.150 | -0.061 | -0.850 | 0.000 | -0.850 | Amensalism |
| *Enterobacter cancerogenus* | *Escherichia coli* | Flight 1 | 2 | 7.602 | 8.331 | 0.000 | 8.331 | 0.000 | 1.000 | -1.000 | 0.000 | -1.000 | 0.000 | Amensalism |
| *Enterobacter cancerogenus* | *Klebsiella oxytoca* | Flight 1 | 2 | 7.602 | 9.403 | 0.000 | 9.403 | 0.000 | 1.000 | -1.000 | 0.000 | -1.000 | 0.000 | Amensalism |
| *Enterobacter cancerogenus* | *Klebsiella pneumoniae* | Flight 1 | 2 | 7.602 | 9.251 | 0.000 | 9.251 | 0.000 | 1.000 | -1.000 | 0.000 | -1.000 | 0.000 | Amensalism |
| *Enterobacter cancerogenus* | *Klebsiella quasipneumoniae strain IF2SW-B3* | Flight 1 | 2 | 7.602 | 9.251 | 0.000 | 9.251 | 0.000 | 1.000 | -1.000 | 0.000 | -1.000 | 0.000 | Amensalism |
| *Enterobacter cancerogenus* | *Klebsiella quasipneumoniae strain IF2SW-P1* | Flight 1 | 2 | 7.602 | 9.251 | 0.000 | 9.251 | 0.000 | 1.000 | -1.000 | 0.000 | -1.000 | 0.000 | Amensalism |
| *Enterobacter cancerogenus* | *Pantoea ananatis* | Flight 1 | 2 | 7.602 | 7.840 | 9.033 | 10.068 | 0.473 | 0.527 | 0.188 | 0.284 | 0.188 | 0.284 | Mutualism |
| *Enterobacter cancerogenus* | *Penicillium rubens* | Flight 1 | 2 | 7.602 | 0.049 | 7.602 | 0.000 | 1.000 | 0.000 | 0.000 | -1.000 | 0.000 | -1.000 | Amensalism |
| *Enterobacter cancerogenus* | *Rhodotorula sp. strain JG-1b* | Flight 1 | 2 | 7.602 | 0.153 | 7.632 | 0.140 | 0.982 | 0.018 | 0.004 | -0.084 | 0.000 | 0.000 | Neutral |
| *Enterobacter cancerogenus* | *Salmonella enterica* | Flight 1 | 2 | 7.602 | 8.478 | 0.000 | 8.478 | 0.000 | 1.000 | -1.000 | 0.000 | -1.000 | 0.000 | Amensalism |
| *Enterobacter cancerogenus* | *Shigella sonnei* | Flight 1 | 2 | 7.602 | 8.331 | 0.000 | 8.331 | 0.000 | 1.000 | -1.000 | 0.000 | -1.000 | 0.000 | Amensalism |
| *Enterobacter cloacae* | *Enterobacter hormaechei* | Flight 1 | 2 | 8.200 | 8.200 | 7.972 | 0.227 | 0.972 | 0.028 | -0.028 | -0.972 | 0.000 | -0.972 | Amensalism |
| *Enterobacter cloacae* | *Enterobacter roggenkampii* | Flight 1 | 2 | 8.200 | 8.200 | 8.200 | 0.000 | 1.000 | 0.000 | 0.000 | -1.000 | 0.000 | -1.000 | Amensalism |
| *Enterobacter cloacae* | *Enterobacter sp. strain NFIX59* | Flight 1 | 2 | 8.200 | 8.398 | 8.318 | 0.084 | 0.990 | 0.010 | 0.014 | -0.990 | 0.000 | -0.990 | Amensalism |
| *Enterobacter cloacae* | *Escherichia coli* | Flight 1 | 2 | 8.200 | 8.331 | 8.218 | 0.157 | 0.981 | 0.019 | 0.002 | -0.981 | 0.000 | -0.981 | Amensalism |
| *Enterobacter cloacae* | *Escherichia coli* | Flight 3 | 1 | 8.182 | 8.314 | 8.201 | 0.156 | 0.981 | 0.019 | 0.002 | -0.981 | 0.000 | -0.981 | Amensalism |
| *Enterobacter cloacae* | *Escherichia coli* | Flight 3 | 5 | 6.106 | 6.230 | 6.137 | 0.114 | 0.982 | 0.018 | 0.005 | -0.982 | 0.000 | -0.982 | Amensalism |
| *Enterobacter cloacae* | *Escherichia coli* | Flight 3 | 7 | 6.106 | 6.230 | 6.137 | 0.114 | 0.982 | 0.018 | 0.005 | -0.982 | 0.000 | -0.982 | Amensalism |
| *Enterobacter cloacae* | *Escherichia coli* | Flight 3 | 8 | 6.106 | 6.230 | 6.137 | 0.114 | 0.982 | 0.018 | 0.005 | -0.982 | 0.000 | -0.982 | Amensalism |
| *Enterobacter cloacae* | *Klebsiella aerogenes strain IIIF7SW-P1* | Flight 3 | 7 | 6.106 | 7.338 | 10.397 | 5.499 | 0.654 | 0.346 | 0.703 | -0.251 | 0.703 | -0.251 | Parasitism |
| *Enterobacter cloacae* | *Klebsiella oxytoca* | Flight 1 | 2 | 8.200 | 9.403 | 9.013 | 0.415 | 0.956 | 0.044 | 0.099 | -0.956 | 0.000 | -0.956 | Amensalism |
| *Enterobacter cloacae* | *Klebsiella pneumoniae* | Flight 1 | 2 | 8.200 | 9.251 | 8.835 | 0.439 | 0.953 | 0.047 | 0.077 | -0.953 | 0.000 | -0.953 | Amensalism |
| *Enterobacter cloacae* | *Klebsiella pneumoniae* | Flight 3 | 1 | 8.182 | 9.051 | 8.653 | 0.402 | 0.956 | 0.044 | 0.057 | -0.956 | 0.000 | -0.956 | Amensalism |
| *Enterobacter cloacae* | *Klebsiella pneumoniae* | Flight 3 | 5 | 6.106 | 6.623 | 6.517 | 0.136 | 0.980 | 0.020 | 0.067 | -0.979 | 0.000 | -0.979 | Amensalism |

| Microorganism A | Microorganism B | Flight | Location | Vbio of A | Vbio of B | Vbio of A in AB | Vbio of B in AB | Ratio of A in AB | Ratio of B in AB | Effect of AB on A | Effect of AB on B | Significant effect on A | Significant effect on B | Interaction type |
|---|---|---|---|---|---|---|---|---|---|---|---|---|---|---|
| *Enterobacter cloacae* | *Klebsiella pneumoniae* | Flight 3 | 7 | 6.106 | 6.623 | 6.517 | 0.136 | 0.980 | 0.020 | 0.067 | -0.979 | 0.000 | -0.979 | Amensalism |
| *Enterobacter cloacae* | *Klebsiella pneumoniae* | Flight 3 | 8 | 6.106 | 6.623 | 6.517 | 0.136 | 0.980 | 0.020 | 0.067 | -0.979 | 0.000 | -0.979 | Amensalism |
| *Enterobacter cloacae* | *Klebsiella quasipneumoniae strain IF2SW-B3* | Flight 1 | 2 | 8.200 | 9.251 | 8.835 | 0.439 | 0.953 | 0.047 | 0.077 | -0.953 | 0.000 | -0.953 | Amensalism |
| *Enterobacter cloacae* | *Klebsiella quasipneumoniae strain IF2SW-P1* | Flight 1 | 2 | 8.200 | 9.251 | 8.835 | 0.439 | 0.953 | 0.047 | 0.077 | -0.953 | 0.000 | -0.953 | Amensalism |
| *Enterobacter cloacae* | *Pantoea agglomerans* | Flight 3 | 1 | 8.182 | 4.640 | 8.632 | 0.596 | 0.935 | 0.065 | 0.055 | -0.872 | 0.000 | -0.872 | Amensalism |
| *Enterobacter cloacae* | *Pantoea agglomerans* | Flight 3 | 5 | 6.106 | 2.777 | 6.772 | 0.367 | 0.949 | 0.051 | 0.109 | -0.868 | 0.109 | -0.868 | Parasitism |
| *Enterobacter cloacae* | *Pantoea agglomerans* | Flight 3 | 7 | 6.106 | 2.777 | 6.772 | 0.367 | 0.949 | 0.051 | 0.109 | -0.868 | 0.109 | -0.868 | Parasitism |
| *Enterobacter cloacae* | *Pantoea agglomerans* | Flight 3 | 8 | 6.106 | 2.777 | 6.772 | 0.367 | 0.949 | 0.051 | 0.109 | -0.868 | 0.109 | -0.868 | Parasitism |
| *Enterobacter cloacae* | *Pantoea ananatis* | Flight 1 | 2 | 8.200 | 7.840 | 9.050 | 10.087 | 0.473 | 0.527 | 0.104 | 0.287 | 0.104 | 0.287 | Mutualism |
| *Enterobacter cloacae* | *Pantoea ananatis* | Flight 3 | 1 | 8.182 | 7.622 | 8.941 | 9.967 | 0.473 | 0.527 | 0.093 | 0.308 | 0.000 | 0.308 | Commensals |
| *Enterobacter cloacae* | *Pantoea ananatis* | Flight 3 | 5 | 6.106 | 5.534 | 6.946 | 7.805 | 0.471 | 0.529 | 0.138 | 0.410 | 0.138 | 0.410 | Mutualism |
| *Enterobacter cloacae* | *Pantoea ananatis* | Flight 3 | 7 | 6.106 | 5.534 | 6.946 | 7.805 | 0.471 | 0.529 | 0.138 | 0.410 | 0.138 | 0.410 | Mutualism |
| *Enterobacter cloacae* | *Pantoea ananatis* | Flight 3 | 8 | 6.106 | 5.534 | 6.946 | 7.805 | 0.471 | 0.529 | 0.138 | 0.410 | 0.138 | 0.410 | Mutualism |
| *Enterobacter cloacae* | *Pantoea conspicua* | Flight 3 | 1 | 8.182 | 4.408 | 8.549 | 0.695 | 0.925 | 0.075 | 0.045 | -0.842 | 0.000 | -0.842 | Amensalism |
| *Enterobacter cloacae* | *Pantoea conspicua* | Flight 3 | 5 | 6.106 | 2.768 | 6.721 | 0.450 | 0.937 | 0.063 | 0.101 | -0.837 | 0.101 | -0.837 | Parasitism |
| *Enterobacter cloacae* | *Pantoea conspicua* | Flight 3 | 7 | 6.106 | 2.768 | 6.721 | 0.450 | 0.937 | 0.063 | 0.101 | -0.837 | 0.101 | -0.837 | Parasitism |
| *Enterobacter cloacae* | *Pantoea conspicua* | Flight 3 | 8 | 6.106 | 2.768 | 6.721 | 0.450 | 0.937 | 0.063 | 0.101 | -0.837 | 0.101 | -0.837 | Parasitism |
| *Enterobacter cloacae* | *Pantoea sp. strain A4* | Flight 3 | 1 | 8.182 | 4.256 | 8.434 | 0.674 | 0.926 | 0.074 | 0.031 | -0.842 | 0.000 | -0.842 | Amensalism |
| *Enterobacter cloacae* | *Pantoea sp. strain A4* | Flight 3 | 5 | 6.106 | 2.593 | 6.600 | 0.351 | 0.949 | 0.051 | 0.081 | -0.864 | 0.000 | -0.864 | Amensalism |
| *Enterobacter cloacae* | *Pantoea sp. strain A4* | Flight 3 | 7 | 6.106 | 2.593 | 6.600 | 0.351 | 0.949 | 0.051 | 0.081 | -0.864 | 0.000 | -0.864 | Amensalism |
| *Enterobacter cloacae* | *Pantoea sp. strain A4* | Flight 3 | 8 | 6.106 | 2.593 | 6.600 | 0.351 | 0.949 | 0.051 | 0.081 | -0.864 | 0.000 | -0.864 | Amensalism |
| *Enterobacter cloacae* | *Pantoea sp. strain At-9b* | Flight 3 | 1 | 8.182 | 3.804 | 8.863 | 0.263 | 0.971 | 0.029 | 0.083 | -0.931 | 0.000 | -0.931 | Amensalism |
| *Enterobacter cloacae* | *Pantoea sp. strain At-9b* | Flight 3 | 5 | 6.106 | 2.173 | 6.736 | 0.253 | 0.964 | 0.036 | 0.103 | -0.884 | 0.103 | -0.884 | Parasitism |
| *Enterobacter cloacae* | *Pantoea sp. strain At-9b* | Flight 3 | 8 | 6.106 | 2.173 | 6.736 | 0.253 | 0.964 | 0.036 | 0.103 | -0.884 | 0.103 | -0.884 | Parasitism |
| *Enterobacter cloacae* | *Pantoea sp. strain FF5* | Flight 3 | 1 | 8.182 | 4.640 | 8.632 | 0.596 | 0.935 | 0.065 | 0.055 | -0.872 | 0.000 | -0.872 | Amensalism |
| *Enterobacter cloacae* | *Pantoea sp. strain FF5* | Flight 3 | 5 | 6.106 | 2.777 | 6.772 | 0.367 | 0.949 | 0.051 | 0.109 | -0.868 | 0.109 | -0.868 | Parasitism |
| *Enterobacter cloacae* | *Pantoea sp. strain FF5* | Flight 3 | 7 | 6.106 | 2.777 | 6.772 | 0.367 | 0.949 | 0.051 | 0.109 | -0.868 | 0.109 | -0.868 | Parasitism |
| *Enterobacter cloacae* | *Pantoea sp. strain FF5* | Flight 3 | 8 | 6.106 | 2.777 | 6.772 | 0.367 | 0.949 | 0.051 | 0.109 | -0.868 | 0.109 | -0.868 | Parasitism |
| *Enterobacter cloacae* | *Pantoea sp. strain IMH* | Flight 3 | 1 | 8.182 | 4.409 | 7.593 | 1.295 | 0.854 | 0.146 | -0.072 | -0.706 | 0.000 | -0.706 | Amensalism |
| *Enterobacter cloacae* | *Pantoea sp. strain IMH* | Flight 3 | 5 | 6.106 | 2.634 | 6.024 | 0.733 | 0.892 | 0.108 | -0.014 | -0.722 | 0.000 | -0.722 | Amensalism |
| *Enterobacter cloacae* | *Pantoea sp. strain IMH* | Flight 3 | 7 | 6.106 | 2.634 | 6.024 | 0.733 | 0.892 | 0.108 | -0.014 | -0.722 | 0.000 | -0.722 | Amensalism |
| *Enterobacter cloacae* | *Pantoea sp. strain IMH* | Flight 3 | 8 | 6.106 | 2.634 | 6.024 | 0.733 | 0.892 | 0.108 | -0.014 | -0.722 | 0.000 | -0.722 | Amensalism |
| *Enterobacter cloacae* | *Pantoea sp. strain NGS-ED-1003* | Flight 3 | 1 | 8.182 | 4.640 | 8.632 | 0.596 | 0.935 | 0.065 | 0.055 | -0.872 | 0.000 | -0.872 | Amensalism |
| *Enterobacter cloacae* | *Pantoea sp. strain NGS-ED-1003* | Flight 3 | 5 | 6.106 | 2.777 | 6.772 | 0.367 | 0.949 | 0.051 | 0.109 | -0.868 | 0.109 | -0.868 | Parasitism |
| *Enterobacter cloacae* | *Pantoea sp. strain NGS-ED-1003* | Flight 3 | 7 | 6.106 | 2.777 | 6.772 | 0.367 | 0.949 | 0.051 | 0.109 | -0.868 | 0.109 | -0.868 | Parasitism |
| *Enterobacter cloacae* | *Pantoea sp. strain NGS-ED-1003* | Flight 3 | 8 | 6.106 | 2.777 | 6.772 | 0.367 | 0.949 | 0.051 | 0.109 | -0.868 | 0.109 | -0.868 | Parasitism |
| *Enterobacter cloacae* | *Pantoea sp. strain OXWO6B1* | Flight 3 | 1 | 8.182 | 4.642 | 8.633 | 0.596 | 0.935 | 0.065 | 0.055 | -0.872 | 0.000 | -0.872 | Amensalism |
| *Enterobacter cloacae* | *Pantoea sp. strain OXWO6B1* | Flight 3 | 5 | 6.106 | 2.779 | 6.772 | 0.367 | 0.949 | 0.051 | 0.109 | -0.868 | 0.109 | -0.868 | Parasitism |
| *Enterobacter cloacae* | *Pantoea sp. strain OXWO6B1* | Flight 3 | 7 | 6.106 | 2.779 | 6.772 | 0.367 | 0.949 | 0.051 | 0.109 | -0.868 | 0.109 | -0.868 | Parasitism |
| *Enterobacter cloacae* | *Pantoea sp. strain OXWO6B1* | Flight 3 | 8 | 6.106 | 2.779 | 6.772 | 0.367 | 0.949 | 0.051 | 0.109 | -0.868 | 0.109 | -0.868 | Parasitism |
| *Enterobacter cloacae* | *Pantoea vagans* | Flight 3 | 1 | 8.182 | 4.642 | 8.633 | 0.596 | 0.935 | 0.065 | 0.055 | -0.872 | 0.000 | -0.872 | Amensalism |
| *Enterobacter cloacae* | *Pantoea vagans* | Flight 3 | 5 | 6.106 | 2.778 | 6.772 | 0.367 | 0.949 | 0.051 | 0.109 | -0.868 | 0.109 | -0.868 | Parasitism |
| *Enterobacter cloacae* | *Pantoea vagans* | Flight 3 | 7 | 6.106 | 2.778 | 6.772 | 0.367 | 0.949 | 0.051 | 0.109 | -0.868 | 0.109 | -0.868 | Parasitism |
| *Enterobacter cloacae* | *Pantoea vagans* | Flight 3 | 8 | 6.106 | 2.778 | 6.772 | 0.367 | 0.949 | 0.051 | 0.109 | -0.868 | 0.109 | -0.868 | Parasitism |
| *Enterobacter cloacae* | *Penicillium rubens* | Flight 1 | 2 | 8.200 | 0.049 | 8.200 | 0.000 | 1.000 | 0.000 | 0.000 | -1.000 | 0.000 | -1.000 | Amensalism |
| *Enterobacter cloacae* | *Rhodotorula sp. strain JG-1b* | Flight 1 | 2 | 8.200 | 0.153 | 8.183 | 0.150 | 0.982 | 0.018 | -0.002 | -0.021 | 0.000 | 0.000 | Neutral |
| *Enterobacter cloacae* | *Rhodotorula sp. strain JG-1b* | Flight 3 | 1 | 8.182 | 0.154 | 8.182 | 0.000 | 1.000 | 0.000 | 0.000 | -1.000 | 0.000 | -1.000 | Amensalism |
| *Enterobacter cloacae* | *Salmonella enterica* | Flight 1 | 2 | 8.200 | 8.478 | 8.836 | 0.439 | 0.953 | 0.047 | 0.078 | -0.948 | 0.000 | -0.948 | Amensalism |
| *Enterobacter cloacae* | *Salmonella enterica* | Flight 3 | 1 | 8.182 | 8.372 | 8.654 | 0.402 | 0.956 | 0.044 | 0.058 | -0.952 | 0.000 | -0.952 | Amensalism |
| *Enterobacter cloacae* | *Salmonella enterica* | Flight 3 | 5 | 6.106 | 5.907 | 6.517 | 0.136 | 0.980 | 0.020 | 0.067 | -0.977 | 0.000 | -0.977 | Amensalism |
| *Enterobacter cloacae* | *Salmonella enterica* | Flight 3 | 7 | 6.106 | 5.907 | 6.517 | 0.136 | 0.980 | 0.020 | 0.067 | -0.977 | 0.000 | -0.977 | Amensalism |
| *Enterobacter cloacae* | *Salmonella enterica* | Flight 3 | 8 | 6.106 | 5.907 | 6.517 | 0.136 | 0.980 | 0.020 | 0.067 | -0.977 | 0.000 | -0.977 | Amensalism |
| *Enterobacter cloacae* | *Shigella sonnei* | Flight 1 | 2 | 8.200 | 8.331 | 8.218 | 0.157 | 0.981 | 0.019 | 0.002 | -0.981 | 0.000 | -0.981 | Amensalism |
| *Enterobacter hormaechei* | *Enterobacter roggenkampii* | Flight 1 | 2 | 8.200 | 8.200 | 7.803 | 0.397 | 0.952 | 0.048 | -0.048 | -0.952 | 0.000 | -0.952 | Amensalism |
| *Enterobacter hormaechei* | *Enterobacter sp. strain NFIX59* | Flight 1 | 2 | 8.200 | 8.398 | 8.318 | 0.084 | 0.990 | 0.010 | 0.014 | -0.990 | 0.000 | -0.990 | Amensalism |
| *Enterobacter hormaechei* | *Escherichia coli* | Flight 1 | 2 | 8.200 | 8.331 | 8.218 | 0.157 | 0.981 | 0.019 | 0.002 | -0.981 | 0.000 | -0.981 | Amensalism |
| *Enterobacter hormaechei* | *Escherichia coli* | Flight 3 | 4 | 6.106 | 6.230 | 6.137 | 0.114 | 0.982 | 0.018 | 0.005 | -0.982 | 0.000 | -0.982 | Amensalism |
| *Enterobacter hormaechei* | *Klebsiella oxytoca* | Flight 1 | 2 | 8.200 | 9.403 | 9.013 | 0.415 | 0.956 | 0.044 | 0.099 | -0.956 | 0.000 | -0.956 | Amensalism |
| *Enterobacter hormaechei* | *Klebsiella pneumoniae* | Flight 1 | 2 | 8.200 | 9.251 | 8.835 | 0.439 | 0.953 | 0.047 | 0.077 | -0.953 | 0.000 | -0.953 | Amensalism |
| *Enterobacter hormaechei* | *Klebsiella pneumoniae* | Flight 3 | 4 | 6.106 | 6.623 | 6.517 | 0.136 | 0.980 | 0.020 | 0.067 | -0.979 | 0.000 | -0.979 | Amensalism |
| *Enterobacter hormaechei* | *Klebsiella quasipneumoniae strain IF2SW-B3* | Flight 1 | 2 | 8.200 | 9.251 | 8.835 | 0.439 | 0.953 | 0.047 | 0.077 | -0.953 | 0.000 | -0.953 | Amensalism |
| *Enterobacter hormaechei* | *Klebsiella quasipneumoniae strain IF2SW-P1* | Flight 1 | 2 | 8.200 | 9.251 | 8.835 | 0.439 | 0.953 | 0.047 | 0.077 | -0.953 | 0.000 | -0.953 | Amensalism |
| *Enterobacter hormaechei* | *Pantoea ananatis* | Flight 1 | 2 | 8.200 | 7.840 | 9.050 | 10.087 | 0.473 | 0.527 | 0.104 | 0.287 | 0.104 | 0.287 | Mutualism |
| *Enterobacter hormaechei* | *Pantoea ananatis* | Flight 3 | 4 | 6.106 | 5.534 | 6.946 | 7.805 | 0.471 | 0.529 | 0.138 | 0.410 | 0.138 | 0.410 | Mutualism |
| *Enterobacter hormaechei* | *Pantoea dispersa* | Flight 3 | 4 | 6.106 | 1.869 | 6.044 | 0.390 | 0.939 | 0.061 | -0.010 | -0.791 | 0.000 | -0.791 | Amensalism |
| *Enterobacter hormaechei* | *Penicillium rubens* | Flight 1 | 2 | 8.200 | 0.049 | 8.200 | 0.000 | 1.000 | 0.000 | 0.000 | -1.000 | 0.000 | -1.000 | Amensalism |

| Microorganism A | Microorganism B | Flight | Location | Vbio of A | Vbio of B | Vbio of A in AB | Vbio of B in AB | Ratio of A in AB | Ratio of B in AB | Effect of AB on A | Effect of AB on B | Significant effect on A | Significant effect on B | Interaction type |
|---|---|---|---|---|---|---|---|---|---|---|---|---|---|---|
| *Enterobacter hormaechei* | *Rahnella aquatilis* | Flight 3 | 4 | 6.106 | 2.624 | 5.889 | 0.424 | 0.933 | 0.067 | -0.036 | -0.838 | 0.000 | -0.838 | Amensalism |
| *Enterobacter hormaechei* | *Rhodotorula sp. strain JG-1b* | Flight 1 | 2 | 8.200 | 0.153 | 8.183 | 0.150 | 0.982 | 0.018 | -0.002 | -0.021 | 0.000 | 0.000 | Neutral |
| *Enterobacter hormaechei* | *Salmonella enterica* | Flight 1 | 2 | 8.200 | 8.478 | 8.680 | 0.410 | 0.955 | 0.045 | 0.059 | -0.952 | 0.000 | -0.952 | Amensalism |
| *Enterobacter hormaechei* | *Shigella sonnei* | Flight 1 | 2 | 8.200 | 8.331 | 8.218 | 0.157 | 0.981 | 0.019 | 0.002 | -0.981 | 0.000 | -0.981 | Amensalism |
| *Enterobacter roggenkampii* | *Enterobacter sp. strain NFIX59* | Flight 1 | 2 | 8.200 | 8.398 | 8.318 | 0.084 | 0.990 | 0.010 | 0.014 | -0.990 | 0.000 | -0.990 | Amensalism |
| *Enterobacter roggenkampii* | *Escherichia coli* | Flight 1 | 2 | 8.200 | 8.331 | 8.218 | 0.157 | 0.981 | 0.019 | 0.002 | -0.981 | 0.000 | -0.981 | Amensalism |
| *Enterobacter roggenkampii* | *Klebsiella oxytoca* | Flight 1 | 2 | 8.200 | 9.403 | 9.013 | 0.415 | 0.956 | 0.044 | 0.099 | -0.956 | 0.000 | -0.956 | Amensalism |
| *Enterobacter roggenkampii* | *Klebsiella pneumoniae* | Flight 1 | 2 | 8.200 | 9.251 | 8.835 | 0.439 | 0.953 | 0.047 | 0.077 | -0.953 | 0.000 | -0.953 | Amensalism |
| *Enterobacter roggenkampii* | *Klebsiella quasipneumoniae strain IF2SW-B3* | Flight 1 | 2 | 8.200 | 9.251 | 8.835 | 0.439 | 0.953 | 0.047 | 0.077 | -0.953 | 0.000 | -0.953 | Amensalism |
| *Enterobacter roggenkampii* | *Klebsiella quasipneumoniae strain IF2SW-P1* | Flight 1 | 2 | 8.200 | 9.251 | 8.835 | 0.439 | 0.953 | 0.047 | 0.077 | -0.953 | 0.000 | -0.953 | Amensalism |
| *Enterobacter roggenkampii* | *Pantoea ananatis* | Flight 1 | 2 | 8.200 | 7.840 | 9.050 | 10.087 | 0.473 | 0.527 | 0.104 | 0.287 | 0.104 | 0.287 | Mutualism |
| *Enterobacter roggenkampii* | *Penicillium rubens* | Flight 1 | 2 | 8.200 | 0.049 | 8.200 | 0.000 | 1.000 | 0.000 | 0.000 | -1.000 | 0.000 | -1.000 | Amensalism |
| *Enterobacter roggenkampii* | *Rhodotorula sp. strain JG-1b* | Flight 1 | 2 | 8.200 | 0.153 | 8.183 | 0.150 | 0.982 | 0.018 | -0.002 | -0.021 | 0.000 | 0.000 | Neutral |
| *Enterobacter roggenkampii* | *Salmonella enterica* | Flight 1 | 2 | 8.200 | 8.478 | 8.836 | 0.439 | 0.953 | 0.047 | 0.078 | -0.948 | 0.000 | -0.948 | Amensalism |
| *Enterobacter roggenkampii* | *Shigella sonnei* | Flight 1 | 2 | 8.200 | 8.331 | 8.218 | 0.157 | 0.981 | 0.019 | 0.002 | -0.981 | 0.000 | -0.981 | Amensalism |
| *Enterobacter sp. strain NFIX59* | *Escherichia coli* | Flight 1 | 2 | 8.398 | 8.331 | 0.086 | 8.490 | 0.010 | 0.990 | -0.990 | 0.019 | -0.990 | 0.000 | Amensalism |
| *Enterobacter sp. strain NFIX59* | *Klebsiella oxytoca* | Flight 1 | 2 | 8.398 | 9.403 | 0.096 | 9.510 | 0.010 | 0.990 | -0.989 | 0.011 | -0.989 | 0.000 | Amensalism |
| *Enterobacter sp. strain NFIX59* | *Klebsiella pneumoniae* | Flight 1 | 2 | 8.398 | 9.251 | 0.095 | 9.359 | 0.010 | 0.990 | -0.989 | 0.012 | -0.989 | 0.000 | Amensalism |
| *Enterobacter sp. strain NFIX59* | *Klebsiella quasipneumoniae strain IF2SW-B3* | Flight 1 | 2 | 8.398 | 9.251 | 0.095 | 9.359 | 0.010 | 0.990 | -0.989 | 0.012 | -0.989 | 0.000 | Amensalism |
| *Enterobacter sp. strain NFIX59* | *Klebsiella quasipneumoniae strain IF2SW-P1* | Flight 1 | 2 | 8.398 | 9.251 | 0.095 | 9.359 | 0.010 | 0.990 | -0.989 | 0.012 | -0.989 | 0.000 | Amensalism |
| *Enterobacter sp. strain NFIX59* | *Pantoea ananatis* | Flight 1 | 2 | 8.398 | 7.840 | 9.048 | 10.085 | 0.473 | 0.527 | 0.077 | 0.286 | 0.000 | 0.286 | Commensals |
| *Enterobacter sp. strain NFIX59* | *Penicillium rubens* | Flight 1 | 2 | 8.398 | 0.049 | 8.398 | 0.000 | 1.000 | 0.000 | 0.000 | -1.000 | 0.000 | -1.000 | Amensalism |
| *Enterobacter sp. strain NFIX59* | *Rhodotorula sp. strain JG-1b* | Flight 1 | 2 | 8.398 | 0.153 | 8.398 | 0.000 | 1.000 | 0.000 | 0.000 | -1.000 | 0.000 | -1.000 | Amensalism |
| *Enterobacter sp. strain NFIX59* | *Salmonella enterica* | Flight 1 | 2 | 8.398 | 8.478 | 0.176 | 9.282 | 0.019 | 0.981 | -0.979 | 0.095 | -0.979 | 0.000 | Amensalism |
| *Enterobacter sp. strain NFIX59* | *Shigella sonnei* | Flight 1 | 2 | 8.398 | 8.331 | 0.086 | 8.490 | 0.010 | 0.990 | -0.990 | 0.019 | -0.990 | 0.000 | Amensalism |
| *Enterococcus avium* | *Enterococcus faecalis* | Flight 3 | 2 | 1.824 | 1.032 | 2.058 | 0.021 | 0.990 | 0.010 | 0.129 | -0.980 | 0.129 | -0.980 | Parasitism |
| *Enterococcus avium* | *Klebsiella pneumoniae strain F3-2P(2*)* | Flight 3 | 2 | 1.824 | 12.151 | 3.431 | 10.472 | 0.247 | 0.753 | 0.881 | -0.138 | 0.881 | -0.138 | Parasitism |
| *Enterococcus avium* | *Paenibacillus polymyxa* | Flight 3 | 2 | 1.824 | 1.781 | 2.779 | 0.028 | 0.990 | 0.010 | 0.524 | -0.984 | 0.524 | -0.984 | Parasitism |
| *Enterococcus avium* | *Penicillium chrysogenum* | Flight 3 | 2 | 1.824 | 0.049 | 2.354 | 0.056 | 0.977 | 0.023 | 0.291 | 0.131 | 0.291 | 0.131 | Mutualism |
| *Enterococcus avium* | *Penicillium flavigenum* | Flight 3 | 2 | 1.824 | 0.097 | 2.198 | 0.029 | 0.987 | 0.013 | 0.205 | -0.702 | 0.205 | -0.702 | Parasitism |
| *Enterococcus avium* | *Penicillium nalgiovense* | Flight 3 | 2 | 1.824 | 0.067 | 2.165 | 0.033 | 0.985 | 0.015 | 0.187 | -0.514 | 0.187 | -0.514 | Parasitism |
| *Enterococcus avium* | *Penicillium rubens* | Flight 3 | 2 | 1.824 | 0.049 | 2.354 | 0.056 | 0.977 | 0.023 | 0.291 | 0.131 | 0.291 | 0.131 | Mutualism |
| *Enterococcus avium* | *Rhodotorula sp. strain JG-1b* | Flight 3 | 2 | 1.824 | 0.156 | 1.412 | 1.069 | 0.569 | 0.431 | -0.226 | 5.842 | -0.226 | 5.842 | Parasitism |
| *Enterococcus avium* | *Staphylococcus aureus* | Flight 3 | 2 | 1.824 | 2.739 | 4.602 | 0.241 | 0.950 | 0.050 | 1.523 | -0.912 | 1.523 | -0.912 | Parasitism |
| *Enterococcus avium* | *Staphylococcus epidermidis* | Flight 3 | 2 | 1.824 | 2.313 | 4.167 | 0.218 | 0.950 | 0.050 | 1.285 | -0.906 | 1.285 | -0.906 | Parasitism |
| *Enterococcus avium* | *Staphylococcus haemolyticus* | Flight 3 | 2 | 1.824 | 2.519 | 4.509 | 0.236 | 0.950 | 0.050 | 1.473 | -0.906 | 1.473 | -0.906 | Parasitism |
| *Enterococcus avium* | *Staphylococcus saprophyticus* | Flight 3 | 2 | 1.824 | 2.363 | 4.314 | 0.226 | 0.950 | 0.050 | 1.365 | -0.904 | 1.365 | -0.904 | Parasitism |
| *Enterococcus avium* | *Staphylococcus sp. strain LCT-H4* | Flight 3 | 2 | 1.824 | 2.388 | 2.018 | 2.698 | 0.428 | 0.572 | 0.107 | 0.130 | 0.107 | 0.130 | Mutualism |
| *Enterococcus avium* | *Staphylococcus warneri* | Flight 3 | 2 | 1.824 | 2.543 | 4.124 | 0.765 | 0.844 | 0.156 | 1.262 | -0.699 | 1.262 | -0.699 | Parasitism |
| *Enterococcus faecalis* | *Klebsiella pneumoniae strain F3-2P(2*)* | Flight 3 | 2 | 1.032 | 12.151 | 4.532 | 13.603 | 0.250 | 0.750 | 3.391 | 0.119 | 3.391 | 0.119 | Mutualism |
| *Enterococcus faecalis* | *Paenibacillus polymyxa* | Flight 3 | 2 | 1.032 | 1.781 | 1.802 | 0.358 | 0.834 | 0.166 | 0.746 | -0.799 | 0.746 | -0.799 | Parasitism |
| *Enterococcus faecalis* | *Penicillium chrysogenum* | Flight 3 | 2 | 1.032 | 0.049 | 1.465 | 0.028 | 0.981 | 0.019 | 0.419 | -0.438 | 0.419 | -0.438 | Parasitism |
| *Enterococcus faecalis* | *Penicillium flavigenum* | Flight 3 | 2 | 1.032 | 0.097 | 1.516 | 0.015 | 0.990 | 0.010 | 0.468 | -0.842 | 0.468 | -0.842 | Parasitism |
| *Enterococcus faecalis* | *Penicillium nalgiovense* | Flight 3 | 2 | 1.032 | 0.067 | 1.459 | 0.020 | 0.987 | 0.013 | 0.414 | -0.707 | 0.414 | -0.707 | Parasitism |
| *Enterococcus faecalis* | *Penicillium rubens* | Flight 3 | 2 | 1.032 | 0.049 | 1.465 | 0.028 | 0.981 | 0.019 | 0.419 | -0.438 | 0.419 | -0.438 | Parasitism |
| *Enterococcus faecalis* | *Rhodotorula sp. strain JG-1b* | Flight 3 | 2 | 1.032 | 0.156 | 0.989 | 0.522 | 0.655 | 0.345 | -0.041 | 2.343 | 0.000 | 2.343 | Commensals |
| *Enterococcus faecalis* | *Staphylococcus aureus* | Flight 3 | 2 | 1.032 | 2.739 | 0.160 | 3.057 | 0.050 | 0.950 | -0.845 | 0.116 | -0.845 | 0.116 | Parasitism |
| *Enterococcus faecalis* | *Staphylococcus epidermidis* | Flight 3 | 2 | 1.032 | 2.313 | 0.218 | 2.608 | 0.077 | 0.923 | -0.788 | 0.128 | -0.788 | 0.128 | Parasitism |
| *Enterococcus faecalis* | *Staphylococcus haemolyticus* | Flight 3 | 2 | 1.032 | 2.519 | 0.151 | 2.882 | 0.050 | 0.950 | -0.854 | 0.144 | -0.854 | 0.144 | Parasitism |
| *Enterococcus faecalis* | *Staphylococcus saprophyticus* | Flight 3 | 2 | 1.032 | 2.363 | 0.141 | 2.682 | 0.050 | 0.950 | -0.864 | 0.135 | -0.864 | 0.135 | Parasitism |
| *Enterococcus faecalis* | *Staphylococcus sp. strain LCT-H4* | Flight 3 | 2 | 1.032 | 2.388 | 0.145 | 2.757 | 0.050 | 0.950 | -0.860 | 0.155 | -0.860 | 0.155 | Parasitism |
| *Enterococcus faecalis* | *Staphylococcus warneri* | Flight 3 | 2 | 1.032 | 2.543 | 0.149 | 2.848 | 0.050 | 0.950 | -0.855 | 0.120 | -0.855 | 0.120 | Parasitism |
| *Escherichia coli* | *Klebsiella aerogenes strain IIIF7SW-P1* | Flight 3 | 7 | 6.230 | 7.338 | 10.685 | 5.212 | 0.672 | 0.328 | 0.715 | -0.290 | 0.715 | -0.290 | Parasitism |
| *Escherichia coli* | *Klebsiella oxytoca* | Flight 1 | 2 | 8.331 | 9.403 | 8.988 | 0.418 | 0.956 | 0.044 | 0.079 | -0.956 | 0.000 | -0.956 | Amensalism |
| *Escherichia coli* | *Klebsiella pneumoniae* | Flight 1 | 2 | 8.331 | 9.251 | 8.988 | 0.418 | 0.956 | 0.044 | 0.079 | -0.955 | 0.000 | -0.955 | Amensalism |
| *Escherichia coli* | *Klebsiella pneumoniae* | Flight 1 | 5 | 8.331 | 9.251 | 8.988 | 0.418 | 0.956 | 0.044 | 0.079 | -0.955 | 0.000 | -0.955 | Amensalism |
| *Escherichia coli* | *Klebsiella pneumoniae* | Flight 2 | 5 | 6.492 | 6.981 | 7.034 | 0.143 | 0.980 | 0.020 | 0.084 | -0.979 | 0.000 | -0.979 | Amensalism |
| *Escherichia coli* | *Klebsiella pneumoniae* | Flight 3 | 1 | 8.314 | 9.051 | 8.809 | 0.376 | 0.959 | 0.041 | 0.059 | -0.959 | 0.000 | -0.959 | Amensalism |
| *Escherichia coli* | *Klebsiella pneumoniae* | Flight 3 | 3 | 6.864 | 7.662 | 7.550 | 0.269 | 0.966 | 0.034 | 0.100 | -0.965 | 0.100 | -0.965 | Parasitism |
| *Escherichia coli* | *Klebsiella pneumoniae* | Flight 3 | 4 | 6.230 | 6.623 | 6.700 | 0.118 | 0.983 | 0.017 | 0.075 | -0.982 | 0.000 | -0.982 | Amensalism |
| *Escherichia coli* | *Klebsiella pneumoniae* | Flight 3 | 5 | 6.230 | 6.623 | 6.700 | 0.118 | 0.983 | 0.017 | 0.075 | -0.982 | 0.000 | -0.982 | Amensalism |
| *Escherichia coli* | *Klebsiella pneumoniae* | Flight 3 | 7 | 6.230 | 6.623 | 6.700 | 0.118 | 0.983 | 0.017 | 0.075 | -0.982 | 0.000 | -0.982 | Amensalism |
| *Escherichia coli* | *Klebsiella pneumoniae* | Flight 3 | 8 | 6.230 | 6.623 | 6.700 | 0.118 | 0.983 | 0.017 | 0.075 | -0.982 | 0.000 | -0.982 | Amensalism |
| *Escherichia coli* | *Klebsiella quasipneumoniae strain IF2SW-B3* | Flight 1 | 2 | 8.331 | 9.251 | 8.988 | 0.418 | 0.956 | 0.044 | 0.079 | -0.955 | 0.000 | -0.955 | Amensalism |
| *Escherichia coli* | *Klebsiella quasipneumoniae strain IF2SW-P1* | Flight 1 | 2 | 8.331 | 9.251 | 8.988 | 0.418 | 0.956 | 0.044 | 0.079 | -0.955 | 0.000 | -0.955 | Amensalism |
| *Escherichia coli* | *Klebsiella quasipneumoniae strain IIIF3SW-P1* | Flight 3 | 3 | 6.864 | 7.662 | 7.550 | 0.269 | 0.966 | 0.034 | 0.100 | -0.965 | 0.100 | -0.965 | Parasitism |

| Microorganism A | Microorganism B | Flight | Location | Vbio of A | Vbio of B | Vbio of A in AB | Vbio of B in AB | Ratio of A in AB | Ratio of B in AB | Effect of AB on A | Effect of AB on B | Significant effect on A | Significant effect on B | Interaction type |
|---|---|---|---|---|---|---|---|---|---|---|---|---|---|---|
| *Escherichia coli* | *Klebsiella sp. strain MS 92-3* | Flight 3 | 3 | 6.864 | 7.756 | 7.747 | 0.270 | 0.966 | 0.034 | 0.129 | -0.965 | 0.129 | -0.965 | Parasitism |
| *Escherichia coli* | *Klebsiella variicola* | Flight 3 | 3 | 6.864 | 7.668 | 7.569 | 0.269 | 0.966 | 0.034 | 0.103 | -0.965 | 0.103 | -0.965 | Parasitism |
| *Escherichia coli* | *Paenibacillus polymyxa strain IIF5SW-B3* | Flight 2 | 5 | 6.492 | 0.765 | 8.763 | 2.594 | 0.772 | 0.228 | 0.350 | 2.390 | 0.350 | 2.390 | Mutualism |
| *Escherichia coli* | *Paenibacillus polymyxa strain IIF5SW-B4* | Flight 2 | 5 | 6.492 | 0.765 | 8.763 | 2.594 | 0.772 | 0.228 | 0.350 | 2.390 | 0.350 | 2.390 | Mutualism |
| *Escherichia coli* | *Pantoea agglomerans* | Flight 1 | 5 | 8.331 | 5.233 | 8.896 | 0.756 | 0.922 | 0.078 | 0.068 | -0.855 | 0.000 | -0.855 | Amensalism |
| *Escherichia coli* | *Pantoea agglomerans* | Flight 2 | 5 | 6.492 | 3.310 | 7.015 | 0.411 | 0.945 | 0.055 | 0.081 | -0.876 | 0.000 | -0.876 | Amensalism |
| *Escherichia coli* | *Pantoea agglomerans* | Flight 3 | 1 | 8.314 | 4.640 | 8.635 | 0.589 | 0.936 | 0.064 | 0.039 | -0.873 | 0.000 | -0.873 | Amensalism |
| *Escherichia coli* | *Pantoea agglomerans* | Flight 3 | 5 | 6.230 | 2.777 | 6.761 | 0.369 | 0.948 | 0.052 | 0.085 | -0.867 | 0.000 | -0.867 | Amensalism |
| *Escherichia coli* | *Pantoea agglomerans* | Flight 3 | 7 | 6.230 | 2.777 | 6.761 | 0.369 | 0.948 | 0.052 | 0.085 | -0.867 | 0.000 | -0.867 | Amensalism |
| *Escherichia coli* | *Pantoea agglomerans* | Flight 3 | 8 | 6.230 | 2.777 | 6.761 | 0.369 | 0.948 | 0.052 | 0.085 | -0.867 | 0.000 | -0.867 | Amensalism |
| *Escherichia coli* | *Pantoea ananatis* | Flight 1 | 2 | 8.331 | 7.840 | 9.036 | 10.072 | 0.473 | 0.527 | 0.085 | 0.285 | 0.000 | 0.285 | Commensals |
| *Escherichia coli* | *Pantoea ananatis* | Flight 1 | 5 | 8.331 | 7.840 | 9.036 | 10.072 | 0.473 | 0.527 | 0.085 | 0.285 | 0.000 | 0.285 | Commensals |
| *Escherichia coli* | *Pantoea ananatis* | Flight 2 | 5 | 6.492 | 6.254 | 7.551 | 8.491 | 0.471 | 0.529 | 0.163 | 0.358 | 0.163 | 0.358 | Mutualism |
| *Escherichia coli* | *Pantoea ananatis* | Flight 3 | 1 | 8.314 | 7.622 | 8.932 | 9.958 | 0.473 | 0.527 | 0.074 | 0.306 | 0.000 | 0.306 | Commensals |
| *Escherichia coli* | *Pantoea ananatis* | Flight 3 | 4 | 6.230 | 5.534 | 6.877 | 7.735 | 0.471 | 0.529 | 0.104 | 0.398 | 0.104 | 0.398 | Mutualism |
| *Escherichia coli* | *Pantoea ananatis* | Flight 3 | 5 | 6.230 | 5.534 | 6.877 | 7.735 | 0.471 | 0.529 | 0.104 | 0.398 | 0.104 | 0.398 | Mutualism |
| *Escherichia coli* | *Pantoea ananatis* | Flight 3 | 7 | 6.230 | 5.534 | 6.877 | 7.735 | 0.471 | 0.529 | 0.104 | 0.398 | 0.104 | 0.398 | Mutualism |
| *Escherichia coli* | *Pantoea ananatis* | Flight 3 | 8 | 6.230 | 5.534 | 6.877 | 7.735 | 0.471 | 0.529 | 0.104 | 0.398 | 0.104 | 0.398 | Mutualism |
| *Escherichia coli* | *Pantoea conspicua* | Flight 1 | 5 | 8.331 | 4.632 | 8.071 | 1.728 | 0.824 | 0.176 | -0.031 | -0.627 | 0.000 | -0.627 | Amensalism |
| *Escherichia coli* | *Pantoea conspicua* | Flight 2 | 5 | 6.492 | 3.267 | 6.766 | 0.805 | 0.894 | 0.106 | 0.042 | -0.753 | 0.000 | -0.753 | Amensalism |
| *Escherichia coli* | *Pantoea conspicua* | Flight 3 | 1 | 8.314 | 4.408 | 8.551 | 0.690 | 0.925 | 0.075 | 0.028 | -0.843 | 0.000 | -0.843 | Amensalism |
| *Escherichia coli* | *Pantoea conspicua* | Flight 3 | 5 | 6.230 | 2.768 | 6.726 | 0.426 | 0.940 | 0.060 | 0.080 | -0.846 | 0.000 | -0.846 | Amensalism |
| *Escherichia coli* | *Pantoea conspicua* | Flight 3 | 7 | 6.230 | 2.768 | 6.726 | 0.426 | 0.940 | 0.060 | 0.080 | -0.846 | 0.000 | -0.846 | Amensalism |
| *Escherichia coli* | *Pantoea conspicua* | Flight 3 | 8 | 6.230 | 2.768 | 6.726 | 0.426 | 0.940 | 0.060 | 0.080 | -0.846 | 0.000 | -0.846 | Amensalism |
| *Escherichia coli* | *Pantoea dispersa* | Flight 3 | 4 | 6.230 | 1.869 | 6.226 | 0.339 | 0.948 | 0.052 | -0.001 | -0.819 | 0.000 | -0.819 | Amensalism |
| *Escherichia coli* | *Pantoea sp. strain 3.5.1* | Flight 1 | 5 | 8.331 | 5.233 | 8.896 | 0.756 | 0.922 | 0.078 | 0.068 | -0.855 | 0.000 | -0.855 | Amensalism |
| *Escherichia coli* | *Pantoea sp. strain 3.5.1* | Flight 2 | 5 | 6.492 | 3.310 | 7.015 | 0.411 | 0.945 | 0.055 | 0.081 | -0.876 | 0.000 | -0.876 | Amensalism |
| *Escherichia coli* | *Pantoea sp. strain A4* | Flight 3 | 1 | 8.314 | 4.256 | 8.635 | 0.589 | 0.936 | 0.064 | 0.039 | -0.862 | 0.000 | -0.862 | Amensalism |
| *Escherichia coli* | *Pantoea sp. strain A4* | Flight 3 | 5 | 6.230 | 2.593 | 6.761 | 0.369 | 0.948 | 0.052 | 0.085 | -0.858 | 0.000 | -0.858 | Amensalism |
| *Escherichia coli* | *Pantoea sp. strain A4* | Flight 3 | 7 | 6.230 | 2.593 | 6.761 | 0.369 | 0.948 | 0.052 | 0.085 | -0.858 | 0.000 | -0.858 | Amensalism |
| *Escherichia coli* | *Pantoea sp. strain A4* | Flight 3 | 8 | 6.230 | 2.593 | 6.761 | 0.369 | 0.948 | 0.052 | 0.085 | -0.858 | 0.000 | -0.858 | Amensalism |
| *Escherichia coli* | *Pantoea sp. strain At-9b* | Flight 3 | 1 | 8.314 | 3.804 | 8.860 | 0.263 | 0.971 | 0.029 | 0.066 | -0.931 | 0.000 | -0.931 | Amensalism |
| *Escherichia coli* | *Pantoea sp. strain At-9b* | Flight 3 | 5 | 6.230 | 2.173 | 6.725 | 0.254 | 0.964 | 0.036 | 0.079 | -0.883 | 0.000 | -0.883 | Amensalism |
| *Escherichia coli* | *Pantoea sp. strain At-9b* | Flight 3 | 8 | 6.230 | 2.173 | 6.725 | 0.254 | 0.964 | 0.036 | 0.079 | -0.883 | 0.000 | -0.883 | Amensalism |
| *Escherichia coli* | *Pantoea sp. strain FF5* | Flight 3 | 1 | 8.314 | 4.640 | 8.635 | 0.589 | 0.936 | 0.064 | 0.039 | -0.873 | 0.000 | -0.873 | Amensalism |
| *Escherichia coli* | *Pantoea sp. strain FF5* | Flight 3 | 5 | 6.230 | 2.777 | 6.761 | 0.369 | 0.948 | 0.052 | 0.085 | -0.867 | 0.000 | -0.867 | Amensalism |
| *Escherichia coli* | *Pantoea sp. strain FF5* | Flight 3 | 7 | 6.230 | 2.777 | 6.761 | 0.369 | 0.948 | 0.052 | 0.085 | -0.867 | 0.000 | -0.867 | Amensalism |
| *Escherichia coli* | *Pantoea sp. strain FF5* | Flight 3 | 8 | 6.230 | 2.777 | 6.761 | 0.369 | 0.948 | 0.052 | 0.085 | -0.867 | 0.000 | -0.867 | Amensalism |
| *Escherichia coli* | *Pantoea sp. strain IMH* | Flight 3 | 1 | 8.314 | 4.409 | 7.594 | 1.291 | 0.855 | 0.145 | -0.087 | -0.707 | 0.000 | -0.707 | Amensalism |
| *Escherichia coli* | *Pantoea sp. strain IMH* | Flight 3 | 5 | 6.230 | 2.634 | 6.015 | 0.710 | 0.894 | 0.106 | -0.035 | -0.730 | 0.000 | -0.730 | Amensalism |
| *Escherichia coli* | *Pantoea sp. strain IMH* | Flight 3 | 7 | 6.230 | 2.634 | 6.015 | 0.710 | 0.894 | 0.106 | -0.035 | -0.730 | 0.000 | -0.730 | Amensalism |
| *Escherichia coli* | *Pantoea sp. strain IMH* | Flight 3 | 8 | 6.230 | 2.634 | 6.015 | 0.710 | 0.894 | 0.106 | -0.035 | -0.730 | 0.000 | -0.730 | Amensalism |
| *Escherichia coli* | *Pantoea sp. strain NGS-ED-1003* | Flight 3 | 1 | 8.314 | 4.640 | 8.635 | 0.589 | 0.936 | 0.064 | 0.039 | -0.873 | 0.000 | -0.873 | Amensalism |
| *Escherichia coli* | *Pantoea sp. strain NGS-ED-1003* | Flight 3 | 5 | 6.230 | 2.777 | 6.761 | 0.369 | 0.948 | 0.052 | 0.085 | -0.867 | 0.000 | -0.867 | Amensalism |
| *Escherichia coli* | *Pantoea sp. strain NGS-ED-1003* | Flight 3 | 7 | 6.230 | 2.777 | 6.761 | 0.369 | 0.948 | 0.052 | 0.085 | -0.867 | 0.000 | -0.867 | Amensalism |
| *Escherichia coli* | *Pantoea sp. strain NGS-ED-1003* | Flight 3 | 8 | 6.230 | 2.777 | 6.761 | 0.369 | 0.948 | 0.052 | 0.085 | -0.867 | 0.000 | -0.867 | Amensalism |
| *Escherichia coli* | *Pantoea sp. strain OXWO6B1* | Flight 3 | 1 | 8.314 | 4.642 | 8.636 | 0.589 | 0.936 | 0.064 | 0.039 | -0.873 | 0.000 | -0.873 | Amensalism |
| *Escherichia coli* | *Pantoea sp. strain OXWO6B1* | Flight 3 | 5 | 6.230 | 2.779 | 6.761 | 0.369 | 0.948 | 0.052 | 0.085 | -0.867 | 0.000 | -0.867 | Amensalism |
| *Escherichia coli* | *Pantoea sp. strain OXWO6B1* | Flight 3 | 7 | 6.230 | 2.779 | 6.761 | 0.369 | 0.948 | 0.052 | 0.085 | -0.867 | 0.000 | -0.867 | Amensalism |
| *Escherichia coli* | *Pantoea sp. strain OXWO6B1* | Flight 3 | 8 | 6.230 | 2.779 | 6.761 | 0.369 | 0.948 | 0.052 | 0.085 | -0.867 | 0.000 | -0.867 | Amensalism |
| *Escherichia coli* | *Pantoea vagans* | Flight 2 | 5 | 6.492 | 3.312 | 7.016 | 0.411 | 0.945 | 0.055 | 0.081 | -0.876 | 0.000 | -0.876 | Amensalism |
| *Escherichia coli* | *Pantoea vagans* | Flight 3 | 1 | 8.314 | 4.642 | 8.636 | 0.589 | 0.936 | 0.064 | 0.039 | -0.873 | 0.000 | -0.873 | Amensalism |
| *Escherichia coli* | *Pantoea vagans* | Flight 3 | 5 | 6.230 | 2.778 | 6.761 | 0.369 | 0.948 | 0.052 | 0.085 | -0.867 | 0.000 | -0.867 | Amensalism |
| *Escherichia coli* | *Pantoea vagans* | Flight 3 | 7 | 6.230 | 2.778 | 6.761 | 0.369 | 0.948 | 0.052 | 0.085 | -0.867 | 0.000 | -0.867 | Amensalism |
| *Escherichia coli* | *Pantoea vagans* | Flight 3 | 8 | 6.230 | 2.778 | 6.761 | 0.369 | 0.948 | 0.052 | 0.085 | -0.867 | 0.000 | -0.867 | Amensalism |
| *Escherichia coli* | *Penicillium rubens* | Flight 1 | 2 | 8.331 | 0.049 | 8.331 | 0.000 | 1.000 | 0.000 | 0.000 | -1.000 | 0.000 | -1.000 | Amensalism |
| *Escherichia coli* | *Penicillium rubens* | Flight 1 | 5 | 8.331 | 0.049 | 8.331 | 0.000 | 1.000 | 0.000 | 0.000 | -1.000 | 0.000 | -1.000 | Amensalism |
| *Escherichia coli* | *Penicillium rubens* | Flight 3 | 3 | 6.864 | 0.023 | 6.864 | 0.000 | 1.000 | 0.000 | 0.000 | -1.000 | 0.000 | -1.000 | Amensalism |
| *Escherichia coli* | *Rahnella aquatilis* | Flight 3 | 4 | 6.230 | 2.624 | 6.083 | 0.333 | 0.948 | 0.052 | -0.024 | -0.873 | 0.000 | -0.873 | Amensalism |
| *Escherichia coli* | *Rhodotorula sp. strain JG-1b* | Flight 1 | 2 | 8.331 | 0.153 | 8.344 | 0.152 | 0.982 | 0.018 | 0.001 | -0.003 | 0.000 | 0.000 | Neutral |
| *Escherichia coli* | *Rhodotorula sp. strain JG-1b* | Flight 1 | 5 | 8.331 | 0.153 | 8.344 | 0.152 | 0.982 | 0.018 | 0.001 | -0.003 | 0.000 | 0.000 | Neutral |
| *Escherichia coli* | *Rhodotorula sp. strain JG-1b* | Flight 3 | 1 | 8.314 | 0.154 | 8.314 | 0.000 | 1.000 | 0.000 | 0.000 | -1.000 | 0.000 | -1.000 | Amensalism |
| *Escherichia coli* | *Salmonella enterica* | Flight 1 | 2 | 8.331 | 8.478 | 8.988 | 0.417 | 0.956 | 0.044 | 0.079 | -0.951 | 0.000 | -0.951 | Amensalism |
| *Escherichia coli* | *Salmonella enterica* | Flight 3 | 1 | 8.314 | 8.372 | 0.120 | 9.061 | 0.013 | 0.987 | -0.986 | 0.082 | -0.986 | 0.000 | Amensalism |
| *Escherichia coli* | *Salmonella enterica* | Flight 3 | 3 | 6.864 | 6.819 | 7.468 | 0.244 | 0.968 | 0.032 | 0.088 | -0.964 | 0.000 | -0.964 | Amensalism |
| *Escherichia coli* | *Salmonella enterica* | Flight 3 | 5 | 6.230 | 5.907 | 6.618 | 0.113 | 0.983 | 0.017 | 0.062 | -0.981 | 0.000 | -0.981 | Amensalism |

| Microorganism A | Microorganism B | Flight | Location | Vbio of A | Vbio of B | Vbio of A in AB | Vbio of B in AB | Ratio of A in AB | Ratio of B in AB | Effect of AB on A | Effect of AB on B | Significant effect on A | Significant effect on B | Interaction type |
|---|---|---|---|---|---|---|---|---|---|---|---|---|---|---|
| *Escherichia coli* | *Salmonella enterica* | Flight 3 | 7 | 6.230 | 5.907 | 6.618 | 0.113 | 0.983 | 0.017 | 0.062 | -0.981 | 0.000 | -0.981 | Amensalism |
| *Escherichia coli* | *Salmonella enterica* | Flight 3 | 8 | 6.230 | 5.907 | 6.618 | 0.113 | 0.983 | 0.017 | 0.062 | -0.981 | 0.000 | -0.981 | Amensalism |
| *Escherichia coli* | *Shigella sonnei* | Flight 1 | 2 | 8.331 | 8.331 | 5.648 | 2.684 | 0.678 | 0.322 | -0.322 | -0.678 | -0.322 | -0.678 | Competitive |
| *Escherichia coli* | *Staphylococcus saprophyticus* | Flight 3 | 3 | 6.864 | 1.481 | 9.322 | 2.828 | 0.767 | 0.233 | 0.358 | 0.909 | 0.358 | 0.909 | Mutualism |
| *Klebsiella aerogenes strain IIIF7SW-P1* | *Klebsiella pneumoniae* | Flight 3 | 7 | 7.338 | 6.623 | 6.330 | 9.565 | 0.398 | 0.602 | -0.137 | 0.444 | -0.137 | 0.444 | Parasitism |
| *Klebsiella aerogenes strain IIIF7SW-P1* | *Pantoea agglomerans* | Flight 3 | 7 | 7.338 | 2.777 | 7.253 | 0.394 | 0.948 | 0.052 | -0.012 | -0.858 | 0.000 | -0.858 | Amensalism |
| *Klebsiella aerogenes strain IIIF7SW-P1* | *Pantoea ananatis* | Flight 3 | 7 | 7.338 | 5.534 | 7.377 | 0.122 | 0.984 | 0.016 | 0.005 | -0.978 | 0.000 | -0.978 | Amensalism |
| *Klebsiella aerogenes strain IIIF7SW-P1* | *Pantoea conspicua* | Flight 3 | 7 | 7.338 | 2.768 | 7.174 | 0.516 | 0.933 | 0.067 | -0.022 | -0.814 | 0.000 | -0.814 | Amensalism |
| *Klebsiella aerogenes strain IIIF7SW-P1* | *Pantoea sp. strain A4* | Flight 3 | 7 | 7.338 | 2.593 | 7.253 | 0.394 | 0.948 | 0.052 | -0.012 | -0.848 | 0.000 | -0.848 | Amensalism |
| *Klebsiella aerogenes strain IIIF7SW-P1* | *Pantoea sp. strain FF5* | Flight 3 | 7 | 7.338 | 2.777 | 7.253 | 0.394 | 0.948 | 0.052 | -0.012 | -0.858 | 0.000 | -0.858 | Amensalism |
| *Klebsiella aerogenes strain IIIF7SW-P1* | *Pantoea sp. strain IMH* | Flight 3 | 7 | 7.338 | 2.634 | 7.092 | 0.506 | 0.933 | 0.067 | -0.034 | -0.808 | 0.000 | -0.808 | Amensalism |
| *Klebsiella aerogenes strain IIIF7SW-P1* | *Pantoea sp. strain NGS-ED-1003* | Flight 3 | 7 | 7.338 | 2.777 | 7.253 | 0.394 | 0.948 | 0.052 | -0.012 | -0.858 | 0.000 | -0.858 | Amensalism |
| *Klebsiella aerogenes strain IIIF7SW-P1* | *Pantoea sp. strain OXWO6B1* | Flight 3 | 7 | 7.338 | 2.779 | 7.253 | 0.394 | 0.948 | 0.052 | -0.012 | -0.858 | 0.000 | -0.858 | Amensalism |
| *Klebsiella aerogenes strain IIIF7SW-P1* | *Pantoea vagans* | Flight 3 | 7 | 7.338 | 2.778 | 7.253 | 0.394 | 0.948 | 0.052 | -0.012 | -0.858 | 0.000 | -0.858 | Amensalism |
| *Klebsiella aerogenes strain IIIF7SW-P1* | *Salmonella enterica* | Flight 3 | 7 | 7.338 | 5.907 | 5.212 | 10.685 | 0.328 | 0.672 | -0.290 | 0.809 | -0.290 | 0.809 | Parasitism |
| *Klebsiella oxytoca* | *Klebsiella pneumoniae* | Flight 1 | 2 | 9.403 | 9.251 | 9.403 | 0.000 | 1.000 | 0.000 | 0.000 | -1.000 | 0.000 | -1.000 | Amensalism |
| *Klebsiella oxytoca* | *Klebsiella quasipneumoniae strain IF2SW-B3* | Flight 1 | 2 | 9.403 | 9.251 | 9.403 | 0.000 | 1.000 | 0.000 | 0.000 | -1.000 | 0.000 | -1.000 | Amensalism |
| *Klebsiella oxytoca* | *Klebsiella quasipneumoniae strain IF2SW-P1* | Flight 1 | 2 | 9.403 | 9.251 | 9.403 | 0.000 | 1.000 | 0.000 | 0.000 | -1.000 | 0.000 | -1.000 | Amensalism |
| *Klebsiella oxytoca* | *Pantoea ananatis* | Flight 1 | 2 | 9.403 | 7.840 | 9.107 | 10.150 | 0.473 | 0.527 | -0.031 | 0.295 | 0.000 | 0.295 | Commensals |
| *Klebsiella oxytoca* | *Penicillium rubens* | Flight 1 | 2 | 9.403 | 0.049 | 9.403 | 0.000 | 1.000 | 0.000 | 0.000 | -1.000 | 0.000 | -1.000 | Amensalism |
| *Klebsiella oxytoca* | *Rhodotorula sp. strain JG-1b* | Flight 1 | 2 | 9.403 | 0.153 | 9.393 | 0.171 | 0.982 | 0.018 | -0.001 | 0.116 | 0.000 | 0.116 | Commensals |
| *Klebsiella oxytoca* | *Salmonella enterica* | Flight 1 | 2 | 9.403 | 8.478 | 0.175 | 9.231 | 0.019 | 0.981 | -0.981 | 0.089 | -0.981 | 0.000 | Amensalism |
| *Klebsiella oxytoca* | *Shigella sonnei* | Flight 1 | 2 | 9.403 | 8.331 | 0.418 | 8.988 | 0.044 | 0.956 | -0.956 | 0.079 | -0.956 | 0.000 | Amensalism |
| *Klebsiella pneumoniae* | *Klebsiella quasipneumoniae strain IF1SW-B2* | Flight 1 | 1 | 9.634 | 9.634 | 0.000 | 9.634 | 0.000 | 1.000 | -1.000 | 0.000 | -1.000 | 0.000 | Amensalism |
| *Klebsiella pneumoniae* | *Klebsiella quasipneumoniae strain IF1SW-P3* | Flight 1 | 1 | 9.634 | 9.634 | 0.000 | 9.634 | 0.000 | 1.000 | -1.000 | 0.000 | -1.000 | 0.000 | Amensalism |
| *Klebsiella pneumoniae* | *Klebsiella quasipneumoniae strain IF1SW-P4* | Flight 1 | 1 | 9.634 | 9.634 | 0.000 | 9.634 | 0.000 | 1.000 | -1.000 | 0.000 | -1.000 | 0.000 | Amensalism |
| *Klebsiella pneumoniae* | *Klebsiella quasipneumoniae strain IF2SW-B3* | Flight 1 | 2 | 9.251 | 9.251 | 0.022 | 9.228 | 0.002 | 0.998 | -0.998 | -0.002 | -0.998 | 0.000 | Amensalism |
| *Klebsiella pneumoniae* | *Klebsiella quasipneumoniae strain IF2SW-P1* | Flight 1 | 2 | 9.251 | 9.251 | 0.030 | 9.220 | 0.003 | 0.997 | -0.997 | -0.003 | -0.997 | 0.000 | Amensalism |
| *Klebsiella pneumoniae* | *Klebsiella quasipneumoniae strain IIIF3SW-P1* | Flight 3 | 3 | 7.662 | 7.662 | 0.046 | 7.616 | 0.006 | 0.994 | -0.994 | -0.006 | -0.994 | 0.000 | Amensalism |
| *Klebsiella pneumoniae* | *Klebsiella sp. strain MS 92-3* | Flight 3 | 3 | 7.662 | 7.756 | 7.769 | 0.078 | 0.990 | 0.010 | 0.014 | -0.990 | 0.000 | -0.990 | Amensalism |
| *Klebsiella pneumoniae* | *Klebsiella variicola* | Flight 3 | 3 | 7.662 | 7.668 | 0.708 | 6.960 | 0.092 | 0.908 | -0.908 | -0.092 | -0.908 | 0.000 | Amensalism |
| *Klebsiella pneumoniae* | *Paenibacillus polymyxa strain IIF5SW-B3* | Flight 2 | 5 | 6.981 | 0.765 | 6.765 | 1.852 | 0.785 | 0.215 | -0.031 | 1.421 | 0.000 | 1.421 | Commensals |
| *Klebsiella pneumoniae* | *Paenibacillus polymyxa strain IIF5SW-B4* | Flight 2 | 5 | 6.981 | 0.765 | 6.765 | 1.852 | 0.785 | 0.215 | -0.031 | 1.421 | 0.000 | 1.421 | Commensals |
| *Klebsiella pneumoniae* | *Pantoea agglomerans* | Flight 1 | 5 | 9.251 | 5.233 | 8.417 | 2.019 | 0.807 | 0.193 | -0.090 | -0.614 | 0.000 | -0.614 | Amensalism |
| *Klebsiella pneumoniae* | *Pantoea agglomerans* | Flight 2 | 5 | 6.981 | 3.310 | 6.637 | 1.165 | 0.851 | 0.149 | -0.049 | -0.648 | 0.000 | -0.648 | Amensalism |
| *Klebsiella pneumoniae* | *Pantoea agglomerans* | Flight 3 | 1 | 9.051 | 4.640 | 7.916 | 1.933 | 0.804 | 0.196 | -0.125 | -0.583 | -0.125 | -0.583 | Competitive |
| *Klebsiella pneumoniae* | *Pantoea agglomerans* | Flight 3 | 5 | 6.623 | 2.777 | 6.333 | 1.109 | 0.851 | 0.149 | -0.044 | -0.601 | 0.000 | -0.601 | Amensalism |
| *Klebsiella pneumoniae* | *Pantoea agglomerans* | Flight 3 | 7 | 6.623 | 2.777 | 6.333 | 1.109 | 0.851 | 0.149 | -0.044 | -0.601 | 0.000 | -0.601 | Amensalism |
| *Klebsiella pneumoniae* | *Pantoea agglomerans* | Flight 3 | 8 | 6.623 | 2.777 | 6.333 | 1.109 | 0.851 | 0.149 | -0.044 | -0.601 | 0.000 | -0.601 | Amensalism |
| *Klebsiella pneumoniae* | *Pantoea ananatis* | Flight 1 | 2 | 9.251 | 7.840 | 9.107 | 10.150 | 0.473 | 0.527 | -0.015 | 0.295 | 0.000 | 0.295 | Commensals |
| *Klebsiella pneumoniae* | *Pantoea ananatis* | Flight 1 | 5 | 9.251 | 7.840 | 9.107 | 10.150 | 0.473 | 0.527 | -0.015 | 0.295 | 0.000 | 0.295 | Commensals |
| *Klebsiella pneumoniae* | *Pantoea ananatis* | Flight 2 | 5 | 6.981 | 6.254 | 7.614 | 8.552 | 0.471 | 0.529 | 0.091 | 0.367 | 0.000 | 0.367 | Commensals |
| *Klebsiella pneumoniae* | *Pantoea ananatis* | Flight 3 | 1 | 9.051 | 7.622 | 8.992 | 10.023 | 0.473 | 0.527 | -0.007 | 0.315 | 0.000 | 0.315 | Commensals |
| *Klebsiella pneumoniae* | *Pantoea ananatis* | Flight 3 | 4 | 6.623 | 5.534 | 6.933 | 7.790 | 0.471 | 0.529 | 0.047 | 0.408 | 0.000 | 0.408 | Commensals |
| *Klebsiella pneumoniae* | *Pantoea ananatis* | Flight 3 | 5 | 6.623 | 5.534 | 6.933 | 7.790 | 0.471 | 0.529 | 0.047 | 0.408 | 0.000 | 0.408 | Commensals |
| *Klebsiella pneumoniae* | *Pantoea ananatis* | Flight 3 | 7 | 6.623 | 5.534 | 6.933 | 7.790 | 0.471 | 0.529 | 0.047 | 0.408 | 0.000 | 0.408 | Commensals |
| *Klebsiella pneumoniae* | *Pantoea ananatis* | Flight 3 | 8 | 6.623 | 5.534 | 6.933 | 7.790 | 0.471 | 0.529 | 0.047 | 0.408 | 0.000 | 0.408 | Commensals |
| *Klebsiella pneumoniae* | *Pantoea conspicua* | Flight 1 | 5 | 9.251 | 4.632 | 8.416 | 2.020 | 0.806 | 0.194 | -0.090 | -0.564 | 0.000 | -0.564 | Amensalism |
| *Klebsiella pneumoniae* | *Pantoea conspicua* | Flight 2 | 5 | 6.981 | 3.267 | 6.637 | 1.165 | 0.851 | 0.149 | -0.049 | -0.643 | 0.000 | -0.643 | Amensalism |
| *Klebsiella pneumoniae* | *Pantoea conspicua* | Flight 3 | 1 | 9.051 | 4.408 | 7.915 | 1.934 | 0.804 | 0.196 | -0.125 | -0.561 | -0.125 | -0.561 | Competitive |
| *Klebsiella pneumoniae* | *Pantoea conspicua* | Flight 3 | 5 | 6.623 | 2.768 | 6.333 | 1.109 | 0.851 | 0.149 | -0.044 | -0.599 | 0.000 | -0.599 | Amensalism |
| *Klebsiella pneumoniae* | *Pantoea conspicua* | Flight 3 | 7 | 6.623 | 2.768 | 6.333 | 1.109 | 0.851 | 0.149 | -0.044 | -0.599 | 0.000 | -0.599 | Amensalism |
| *Klebsiella pneumoniae* | *Pantoea conspicua* | Flight 3 | 8 | 6.623 | 2.768 | 6.333 | 1.109 | 0.851 | 0.149 | -0.044 | -0.599 | 0.000 | -0.599 | Amensalism |
| *Klebsiella pneumoniae* | *Pantoea dispersa* | Flight 3 | 4 | 6.623 | 1.869 | 5.887 | 1.060 | 0.847 | 0.153 | -0.111 | -0.433 | -0.111 | -0.433 | Competitive |
| *Klebsiella pneumoniae* | *Pantoea sp. strain 3.5.1* | Flight 1 | 5 | 9.251 | 5.233 | 8.417 | 2.019 | 0.807 | 0.193 | -0.090 | -0.614 | 0.000 | -0.614 | Amensalism |
| *Klebsiella pneumoniae* | *Pantoea sp. strain 3.5.1* | Flight 2 | 5 | 6.981 | 3.310 | 6.637 | 1.165 | 0.851 | 0.149 | -0.049 | -0.648 | 0.000 | -0.648 | Amensalism |
| *Klebsiella pneumoniae* | *Pantoea sp. strain A4* | Flight 3 | 1 | 9.051 | 4.256 | 7.819 | 1.966 | 0.799 | 0.201 | -0.136 | -0.538 | -0.136 | -0.538 | Competitive |
| *Klebsiella pneumoniae* | *Pantoea sp. strain A4* | Flight 3 | 5 | 6.623 | 2.593 | 6.258 | 1.017 | 0.860 | 0.140 | -0.055 | -0.608 | 0.000 | -0.608 | Amensalism |
| *Klebsiella pneumoniae* | *Pantoea sp. strain A4* | Flight 3 | 7 | 6.623 | 2.593 | 6.258 | 1.017 | 0.860 | 0.140 | -0.055 | -0.608 | 0.000 | -0.608 | Amensalism |
| *Klebsiella pneumoniae* | *Pantoea sp. strain A4* | Flight 3 | 8 | 6.623 | 2.593 | 6.258 | 1.017 | 0.860 | 0.140 | -0.055 | -0.608 | 0.000 | -0.608 | Amensalism |
| *Klebsiella pneumoniae* | *Pantoea sp. strain At-9b* | Flight 3 | 1 | 9.051 | 3.804 | 9.295 | 0.188 | 0.980 | 0.020 | 0.027 | -0.951 | 0.000 | -0.951 | Amensalism |
| *Klebsiella pneumoniae* | *Pantoea sp. strain At-9b* | Flight 3 | 5 | 6.623 | 2.173 | 7.128 | 0.172 | 0.976 | 0.024 | 0.076 | -0.921 | 0.000 | -0.921 | Amensalism |
| *Klebsiella pneumoniae* | *Pantoea sp. strain At-9b* | Flight 3 | 8 | 6.623 | 2.173 | 7.128 | 0.172 | 0.976 | 0.024 | 0.076 | -0.921 | 0.000 | -0.921 | Amensalism |
| *Klebsiella pneumoniae* | *Pantoea sp. strain FF5* | Flight 3 | 1 | 9.051 | 4.640 | 7.916 | 1.933 | 0.804 | 0.196 | -0.125 | -0.583 | -0.125 | -0.583 | Competitive |
| *Klebsiella pneumoniae* | *Pantoea sp. strain FF5* | Flight 3 | 5 | 6.623 | 2.777 | 6.333 | 1.109 | 0.851 | 0.149 | -0.044 | -0.601 | 0.000 | -0.601 | Amensalism |
| *Klebsiella pneumoniae* | *Pantoea sp. strain FF5* | Flight 3 | 7 | 6.623 | 2.777 | 6.333 | 1.109 | 0.851 | 0.149 | -0.044 | -0.601 | 0.000 | -0.601 | Amensalism |

| Microorganism A | Microorganism B | Flight | Location | Vbio of A | Vbio of B | Vbio of A in AB | Vbio of B in AB | Ratio of A in AB | Ratio of B in AB | Effect of AB on A | Effect of AB on B | Significant effect on A | Significant effect on B | Interaction type |
|---|---|---|---|---|---|---|---|---|---|---|---|---|---|---|
| *Klebsiella pneumoniae* | *Pantoea sp. strain FF5* | Flight 3 | 8 | 6.623 | 2.777 | 6.333 | 1.109 | 0.851 | 0.149 | -0.044 | -0.601 | 0.000 | -0.601 | Amensalism |
| *Klebsiella pneumoniae* | *Pantoea sp. strain IMH* | Flight 3 | 1 | 9.051 | 4.409 | 7.659 | 1.939 | 0.798 | 0.202 | -0.154 | -0.560 | -0.154 | -0.560 | Competitive |
| *Klebsiella pneumoniae* | *Pantoea sp. strain IMH* | Flight 3 | 5 | 6.623 | 2.634 | 5.887 | 1.059 | 0.847 | 0.153 | -0.111 | -0.598 | -0.111 | -0.598 | Competitive |
| *Klebsiella pneumoniae* | *Pantoea sp. strain IMH* | Flight 3 | 7 | 6.623 | 2.634 | 5.887 | 1.059 | 0.847 | 0.153 | -0.111 | -0.598 | -0.111 | -0.598 | Competitive |
| *Klebsiella pneumoniae* | *Pantoea sp. strain IMH* | Flight 3 | 8 | 6.623 | 2.634 | 5.887 | 1.059 | 0.847 | 0.153 | -0.111 | -0.598 | -0.111 | -0.598 | Competitive |
| *Klebsiella pneumoniae* | *Pantoea sp. strain NGS-ED-1003* | Flight 3 | 1 | 9.051 | 4.640 | 7.916 | 1.933 | 0.804 | 0.196 | -0.125 | -0.583 | -0.125 | -0.583 | Competitive |
| *Klebsiella pneumoniae* | *Pantoea sp. strain NGS-ED-1003* | Flight 3 | 5 | 6.623 | 2.777 | 6.333 | 1.109 | 0.851 | 0.149 | -0.044 | -0.601 | 0.000 | -0.601 | Amensalism |
| *Klebsiella pneumoniae* | *Pantoea sp. strain NGS-ED-1003* | Flight 3 | 7 | 6.623 | 2.777 | 6.333 | 1.109 | 0.851 | 0.149 | -0.044 | -0.601 | 0.000 | -0.601 | Amensalism |
| *Klebsiella pneumoniae* | *Pantoea sp. strain NGS-ED-1003* | Flight 3 | 8 | 6.623 | 2.777 | 6.333 | 1.109 | 0.851 | 0.149 | -0.044 | -0.601 | 0.000 | -0.601 | Amensalism |
| *Klebsiella pneumoniae* | *Pantoea sp. strain OXWO6B1* | Flight 3 | 1 | 9.051 | 4.642 | 7.917 | 1.933 | 0.804 | 0.196 | -0.125 | -0.584 | -0.125 | -0.584 | Competitive |
| *Klebsiella pneumoniae* | *Pantoea sp. strain OXWO6B1* | Flight 3 | 5 | 6.623 | 2.779 | 6.333 | 1.109 | 0.851 | 0.149 | -0.044 | -0.601 | 0.000 | -0.601 | Amensalism |
| *Klebsiella pneumoniae* | *Pantoea sp. strain OXWO6B1* | Flight 3 | 7 | 6.623 | 2.779 | 6.333 | 1.109 | 0.851 | 0.149 | -0.044 | -0.601 | 0.000 | -0.601 | Amensalism |
| *Klebsiella pneumoniae* | *Pantoea sp. strain OXWO6B1* | Flight 3 | 8 | 6.623 | 2.779 | 6.333 | 1.109 | 0.851 | 0.149 | -0.044 | -0.601 | 0.000 | -0.601 | Amensalism |
| *Klebsiella pneumoniae* | *Pantoea vagans* | Flight 2 | 5 | 6.981 | 3.312 | 6.637 | 1.165 | 0.851 | 0.149 | -0.049 | -0.648 | 0.000 | -0.648 | Amensalism |
| *Klebsiella pneumoniae* | *Pantoea vagans* | Flight 3 | 1 | 9.051 | 4.642 | 7.917 | 1.933 | 0.804 | 0.196 | -0.125 | -0.584 | -0.125 | -0.584 | Competitive |
| *Klebsiella pneumoniae* | *Pantoea vagans* | Flight 3 | 5 | 6.623 | 2.778 | 6.333 | 1.109 | 0.851 | 0.149 | -0.044 | -0.601 | 0.000 | -0.601 | Amensalism |
| *Klebsiella pneumoniae* | *Pantoea vagans* | Flight 3 | 7 | 6.623 | 2.778 | 6.333 | 1.109 | 0.851 | 0.149 | -0.044 | -0.601 | 0.000 | -0.601 | Amensalism |
| *Klebsiella pneumoniae* | *Pantoea vagans* | Flight 3 | 8 | 6.623 | 2.778 | 6.333 | 1.109 | 0.851 | 0.149 | -0.044 | -0.601 | 0.000 | -0.601 | Amensalism |
| *Klebsiella pneumoniae* | *Penicillium chrysogenum* | Flight 1 | 1 | 9.634 | 0.071 | 9.634 | 0.000 | 1.000 | 0.000 | 0.000 | -1.000 | 0.000 | -1.000 | Amensalism |
| *Klebsiella pneumoniae* | *Penicillium rubens* | Flight 1 | 1 | 9.634 | 0.071 | 9.634 | 0.000 | 1.000 | 0.000 | 0.000 | -1.000 | 0.000 | -1.000 | Amensalism |
| *Klebsiella pneumoniae* | *Penicillium rubens* | Flight 1 | 2 | 9.251 | 0.049 | 9.251 | 0.000 | 1.000 | 0.000 | 0.000 | -1.000 | 0.000 | -1.000 | Amensalism |
| *Klebsiella pneumoniae* | *Penicillium rubens* | Flight 1 | 5 | 9.251 | 0.049 | 9.251 | 0.000 | 1.000 | 0.000 | 0.000 | -1.000 | 0.000 | -1.000 | Amensalism |
| *Klebsiella pneumoniae* | *Penicillium rubens* | Flight 3 | 3 | 7.662 | 0.023 | 7.662 | 0.000 | 1.000 | 0.000 | 0.000 | -1.000 | 0.000 | -1.000 | Amensalism |
| *Klebsiella pneumoniae* | *Rahnella aquatilis* | Flight 3 | 4 | 6.623 | 2.624 | 5.809 | 1.050 | 0.847 | 0.153 | -0.123 | -0.600 | -0.123 | -0.600 | Competitive |
| *Klebsiella pneumoniae* | *Rhodotorula sp. strain JG-1b* | Flight 1 | 1 | 9.634 | 0.154 | 9.897 | 0.251 | 0.975 | 0.025 | 0.027 | 0.627 | 0.000 | 0.627 | Commensals |
| *Klebsiella pneumoniae* | *Rhodotorula sp. strain JG-1b* | Flight 1 | 2 | 9.251 | 0.153 | 9.244 | 0.168 | 0.982 | 0.018 | -0.001 | 0.099 | 0.000 | 0.000 | Neutral |
| *Klebsiella pneumoniae* | *Rhodotorula sp. strain JG-1b* | Flight 1 | 5 | 9.251 | 0.153 | 9.244 | 0.168 | 0.982 | 0.018 | -0.001 | 0.099 | 0.000 | 0.000 | Neutral |
| *Klebsiella pneumoniae* | *Rhodotorula sp. strain JG-1b* | Flight 3 | 1 | 9.051 | 0.154 | 9.051 | 0.000 | 1.000 | 0.000 | 0.000 | -1.000 | 0.000 | -1.000 | Amensalism |
| *Klebsiella pneumoniae* | *Rhodotorula toruloides* | Flight 1 | 1 | 9.634 | 2.346 | 9.262 | 4.160 | 0.690 | 0.310 | -0.039 | 0.773 | 0.000 | 0.773 | Commensals |
| *Klebsiella pneumoniae* | *Salmonella enterica* | Flight 1 | 2 | 9.251 | 8.478 | 0.172 | 9.081 | 0.019 | 0.981 | -0.981 | 0.071 | -0.981 | 0.000 | Amensalism |
| *Klebsiella pneumoniae* | *Salmonella enterica* | Flight 3 | 1 | 9.051 | 8.372 | 0.091 | 8.964 | 0.010 | 0.990 | -0.990 | 0.071 | -0.990 | 0.000 | Amensalism |
| *Klebsiella pneumoniae* | *Salmonella enterica* | Flight 3 | 3 | 7.662 | 6.819 | 0.144 | 7.520 | 0.019 | 0.981 | -0.981 | 0.103 | -0.981 | 0.103 | Parasitism |
| *Klebsiella pneumoniae* | *Salmonella enterica* | Flight 3 | 5 | 6.623 | 5.907 | 0.126 | 6.527 | 0.019 | 0.981 | -0.981 | 0.105 | -0.981 | 0.105 | Parasitism |
| *Klebsiella pneumoniae* | *Salmonella enterica* | Flight 3 | 7 | 6.623 | 5.907 | 0.126 | 6.527 | 0.019 | 0.981 | -0.981 | 0.105 | -0.981 | 0.105 | Parasitism |
| *Klebsiella pneumoniae* | *Salmonella enterica* | Flight 3 | 8 | 6.623 | 5.907 | 0.126 | 6.527 | 0.019 | 0.981 | -0.981 | 0.105 | -0.981 | 0.105 | Parasitism |
| *Klebsiella pneumoniae* | *Shigella sonnei* | Flight 1 | 2 | 9.251 | 8.331 | 0.418 | 8.988 | 0.044 | 0.956 | -0.955 | 0.079 | -0.955 | 0.000 | Amensalism |
| *Klebsiella pneumoniae* | *Staphylococcus saprophyticus* | Flight 3 | 3 | 7.662 | 1.481 | 9.228 | 2.942 | 0.758 | 0.242 | 0.204 | 0.986 | 0.204 | 0.986 | Mutualism |
| *Klebsiella pneumoniae strain F3-2P(2*)* | *Paenibacillus polymyxa* | Flight 3 | 2 | 12.151 | 1.781 | 10.147 | 3.528 | 0.742 | 0.258 | -0.165 | 0.981 | -0.165 | 0.981 | Parasitism |
| *Klebsiella pneumoniae strain F3-2P(2*)* | *Penicillium chrysogenum* | Flight 3 | 2 | 12.151 | 0.049 | 12.151 | 0.000 | 1.000 | 0.000 | 0.000 | -1.000 | 0.000 | -1.000 | Amensalism |
| *Klebsiella pneumoniae strain F3-2P(2*)* | *Penicillium flavigenum* | Flight 3 | 2 | 12.151 | 0.097 | 12.151 | 0.000 | 1.000 | 0.000 | 0.000 | -1.000 | 0.000 | -1.000 | Amensalism |
| *Klebsiella pneumoniae strain F3-2P(2*)* | *Penicillium nalgiovense* | Flight 3 | 2 | 12.151 | 0.067 | 12.151 | 0.000 | 1.000 | 0.000 | 0.000 | -1.000 | 0.000 | -1.000 | Amensalism |
| *Klebsiella pneumoniae strain F3-2P(2*)* | *Penicillium rubens* | Flight 3 | 2 | 12.151 | 0.049 | 12.151 | 0.000 | 1.000 | 0.000 | 0.000 | -1.000 | 0.000 | -1.000 | Amensalism |
| *Klebsiella pneumoniae strain F3-2P(2*)* | *Rhodotorula sp. strain JG-1b* | Flight 3 | 2 | 12.151 | 0.156 | 11.922 | 0.301 | 0.975 | 0.025 | -0.019 | 0.925 | 0.000 | 0.925 | Commensals |
| *Klebsiella pneumoniae strain F3-2P(2*)* | *Staphylococcus aureus* | Flight 3 | 2 | 12.151 | 2.739 | 13.549 | 4.789 | 0.739 | 0.261 | 0.115 | 0.748 | 0.115 | 0.748 | Mutualism |
| *Klebsiella pneumoniae strain F3-2P(2*)* | *Staphylococcus epidermidis* | Flight 3 | 2 | 12.151 | 2.313 | 13.474 | 4.700 | 0.741 | 0.259 | 0.109 | 1.032 | 0.109 | 1.032 | Mutualism |
| *Klebsiella pneumoniae strain F3-2P(2*)* | *Staphylococcus haemolyticus* | Flight 3 | 2 | 12.151 | 2.519 | 13.679 | 4.780 | 0.741 | 0.259 | 0.126 | 0.898 | 0.126 | 0.898 | Mutualism |
| *Klebsiella pneumoniae strain F3-2P(2*)* | *Staphylococcus saprophyticus* | Flight 3 | 2 | 12.151 | 2.363 | 13.248 | 4.715 | 0.738 | 0.262 | 0.090 | 0.995 | 0.000 | 0.995 | Commensals |
| *Klebsiella pneumoniae strain F3-2P(2*)* | *Staphylococcus sp. strain LCT-H4* | Flight 3 | 2 | 12.151 | 2.388 | 13.459 | 4.562 | 0.747 | 0.253 | 0.108 | 0.910 | 0.108 | 0.910 | Mutualism |
| *Klebsiella pneumoniae strain F3-2P(2*)* | *Staphylococcus warneri* | Flight 3 | 2 | 12.151 | 2.543 | 16.439 | 4.820 | 0.773 | 0.227 | 0.353 | 0.896 | 0.353 | 0.896 | Mutualism |
| *Klebsiella quasipneumoniae strain IF1SW-B2* | *Klebsiella quasipneumoniae strain IF1SW-P3* | Flight 1 | 1 | 9.634 | 9.634 | 4.817 | 4.817 | 0.500 | 0.500 | -0.500 | -0.500 | -0.500 | -0.500 | Competitive |
| *Klebsiella quasipneumoniae strain IF1SW-B2* | *Klebsiella quasipneumoniae strain IF1SW-P4* | Flight 1 | 1 | 9.634 | 9.634 | 4.817 | 4.817 | 0.500 | 0.500 | -0.500 | -0.500 | -0.500 | -0.500 | Competitive |
| *Klebsiella quasipneumoniae strain IF1SW-B2* | *Penicillium chrysogenum* | Flight 1 | 1 | 9.634 | 0.071 | 9.634 | 0.000 | 1.000 | 0.000 | 0.000 | -1.000 | 0.000 | -1.000 | Amensalism |
| *Klebsiella quasipneumoniae strain IF1SW-B2* | *Penicillium rubens* | Flight 1 | 1 | 9.634 | 0.071 | 9.634 | 0.000 | 1.000 | 0.000 | 0.000 | -1.000 | 0.000 | -1.000 | Amensalism |
| *Klebsiella quasipneumoniae strain IF1SW-B2* | *Rhodotorula sp. strain JG-1b* | Flight 1 | 1 | 9.634 | 0.154 | 9.897 | 0.251 | 0.975 | 0.025 | 0.027 | 0.627 | 0.000 | 0.627 | Commensals |
| *Klebsiella quasipneumoniae strain IF1SW-B2* | *Rhodotorula toruloides* | Flight 1 | 1 | 9.634 | 2.346 | 9.262 | 4.160 | 0.690 | 0.310 | -0.039 | 0.773 | 0.000 | 0.773 | Commensals |
| *Klebsiella quasipneumoniae strain IF1SW-P3* | *Klebsiella quasipneumoniae strain IF1SW-P4* | Flight 1 | 1 | 9.634 | 9.634 | 4.817 | 4.817 | 0.500 | 0.500 | -0.500 | -0.500 | -0.500 | -0.500 | Competitive |
| *Klebsiella quasipneumoniae strain IF1SW-P3* | *Penicillium chrysogenum* | Flight 1 | 1 | 9.634 | 0.071 | 9.634 | 0.000 | 1.000 | 0.000 | 0.000 | -1.000 | 0.000 | -1.000 | Amensalism |
| *Klebsiella quasipneumoniae strain IF1SW-P3* | *Penicillium rubens* | Flight 1 | 1 | 9.634 | 0.071 | 9.634 | 0.000 | 1.000 | 0.000 | 0.000 | -1.000 | 0.000 | -1.000 | Amensalism |
| *Klebsiella quasipneumoniae strain IF1SW-P3* | *Rhodotorula sp. strain JG-1b* | Flight 1 | 1 | 9.634 | 0.154 | 9.897 | 0.251 | 0.975 | 0.025 | 0.027 | 0.627 | 0.000 | 0.627 | Commensals |
| *Klebsiella quasipneumoniae strain IF1SW-P3* | *Rhodotorula toruloides* | Flight 1 | 1 | 9.634 | 2.346 | 9.262 | 4.160 | 0.690 | 0.310 | -0.039 | 0.773 | 0.000 | 0.773 | Commensals |
| *Klebsiella quasipneumoniae strain IF1SW-P4* | *Penicillium chrysogenum* | Flight 1 | 1 | 9.634 | 0.071 | 9.634 | 0.000 | 1.000 | 0.000 | 0.000 | -1.000 | 0.000 | -1.000 | Amensalism |
| *Klebsiella quasipneumoniae strain IF1SW-P4* | *Penicillium rubens* | Flight 1 | 1 | 9.634 | 0.071 | 9.634 | 0.000 | 1.000 | 0.000 | 0.000 | -1.000 | 0.000 | -1.000 | Amensalism |
| *Klebsiella quasipneumoniae strain IF1SW-P4* | *Rhodotorula sp. strain JG-1b* | Flight 1 | 1 | 9.634 | 0.154 | 9.897 | 0.251 | 0.975 | 0.025 | 0.027 | 0.627 | 0.000 | 0.627 | Commensals |
| *Klebsiella quasipneumoniae strain IF1SW-P4* | *Rhodotorula toruloides* | Flight 1 | 1 | 9.634 | 2.346 | 9.262 | 4.160 | 0.690 | 0.310 | -0.039 | 0.773 | 0.000 | 0.773 | Commensals |
| *Klebsiella quasipneumoniae strain IF2SW-B3* | *Klebsiella quasipneumoniae strain IF2SW-P1* | Flight 1 | 2 | 9.251 | 9.251 | 4.672 | 4.579 | 0.505 | 0.495 | -0.495 | -0.505 | -0.495 | -0.505 | Competitive |
| *Klebsiella quasipneumoniae strain IF2SW-B3* | *Pantoea ananatis* | Flight 1 | 2 | 9.251 | 7.840 | 9.108 | 10.150 | 0.473 | 0.527 | -0.015 | 0.295 | 0.000 | 0.295 | Commensals |

| Microorganism A | Microorganism B | Flight | Location | Vbio of A | Vbio of B | Vbio of A in AB | Vbio of B in AB | Ratio of A in AB | Ratio of B in AB | Effect of AB on A | Effect of AB on B | Significant effect on A | Significant effect on B | Interaction type |
|---|---|---|---|---|---|---|---|---|---|---|---|---|---|---|
| *Klebsiella quasipneumoniae strain IF2SW-B3* | *Penicillium rubens* | Flight 1 | 2 | 9.251 | 0.049 | 9.251 | 0.000 | 1.000 | 0.000 | 0.000 | -1.000 | 0.000 | -1.000 | Amensalism |
| *Klebsiella quasipneumoniae strain IF2SW-B3* | *Rhodotorula sp. strain JG-1b* | Flight 1 | 2 | 9.251 | 0.153 | 9.244 | 0.168 | 0.982 | 0.018 | -0.001 | 0.099 | 0.000 | 0.000 | Neutral |
| *Klebsiella quasipneumoniae strain IF2SW-B3* | *Salmonella enterica* | Flight 1 | 2 | 9.251 | 8.478 | 0.172 | 9.081 | 0.019 | 0.981 | -0.981 | 0.071 | -0.981 | 0.000 | Amensalism |
| *Klebsiella quasipneumoniae strain IF2SW-B3* | *Shigella sonnei* | Flight 1 | 2 | 9.251 | 8.331 | 0.418 | 8.988 | 0.044 | 0.956 | -0.955 | 0.079 | -0.955 | 0.000 | Amensalism |
| *Klebsiella quasipneumoniae strain IF2SW-P1* | *Pantoea ananatis* | Flight 1 | 2 | 9.251 | 7.840 | 9.108 | 10.150 | 0.473 | 0.527 | -0.015 | 0.295 | 0.000 | 0.295 | Commensals |
| *Klebsiella quasipneumoniae strain IF2SW-P1* | *Penicillium rubens* | Flight 1 | 2 | 9.251 | 0.049 | 9.251 | 0.000 | 1.000 | 0.000 | 0.000 | -1.000 | 0.000 | -1.000 | Amensalism |
| *Klebsiella quasipneumoniae strain IF2SW-P1* | *Rhodotorula sp. strain JG-1b* | Flight 1 | 2 | 9.251 | 0.153 | 9.244 | 0.168 | 0.982 | 0.018 | -0.001 | 0.099 | 0.000 | 0.000 | Neutral |
| *Klebsiella quasipneumoniae strain IF2SW-P1* | *Salmonella enterica* | Flight 1 | 2 | 9.251 | 8.478 | 0.172 | 9.081 | 0.019 | 0.981 | -0.981 | 0.071 | -0.981 | 0.000 | Amensalism |
| *Klebsiella quasipneumoniae strain IF2SW-P1* | *Shigella sonnei* | Flight 1 | 2 | 9.251 | 8.331 | 0.418 | 8.988 | 0.044 | 0.956 | -0.955 | 0.079 | -0.955 | 0.000 | Amensalism |
| *Klebsiella quasipneumoniae strain IIIF3SW-P1* | *Klebsiella sp. strain MS 92-3* | Flight 3 | 3 | 7.662 | 7.756 | 7.769 | 0.078 | 0.990 | 0.010 | 0.014 | -0.990 | 0.000 | -0.990 | Amensalism |
| *Klebsiella quasipneumoniae strain IIIF3SW-P1* | *Klebsiella variicola* | Flight 3 | 3 | 7.662 | 7.668 | 7.641 | 0.027 | 0.996 | 0.004 | -0.003 | -0.996 | 0.000 | -0.996 | Amensalism |
| *Klebsiella quasipneumoniae strain IIIF3SW-P1* | *Penicillium rubens* | Flight 3 | 3 | 7.662 | 0.023 | 7.662 | 0.000 | 1.000 | 0.000 | 0.000 | -1.000 | 0.000 | -1.000 | Amensalism |
| *Klebsiella quasipneumoniae strain IIIF3SW-P1* | *Salmonella enterica* | Flight 3 | 3 | 7.662 | 6.819 | 0.144 | 7.520 | 0.019 | 0.981 | -0.981 | 0.103 | -0.981 | 0.103 | Parasitism |
| *Klebsiella quasipneumoniae strain IIIF3SW-P1* | *Staphylococcus saprophyticus* | Flight 3 | 3 | 7.662 | 1.481 | 9.228 | 2.942 | 0.758 | 0.242 | 0.204 | 0.986 | 0.204 | 0.986 | Mutualism |
| *Klebsiella sp. strain MS 92-3* | *Klebsiella variicola* | Flight 3 | 3 | 7.756 | 7.668 | 0.078 | 7.769 | 0.010 | 0.990 | -0.990 | 0.013 | -0.990 | 0.000 | Amensalism |
| *Klebsiella sp. strain MS 92-3* | *Penicillium rubens* | Flight 3 | 3 | 7.756 | 0.023 | 7.756 | 0.000 | 1.000 | 0.000 | 0.000 | -1.000 | 0.000 | -1.000 | Amensalism |
| *Klebsiella sp. strain MS 92-3* | *Salmonella enterica* | Flight 3 | 3 | 7.756 | 6.819 | 0.148 | 7.716 | 0.019 | 0.981 | -0.981 | 0.131 | -0.981 | 0.131 | Parasitism |
| *Klebsiella sp. strain MS 92-3* | *Staphylococcus saprophyticus* | Flight 3 | 3 | 7.756 | 1.481 | 9.353 | 2.928 | 0.762 | 0.238 | 0.206 | 0.976 | 0.206 | 0.976 | Mutualism |
| *Klebsiella variicola* | *Penicillium rubens* | Flight 3 | 3 | 7.668 | 0.023 | 7.668 | 0.000 | 1.000 | 0.000 | 0.000 | -1.000 | 0.000 | -1.000 | Amensalism |
| *Klebsiella variicola* | *Salmonella enterica* | Flight 3 | 3 | 7.668 | 6.819 | 0.145 | 7.540 | 0.019 | 0.981 | -0.981 | 0.106 | -0.981 | 0.106 | Parasitism |
| *Klebsiella variicola* | *Staphylococcus saprophyticus* | Flight 3 | 3 | 7.668 | 1.481 | 9.228 | 2.942 | 0.758 | 0.242 | 0.203 | 0.986 | 0.203 | 0.986 | Mutualism |
| *Paenibacillus polymyxa* | *Penicillium chrysogenum* | Flight 3 | 2 | 1.781 | 0.049 | 2.053 | 0.034 | 0.984 | 0.016 | 0.153 | -0.312 | 0.153 | -0.312 | Parasitism |
| *Paenibacillus polymyxa* | *Penicillium flavigenum* | Flight 3 | 2 | 1.781 | 0.097 | 2.021 | 0.020 | 0.990 | 0.010 | 0.135 | -0.789 | 0.135 | -0.789 | Parasitism |
| *Paenibacillus polymyxa* | *Penicillium nalgiovense* | Flight 3 | 2 | 1.781 | 0.067 | 1.965 | 0.020 | 0.990 | 0.010 | 0.103 | -0.704 | 0.103 | -0.704 | Parasitism |
| *Paenibacillus polymyxa* | *Penicillium rubens* | Flight 3 | 2 | 1.781 | 0.049 | 2.053 | 0.034 | 0.984 | 0.016 | 0.153 | -0.312 | 0.153 | -0.312 | Parasitism |
| *Paenibacillus polymyxa* | *Rhodotorula sp. strain JG-1b* | Flight 3 | 2 | 1.781 | 0.156 | 1.260 | 1.392 | 0.475 | 0.525 | -0.292 | 7.909 | -0.292 | 7.909 | Parasitism |
| *Paenibacillus polymyxa* | *Staphylococcus aureus* | Flight 3 | 2 | 1.781 | 2.739 | 0.039 | 3.094 | 0.012 | 0.988 | -0.978 | 0.129 | -0.978 | 0.129 | Parasitism |
| *Paenibacillus polymyxa* | *Staphylococcus epidermidis* | Flight 3 | 2 | 1.781 | 2.313 | 0.090 | 2.947 | 0.030 | 0.970 | -0.949 | 0.274 | -0.949 | 0.274 | Parasitism |
| *Paenibacillus polymyxa* | *Staphylococcus haemolyticus* | Flight 3 | 2 | 1.781 | 2.519 | 0.258 | 2.894 | 0.082 | 0.918 | -0.855 | 0.149 | -0.855 | 0.149 | Parasitism |
| *Paenibacillus polymyxa* | *Staphylococcus saprophyticus* | Flight 3 | 2 | 1.781 | 2.363 | 0.036 | 2.788 | 0.013 | 0.987 | -0.980 | 0.180 | -0.980 | 0.180 | Parasitism |
| *Paenibacillus polymyxa* | *Staphylococcus sp. strain LCT-H4* | Flight 3 | 2 | 1.781 | 2.388 | 0.028 | 2.796 | 0.010 | 0.990 | -0.984 | 0.171 | -0.984 | 0.171 | Parasitism |
| *Paenibacillus polymyxa* | *Staphylococcus warneri* | Flight 3 | 2 | 1.781 | 2.543 | 0.094 | 3.058 | 0.030 | 0.970 | -0.947 | 0.203 | -0.947 | 0.203 | Parasitism |
| *Paenibacillus polymyxa strain IIF5SW-B3* | *Paenibacillus polymyxa strain IIF5SW-B4* | Flight 2 | 5 | 0.765 | 0.765 | 0.371 | 0.394 | 0.485 | 0.515 | -0.515 | -0.485 | -0.515 | -0.485 | Competitive |
| *Paenibacillus polymyxa strain IIF5SW-B3* | *Pantoea agglomerans* | Flight 2 | 5 | 0.765 | 3.310 | 0.959 | 3.187 | 0.231 | 0.769 | 0.253 | -0.037 | 0.253 | 0.000 | Commensals |
| *Paenibacillus polymyxa strain IIF5SW-B3* | *Pantoea ananatis* | Flight 2 | 5 | 0.765 | 6.254 | 1.664 | 6.846 | 0.196 | 0.804 | 1.175 | 0.095 | 1.175 | 0.000 | Commensals |
| *Paenibacillus polymyxa strain IIF5SW-B3* | *Pantoea conspicua* | Flight 2 | 5 | 0.765 | 3.267 | 0.797 | 3.280 | 0.196 | 0.804 | 0.042 | 0.004 | 0.000 | 0.000 | Neutral |
| *Paenibacillus polymyxa strain IIF5SW-B3* | *Pantoea sp. strain 3.5.1* | Flight 2 | 5 | 0.765 | 3.310 | 0.959 | 3.187 | 0.231 | 0.769 | 0.253 | -0.037 | 0.253 | 0.000 | Commensals |
| *Paenibacillus polymyxa strain IIF5SW-B3* | *Pantoea vagans* | Flight 2 | 5 | 0.765 | 3.312 | 0.959 | 3.189 | 0.231 | 0.769 | 0.253 | -0.037 | 0.253 | 0.000 | Commensals |
| *Paenibacillus polymyxa strain IIF5SW-B4* | *Pantoea agglomerans* | Flight 2 | 5 | 0.765 | 3.310 | 0.959 | 3.187 | 0.231 | 0.769 | 0.253 | -0.037 | 0.253 | 0.000 | Commensals |
| *Paenibacillus polymyxa strain IIF5SW-B4* | *Pantoea ananatis* | Flight 2 | 5 | 0.765 | 6.254 | 1.664 | 6.846 | 0.196 | 0.804 | 1.175 | 0.095 | 1.175 | 0.000 | Commensals |
| *Paenibacillus polymyxa strain IIF5SW-B4* | *Pantoea conspicua* | Flight 2 | 5 | 0.765 | 3.267 | 0.797 | 3.280 | 0.196 | 0.804 | 0.042 | 0.004 | 0.000 | 0.000 | Neutral |
| *Paenibacillus polymyxa strain IIF5SW-B4* | *Pantoea sp. strain 3.5.1* | Flight 2 | 5 | 0.765 | 3.310 | 0.959 | 3.187 | 0.231 | 0.769 | 0.253 | -0.037 | 0.253 | 0.000 | Commensals |
| *Paenibacillus polymyxa strain IIF5SW-B4* | *Pantoea vagans* | Flight 2 | 5 | 0.765 | 3.312 | 0.959 | 3.189 | 0.231 | 0.769 | 0.253 | -0.037 | 0.253 | 0.000 | Commensals |
| *Pantoea agglomerans* | *Pantoea ananatis* | Flight 1 | 5 | 5.233 | 7.840 | 0.657 | 8.712 | 0.070 | 0.930 | -0.874 | 0.111 | -0.874 | 0.111 | Parasitism |
| *Pantoea agglomerans* | *Pantoea ananatis* | Flight 2 | 5 | 3.310 | 6.254 | 0.187 | 7.234 | 0.025 | 0.975 | -0.943 | 0.157 | -0.943 | 0.157 | Parasitism |
| *Pantoea agglomerans* | *Pantoea ananatis* | Flight 3 | 1 | 4.640 | 7.622 | 0.485 | 8.477 | 0.054 | 0.946 | -0.895 | 0.112 | -0.895 | 0.112 | Parasitism |
| *Pantoea agglomerans* | *Pantoea ananatis* | Flight 3 | 5 | 2.777 | 5.534 | 0.299 | 6.568 | 0.044 | 0.956 | -0.892 | 0.187 | -0.892 | 0.187 | Parasitism |
| *Pantoea agglomerans* | *Pantoea ananatis* | Flight 3 | 7 | 2.777 | 5.534 | 0.299 | 6.568 | 0.044 | 0.956 | -0.892 | 0.187 | -0.892 | 0.187 | Parasitism |
| *Pantoea agglomerans* | *Pantoea ananatis* | Flight 3 | 8 | 2.777 | 5.534 | 0.299 | 6.568 | 0.044 | 0.956 | -0.892 | 0.187 | -0.892 | 0.187 | Parasitism |
| *Pantoea agglomerans* | *Pantoea conspicua* | Flight 1 | 5 | 5.233 | 4.632 | 5.131 | 0.102 | 0.981 | 0.019 | -0.019 | -0.978 | 0.000 | -0.978 | Amensalism |
| *Pantoea agglomerans* | *Pantoea conspicua* | Flight 2 | 5 | 3.310 | 3.267 | 0.025 | 3.286 | 0.008 | 0.992 | -0.992 | 0.006 | -0.992 | 0.000 | Amensalism |
| *Pantoea agglomerans* | *Pantoea conspicua* | Flight 3 | 1 | 4.640 | 4.408 | 4.640 | 0.000 | 1.000 | 0.000 | 0.000 | -1.000 | 0.000 | -1.000 | Amensalism |
| *Pantoea agglomerans* | *Pantoea conspicua* | Flight 3 | 5 | 2.777 | 2.768 | 0.001 | 2.776 | 0.001 | 0.999 | -0.999 | 0.003 | -0.999 | 0.000 | Amensalism |
| *Pantoea agglomerans* | *Pantoea conspicua* | Flight 3 | 7 | 2.777 | 2.768 | 0.001 | 2.776 | 0.001 | 0.999 | -0.999 | 0.003 | -0.999 | 0.000 | Amensalism |
| *Pantoea agglomerans* | *Pantoea conspicua* | Flight 3 | 8 | 2.777 | 2.768 | 0.001 | 2.776 | 0.001 | 0.999 | -0.999 | 0.003 | -0.999 | 0.000 | Amensalism |
| *Pantoea agglomerans* | *Pantoea sp. strain 3.5.1* | Flight 1 | 5 | 5.233 | 5.233 | 5.233 | 0.000 | 1.000 | 0.000 | 0.000 | -1.000 | 0.000 | -1.000 | Amensalism |
| *Pantoea agglomerans* | *Pantoea sp. strain 3.5.1* | Flight 2 | 5 | 3.310 | 3.310 | 3.310 | 0.000 | 1.000 | 0.000 | 0.000 | -1.000 | 0.000 | -1.000 | Amensalism |
| *Pantoea agglomerans* | *Pantoea sp. strain A4* | Flight 3 | 1 | 4.640 | 4.256 | 0.053 | 4.596 | 0.011 | 0.989 | -0.988 | 0.080 | -0.988 | 0.000 | Amensalism |
| *Pantoea agglomerans* | *Pantoea sp. strain A4* | Flight 3 | 5 | 2.777 | 2.593 | 0.033 | 2.749 | 0.012 | 0.988 | -0.988 | 0.060 | -0.988 | 0.000 | Amensalism |
| *Pantoea agglomerans* | *Pantoea sp. strain A4* | Flight 3 | 7 | 2.777 | 2.593 | 0.033 | 2.749 | 0.012 | 0.988 | -0.988 | 0.060 | -0.988 | 0.000 | Amensalism |
| *Pantoea agglomerans* | *Pantoea sp. strain A4* | Flight 3 | 8 | 2.777 | 2.593 | 0.033 | 2.749 | 0.012 | 0.988 | -0.988 | 0.060 | -0.988 | 0.000 | Amensalism |
| *Pantoea agglomerans* | *Pantoea sp. strain At-9b* | Flight 3 | 1 | 4.640 | 3.804 | 0.443 | 4.199 | 0.095 | 0.905 | -0.904 | 0.104 | -0.904 | 0.104 | Parasitism |
| *Pantoea agglomerans* | *Pantoea sp. strain At-9b* | Flight 3 | 5 | 2.777 | 2.173 | 0.156 | 2.622 | 0.056 | 0.944 | -0.944 | 0.207 | -0.944 | 0.207 | Parasitism |
| *Pantoea agglomerans* | *Pantoea sp. strain At-9b* | Flight 3 | 8 | 2.777 | 2.173 | 0.156 | 2.622 | 0.056 | 0.944 | -0.944 | 0.207 | -0.944 | 0.207 | Parasitism |
| *Pantoea agglomerans* | *Pantoea sp. strain FF5* | Flight 3 | 1 | 4.640 | 4.640 | 4.640 | 0.000 | 1.000 | 0.000 | 0.000 | -1.000 | 0.000 | -1.000 | Amensalism |
| *Pantoea agglomerans* | *Pantoea sp. strain FF5* | Flight 3 | 5 | 2.777 | 2.777 | 2.777 | 0.000 | 1.000 | 0.000 | 0.000 | -1.000 | 0.000 | -1.000 | Amensalism |

| Microorganism A | Microorganism B | Flight | Location | Vbio of A | Vbio of B | Vbio of A in AB | Vbio of B in AB | Ratio of A in AB | Ratio of B in AB | Effect of AB on A | Effect of AB on B | Significant effect on A | Significant effect on B | Interaction type |
|---|---|---|---|---|---|---|---|---|---|---|---|---|---|---|
| *Pantoea agglomerans* | *Pantoea sp. strain FF5* | Flight 3 | 7 | 2.777 | 2.777 | 2.777 | 0.000 | 1.000 | 0.000 | 0.000 | -1.000 | 0.000 | -1.000 | Amensalism |
| *Pantoea agglomerans* | *Pantoea sp. strain FF5* | Flight 3 | 8 | 2.777 | 2.777 | 2.777 | 0.000 | 1.000 | 0.000 | 0.000 | -1.000 | 0.000 | -1.000 | Amensalism |
| *Pantoea agglomerans* | *Pantoea sp. strain IMH* | Flight 3 | 1 | 4.640 | 4.409 | 0.081 | 4.584 | 0.017 | 0.983 | -0.982 | 0.040 | -0.982 | 0.000 | Amensalism |
| *Pantoea agglomerans* | *Pantoea sp. strain IMH* | Flight 3 | 5 | 2.777 | 2.634 | 0.029 | 2.791 | 0.010 | 0.990 | -0.990 | 0.059 | -0.990 | 0.000 | Amensalism |
| *Pantoea agglomerans* | *Pantoea sp. strain IMH* | Flight 3 | 7 | 2.777 | 2.634 | 0.029 | 2.791 | 0.010 | 0.990 | -0.990 | 0.059 | -0.990 | 0.000 | Amensalism |
| *Pantoea agglomerans* | *Pantoea sp. strain IMH* | Flight 3 | 8 | 2.777 | 2.634 | 0.029 | 2.791 | 0.010 | 0.990 | -0.990 | 0.059 | -0.990 | 0.000 | Amensalism |
| *Pantoea agglomerans* | *Pantoea sp. strain NGS-ED-1003* | Flight 3 | 1 | 4.640 | 4.640 | 4.640 | 0.000 | 1.000 | 0.000 | 0.000 | -1.000 | 0.000 | -1.000 | Amensalism |
| *Pantoea agglomerans* | *Pantoea sp. strain NGS-ED-1003* | Flight 3 | 5 | 2.777 | 2.777 | 2.777 | 0.000 | 1.000 | 0.000 | 0.000 | -1.000 | 0.000 | -1.000 | Amensalism |
| *Pantoea agglomerans* | *Pantoea sp. strain NGS-ED-1003* | Flight 3 | 7 | 2.777 | 2.777 | 2.777 | 0.000 | 1.000 | 0.000 | 0.000 | -1.000 | 0.000 | -1.000 | Amensalism |
| *Pantoea agglomerans* | *Pantoea sp. strain NGS-ED-1003* | Flight 3 | 8 | 2.777 | 2.777 | 2.777 | 0.000 | 1.000 | 0.000 | 0.000 | -1.000 | 0.000 | -1.000 | Amensalism |
| *Pantoea agglomerans* | *Pantoea sp. strain OXWO6B1* | Flight 3 | 1 | 4.640 | 4.642 | 0.000 | 4.642 | 0.000 | 1.000 | -1.000 | 0.000 | -1.000 | 0.000 | Amensalism |
| *Pantoea agglomerans* | *Pantoea sp. strain OXWO6B1* | Flight 3 | 5 | 2.777 | 2.779 | 0.000 | 2.779 | 0.000 | 1.000 | -1.000 | 0.000 | -1.000 | 0.000 | Amensalism |
| *Pantoea agglomerans* | *Pantoea sp. strain OXWO6B1* | Flight 3 | 7 | 2.777 | 2.779 | 0.000 | 2.779 | 0.000 | 1.000 | -1.000 | 0.000 | -1.000 | 0.000 | Amensalism |
| *Pantoea agglomerans* | *Pantoea sp. strain OXWO6B1* | Flight 3 | 8 | 2.777 | 2.779 | 0.000 | 2.779 | 0.000 | 1.000 | -1.000 | 0.000 | -1.000 | 0.000 | Amensalism |
| *Pantoea agglomerans* | *Pantoea vagans* | Flight 2 | 5 | 3.310 | 3.312 | 0.000 | 3.312 | 0.000 | 1.000 | -1.000 | 0.000 | -1.000 | 0.000 | Amensalism |
| *Pantoea agglomerans* | *Pantoea vagans* | Flight 3 | 1 | 4.640 | 4.642 | 0.000 | 4.642 | 0.000 | 1.000 | -1.000 | 0.000 | -1.000 | 0.000 | Amensalism |
| *Pantoea agglomerans* | *Pantoea vagans* | Flight 3 | 5 | 2.777 | 2.778 | 0.000 | 2.778 | 0.000 | 1.000 | -1.000 | 0.000 | -1.000 | 0.000 | Amensalism |
| *Pantoea agglomerans* | *Pantoea vagans* | Flight 3 | 7 | 2.777 | 2.778 | 0.000 | 2.778 | 0.000 | 1.000 | -1.000 | 0.000 | -1.000 | 0.000 | Amensalism |
| *Pantoea agglomerans* | *Pantoea vagans* | Flight 3 | 8 | 2.777 | 2.778 | 0.000 | 2.778 | 0.000 | 1.000 | -1.000 | 0.000 | -1.000 | 0.000 | Amensalism |
| *Pantoea agglomerans* | *Penicillium rubens* | Flight 1 | 5 | 5.233 | 0.049 | 5.233 | 0.000 | 1.000 | 0.000 | 0.000 | -1.000 | 0.000 | -1.000 | Amensalism |
| *Pantoea agglomerans* | *Rhodotorula sp. strain JG-1b* | Flight 1 | 5 | 5.233 | 0.153 | 5.233 | 0.000 | 1.000 | 0.000 | 0.000 | -1.000 | 0.000 | -1.000 | Amensalism |
| *Pantoea agglomerans* | *Rhodotorula sp. strain JG-1b* | Flight 3 | 1 | 4.640 | 0.154 | 4.640 | 0.000 | 1.000 | 0.000 | 0.000 | -1.000 | 0.000 | -1.000 | Amensalism |
| *Pantoea agglomerans* | *Salmonella enterica* | Flight 3 | 1 | 4.640 | 8.372 | 1.645 | 7.507 | 0.180 | 0.820 | -0.645 | -0.103 | -0.645 | -0.103 | Competitive |
| *Pantoea agglomerans* | *Salmonella enterica* | Flight 3 | 5 | 2.777 | 5.907 | 0.928 | 5.700 | 0.140 | 0.860 | -0.666 | -0.035 | -0.666 | 0.000 | Amensalism |
| *Pantoea agglomerans* | *Salmonella enterica* | Flight 3 | 7 | 2.777 | 5.907 | 0.928 | 5.700 | 0.140 | 0.860 | -0.666 | -0.035 | -0.666 | 0.000 | Amensalism |
| *Pantoea agglomerans* | *Salmonella enterica* | Flight 3 | 8 | 2.777 | 5.907 | 0.928 | 5.700 | 0.140 | 0.860 | -0.666 | -0.035 | -0.666 | 0.000 | Amensalism |
| *Pantoea ananatis* | *Pantoea conspicua* | Flight 1 | 5 | 7.840 | 4.632 | 8.054 | 1.471 | 0.846 | 0.154 | 0.027 | -0.683 | 0.000 | -0.683 | Amensalism |
| *Pantoea ananatis* | *Pantoea conspicua* | Flight 2 | 5 | 6.254 | 3.267 | 7.234 | 0.187 | 0.975 | 0.025 | 0.157 | -0.943 | 0.157 | -0.943 | Parasitism |
| *Pantoea ananatis* | *Pantoea conspicua* | Flight 3 | 1 | 7.622 | 4.408 | 8.274 | 0.731 | 0.919 | 0.081 | 0.086 | -0.834 | 0.000 | -0.834 | Amensalism |
| *Pantoea ananatis* | *Pantoea conspicua* | Flight 3 | 5 | 5.534 | 2.768 | 6.477 | 0.453 | 0.935 | 0.065 | 0.170 | -0.836 | 0.170 | -0.836 | Parasitism |
| *Pantoea ananatis* | *Pantoea conspicua* | Flight 3 | 7 | 5.534 | 2.768 | 6.477 | 0.453 | 0.935 | 0.065 | 0.170 | -0.836 | 0.170 | -0.836 | Parasitism |
| *Pantoea ananatis* | *Pantoea conspicua* | Flight 3 | 8 | 5.534 | 2.768 | 6.477 | 0.453 | 0.935 | 0.065 | 0.170 | -0.836 | 0.170 | -0.836 | Parasitism |
| *Pantoea ananatis* | *Pantoea dispersa* | Flight 3 | 4 | 5.534 | 1.869 | 5.406 | 0.298 | 0.948 | 0.052 | -0.023 | -0.841 | 0.000 | -0.841 | Amensalism |
| *Pantoea ananatis* | *Pantoea sp. strain 3.5.1* | Flight 1 | 5 | 7.840 | 5.233 | 8.712 | 0.657 | 0.930 | 0.070 | 0.111 | -0.874 | 0.111 | -0.874 | Parasitism |
| *Pantoea ananatis* | *Pantoea sp. strain 3.5.1* | Flight 2 | 5 | 6.254 | 3.310 | 7.234 | 0.187 | 0.975 | 0.025 | 0.157 | -0.943 | 0.157 | -0.943 | Parasitism |
| *Pantoea ananatis* | *Pantoea sp. strain A4* | Flight 3 | 1 | 7.622 | 4.256 | 8.038 | 0.465 | 0.945 | 0.055 | 0.055 | -0.891 | 0.000 | -0.891 | Amensalism |
| *Pantoea ananatis* | *Pantoea sp. strain A4* | Flight 3 | 5 | 5.534 | 2.593 | 6.044 | 0.308 | 0.951 | 0.049 | 0.092 | -0.881 | 0.000 | -0.881 | Amensalism |
| *Pantoea ananatis* | *Pantoea sp. strain A4* | Flight 3 | 7 | 5.534 | 2.593 | 6.044 | 0.308 | 0.951 | 0.049 | 0.092 | -0.881 | 0.000 | -0.881 | Amensalism |
| *Pantoea ananatis* | *Pantoea sp. strain A4* | Flight 3 | 8 | 5.534 | 2.593 | 6.044 | 0.308 | 0.951 | 0.049 | 0.092 | -0.881 | 0.000 | -0.881 | Amensalism |
| *Pantoea ananatis* | *Pantoea sp. strain At-9b* | Flight 3 | 1 | 7.622 | 3.804 | 7.622 | 0.000 | 1.000 | 0.000 | 0.000 | -1.000 | 0.000 | -1.000 | Amensalism |
| *Pantoea ananatis* | *Pantoea sp. strain At-9b* | Flight 3 | 5 | 5.534 | 2.173 | 5.534 | 0.000 | 1.000 | 0.000 | 0.000 | -1.000 | 0.000 | -1.000 | Amensalism |
| *Pantoea ananatis* | *Pantoea sp. strain At-9b* | Flight 3 | 8 | 5.534 | 2.173 | 5.534 | 0.000 | 1.000 | 0.000 | 0.000 | -1.000 | 0.000 | -1.000 | Amensalism |
| *Pantoea ananatis* | *Pantoea sp. strain FF5* | Flight 3 | 1 | 7.622 | 4.640 | 8.477 | 0.485 | 0.946 | 0.054 | 0.112 | -0.895 | 0.112 | -0.895 | Parasitism |
| *Pantoea ananatis* | *Pantoea sp. strain FF5* | Flight 3 | 5 | 5.534 | 2.777 | 6.568 | 0.299 | 0.956 | 0.044 | 0.187 | -0.892 | 0.187 | -0.892 | Parasitism |
| *Pantoea ananatis* | *Pantoea sp. strain FF5* | Flight 3 | 7 | 5.534 | 2.777 | 6.568 | 0.299 | 0.956 | 0.044 | 0.187 | -0.892 | 0.187 | -0.892 | Parasitism |
| *Pantoea ananatis* | *Pantoea sp. strain FF5* | Flight 3 | 8 | 5.534 | 2.777 | 6.568 | 0.299 | 0.956 | 0.044 | 0.187 | -0.892 | 0.187 | -0.892 | Parasitism |
| *Pantoea ananatis* | *Pantoea sp. strain IMH* | Flight 3 | 1 | 7.622 | 4.409 | 8.293 | 0.734 | 0.919 | 0.081 | 0.088 | -0.834 | 0.000 | -0.834 | Amensalism |
| *Pantoea ananatis* | *Pantoea sp. strain IMH* | Flight 3 | 5 | 5.534 | 2.634 | 6.489 | 0.455 | 0.935 | 0.065 | 0.172 | -0.827 | 0.172 | -0.827 | Parasitism |
| *Pantoea ananatis* | *Pantoea sp. strain IMH* | Flight 3 | 7 | 5.534 | 2.634 | 6.489 | 0.455 | 0.935 | 0.065 | 0.172 | -0.827 | 0.172 | -0.827 | Parasitism |
| *Pantoea ananatis* | *Pantoea sp. strain IMH* | Flight 3 | 8 | 5.534 | 2.634 | 6.489 | 0.455 | 0.935 | 0.065 | 0.172 | -0.827 | 0.172 | -0.827 | Parasitism |
| *Pantoea ananatis* | *Pantoea sp. strain NGS-ED-1003* | Flight 3 | 1 | 7.622 | 4.640 | 8.477 | 0.485 | 0.946 | 0.054 | 0.112 | -0.895 | 0.112 | -0.895 | Parasitism |
| *Pantoea ananatis* | *Pantoea sp. strain NGS-ED-1003* | Flight 3 | 5 | 5.534 | 2.777 | 6.568 | 0.299 | 0.956 | 0.044 | 0.187 | -0.892 | 0.187 | -0.892 | Parasitism |
| *Pantoea ananatis* | *Pantoea sp. strain NGS-ED-1003* | Flight 3 | 7 | 5.534 | 2.777 | 6.568 | 0.299 | 0.956 | 0.044 | 0.187 | -0.892 | 0.187 | -0.892 | Parasitism |
| *Pantoea ananatis* | *Pantoea sp. strain NGS-ED-1003* | Flight 3 | 8 | 5.534 | 2.777 | 6.568 | 0.299 | 0.956 | 0.044 | 0.187 | -0.892 | 0.187 | -0.892 | Parasitism |
| *Pantoea ananatis* | *Pantoea sp. strain OXWO6B1* | Flight 3 | 1 | 7.622 | 4.642 | 8.477 | 0.485 | 0.946 | 0.054 | 0.112 | -0.895 | 0.112 | -0.895 | Parasitism |
| *Pantoea ananatis* | *Pantoea sp. strain OXWO6B1* | Flight 3 | 5 | 5.534 | 2.779 | 6.569 | 0.299 | 0.956 | 0.044 | 0.187 | -0.892 | 0.187 | -0.892 | Parasitism |
| *Pantoea ananatis* | *Pantoea sp. strain OXWO6B1* | Flight 3 | 7 | 5.534 | 2.779 | 6.569 | 0.299 | 0.956 | 0.044 | 0.187 | -0.892 | 0.187 | -0.892 | Parasitism |
| *Pantoea ananatis* | *Pantoea sp. strain OXWO6B1* | Flight 3 | 8 | 5.534 | 2.779 | 6.569 | 0.299 | 0.956 | 0.044 | 0.187 | -0.892 | 0.187 | -0.892 | Parasitism |
| *Pantoea ananatis* | *Pantoea vagans* | Flight 2 | 5 | 6.254 | 3.312 | 7.234 | 0.187 | 0.975 | 0.025 | 0.157 | -0.943 | 0.157 | -0.943 | Parasitism |
| *Pantoea ananatis* | *Pantoea vagans* | Flight 3 | 1 | 7.622 | 4.642 | 8.477 | 0.485 | 0.946 | 0.054 | 0.112 | -0.895 | 0.112 | -0.895 | Parasitism |
| *Pantoea ananatis* | *Pantoea vagans* | Flight 3 | 5 | 5.534 | 2.778 | 6.569 | 0.299 | 0.956 | 0.044 | 0.187 | -0.892 | 0.187 | -0.892 | Parasitism |
| *Pantoea ananatis* | *Pantoea vagans* | Flight 3 | 7 | 5.534 | 2.778 | 6.569 | 0.299 | 0.956 | 0.044 | 0.187 | -0.892 | 0.187 | -0.892 | Parasitism |
| *Pantoea ananatis* | *Pantoea vagans* | Flight 3 | 8 | 5.534 | 2.778 | 6.569 | 0.299 | 0.956 | 0.044 | 0.187 | -0.892 | 0.187 | -0.892 | Parasitism |
| *Pantoea ananatis* | *Penicillium rubens* | Flight 1 | 2 | 7.840 | 0.049 | 7.480 | 0.000 | 1.000 | 0.000 | 0.000 | -1.000 | 0.000 | -1.000 | Amensalism |
| *Pantoea ananatis* | *Penicillium rubens* | Flight 1 | 5 | 7.840 | 0.049 | 7.840 | 0.000 | 1.000 | 0.000 | 0.000 | -1.000 | 0.000 | -1.000 | Amensalism |
| *Pantoea ananatis* | *Rahnella aquatilis* | Flight 3 | 4 | 5.534 | 2.624 | 0.361 | 6.602 | 0.052 | 0.948 | -0.935 | 1.516 | -0.935 | 1.516 | Parasitism |

| Microorganism A | Microorganism B | Flight | Location | Vbio of A | Vbio of B | Vbio of A in AB | Vbio of B in AB | Ratio of A in AB | Ratio of B in AB | Effect of AB on A | Effect of AB on B | Significant effect on A | Significant effect on B | Interaction type |
|---|---|---|---|---|---|---|---|---|---|---|---|---|---|---|
| *Pantoea ananatis* | *Rhodotorula sp. strain JG-1b* | Flight 1 | 2 | 7.840 | 0.153 | 7.840 | 0.000 | 1.000 | 0.000 | 0.000 | -1.000 | 0.000 | -1.000 | Amensalism |
| *Pantoea ananatis* | *Rhodotorula sp. strain JG-1b* | Flight 1 | 5 | 7.840 | 0.153 | 7.840 | 0.000 | 1.000 | 0.000 | 0.000 | -1.000 | 0.000 | -1.000 | Amensalism |
| *Pantoea ananatis* | *Rhodotorula sp. strain JG-1b* | Flight 3 | 1 | 7.622 | 0.154 | 7.622 | 0.000 | 1.000 | 0.000 | 0.000 | -1.000 | 0.000 | -1.000 | Amensalism |
| *Pantoea ananatis* | *Salmonella enterica* | Flight 1 | 2 | 7.840 | 8.478 | 10.152 | 9.108 | 0.527 | 0.473 | 0.295 | 0.074 | 0.295 | 0.000 | Commensals |
| *Pantoea ananatis* | *Salmonella enterica* | Flight 3 | 1 | 7.622 | 8.372 | 10.025 | 8.993 | 0.527 | 0.473 | 0.315 | 0.074 | 0.315 | 0.000 | Commensals |
| *Pantoea ananatis* | *Salmonella enterica* | Flight 3 | 5 | 5.534 | 5.907 | 7.753 | 6.893 | 0.529 | 0.471 | 0.401 | 0.167 | 0.401 | 0.167 | Mutualism |
| *Pantoea ananatis* | *Salmonella enterica* | Flight 3 | 7 | 5.534 | 5.907 | 7.753 | 6.893 | 0.529 | 0.471 | 0.401 | 0.167 | 0.401 | 0.167 | Mutualism |
| *Pantoea ananatis* | *Salmonella enterica* | Flight 3 | 8 | 5.534 | 5.907 | 7.753 | 6.893 | 0.529 | 0.471 | 0.401 | 0.167 | 0.401 | 0.167 | Mutualism |
| *Pantoea ananatis* | *Shigella sonnei* | Flight 1 | 2 | 7.840 | 8.331 | 10.072 | 9.036 | 0.527 | 0.473 | 0.285 | 0.085 | 0.285 | 0.000 | Commensals |
| *Pantoea conspicua* | *Pantoea sp. strain 3.5.1* | Flight 1 | 5 | 4.632 | 5.233 | 4.879 | 0.353 | 0.932 | 0.068 | 0.053 | -0.932 | 0.000 | -0.932 | Amensalism |
| *Pantoea conspicua* | *Pantoea sp. strain 3.5.1* | Flight 2 | 5 | 3.267 | 3.310 | 3.286 | 0.025 | 0.992 | 0.008 | 0.006 | -0.992 | 0.000 | -0.992 | Amensalism |
| *Pantoea conspicua* | *Pantoea sp. strain A4* | Flight 3 | 1 | 4.408 | 4.256 | 0.070 | 4.580 | 0.015 | 0.985 | -0.984 | 0.076 | -0.984 | 0.000 | Amensalism |
| *Pantoea conspicua* | *Pantoea sp. strain A4* | Flight 3 | 5 | 2.768 | 2.593 | 0.039 | 2.743 | 0.014 | 0.986 | -0.986 | 0.058 | -0.986 | 0.000 | Amensalism |
| *Pantoea conspicua* | *Pantoea sp. strain A4* | Flight 3 | 7 | 2.768 | 2.593 | 0.039 | 2.743 | 0.014 | 0.986 | -0.986 | 0.058 | -0.986 | 0.000 | Amensalism |
| *Pantoea conspicua* | *Pantoea sp. strain A4* | Flight 3 | 8 | 2.768 | 2.593 | 0.039 | 2.743 | 0.014 | 0.986 | -0.986 | 0.058 | -0.986 | 0.000 | Amensalism |
| *Pantoea conspicua* | *Pantoea sp. strain At-9b* | Flight 3 | 1 | 4.408 | 3.804 | 0.443 | 4.199 | 0.096 | 0.904 | -0.899 | 0.104 | -0.899 | 0.104 | Parasitism |
| *Pantoea conspicua* | *Pantoea sp. strain At-9b* | Flight 3 | 5 | 2.768 | 2.173 | 0.156 | 2.622 | 0.056 | 0.944 | -0.943 | 0.207 | -0.943 | 0.207 | Parasitism |
| *Pantoea conspicua* | *Pantoea sp. strain At-9b* | Flight 3 | 8 | 2.768 | 2.173 | 0.156 | 2.622 | 0.056 | 0.944 | -0.943 | 0.207 | -0.943 | 0.207 | Parasitism |
| *Pantoea conspicua* | *Pantoea sp. strain FF5* | Flight 3 | 1 | 4.408 | 4.640 | 0.000 | 4.640 | 0.000 | 1.000 | -1.000 | 0.000 | -1.000 | 0.000 | Amensalism |
| *Pantoea conspicua* | *Pantoea sp. strain FF5* | Flight 3 | 5 | 2.768 | 2.777 | 2.776 | 0.001 | 0.999 | 0.001 | 0.003 | -0.999 | 0.000 | -0.999 | Amensalism |
| *Pantoea conspicua* | *Pantoea sp. strain FF5* | Flight 3 | 7 | 2.768 | 2.777 | 2.776 | 0.001 | 0.999 | 0.001 | 0.003 | -0.999 | 0.000 | -0.999 | Amensalism |
| *Pantoea conspicua* | *Pantoea sp. strain FF5* | Flight 3 | 8 | 2.768 | 2.777 | 2.776 | 0.001 | 0.999 | 0.001 | 0.003 | -0.999 | 0.000 | -0.999 | Amensalism |
| *Pantoea conspicua* | *Pantoea sp. strain IMH* | Flight 3 | 1 | 4.408 | 4.409 | 0.081 | 4.584 | 0.017 | 0.983 | -0.982 | 0.040 | -0.982 | 0.000 | Amensalism |
| *Pantoea conspicua* | *Pantoea sp. strain IMH* | Flight 3 | 5 | 2.768 | 2.634 | 0.029 | 2.791 | 0.010 | 0.990 | -0.990 | 0.059 | -0.990 | 0.000 | Amensalism |
| *Pantoea conspicua* | *Pantoea sp. strain IMH* | Flight 3 | 7 | 2.768 | 2.634 | 0.029 | 2.791 | 0.010 | 0.990 | -0.990 | 0.059 | -0.990 | 0.000 | Amensalism |
| *Pantoea conspicua* | *Pantoea sp. strain IMH* | Flight 3 | 8 | 2.768 | 2.634 | 0.029 | 2.791 | 0.010 | 0.990 | -0.990 | 0.059 | -0.990 | 0.000 | Amensalism |
| *Pantoea conspicua* | *Pantoea sp. strain NGS-ED-1003* | Flight 3 | 1 | 4.408 | 4.640 | 0.000 | 4.640 | 0.000 | 1.000 | -1.000 | 0.000 | -1.000 | 0.000 | Amensalism |
| *Pantoea conspicua* | *Pantoea sp. strain NGS-ED-1003* | Flight 3 | 5 | 2.768 | 2.777 | 2.776 | 0.001 | 0.999 | 0.001 | 0.003 | -0.999 | 0.000 | -0.999 | Amensalism |
| *Pantoea conspicua* | *Pantoea sp. strain NGS-ED-1003* | Flight 3 | 7 | 2.768 | 2.777 | 2.776 | 0.001 | 0.999 | 0.001 | 0.003 | -0.999 | 0.000 | -0.999 | Amensalism |
| *Pantoea conspicua* | *Pantoea sp. strain NGS-ED-1003* | Flight 3 | 8 | 2.768 | 2.777 | 2.776 | 0.001 | 0.999 | 0.001 | 0.003 | -0.999 | 0.000 | -0.999 | Amensalism |
| *Pantoea conspicua* | *Pantoea sp. strain OXWO6B1* | Flight 3 | 1 | 4.408 | 4.642 | 0.000 | 4.642 | 0.000 | 1.000 | -1.000 | 0.000 | -1.000 | 0.000 | Amensalism |
| *Pantoea conspicua* | *Pantoea sp. strain OXWO6B1* | Flight 3 | 5 | 2.768 | 2.779 | 0.000 | 2.779 | 0.000 | 1.000 | -1.000 | 0.000 | -1.000 | 0.000 | Amensalism |
| *Pantoea conspicua* | *Pantoea sp. strain OXWO6B1* | Flight 3 | 7 | 2.768 | 2.779 | 0.000 | 2.779 | 0.000 | 1.000 | -1.000 | 0.000 | -1.000 | 0.000 | Amensalism |
| *Pantoea conspicua* | *Pantoea sp. strain OXWO6B1* | Flight 3 | 8 | 2.768 | 2.779 | 0.000 | 2.779 | 0.000 | 1.000 | -1.000 | 0.000 | -1.000 | 0.000 | Amensalism |
| *Pantoea conspicua* | *Pantoea vagans* | Flight 2 | 5 | 3.267 | 3.312 | 0.000 | 3.312 | 0.000 | 1.000 | -1.000 | 0.000 | -1.000 | 0.000 | Amensalism |
| *Pantoea conspicua* | *Pantoea vagans* | Flight 3 | 1 | 4.408 | 4.642 | 0.000 | 4.642 | 0.000 | 1.000 | -1.000 | 0.000 | -1.000 | 0.000 | Amensalism |
| *Pantoea conspicua* | *Pantoea vagans* | Flight 3 | 5 | 2.768 | 2.778 | 0.000 | 2.778 | 0.000 | 1.000 | -1.000 | 0.000 | -1.000 | 0.000 | Amensalism |
| *Pantoea conspicua* | *Pantoea vagans* | Flight 3 | 7 | 2.768 | 2.778 | 0.000 | 2.778 | 0.000 | 1.000 | -1.000 | 0.000 | -1.000 | 0.000 | Amensalism |
| *Pantoea conspicua* | *Pantoea vagans* | Flight 3 | 8 | 2.768 | 2.778 | 0.000 | 2.778 | 0.000 | 1.000 | -1.000 | 0.000 | -1.000 | 0.000 | Amensalism |
| *Pantoea conspicua* | *Penicillium rubens* | Flight 1 | 5 | 4.632 | 0.049 | 4.632 | 0.000 | 1.000 | 0.000 | 0.000 | -1.000 | 0.000 | -1.000 | Amensalism |
| *Pantoea conspicua* | *Rhodotorula sp. strain JG-1b* | Flight 1 | 5 | 4.632 | 0.153 | 4.632 | 0.000 | 1.000 | 0.000 | 0.000 | -1.000 | 0.000 | -1.000 | Amensalism |
| *Pantoea conspicua* | *Rhodotorula sp. strain JG-1b* | Flight 3 | 1 | 4.408 | 0.154 | 4.408 | 0.000 | 1.000 | 0.000 | 0.000 | -1.000 | 0.000 | -1.000 | Amensalism |
| *Pantoea conspicua* | *Salmonella enterica* | Flight 3 | 1 | 4.408 | 8.372 | 1.645 | 7.506 | 0.180 | 0.820 | -0.627 | -0.103 | -0.627 | -0.103 | Competitive |
| *Pantoea conspicua* | *Salmonella enterica* | Flight 3 | 5 | 2.768 | 5.907 | 0.928 | 5.700 | 0.140 | 0.860 | -0.665 | -0.035 | -0.665 | 0.000 | Amensalism |
| *Pantoea conspicua* | *Salmonella enterica* | Flight 3 | 7 | 2.768 | 5.907 | 0.928 | 5.700 | 0.140 | 0.860 | -0.665 | -0.035 | -0.665 | 0.000 | Amensalism |
| *Pantoea conspicua* | *Salmonella enterica* | Flight 3 | 8 | 2.768 | 5.907 | 0.928 | 5.700 | 0.140 | 0.860 | -0.665 | -0.035 | -0.665 | 0.000 | Amensalism |
| *Pantoea dispersa* | *Rahnella aquatilis* | Flight 3 | 4 | 1.869 | 2.624 | 0.027 | 2.710 | 0.010 | 0.990 | -0.985 | 0.033 | -0.985 | 0.000 | Amensalism |
| *Pantoea sp. strain 3.5.1* | *Pantoea vagans* | Flight 2 | 5 | 3.310 | 3.312 | 0.000 | 3.312 | 0.000 | 1.000 | -1.000 | 0.000 | -1.000 | 0.000 | Amensalism |
| *Pantoea sp. strain 3.5.1* | *Penicillium rubens* | Flight 1 | 5 | 5.233 | 0.049 | 5.233 | 0.000 | 1.000 | 0.000 | 0.000 | -1.000 | 0.000 | -1.000 | Amensalism |
| *Pantoea sp. strain 3.5.1* | *Rhodotorula sp. strain JG-1b* | Flight 1 | 5 | 5.233 | 0.153 | 5.233 | 0.000 | 1.000 | 0.000 | 0.000 | -1.000 | 0.000 | -1.000 | Amensalism |
| *Pantoea sp. strain A4* | *Pantoea sp. strain At-9b* | Flight 3 | 1 | 4.256 | 3.804 | 0.124 | 4.191 | 0.029 | 0.971 | -0.971 | 0.102 | -0.971 | 0.102 | Parasitism |
| *Pantoea sp. strain A4* | *Pantoea sp. strain At-9b* | Flight 3 | 5 | 2.593 | 2.173 | 0.116 | 2.478 | 0.045 | 0.955 | -0.955 | 0.140 | -0.955 | 0.140 | Parasitism |
| *Pantoea sp. strain A4* | *Pantoea sp. strain At-9b* | Flight 3 | 8 | 2.593 | 2.173 | 0.116 | 2.478 | 0.045 | 0.955 | -0.955 | 0.140 | -0.955 | 0.140 | Parasitism |
| *Pantoea sp. strain A4* | *Pantoea sp. strain FF5* | Flight 3 | 1 | 4.256 | 4.640 | 4.596 | 0.053 | 0.989 | 0.011 | 0.080 | -0.988 | 0.000 | -0.988 | Amensalism |
| *Pantoea sp. strain A4* | *Pantoea sp. strain FF5* | Flight 3 | 5 | 2.593 | 2.777 | 2.753 | 0.030 | 0.989 | 0.011 | 0.062 | -0.989 | 0.000 | -0.989 | Amensalism |
| *Pantoea sp. strain A4* | *Pantoea sp. strain FF5* | Flight 3 | 7 | 2.593 | 2.777 | 2.753 | 0.030 | 0.989 | 0.011 | 0.062 | -0.989 | 0.000 | -0.989 | Amensalism |
| *Pantoea sp. strain A4* | *Pantoea sp. strain FF5* | Flight 3 | 8 | 2.593 | 2.777 | 2.753 | 0.030 | 0.989 | 0.011 | 0.062 | -0.989 | 0.000 | -0.989 | Amensalism |
| *Pantoea sp. strain A4* | *Pantoea sp. strain IMH* | Flight 3 | 1 | 4.256 | 4.409 | 0.081 | 4.584 | 0.017 | 0.983 | -0.981 | 0.040 | -0.981 | 0.000 | Amensalism |
| *Pantoea sp. strain A4* | *Pantoea sp. strain IMH* | Flight 3 | 5 | 2.593 | 2.634 | 0.029 | 2.791 | 0.010 | 0.990 | -0.989 | 0.059 | -0.989 | 0.000 | Amensalism |
| *Pantoea sp. strain A4* | *Pantoea sp. strain IMH* | Flight 3 | 7 | 2.593 | 2.634 | 0.029 | 2.791 | 0.010 | 0.990 | -0.989 | 0.059 | -0.989 | 0.000 | Amensalism |
| *Pantoea sp. strain A4* | *Pantoea sp. strain IMH* | Flight 3 | 8 | 2.593 | 2.634 | 0.029 | 2.791 | 0.010 | 0.990 | -0.989 | 0.059 | -0.989 | 0.000 | Amensalism |
| *Pantoea sp. strain A4* | *Pantoea sp. strain NGS-ED-1003* | Flight 3 | 1 | 4.256 | 4.640 | 4.596 | 0.053 | 0.988 | 0.012 | 0.080 | -0.988 | 0.000 | -0.988 | Amensalism |
| *Pantoea sp. strain A4* | *Pantoea sp. strain NGS-ED-1003* | Flight 3 | 5 | 2.593 | 2.777 | 2.753 | 0.029 | 0.989 | 0.011 | 0.062 | -0.989 | 0.000 | -0.989 | Amensalism |
| *Pantoea sp. strain A4* | *Pantoea sp. strain NGS-ED-1003* | Flight 3 | 7 | 2.593 | 2.777 | 2.753 | 0.029 | 0.989 | 0.011 | 0.062 | -0.989 | 0.000 | -0.989 | Amensalism |
| *Pantoea sp. strain A4* | *Pantoea sp. strain NGS-ED-1003* | Flight 3 | 8 | 2.593 | 2.777 | 2.753 | 0.029 | 0.989 | 0.011 | 0.062 | -0.989 | 0.000 | -0.989 | Amensalism |
| *Pantoea sp. strain A4* | *Pantoea sp. strain OXWO6B1* | Flight 3 | 1 | 4.256 | 4.642 | 0.002 | 4.649 | 0.000 | 1.000 | -0.999 | 0.002 | -0.999 | 0.000 | Amensalism |

| Microorganism A | Microorganism B | Flight | Location | Vbio of A | Vbio of B | Vbio of A in AB | Vbio of B in AB | Ratio of A in AB | Ratio of B in AB | Effect of AB on A | Effect of AB on B | Significant effect on A | Significant effect on B | Interaction type |
|---|---|---|---|---|---|---|---|---|---|---|---|---|---|---|
| *Pantoea sp. strain A4* | *Pantoea sp. strain OXWO6B1* | Flight 3 | 5 | 2.593 | 2.779 | 0.002 | 2.782 | 0.001 | 0.999 | -0.999 | 0.001 | -0.999 | 0.000 | Amensalism |
| *Pantoea sp. strain A4* | *Pantoea sp. strain OXWO6B1* | Flight 3 | 7 | 2.593 | 2.779 | 0.002 | 2.782 | 0.001 | 0.999 | -0.999 | 0.001 | -0.999 | 0.000 | Amensalism |
| *Pantoea sp. strain A4* | *Pantoea sp. strain OXWO6B1* | Flight 3 | 8 | 2.593 | 2.779 | 0.002 | 2.782 | 0.001 | 0.999 | -0.999 | 0.001 | -0.999 | 0.000 | Amensalism |
| *Pantoea sp. strain A4* | *Pantoea vagans* | Flight 3 | 1 | 4.256 | 4.642 | 0.002 | 4.649 | 0.000 | 1.000 | -0.999 | 0.002 | -0.999 | 0.000 | Amensalism |
| *Pantoea sp. strain A4* | *Pantoea vagans* | Flight 3 | 5 | 2.593 | 2.778 | 0.002 | 2.782 | 0.001 | 0.999 | -0.999 | 0.001 | -0.999 | 0.000 | Amensalism |
| *Pantoea sp. strain A4* | *Pantoea vagans* | Flight 3 | 7 | 2.593 | 2.778 | 0.002 | 2.782 | 0.001 | 0.999 | -0.999 | 0.001 | -0.999 | 0.000 | Amensalism |
| *Pantoea sp. strain A4* | *Pantoea vagans* | Flight 3 | 8 | 2.593 | 2.778 | 0.002 | 2.782 | 0.001 | 0.999 | -0.999 | 0.001 | -0.999 | 0.000 | Amensalism |
| *Pantoea sp. strain A4* | *Rhodotorula sp. strain JG-1b* | Flight 3 | 1 | 4.256 | 0.154 | 4.256 | 0.000 | 1.000 | 0.000 | 0.000 | -1.000 | 0.000 | -1.000 | Amensalism |
| *Pantoea sp. strain A4* | *Salmonella enterica* | Flight 3 | 1 | 4.256 | 8.372 | 1.645 | 7.507 | 0.180 | 0.820 | -0.613 | -0.103 | -0.613 | -0.103 | Competitive |
| *Pantoea sp. strain A4* | *Salmonella enterica* | Flight 3 | 5 | 2.593 | 5.907 | 0.928 | 5.700 | 0.140 | 0.860 | -0.642 | -0.035 | -0.642 | 0.000 | Amensalism |
| *Pantoea sp. strain A4* | *Salmonella enterica* | Flight 3 | 7 | 2.593 | 5.907 | 0.928 | 5.700 | 0.140 | 0.860 | -0.642 | -0.035 | -0.642 | 0.000 | Amensalism |
| *Pantoea sp. strain A4* | *Salmonella enterica* | Flight 3 | 8 | 2.593 | 5.907 | 0.928 | 5.700 | 0.140 | 0.860 | -0.642 | -0.035 | -0.642 | 0.000 | Amensalism |
| *Pantoea sp. strain At-9b* | *Pantoea sp. strain FF5* | Flight 3 | 1 | 3.804 | 4.640 | 4.199 | 0.443 | 0.905 | 0.095 | 0.104 | -0.904 | 0.104 | -0.904 | Parasitism |
| *Pantoea sp. strain At-9b* | *Pantoea sp. strain FF5* | Flight 3 | 5 | 2.173 | 2.777 | 2.622 | 0.156 | 0.944 | 0.056 | 0.207 | -0.944 | 0.207 | -0.944 | Parasitism |
| *Pantoea sp. strain At-9b* | *Pantoea sp. strain FF5* | Flight 3 | 8 | 2.173 | 2.777 | 2.622 | 0.156 | 0.944 | 0.056 | 0.207 | -0.944 | 0.207 | -0.944 | Parasitism |
| *Pantoea sp. strain At-9b* | *Pantoea sp. strain IMH* | Flight 3 | 1 | 3.804 | 4.409 | 0.081 | 4.584 | 0.017 | 0.983 | -0.979 | 0.040 | -0.979 | 0.000 | Amensalism |
| *Pantoea sp. strain At-9b* | *Pantoea sp. strain IMH* | Flight 3 | 5 | 2.173 | 2.634 | 0.029 | 2.791 | 0.010 | 0.990 | -0.987 | 0.059 | -0.987 | 0.000 | Amensalism |
| *Pantoea sp. strain At-9b* | *Pantoea sp. strain IMH* | Flight 3 | 8 | 2.173 | 2.634 | 0.029 | 2.791 | 0.010 | 0.990 | -0.987 | 0.059 | -0.987 | 0.000 | Amensalism |
| *Pantoea sp. strain At-9b* | *Pantoea sp. strain NGS-ED-1003* | Flight 3 | 1 | 3.804 | 4.640 | 4.199 | 0.443 | 0.905 | 0.095 | 0.104 | -0.904 | 0.104 | -0.904 | Parasitism |
| *Pantoea sp. strain At-9b* | *Pantoea sp. strain NGS-ED-1003* | Flight 3 | 5 | 2.173 | 2.777 | 2.622 | 0.156 | 0.944 | 0.056 | 0.207 | -0.944 | 0.207 | -0.944 | Parasitism |
| *Pantoea sp. strain At-9b* | *Pantoea sp. strain NGS-ED-1003* | Flight 3 | 8 | 2.173 | 2.777 | 2.622 | 0.156 | 0.944 | 0.056 | 0.207 | -0.944 | 0.207 | -0.944 | Parasitism |
| *Pantoea sp. strain At-9b* | *Pantoea sp. strain OXWO6B1* | Flight 3 | 1 | 3.804 | 4.642 | 0.000 | 4.642 | 0.000 | 1.000 | -1.000 | 0.000 | -1.000 | 0.000 | Amensalism |
| *Pantoea sp. strain At-9b* | *Pantoea sp. strain OXWO6B1* | Flight 3 | 5 | 2.173 | 2.779 | 0.000 | 2.779 | 0.000 | 1.000 | -1.000 | 0.000 | -1.000 | 0.000 | Amensalism |
| *Pantoea sp. strain At-9b* | *Pantoea sp. strain OXWO6B1* | Flight 3 | 8 | 2.173 | 2.779 | 0.000 | 2.779 | 0.000 | 1.000 | -1.000 | 0.000 | -1.000 | 0.000 | Amensalism |
| *Pantoea sp. strain At-9b* | *Pantoea vagans* | Flight 3 | 1 | 3.804 | 4.642 | 4.199 | 0.443 | 0.905 | 0.095 | 0.104 | -0.905 | 0.104 | -0.905 | Parasitism |
| *Pantoea sp. strain At-9b* | *Pantoea vagans* | Flight 3 | 5 | 2.173 | 2.778 | 2.622 | 0.156 | 0.944 | 0.056 | 0.207 | -0.944 | 0.207 | -0.944 | Parasitism |
| *Pantoea sp. strain At-9b* | *Pantoea vagans* | Flight 3 | 8 | 2.173 | 2.778 | 2.622 | 0.156 | 0.944 | 0.056 | 0.207 | -0.944 | 0.207 | -0.944 | Parasitism |
| *Pantoea sp. strain At-9b* | *Rhodotorula sp. strain JG-1b* | Flight 3 | 1 | 3.804 | 0.154 | 3.804 | 0.000 | 1.000 | 0.000 | 0.000 | -1.000 | 0.000 | -1.000 | Amensalism |
| *Pantoea sp. strain At-9b* | *Salmonella enterica* | Flight 3 | 1 | 3.804 | 8.372 | 0.185 | 8.633 | 0.021 | 0.979 | -0.951 | 0.031 | -0.951 | 0.000 | Amensalism |
| *Pantoea sp. strain At-9b* | *Salmonella enterica* | Flight 3 | 5 | 2.173 | 5.907 | 0.199 | 6.380 | 0.030 | 0.970 | -0.908 | 0.080 | -0.908 | 0.000 | Amensalism |
| *Pantoea sp. strain At-9b* | *Salmonella enterica* | Flight 3 | 8 | 2.173 | 5.907 | 0.199 | 6.380 | 0.030 | 0.970 | -0.908 | 0.080 | -0.908 | 0.000 | Amensalism |
| *Pantoea sp. strain FF5* | *Pantoea sp. strain IMH* | Flight 3 | 1 | 4.640 | 4.409 | 0.081 | 4.584 | 0.017 | 0.983 | -0.982 | 0.040 | -0.982 | 0.000 | Amensalism |
| *Pantoea sp. strain FF5* | *Pantoea sp. strain IMH* | Flight 3 | 5 | 2.777 | 2.634 | 0.029 | 2.791 | 0.010 | 0.990 | -0.990 | 0.059 | -0.990 | 0.000 | Amensalism |
| *Pantoea sp. strain FF5* | *Pantoea sp. strain IMH* | Flight 3 | 7 | 2.777 | 2.634 | 0.029 | 2.791 | 0.010 | 0.990 | -0.990 | 0.059 | -0.990 | 0.000 | Amensalism |
| *Pantoea sp. strain FF5* | *Pantoea sp. strain IMH* | Flight 3 | 8 | 2.777 | 2.634 | 0.029 | 2.791 | 0.010 | 0.990 | -0.990 | 0.059 | -0.990 | 0.000 | Amensalism |
| *Pantoea sp. strain FF5* | *Pantoea sp. strain NGS-ED-1003* | Flight 3 | 1 | 4.640 | 4.640 | 0.000 | 4.640 | 0.000 | 1.000 | -1.000 | 0.000 | -1.000 | 0.000 | Amensalism |
| *Pantoea sp. strain FF5* | *Pantoea sp. strain NGS-ED-1003* | Flight 3 | 5 | 2.777 | 2.777 | 1.389 | 1.389 | 0.500 | 0.500 | -0.500 | -0.500 | -0.500 | -0.500 | Competitive |
| *Pantoea sp. strain FF5* | *Pantoea sp. strain NGS-ED-1003* | Flight 3 | 7 | 2.777 | 2.777 | 1.389 | 1.389 | 0.500 | 0.500 | -0.500 | -0.500 | -0.500 | -0.500 | Competitive |
| *Pantoea sp. strain FF5* | *Pantoea sp. strain NGS-ED-1003* | Flight 3 | 8 | 2.777 | 2.777 | 1.389 | 1.389 | 0.500 | 0.500 | -0.500 | -0.500 | -0.500 | -0.500 | Competitive |
| *Pantoea sp. strain FF5* | *Pantoea sp. strain OXWO6B1* | Flight 3 | 1 | 4.640 | 4.642 | 0.000 | 4.642 | 0.000 | 1.000 | -1.000 | 0.000 | -1.000 | 0.000 | Amensalism |
| *Pantoea sp. strain FF5* | *Pantoea sp. strain OXWO6B1* | Flight 3 | 5 | 2.777 | 2.779 | 0.000 | 2.779 | 0.000 | 1.000 | -1.000 | 0.000 | -1.000 | 0.000 | Amensalism |
| *Pantoea sp. strain FF5* | *Pantoea sp. strain OXWO6B1* | Flight 3 | 7 | 2.777 | 2.779 | 0.000 | 2.779 | 0.000 | 1.000 | -1.000 | 0.000 | -1.000 | 0.000 | Amensalism |
| *Pantoea sp. strain FF5* | *Pantoea sp. strain OXWO6B1* | Flight 3 | 8 | 2.777 | 2.779 | 0.000 | 2.779 | 0.000 | 1.000 | -1.000 | 0.000 | -1.000 | 0.000 | Amensalism |
| *Pantoea sp. strain FF5* | *Pantoea vagans* | Flight 3 | 1 | 4.640 | 4.642 | 0.000 | 4.642 | 0.000 | 1.000 | -1.000 | 0.000 | -1.000 | 0.000 | Amensalism |
| *Pantoea sp. strain FF5* | *Pantoea vagans* | Flight 3 | 5 | 2.777 | 2.778 | 0.000 | 2.778 | 0.000 | 1.000 | -1.000 | 0.000 | -1.000 | 0.000 | Amensalism |
| *Pantoea sp. strain FF5* | *Pantoea vagans* | Flight 3 | 7 | 2.777 | 2.778 | 0.000 | 2.778 | 0.000 | 1.000 | -1.000 | 0.000 | -1.000 | 0.000 | Amensalism |
| *Pantoea sp. strain FF5* | *Pantoea vagans* | Flight 3 | 8 | 2.777 | 2.778 | 0.000 | 2.778 | 0.000 | 1.000 | -1.000 | 0.000 | -1.000 | 0.000 | Amensalism |
| *Pantoea sp. strain FF5* | *Rhodotorula sp. strain JG-1b* | Flight 3 | 1 | 4.640 | 0.154 | 4.640 | 0.000 | 1.000 | 0.000 | 0.000 | -1.000 | 0.000 | -1.000 | Amensalism |
| *Pantoea sp. strain FF5* | *Salmonella enterica* | Flight 3 | 1 | 4.640 | 8.372 | 1.645 | 7.507 | 0.180 | 0.820 | -0.645 | -0.103 | -0.645 | -0.103 | Competitive |
| *Pantoea sp. strain FF5* | *Salmonella enterica* | Flight 3 | 5 | 2.777 | 5.907 | 0.928 | 5.700 | 0.140 | 0.860 | -0.666 | -0.035 | -0.666 | 0.000 | Amensalism |
| *Pantoea sp. strain FF5* | *Salmonella enterica* | Flight 3 | 7 | 2.777 | 5.907 | 0.928 | 5.700 | 0.140 | 0.860 | -0.666 | -0.035 | -0.666 | 0.000 | Amensalism |
| *Pantoea sp. strain FF5* | *Salmonella enterica* | Flight 3 | 8 | 2.777 | 5.907 | 0.928 | 5.700 | 0.140 | 0.860 | -0.666 | -0.035 | -0.666 | 0.000 | Amensalism |
| *Pantoea sp. strain IMH* | *Pantoea sp. strain NGS-ED-1003* | Flight 3 | 1 | 4.409 | 4.640 | 4.584 | 0.081 | 0.983 | 0.017 | 0.040 | -0.982 | 0.000 | -0.982 | Amensalism |
| *Pantoea sp. strain IMH* | *Pantoea sp. strain NGS-ED-1003* | Flight 3 | 5 | 2.634 | 2.777 | 2.791 | 0.029 | 0.990 | 0.010 | 0.059 | -0.990 | 0.000 | -0.990 | Amensalism |
| *Pantoea sp. strain IMH* | *Pantoea sp. strain NGS-ED-1003* | Flight 3 | 7 | 2.634 | 2.777 | 2.791 | 0.029 | 0.990 | 0.010 | 0.059 | -0.990 | 0.000 | -0.990 | Amensalism |
| *Pantoea sp. strain IMH* | *Pantoea sp. strain NGS-ED-1003* | Flight 3 | 8 | 2.634 | 2.777 | 2.791 | 0.029 | 0.990 | 0.010 | 0.059 | -0.990 | 0.000 | -0.990 | Amensalism |
| *Pantoea sp. strain IMH* | *Pantoea sp. strain OXWO6B1* | Flight 3 | 1 | 4.409 | 4.642 | 4.584 | 0.081 | 0.983 | 0.017 | 0.040 | -0.982 | 0.000 | -0.982 | Amensalism |
| *Pantoea sp. strain IMH* | *Pantoea sp. strain OXWO6B1* | Flight 3 | 5 | 2.634 | 2.779 | 2.791 | 0.029 | 0.990 | 0.010 | 0.059 | -0.990 | 0.000 | -0.990 | Amensalism |
| *Pantoea sp. strain IMH* | *Pantoea sp. strain OXWO6B1* | Flight 3 | 7 | 2.634 | 2.779 | 2.791 | 0.029 | 0.990 | 0.010 | 0.059 | -0.990 | 0.000 | -0.990 | Amensalism |
| *Pantoea sp. strain IMH* | *Pantoea sp. strain OXWO6B1* | Flight 3 | 8 | 2.634 | 2.779 | 2.791 | 0.029 | 0.990 | 0.010 | 0.059 | -0.990 | 0.000 | -0.990 | Amensalism |
| *Pantoea sp. strain IMH* | *Pantoea vagans* | Flight 3 | 1 | 4.409 | 4.642 | 4.584 | 0.081 | 0.983 | 0.017 | 0.040 | -0.982 | 0.000 | -0.982 | Amensalism |
| *Pantoea sp. strain IMH* | *Pantoea vagans* | Flight 3 | 5 | 2.634 | 2.778 | 2.791 | 0.029 | 0.990 | 0.010 | 0.059 | -0.990 | 0.000 | -0.990 | Amensalism |
| *Pantoea sp. strain IMH* | *Pantoea vagans* | Flight 3 | 7 | 2.634 | 2.778 | 2.791 | 0.029 | 0.990 | 0.010 | 0.059 | -0.990 | 0.000 | -0.990 | Amensalism |
| *Pantoea sp. strain IMH* | *Pantoea vagans* | Flight 3 | 8 | 2.634 | 2.778 | 2.791 | 0.029 | 0.990 | 0.010 | 0.059 | -0.990 | 0.000 | -0.990 | Amensalism |
| *Pantoea sp. strain IMH* | *Rhodotorula sp. strain JG-1b* | Flight 3 | 1 | 4.409 | 0.154 | 4.409 | 0.000 | 1.000 | 0.000 | 0.000 | -1.000 | 0.000 | -1.000 | Amensalism |
| *Pantoea sp. strain IMH* | *Salmonella enterica* | Flight 3 | 1 | 4.409 | 8.372 | 1.641 | 7.020 | 0.190 | 0.810 | -0.628 | -0.161 | -0.628 | -0.161 | Competitive |

| Microorganism A | Microorganism B | Flight | Location | Vbio of A | Vbio of B | Vbio of A in AB | Vbio of B in AB | Ratio of A in AB | Ratio of B in AB | Effect of AB on A | Effect of AB on B | Significant effect on A | Significant effect on B | Interaction type |
|---|---|---|---|---|---|---|---|---|---|---|---|---|---|---|
| *Pantoea sp. strain IMH* | *Salmonella enterica* | Flight 3 | 5 | 2.634 | 5.907 | 0.878 | 5.243 | 0.143 | 0.857 | -0.667 | -0.112 | -0.667 | -0.112 | Competitive |
| *Pantoea sp. strain IMH* | *Salmonella enterica* | Flight 3 | 7 | 2.634 | 5.907 | 0.878 | 5.243 | 0.143 | 0.857 | -0.667 | -0.112 | -0.667 | -0.112 | Competitive |
| *Pantoea sp. strain IMH* | *Salmonella enterica* | Flight 3 | 8 | 2.634 | 5.907 | 0.878 | 5.243 | 0.143 | 0.857 | -0.667 | -0.112 | -0.667 | -0.112 | Competitive |
| *Pantoea sp. strain NGS-ED-1003* | *Pantoea sp. strain OXWO6B1* | Flight 3 | 1 | 4.640 | 4.642 | 0.000 | 4.642 | 0.000 | 1.000 | -1.000 | 0.000 | -1.000 | 0.000 | Amensalism |
| *Pantoea sp. strain NGS-ED-1003* | *Pantoea sp. strain OXWO6B1* | Flight 3 | 5 | 2.777 | 2.779 | 0.000 | 2.779 | 0.000 | 1.000 | -1.000 | 0.000 | -1.000 | 0.000 | Amensalism |
| *Pantoea sp. strain NGS-ED-1003* | *Pantoea sp. strain OXWO6B1* | Flight 3 | 7 | 2.777 | 2.779 | 0.000 | 2.779 | 0.000 | 1.000 | -1.000 | 0.000 | -1.000 | 0.000 | Amensalism |
| *Pantoea sp. strain NGS-ED-1003* | *Pantoea sp. strain OXWO6B1* | Flight 3 | 8 | 2.777 | 2.779 | 0.000 | 2.779 | 0.000 | 1.000 | -1.000 | 0.000 | -1.000 | 0.000 | Amensalism |
| *Pantoea sp. strain NGS-ED-1003* | *Pantoea vagans* | Flight 3 | 1 | 4.640 | 4.642 | 0.000 | 4.642 | 0.000 | 1.000 | -1.000 | 0.000 | -1.000 | 0.000 | Amensalism |
| *Pantoea sp. strain NGS-ED-1003* | *Pantoea vagans* | Flight 3 | 5 | 2.777 | 2.778 | 0.000 | 2.778 | 0.000 | 1.000 | -1.000 | 0.000 | -1.000 | 0.000 | Amensalism |
| *Pantoea sp. strain NGS-ED-1003* | *Pantoea vagans* | Flight 3 | 7 | 2.777 | 2.778 | 0.000 | 2.778 | 0.000 | 1.000 | -1.000 | 0.000 | -1.000 | 0.000 | Amensalism |
| *Pantoea sp. strain NGS-ED-1003* | *Pantoea vagans* | Flight 3 | 8 | 2.777 | 2.778 | 0.000 | 2.778 | 0.000 | 1.000 | -1.000 | 0.000 | -1.000 | 0.000 | Amensalism |
| *Pantoea sp. strain NGS-ED-1003* | *Rhodotorula sp. strain JG-1b* | Flight 3 | 1 | 4.640 | 0.154 | 4.640 | 0.000 | 1.000 | 0.000 | 0.000 | -1.000 | 0.000 | -1.000 | Amensalism |
| *Pantoea sp. strain NGS-ED-1003* | *Salmonella enterica* | Flight 3 | 1 | 4.640 | 8.372 | 1.645 | 7.507 | 0.180 | 0.820 | -0.645 | -0.103 | -0.645 | -0.103 | Competitive |
| *Pantoea sp. strain NGS-ED-1003* | *Salmonella enterica* | Flight 3 | 5 | 2.777 | 5.907 | 0.928 | 5.700 | 0.140 | 0.860 | -0.666 | -0.035 | -0.666 | 0.000 | Amensalism |
| *Pantoea sp. strain NGS-ED-1003* | *Salmonella enterica* | Flight 3 | 7 | 2.777 | 5.907 | 0.928 | 5.700 | 0.140 | 0.860 | -0.666 | -0.035 | -0.666 | 0.000 | Amensalism |
| *Pantoea sp. strain NGS-ED-1003* | *Salmonella enterica* | Flight 3 | 8 | 2.777 | 5.907 | 0.928 | 5.700 | 0.140 | 0.860 | -0.666 | -0.035 | -0.666 | 0.000 | Amensalism |
| *Pantoea sp. strain OXWO6B1* | *Pantoea vagans* | Flight 3 | 1 | 4.642 | 4.642 | 4.642 | 0.000 | 1.000 | 0.000 | 0.000 | -1.000 | 0.000 | -1.000 | Amensalism |
| *Pantoea sp. strain OXWO6B1* | *Pantoea vagans* | Flight 3 | 5 | 2.779 | 2.778 | 2.779 | 0.000 | 1.000 | 0.000 | 0.000 | -1.000 | 0.000 | -1.000 | Amensalism |
| *Pantoea sp. strain OXWO6B1* | *Pantoea vagans* | Flight 3 | 7 | 2.779 | 2.778 | 2.779 | 0.000 | 1.000 | 0.000 | 0.000 | -1.000 | 0.000 | -1.000 | Amensalism |
| *Pantoea sp. strain OXWO6B1* | *Pantoea vagans* | Flight 3 | 8 | 2.779 | 2.778 | 2.779 | 0.000 | 1.000 | 0.000 | 0.000 | -1.000 | 0.000 | -1.000 | Amensalism |
| *Pantoea sp. strain OXWO6B1* | *Rhodotorula sp. strain JG-1b* | Flight 3 | 1 | 4.642 | 0.154 | 4.642 | 0.000 | 1.000 | 0.000 | 0.000 | -1.000 | 0.000 | -1.000 | Amensalism |
| *Pantoea sp. strain OXWO6B1* | *Salmonella enterica* | Flight 3 | 1 | 4.642 | 8.372 | 1.645 | 7.507 | 0.180 | 0.820 | -0.646 | -0.103 | -0.646 | -0.103 | Competitive |
| *Pantoea sp. strain OXWO6B1* | *Salmonella enterica* | Flight 3 | 5 | 2.779 | 5.907 | 0.928 | 5.700 | 0.140 | 0.860 | -0.666 | -0.035 | -0.666 | 0.000 | Amensalism |
| *Pantoea sp. strain OXWO6B1* | *Salmonella enterica* | Flight 3 | 7 | 2.779 | 5.907 | 0.928 | 5.700 | 0.140 | 0.860 | -0.666 | -0.035 | -0.666 | 0.000 | Amensalism |
| *Pantoea sp. strain OXWO6B1* | *Salmonella enterica* | Flight 3 | 8 | 2.779 | 5.907 | 0.928 | 5.700 | 0.140 | 0.860 | -0.666 | -0.035 | -0.666 | 0.000 | Amensalism |
| *Pantoea vagans* | *Rhodotorula sp. strain JG-1b* | Flight 3 | 1 | 4.642 | 0.154 | 4.642 | 0.000 | 1.000 | 0.000 | 0.000 | -1.000 | 0.000 | -1.000 | Amensalism |
| *Pantoea vagans* | *Salmonella enterica* | Flight 3 | 1 | 4.642 | 8.372 | 1.645 | 7.507 | 0.180 | 0.820 | -0.646 | -0.103 | -0.646 | -0.103 | Competitive |
| *Pantoea vagans* | *Salmonella enterica* | Flight 3 | 5 | 2.778 | 5.907 | 0.928 | 5.700 | 0.140 | 0.860 | -0.666 | -0.035 | -0.666 | 0.000 | Amensalism |
| *Pantoea vagans* | *Salmonella enterica* | Flight 3 | 7 | 2.778 | 5.907 | 0.928 | 5.700 | 0.140 | 0.860 | -0.666 | -0.035 | -0.666 | 0.000 | Amensalism |
| *Pantoea vagans* | *Salmonella enterica* | Flight 3 | 8 | 2.778 | 5.907 | 0.928 | 5.700 | 0.140 | 0.860 | -0.666 | -0.035 | -0.666 | 0.000 | Amensalism |
| *Penicillium chrysogenum* | *Penicillium flavigenum* | Flight 3 | 2 | 0.049 | 0.097 | 0.001 | 0.109 | 0.012 | 0.988 | -0.974 | 0.124 | -0.974 | 0.124 | Parasitism |
| *Penicillium chrysogenum* | *Penicillium nalgiovense* | Flight 3 | 2 | 0.049 | 0.067 | 0.002 | 0.067 | 0.033 | 0.967 | -0.954 | 0.002 | -0.954 | 0.000 | Amensalism |
| *Penicillium chrysogenum* | *Penicillium rubens* | Flight 1 | 1 | 0.071 | 0.071 | 0.000 | 0.071 | 0.000 | 1.000 | -1.000 | 0.000 | -1.000 | 0.000 | Amensalism |
| *Penicillium chrysogenum* | *Penicillium rubens* | Flight 3 | 2 | 0.049 | 0.049 | 0.000 | 0.049 | 0.000 | 1.000 | -1.000 | 0.000 | -1.000 | 0.000 | Amensalism |
| *Penicillium chrysogenum* | *Rhodotorula sp. strain JG-1b* | Flight 1 | 1 | 0.071 | 0.154 | 0.007 | 0.307 | 0.022 | 0.978 | -0.901 | 0.987 | -0.901 | 0.987 | Parasitism |
| *Penicillium chrysogenum* | *Rhodotorula sp. strain JG-1b* | Flight 3 | 2 | 0.049 | 0.156 | 0.005 | 0.238 | 0.020 | 0.980 | -0.901 | 0.527 | -0.901 | 0.527 | Parasitism |
| *Penicillium chrysogenum* | *Rhodotorula toruloides* | Flight 1 | 1 | 0.071 | 2.346 | 0.000 | 2.346 | 0.000 | 1.000 | -1.000 | 0.000 | -1.000 | 0.000 | Amensalism |
| *Penicillium chrysogenum* | *Staphylococcus aureus* | Flight 3 | 2 | 0.049 | 2.739 | 0.000 | 2.740 | 0.000 | 1.000 | -1.000 | 0.000 | -1.000 | 0.000 | Amensalism |
| *Penicillium chrysogenum* | *Staphylococcus epidermidis* | Flight 3 | 2 | 0.049 | 2.313 | 0.000 | 2.313 | 0.000 | 1.000 | -1.000 | 0.000 | -1.000 | 0.000 | Amensalism |
| *Penicillium chrysogenum* | *Staphylococcus haemolyticus* | Flight 3 | 2 | 0.049 | 2.519 | 0.000 | 2.519 | 0.000 | 1.000 | -1.000 | 0.000 | -1.000 | 0.000 | Amensalism |
| *Penicillium chrysogenum* | *Staphylococcus saprophyticus* | Flight 3 | 2 | 0.049 | 2.363 | 0.000 | 2.363 | 0.000 | 0.986 | -1.000 | 0.000 | -1.000 | 0.000 | Amensalism |
| *Penicillium chrysogenum* | *Staphylococcus sp. strain LCT-H4* | Flight 3 | 2 | 0.049 | 2.388 | 0.000 | 2.388 | 0.000 | 0.997 | -1.000 | 0.000 | -1.000 | 0.000 | Amensalism |
| *Penicillium chrysogenum* | *Staphylococcus warneri* | Flight 3 | 2 | 0.049 | 2.543 | 0.000 | 2.543 | 0.000 | 1.000 | -1.000 | 0.000 | -1.000 | 0.000 | Amensalism |
| *Penicillium flavigenum* | *Penicillium nalgiovense* | Flight 3 | 2 | 0.097 | 0.067 | 0.097 | 0.000 | 1.000 | 0.000 | 0.000 | -1.000 | 0.000 | -1.000 | Amensalism |
| *Penicillium flavigenum* | *Penicillium rubens* | Flight 3 | 2 | 0.097 | 0.049 | 0.109 | 0.001 | 0.988 | 0.012 | 0.124 | -0.974 | 0.124 | -0.974 | Parasitism |
| *Penicillium flavigenum* | *Rhodotorula sp. strain JG-1b* | Flight 3 | 2 | 0.097 | 0.156 | 0.002 | 0.232 | 0.010 | 0.990 | -0.976 | 0.486 | -0.976 | 0.486 | Parasitism |
| *Penicillium flavigenum* | *Staphylococcus aureus* | Flight 3 | 2 | 0.097 | 2.739 | 0.036 | 3.136 | 0.011 | 0.989 | -0.626 | 0.145 | -0.626 | 0.145 | Parasitism |
| *Penicillium flavigenum* | *Staphylococcus epidermidis* | Flight 3 | 2 | 0.097 | 2.313 | 0.030 | 2.680 | 0.011 | 0.989 | -0.695 | 0.159 | -0.695 | 0.159 | Parasitism |
| *Penicillium flavigenum* | *Staphylococcus haemolyticus* | Flight 3 | 2 | 0.097 | 2.519 | 0.032 | 2.861 | 0.011 | 0.989 | -0.669 | 0.136 | -0.669 | 0.136 | Parasitism |
| *Penicillium flavigenum* | *Staphylococcus saprophyticus* | Flight 3 | 2 | 0.097 | 2.363 | 0.030 | 2.742 | 0.011 | 0.989 | -0.686 | 0.161 | -0.686 | 0.161 | Parasitism |
| *Penicillium flavigenum* | *Staphylococcus sp. strain LCT-H4* | Flight 3 | 2 | 0.097 | 2.388 | 0.028 | 2.759 | 0.010 | 0.990 | -0.712 | 0.155 | -0.712 | 0.155 | Parasitism |
| *Penicillium flavigenum* | *Staphylococcus warneri* | Flight 3 | 2 | 0.097 | 2.543 | 0.033 | 2.932 | 0.011 | 0.989 | -0.658 | 0.153 | -0.658 | 0.153 | Parasitism |
| *Penicillium nalgiovense* | *Penicillium rubens* | Flight 3 | 2 | 0.067 | 0.049 | 0.067 | 0.002 | 0.967 | 0.033 | 0.002 | -0.954 | 0.000 | -0.954 | Amensalism |
| *Penicillium nalgiovense* | *Rhodotorula sp. strain JG-1b* | Flight 3 | 2 | 0.067 | 0.156 | 0.002 | 0.231 | 0.010 | 0.990 | -0.965 | 0.478 | -0.965 | 0.478 | Parasitism |
| *Penicillium nalgiovense* | *Staphylococcus aureus* | Flight 3 | 2 | 0.067 | 2.739 | 0.000 | 2.739 | 0.000 | 1.000 | -1.000 | 0.000 | -1.000 | 0.000 | Amensalism |
| *Penicillium nalgiovense* | *Staphylococcus epidermidis* | Flight 3 | 2 | 0.067 | 2.313 | 0.000 | 2.313 | 0.000 | 0.999 | -1.000 | 0.000 | -1.000 | 0.000 | Amensalism |
| *Penicillium nalgiovense* | *Staphylococcus haemolyticus* | Flight 3 | 2 | 0.067 | 2.519 | 0.000 | 2.519 | 0.000 | 1.000 | -1.000 | 0.000 | -1.000 | 0.000 | Amensalism |
| *Penicillium nalgiovense* | *Staphylococcus saprophyticus* | Flight 3 | 2 | 0.067 | 2.363 | 0.000 | 2.363 | 0.000 | 1.000 | -1.000 | 0.000 | -1.000 | 0.000 | Amensalism |
| *Penicillium nalgiovense* | *Staphylococcus sp. strain LCT-H4* | Flight 3 | 2 | 0.067 | 2.388 | 0.000 | 2.388 | 0.000 | 1.000 | -1.000 | 0.000 | -1.000 | 0.000 | Amensalism |
| *Penicillium nalgiovense* | *Staphylococcus warneri* | Flight 3 | 2 | 0.067 | 2.543 | 0.000 | 2.543 | 0.000 | 1.000 | -1.000 | 0.000 | -1.000 | 0.000 | Amensalism |
| *Penicillium rubens* | *Rhodotorula sp. strain JG-1b* | Flight 1 | 1 | 0.071 | 0.154 | 0.007 | 0.307 | 0.022 | 0.978 | -0.901 | 0.987 | -0.901 | 0.987 | Parasitism |
| *Penicillium rubens* | *Rhodotorula sp. strain JG-1b* | Flight 1 | 2 | 0.049 | 0.153 | 0.005 | 0.236 | 0.020 | 0.980 | -0.902 | 0.546 | -0.902 | 0.546 | Parasitism |
| *Penicillium rubens* | *Rhodotorula sp. strain JG-1b* | Flight 1 | 5 | 0.049 | 0.153 | 0.005 | 0.236 | 0.020 | 0.980 | -0.902 | 0.546 | -0.902 | 0.546 | Parasitism |
| *Penicillium rubens* | *Rhodotorula sp. strain JG-1b* | Flight 3 | 2 | 0.049 | 0.156 | 0.005 | 0.238 | 0.020 | 0.980 | -0.901 | 0.527 | -0.901 | 0.527 | Parasitism |
| *Penicillium rubens* | *Rhodotorula toruloides* | Flight 1 | 1 | 0.071 | 2.346 | 0.000 | 2.346 | 0.000 | 1.000 | -1.000 | 0.000 | -1.000 | 0.000 | Amensalism |
| *Penicillium rubens* | *Salmonella enterica* | Flight 1 | 2 | 0.049 | 8.478 | 0.000 | 8.478 | 0.000 | 1.000 | -1.000 | 0.000 | -1.000 | 0.000 | Amensalism |

| Microorganism A | Microorganism B | Flight | Location | Vbio of A | Vbio of B | Vbio of A in AB | Vbio of B in AB | Ratio of A in AB | Ratio of B in AB | Effect of AB on A | Effect of AB on B | Significant effect on A | Significant effect on B | Interaction type |
|---|---|---|---|---|---|---|---|---|---|---|---|---|---|---|
| *Penicillium rubens* | *Salmonella enterica* | Flight 3 | 3 | 0.023 | 6.819 | 0.000 | 6.819 | 0.000 | 1.000 | -1.000 | 0.000 | -1.000 | 0.000 | Amensalism |
| *Penicillium rubens* | *Shigella sonnei* | Flight 1 | 2 | 0.049 | 8.331 | 0.000 | 8.331 | 0.000 | 1.000 | -1.000 | 0.000 | -1.000 | 0.000 | Amensalism |
| *Penicillium rubens* | *Staphylococcus aureus* | Flight 3 | 2 | 0.049 | 2.739 | 0.000 | 2.739 | 0.000 | 1.000 | -1.000 | 0.000 | -1.000 | 0.000 | Amensalism |
| *Penicillium rubens* | *Staphylococcus epidermidis* | Flight 3 | 2 | 0.049 | 2.313 | 0.000 | 2.313 | 0.000 | 1.000 | -1.000 | 0.000 | -1.000 | 0.000 | Amensalism |
| *Penicillium rubens* | *Staphylococcus haemolyticus* | Flight 3 | 2 | 0.049 | 2.519 | 0.000 | 2.519 | 0.000 | 1.000 | -1.000 | 0.000 | -1.000 | 0.000 | Amensalism |
| *Penicillium rubens* | *Staphylococcus saprophyticus* | Flight 3 | 2 | 0.049 | 2.363 | 0.000 | 2.363 | 0.000 | 0.986 | -1.000 | 0.000 | -1.000 | 0.000 | Amensalism |
| *Penicillium rubens* | *Staphylococcus saprophyticus* | Flight 3 | 3 | 0.023 | 1.481 | 0.027 | 1.587 | 0.017 | 0.983 | 0.177 | 0.071 | 0.177 | 0.000 | Commensals |
| *Penicillium rubens* | *Staphylococcus sp. strain LCT-H4* | Flight 3 | 2 | 0.049 | 2.388 | 0.000 | 2.388 | 0.000 | 0.997 | -1.000 | 0.000 | -1.000 | 0.000 | Amensalism |
| *Penicillium rubens* | *Staphylococcus warneri* | Flight 3 | 2 | 0.049 | 2.543 | 0.000 | 2.543 | 0.000 | 1.000 | -1.000 | 0.000 | -1.000 | 0.000 | Amensalism |
| *Rhodotorula sp. strain JG-1b* | *Rhodotorula toruloides* | Flight 1 | 1 | 0.154 | 2.346 | 0.147 | 2.355 | 0.059 | 0.941 | -0.051 | 0.004 | 0.000 | 0.000 | Neutral |
| *Rhodotorula sp. strain JG-1b* | *Salmonella enterica* | Flight 1 | 2 | 0.153 | 8.478 | 0.155 | 8.489 | 0.018 | 0.982 | 0.014 | 0.001 | 0.000 | 0.000 | Neutral |
| *Rhodotorula sp. strain JG-1b* | *Salmonella enterica* | Flight 3 | 1 | 0.154 | 8.372 | 0.000 | 8.372 | 0.000 | 1.000 | -1.000 | 0.000 | -1.000 | 0.000 | Amensalism |
| *Rhodotorula sp. strain JG-1b* | *Shigella sonnei* | Flight 1 | 2 | 0.153 | 8.331 | 0.152 | 8.344 | 0.018 | 0.982 | -0.003 | 0.001 | 0.000 | 0.000 | Neutral |
| *Rhodotorula sp. strain JG-1b* | *Staphylococcus aureus* | Flight 3 | 2 | 0.156 | 2.739 | 0.601 | 2.836 | 0.175 | 0.825 | 2.850 | 0.035 | 2.850 | 0.000 | Commensals |
| *Rhodotorula sp. strain JG-1b* | *Staphylococcus epidermidis* | Flight 3 | 2 | 0.156 | 2.313 | 0.601 | 2.540 | 0.191 | 0.809 | 2.850 | 0.098 | 2.850 | 0.000 | Commensals |
| *Rhodotorula sp. strain JG-1b* | *Staphylococcus haemolyticus* | Flight 3 | 2 | 0.156 | 2.519 | 0.601 | 2.719 | 0.181 | 0.819 | 2.850 | 0.079 | 2.850 | 0.000 | Commensals |
| *Rhodotorula sp. strain JG-1b* | *Staphylococcus saprophyticus* | Flight 3 | 2 | 0.156 | 2.363 | 0.601 | 2.526 | 0.192 | 0.808 | 2.850 | 0.069 | 2.850 | 0.000 | Commensals |
| *Rhodotorula sp. strain JG-1b* | *Staphylococcus sp. strain LCT-H4* | Flight 3 | 2 | 0.156 | 2.388 | 0.601 | 2.551 | 0.191 | 0.809 | 2.850 | 0.068 | 2.850 | 0.000 | Commensals |
| *Rhodotorula sp. strain JG-1b* | *Staphylococcus warneri* | Flight 3 | 2 | 0.156 | 2.543 | 0.601 | 2.630 | 0.186 | 0.814 | 2.850 | 0.034 | 2.850 | 0.000 | Commensals |
| *Salmonella enterica* | *Shigella sonnei* | Flight 1 | 2 | 8.478 | 8.331 | 0.417 | 8.988 | 0.044 | 0.956 | -0.951 | 0.079 | -0.951 | 0.000 | Amensalism |
| *Salmonella enterica* | *Staphylococcus saprophyticus* | Flight 3 | 3 | 6.819 | 1.481 | 8.723 | 2.722 | 0.762 | 0.238 | 0.279 | 0.837 | 0.279 | 0.837 | Mutualism |
| *Staphylococcus aureus* | *Staphylococcus epidermidis* | Flight 3 | 2 | 2.739 | 2.313 | 0.105 | 2.764 | 0.037 | 0.963 | -0.962 | 0.195 | -0.962 | 0.195 | Parasitism |
| *Staphylococcus aureus* | *Staphylococcus haemolyticus* | Flight 3 | 2 | 2.739 | 2.519 | 0.117 | 2.756 | 0.041 | 0.959 | -0.957 | 0.094 | -0.957 | 0.000 | Amensalism |
| *Staphylococcus aureus* | *Staphylococcus saprophyticus* | Flight 3 | 2 | 2.739 | 2.363 | 0.061 | 2.972 | 0.020 | 0.980 | -0.978 | 0.258 | -0.978 | 0.258 | Parasitism |
| *Staphylococcus aureus* | *Staphylococcus sp. strain LCT-H4* | Flight 3 | 2 | 2.739 | 2.388 | 0.033 | 3.253 | 0.010 | 0.990 | -0.988 | 0.362 | -0.988 | 0.362 | Parasitism |
| *Staphylococcus aureus* | *Staphylococcus warneri* | Flight 3 | 2 | 2.739 | 2.543 | 0.029 | 2.854 | 0.010 | 0.990 | -0.989 | 0.122 | -0.989 | 0.122 | Parasitism |
| *Staphylococcus epidermidis* | *Staphylococcus haemolyticus* | Flight 3 | 2 | 2.313 | 2.519 | 2.711 | 0.061 | 0.978 | 0.022 | 0.172 | -0.976 | 0.172 | -0.976 | Parasitism |
| *Staphylococcus epidermidis* | *Staphylococcus saprophyticus* | Flight 3 | 2 | 2.313 | 2.363 | 2.566 | 0.086 | 0.968 | 0.032 | 0.109 | -0.964 | 0.109 | -0.964 | Parasitism |
| *Staphylococcus epidermidis* | *Staphylococcus sp. strain LCT-H4* | Flight 3 | 2 | 2.313 | 2.388 | 3.071 | 0.100 | 0.968 | 0.032 | 0.328 | -0.958 | 0.328 | -0.958 | Parasitism |
| *Staphylococcus epidermidis* | *Staphylococcus warneri* | Flight 3 | 2 | 2.313 | 2.543 | 2.689 | 0.079 | 0.971 | 0.029 | 0.163 | -0.969 | 0.163 | -0.969 | Parasitism |
| *Staphylococcus haemolyticus* | *Staphylococcus saprophyticus* | Flight 3 | 2 | 2.519 | 2.363 | 0.119 | 2.878 | 0.040 | 0.960 | -0.953 | 0.218 | -0.953 | 0.218 | Parasitism |
| *Staphylococcus haemolyticus* | *Staphylococcus sp. strain LCT-H4* | Flight 3 | 2 | 2.519 | 2.388 | 0.033 | 3.253 | 0.010 | 0.990 | -0.987 | 0.362 | -0.987 | 0.362 | Parasitism |
| *Staphylococcus haemolyticus* | *Staphylococcus warneri* | Flight 3 | 2 | 2.519 | 2.543 | 0.028 | 2.797 | 0.010 | 0.990 | -0.989 | 0.100 | -0.989 | 0.100 | Parasitism |
| *Staphylococcus saprophyticus* | *Staphylococcus sp. strain LCT-H4* | Flight 3 | 2 | 2.363 | 2.388 | 0.025 | 2.506 | 0.010 | 0.990 | -0.989 | 0.049 | -0.989 | 0.000 | Amensalism |
| *Staphylococcus saprophyticus* | *Staphylococcus warneri* | Flight 3 | 2 | 2.363 | 2.543 | 0.089 | 2.735 | 0.032 | 0.968 | -0.962 | 0.076 | -0.962 | 0.000 | Amensalism |
| *Staphylococcus sp. strain LCT-H4* | *Staphylococcus warneri* | Flight 3 | 2 | 2.388 | 2.543 | 1.557 | 1.557 | 0.500 | 0.500 | -0.348 | -0.388 | -0.348 | -0.388 | Competitive |
